# Modular, synthetic chromosomes as new tools for large scale engineering of metabolism

**DOI:** 10.1101/2021.10.04.462994

**Authors:** Eline D. Postma, Else-Jasmijn Hassing, Venda Mangkusaputra, Jordi Geelhoed, Pilar de la Torre, Marcel van den Broek, Christiaan Mooiman, Martin Pabst, Jean-Marc Daran, Pascale Daran-Lapujade

## Abstract

The construction of powerful cell factories requires intensive genetic engineering for the addition of new functionalities and the remodeling of native pathways and processes. The present study demonstrates the feasibility of extensive genome reprogramming using modular, specialized *de novo*-assembled neochromosomes in yeast. The *in vivo* assembly of linear and circular neochromosomes, carrying 20 native and 21 heterologous genes, enabled the first *de novo* production in a microbial cell factory of anthocyanins, plant compounds with a broad range pharmacological properties. Turned into exclusive expression platforms for heterologous and essential metabolic routes, the neochromosomes mimic native chromosomes regarding mitotic and genetic stability, copy number, harmlessness for the host and editability by CRISPR/Cas9. This study paves the way for future microbial cell factories with modular genomes in which core metabolic networks, localized on satellite, specialized neochromosomes can be swapped for alternative configurations and serve as landing pads for the addition of functionalities.

## Introduction

While microbial cell factories have a great potential for the sustainable production of fuels, chemicals and therapeutics, microbial processes are often less economically attractive than chemical, oil-based processes ^1, 2^. Increasing the cost efficiency of microbial cell factories requires high product titer, rate and yield, features only attainable by intensive genetic engineering of the microbial host via costly strain construction programs. These programs focus on the transplantation of new functionalities, as well as on the reprogramming of the microbial host metabolic networks in which the new functionalities are plugged. These networks of biochemical reactions are encoded by several hundreds of genes scattered over large mosaic microbial genomes. Even considering the CRISPR/Cas9 revolution, rewiring these biochemical networks remains a daunting challenge, in particular for eukaryotic cell factories characterized by a high degree of genetic redundancy ^3^. Microbial platforms in which core biochemical networks can be remodeled at will have a great role to play in the construction of powerful cell factories, but also to reach a deep fundamental understanding of these biochemical networks and their regulation.

The pathway swapping concept was developed to address this persistent challenge ^4^. This modular concept is based on the genetic reduction and clustering of genes belonging to a metabolic pathway, so the pathway can be easily swapped by any other design. This concept was demonstrated in the model, and industrial yeast *Saccharomyces cerevisiae*, using glycolysis and alcoholic fermentation, a 12-step pathway catalyzed by a set of 26 enzymes, as an example. In the SwYG (Switchable Yeast Glycolysis) strain, this set of 26 genes was reduced to 13 and relocalized to a single chromosomal locus, enabling the facile remodeling of the entire pathway ^4^. Scaling up pathway swapping to the set of core biochemical reactions required for most biotechnological applications (i.e. central carbon metabolism) would offer an unprecedented ability to deeply reprogram cell factories metabolism. However, it would require the integration of hundreds of genes on existing chromosomes, which is a potential source of genetic instability. This problem can be addressed by the implementation of supernumerary, *de novo* assembled synthetic chromosomes (named NeoChrs) as orthogonal expression platforms for genome remodeling ^5^.

The present study explores the potential of combining pathway swapping with NeoChrs to simultaneously equip *S. cerevisiae* with new heterologous routes and remodel native metabolic networks. *De novo* production from glucose of the anthocyanin pelargonidin 3-O-glucoside (P3G), a food and industrial dye, was attempted in *S. cerevisiae* as proof of principle ^6^. Although already demonstrated in *S. cerevisiae*, the synthesis of anthocyanins is highly inefficient in all microbial platforms tested hitherto ^6, 7, 8^. Requiring the channeling of carbon through 27 core metabolic catalytic steps in the glycolytic, pentose phosphate, shikimate and aromatic amino acids biosynthesis pathway, as well as 10 reactions native to plants (Fig. 1 and Supplementary Fig. 1), P3G is a perfect paradigm to demonstrate the potential of NeoChrs for metabolic engineering. Modular NeoChrs in linear and circular form harboring yeast native, bacterial and plant genes required for P3G *de novo* synthesis were designed and constructed (Fig. 1 and Supplementary Fig. 1). The resulting strains were tested by in-depth genetic and physiological characterization, and compared to strains carrying test NeoChrs of equivalent size and mostly composed of non-coding DNA. Ultimately, the ability of NeoChrs to serve as landing pad for large pathways and to be edited by CRISPR/Cas9 for metabolic engineering purposes could be demonstrated.

**Figure 1.**
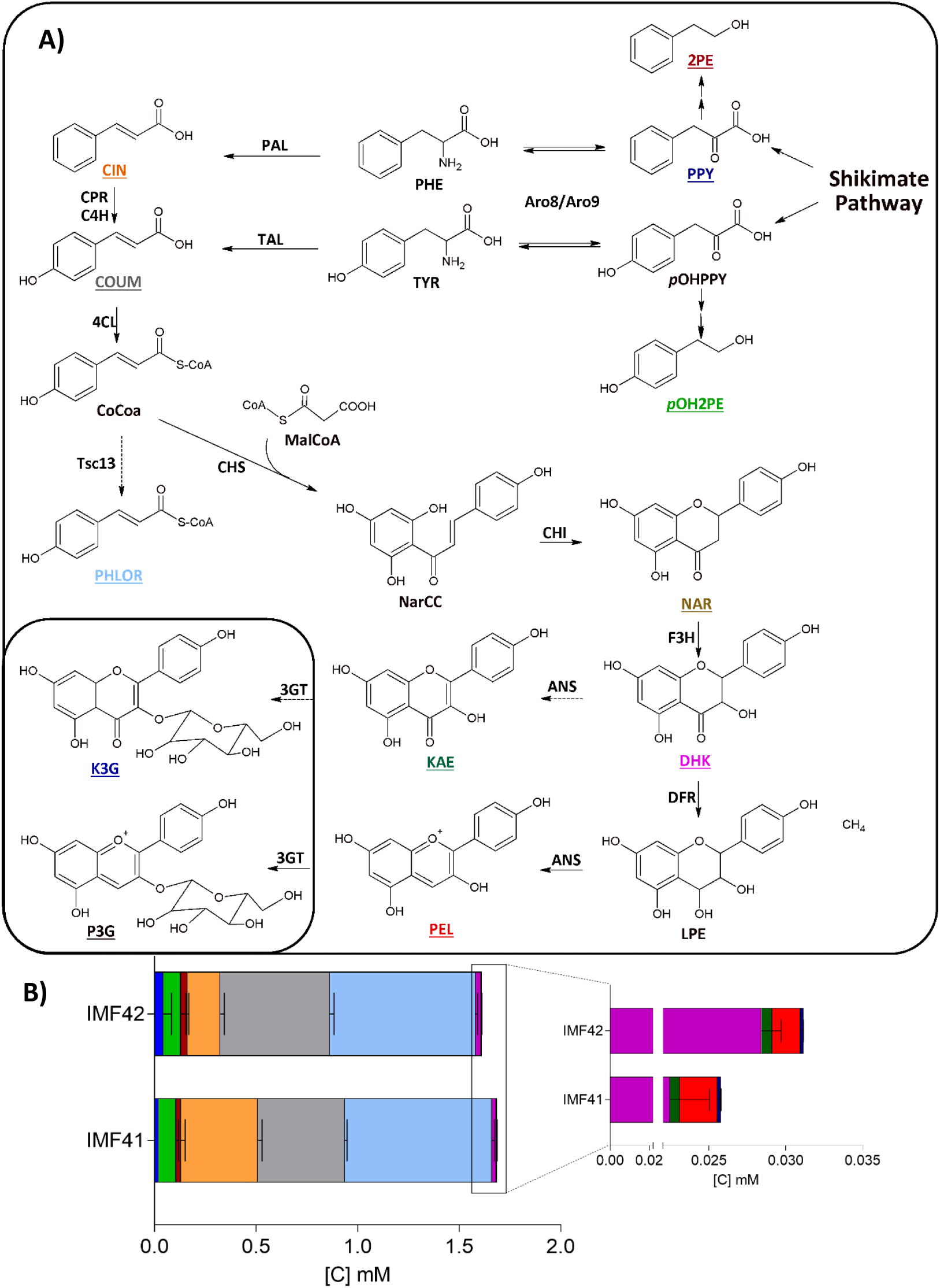
de novo anthocyanin production using NeoChrs. **A)** Schematic overview of the anthocyanin production pathway. Compounds that can be measured by HPLC or GC are underlined, coumaroyl-CoA, narigenin-chalcone, and leucopelargonidin were not measured. The end products of the pathway are indicated in a frame. **B)** Extracellular production of aromatic intermediates of the anthocyanin pathway by engineered *S. cerevisiae* strains IMF41 (Circular NeoChr) and IMF42 (Linear NeoChr). The yeast strains were grown in shake-flask cultures with chemically defined minimal medium with 20 g L^-1^ glucose as carbon source and urea as nitrogen source for 72 hours. The colour of the bars corresponds to the colour of the metabolites in panel A. Phenylpyruvate (blue), *p*-hydroxyphenylethanol (green), 2-phenylethanol (dark red), cinnamic acid (orange), coumaric acid (grey), phloretic acid (sky blue), dihydrokaempferol (purple), kaempferol (dark green), pelargonidin (red) and kaempferol 3-O-glucoside (dark blue). Naringenin and pelargonidin 3-O-glucoside were not detected. The right panel shows a magnification of the data of the metabolites downstream naringenin. The data represents the average ± mean deviation of independent biological triplicates.

## Results

### Neochromosome genetic design: does circular or linear configuration matter?

In *S. cerevisiae* the highly efficient homology directed repair machinery can be exploited for the modular assembly of tailored synthetic chromosomes ^5^. The application of these *de novo* assembled NeoChrs as orthogonal expression platform for pathway engineering requires the fulfilment of several core, size-independent, properties: efficiency and fidelity of assembly, stability in terms of sequence fidelity as well as copy number during replication and segregation, and finally absence of toxicity towards the microbial host. These properties were well met in one particular NeoChr design in strain IMF23 constructed by Postma *et al*. ^5^, offering a promising proof of principle. The chromosomes in this study however were circular, and mimicking the linear configuration of yeast native chromosomes might further improve these core properties^9, 10^. To test this hypothesis, a linear, 100 kb NeoChr was assembled *in vivo* from transcription-unit sized DNA parts and compared to the previously published, circular NeoChr.

The design of the linear test NeoChr in the present study was identical to the 100 kb circular NeoChr design in strain IMF23, consisting of 43 DNA fragments of 2.5 kb ^5^. To summarize, 36 of the DNA fragments used were non-coding in yeast and originated from *Escherichia coli*, the remaining 7 fragments encompassed selection markers, fluorescent reporters and the elements required for chromosome replication and segregation (Fig. 2). Direct assembly of linear chromosomes of this scale has never been attempted before, therefore particular attention was given to the DNA parts framing the chromosomes, containing the telomeric regions. The telomerator containing short telomere seed regions (TeSS) flanking an I-SceI recognition sequence (embedded as intron in a functional *URA3* selection marker), was previously shown to lead to stable linear chromosomes upon *in vivo* digestion ^11, 12^. The same TeSS were used for the direct assembly of linear chromosomes. This design, composed of 44 fragments and identical TeSS regions on the right and left arm of the NeoChr was named NeoChr10. To evaluate the risk of unwanted recombination events between the two identical terminal TeSS fragments, which might cause circularization during *in vivo* assembly of all the DNA parts, a second design was tested. In this second design, one of the TeSS was mutagenized to prevent homologous recombination with the original TeSS, leading to NeoChr11. With the exception of the telomerator module (carrying the *URA3* selection marker), the design of the linear chromosomes (NeoChr10 and 11) and circular chromosome (NeoChr12) was identical, which enabled the direct comparison of the core properties between linear and circular chromosomes. The identification of correctly assembled chromosomes was performed by screening for expression of fluorescent markers by fluorescence-activated cell sorting (FACS), for chromosome size by pulsed field electrophoresis (PFE) and for DNA sequence by whole genome, long-read sequencing (Table 1 and Supplementary Fig. 2-3).

**Figure 2.**
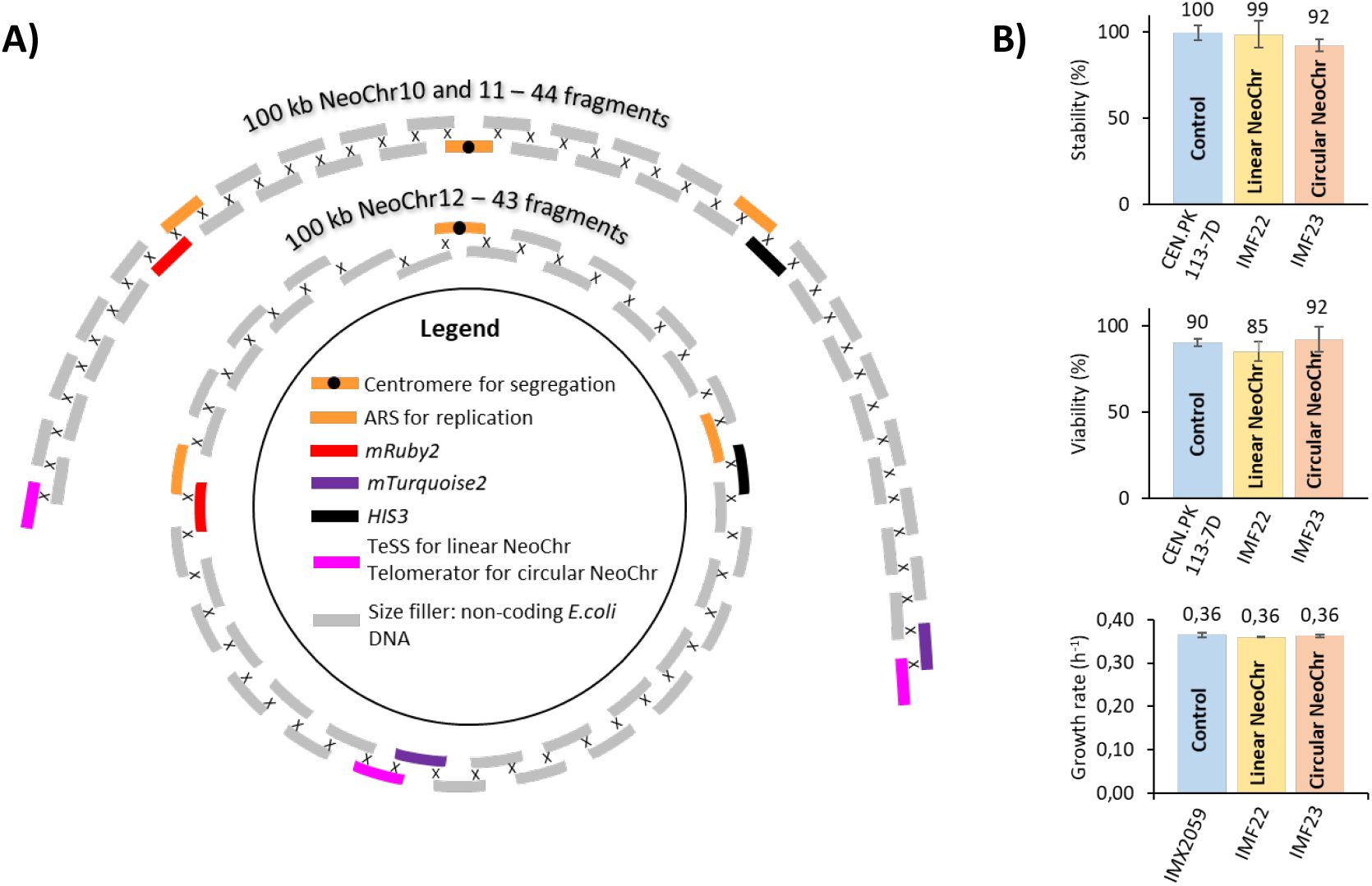
Construction and physiological characterization of test Neochromosomes. **A)** Schematic representation of the *in vivo* assembly design for linear and circular chromosomes. **B**) Physiological characterization of strains with a 100 kb *de novo* assembled linear (IMF22) and circular (IMF23) test NeoChr. **Top graph**, stability of NeoChrs calculated as ratio of the number of colonies on selective medium (with respect to the NeoChr) divided by the number of colonies on non-selective medium. Data represent the average and standard deviation of two days of measurements (day 1 and day 4) of culture duplicates. **Middle graph**, viability of strains, counted as the ratio between the number of colonies growing on non-selective YPD medium and the total number of individually plated cells. Bars represent the average and standard deviation of two days of measurements (day 1 and day 4) of culture duplicates. **Bottom graph**, specific growth rate of strains grown on selective SMD medium. Growth rates represent the average and standard deviation of six biological replicates for IMX2059 and IMF23 and two biological replicates for IMF22. None of the measurements show significant differences between strains (one-way ANOVA with Post-Hoc Tukey-Kramer, *p*<0.05).

**Table 1.**
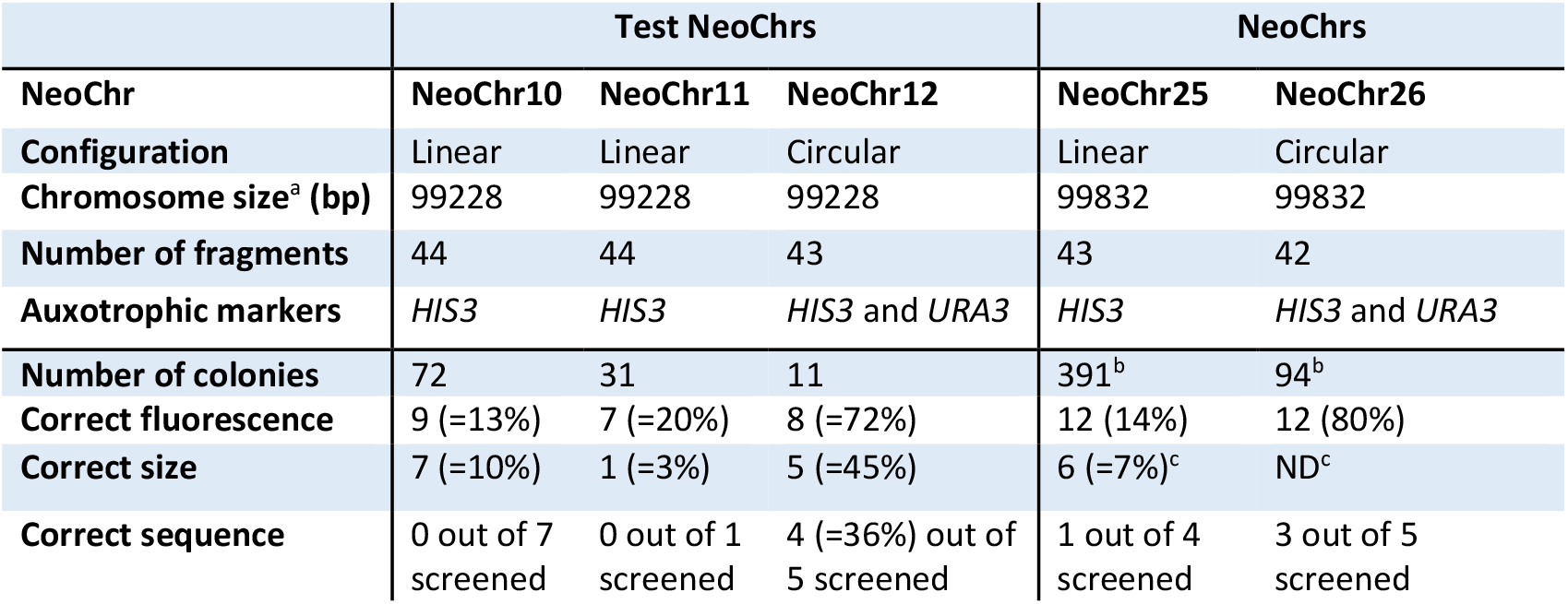
NeoChrs assembly efficiency and fidelity. NeoChrs assembly fidelity tested by the screening pipeline. Correct sequence means that all fragments are present in the correct configuration with respect to the *in silico* design. ^a^ excluding telomeres. ^b^ due to a bacterial infection on plates, the total number of colonies is not reliable. Yeast colonies were subsequently isolated by microscopy based on fluorescence (88 for Neochr25 and 15 for Neochr26). ^c^ For NeoChr25 all 12 colonies were screened by PFE, 6 colonies showed correct size while no band was observed for the other colonies. ND: not determined.

|  | Test NeoChrs |  |  | NeoChrs |  |
| --- | --- | --- | --- | --- | --- |
| NeoChr | NeoChr10 | NeoChr11 | NeoChr12 | NeoChr25 | NeoChr26 |
| Configuration | Linear | Linear | Circular | Linear | Circular |
| Chromosome size<sup>a</sup> (bp) | 99228 | 99228 | 99228 | 99832 | 99832 |
| Number of fragments | 44 | 44 | 43 | 43 | 42 |
| Auxotrophic markers | *HIS3* | *HIS3* | *HIS3* and *URA3* | *HIS3* | *HIS3* and *URA3* |
| Number of colonies | 72 | 31 | 11 | 391<sup>b</sup> | 94<sup>b</sup> |
| Correct fluorescence | 9 (=13%) | 7 (=20%) | 8 (=72%) | 12 (14%) | 12 (80%) |
| Correct size | 7 (=10%) | 1 (=3%) | 5 (=45%) | 6 (=7%)<sup>c</sup> | ND<sup>c</sup> |
| Correct sequence | 0 out of 7 screened | 0 out of 1 screened | 4 (=36%) out of 5 screened | 1 out of 4 screened | 3 out of 5 screened |

As reported by Postma *et al*. ^5^, 36% of the transformants with circular chromosomes were true to the *in silico* design. Conversely, irrespective of the telomere sequence used, no linear NeoChrs faithful to the original design were found. Seven transformants of the linear NeoChr10 and one transformant of the NeoChr11, with correct size according to PFE analysis (Table 1) were sequenced by Nanopore technology. Presence of intact telomeres for all NeoChrs and absence of circularization for both NeoChr10 and 11, demonstrated that the short TeSS supplied were sufficient for the formation of functional telomeres and that homology between the two telomeric fragments was not a hurdle for the direct assembly of linear chromosomes. A wide variety of configurations was observed for these assembled linear NeoChrs (Supplementary Fig. 4), ranging from the absence of a single fragment (NeoChr10.13 and 10.47), a combination of missing and duplicated fragments (e.g. NeoChr10.62) and more complex configurations with missing, duplicated or inverted fragments and swapped regions (NeoChr10.54 and 11.19). These misassemblies revealed the difficulty encountered by yeast cells to assemble all supplied DNA parts, and demonstrated the intervention of non-homologous end joining, while homology directed repair is typically the preferred mode of DNA double strand break repair in *S. cerevisiae* ^13, 14, 15^. The large impact of linear versus circular configuration, despite otherwise identical design, on *in vivo* chromosome assembly possibly revealed differences in the accessibility of telomerator and TeSS fragments in the nucleus for repair and assembly with the other fragments. Nevertheless, two NeoChrs, NeoChr10.13 and 10.47, displayed a remarkably high degree of fidelity with the *in silico* design, with a single fragment (7A) missing while the remaining 43 fragments were correctly assembled. Overall, keeping in mind that circular chromosomes carried an additional auxotrophic marker as compared to linear chromosomes, these results demonstrated that circular chromosomes are superior in terms of assembly efficiency and fidelity as compared to linear chromosomes.

The physiology of strains IMF22, carrying the linear NeoChr10.13, and IMF23, harboring a correct circular chromosome NeoChr12, was compared (Fig. 2B). Both strains grew as fast as the control strain on minimal, chemically defined medium and displayed the same viability on complex medium. During propagation of the strains over four days (ca. 25 generations), the fraction of the population containing both linear and circular chromosomes (measured as the ratio of colonies on selective over non-selective medium) remained unaltered and similar to the control strain, demonstrating the stability of the NeoChrs during cell division. Moreover, fluorescence analysis and sequencing showed that both NeoChr-configurations were present in a single copy per cell (Supplementary Fig. 5-6). To conclude, both linear and circular *de novo* assembled 100 kb NeoChrs were present in one copy per cell, stable and innocuous to their host. As the linear or circular nature of the test NeoChrs did not visibly affect the phenotype of the yeast strains, both configurations were further tested as platforms for metabolic engineering.

### In vivo, de novo *modular assembly of specialized NeoChrs for anthocyanin synthesis*

Following the construction strategy as described above, linear and circular NeoChrs, which were expressing the genes native to yeast, bacteria and plants required for P3G production, were designed (Fig. 3). The precursors for P3G synthesis natively produced in yeast are L-tyrosine and L-phenylalanine (Fig. 1). For *de novo* P3G synthesis from glucose, carbon flows through glycolysis, the pentose phosphate pathway (PPP), cytosolic acetyl-CoA and malonyl-CoA synthesis, the shikimate pathway and aromatic amino acid biosynthesis (Fig. 1). The genes encoding enzymes in these pathways are scattered over the 16 yeast chromosomes and many have a high degree of genetic redundancy, making the remodeling of these metabolic routes extremely challenging. Expanding on the pathway swapping concept, the strain construction strategy was built on SwYG, a strain harboring a single locus, minimized glycolysis and fermentation pathway (13 genes involved in the conversion of glucose to ethanol ^4^). This strain was further engineered by the genetic reduction of the pentose phosphate pathway (removal of the four minor paralogs *NQM1, GND2, TKL2, SOL4* ^16^) with the goal of relocating the genes encoding glycolysis, ethanol fermentation and the PPP from native chromosomes to the specialized NeoChrs (Fig. 3-4 and Supplementary Fig. 1). The biosynthesis of amino acids is tightly regulated in *S. cerevisiae*, particularly via feedback allosteric control ^17^. Therefore, to improve the supply of tyrosine and phenylalanine for P3G synthesis, the entire pathways for aromatic amino acids synthesis from *E. coli* (10 genes) including key feedback resistant alleles (*coEcaroG*^*fbr*^ ^18^, *coEtyrA*^*fbr*^ ^19^, *coEpheA*^*fbr*^ ^20^) were integrated in the design of the specialized NeoChrs. Finally, as aromatic fusel alcohols and acids are undesired by-products during flavonoid production ^21^, the 2-oxo acid decarboxylases responsible for their production (Pdc5, Pdc6 and Aro10) had to be removed ^22, 23^. In the SwYG strain, *PDC5* and *PDC6*, homologues of the major pyruvate decarboxylase encoded by *PDC1*, were already deleted. *ARO10*, that encodes a 2-oxo acid decarboxalyse with broad substrate specificity, was therefore deleted as well from the SwYG strain (Fig. 4A).

**Figure 3.**
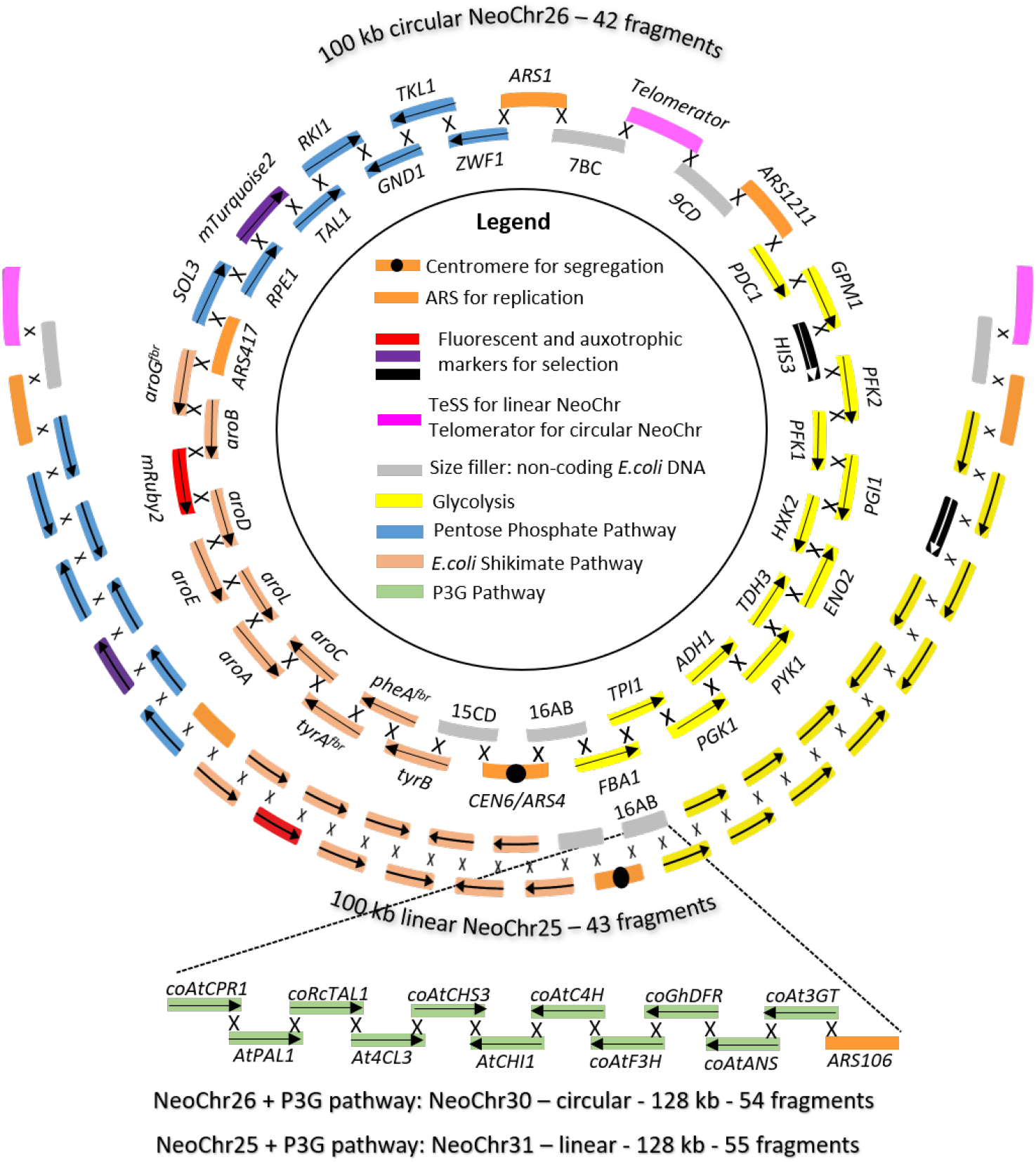
Genetic design of NeoChr for anthocyanin production. *In silico* design of NeoChr25, NeoChr26, NeoChr30 and NeoChr31. The arrows indicate the directionality of transcription. The names of the genes and auxiliary parts are indicated on the circular chromosome. All genes from *E. coli* were codon-optimized. The linear NeoChr differs from the circular NeoChr by the presence of two TeSS ends instead of the telomerator (carrying the *URA3* selection marker, pink parts). The eleven plant genes required for P3G biosynthesis were integrated into chunk 16AB from NeoChr25 and NeoChr26, resulting in linear NeoChr30 and circular NeoChr31 respectively.

**Figure 4.**
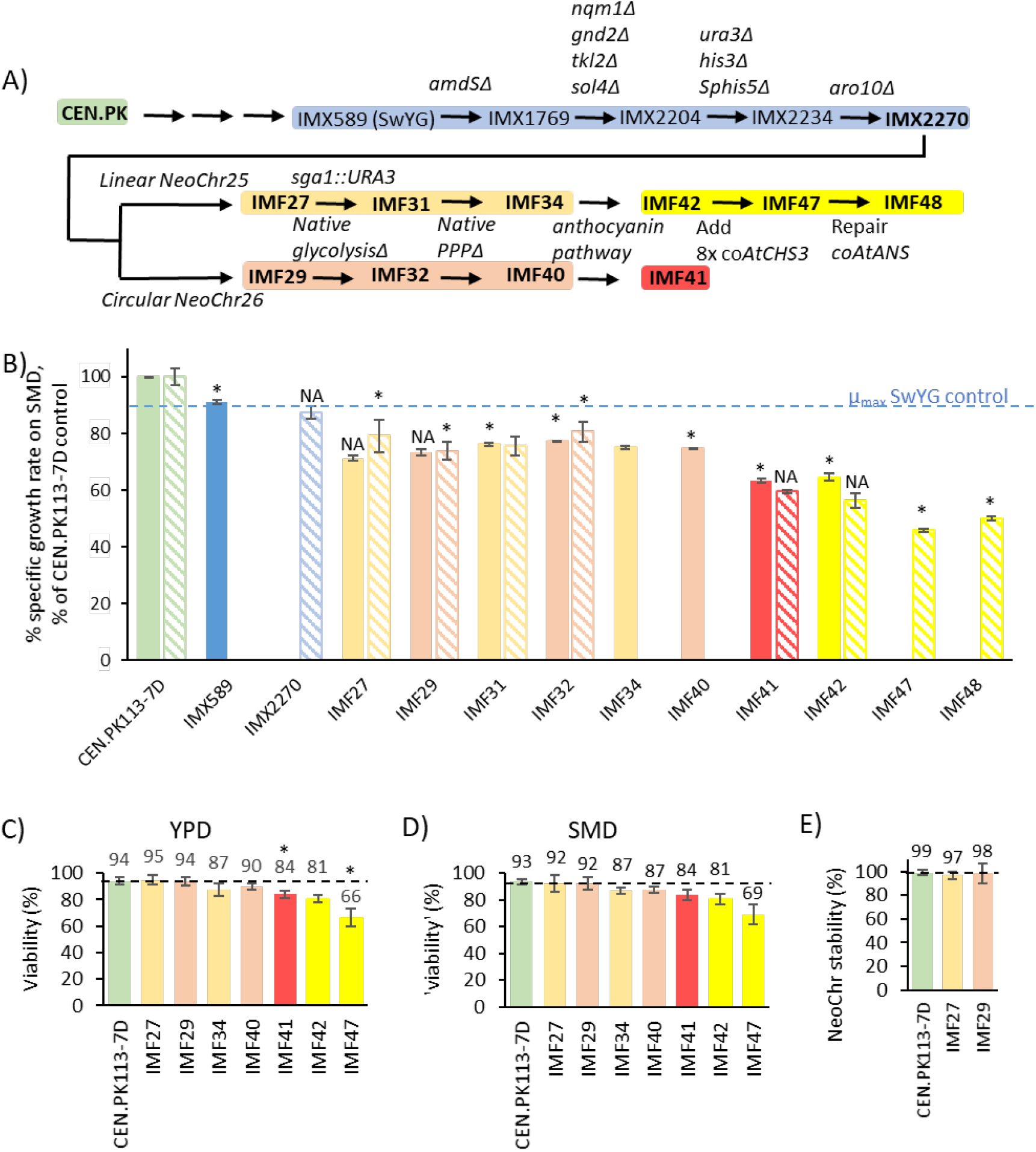
Construction and Physiological characterization of strains with NeoChrs engineered for anthocyanin production. **A**, overview of the strains constructed in this study for pathway engineering. **B**, Specific growth rate on SMD medium of engineered strains expressed as % of the control CEN.PK113-7D. The same colour-coding has been used as for panel A. Solid-filled bars: average and standard deviations of biological duplicates grown in shake flasks. Pattern-filled bars: average and standard deviations of three to six biological replicates grown in plate reader. For strain IMF27, uracil was supplemented to the medium. For IMX2270 uracil and histidine were supplemented to the medium. **C** and **D**, viability calculated as the number of colonies on YPD (C) or SMD (D) divided by the total number of plated cells (384 FACS-sorted cells). **E**, stability of the NeoChrs measured as number of transformants on selective medium (SMD (supplemented with Uracil)) divided by the number of colonies on non-selective medium (YPD). For panels C, D and E, bars represent the average and standard deviation of biological duplicates measured on two days (day 1 and day 4). Significant differences with respect to the first ancestor in the same graph are indicated with an asterisk (two-tailed paired homoscedastic t-test P< 0.05). In Panel B NA indicates data for which statistics could not be calculated because the ancestor was grown in a different set-up than the tested strain (shake-flask or plate reader).

To summarize the NeoChr design for P3G synthesis, the specialized synthetic chromosomes were intended as exclusive expression platforms for the glycolytic, fermentative, pentose phosphate pathways (20 genes), the bacterial shikimate and amino acid biosynthesis pathway (10 genes), and the 11 plant genes involved in the synthesis of P3G from tyrosine and phenylalanine.

The genetic design of the specialized NeoChrs was particularly challenging. This study presents the very first attempt of pathway construction of this magnitude, and information to rationally design an optimal configuration for NeoChr assembly with high efficiency, stability and expression is scarce. The first design consideration was high fidelity assembly and maintenance of the Neochrs. While the present study embraces the remarkable ability of *S. cerevisiae* to recombine homologous sequences, homology directed repair (HDR) might cause unwanted recombinations within the NeoChr during assembly and propagation. To prevent unwanted recombinations, homology within the NeoChr was kept to a minimum. However, maintaining a low homology for promoter and terminators regions for 41 transcription units was challenging. The 20 yeast genes were framed by their native promoter and terminator, however the 21 bacterial and plant genes also required terminators and strong constitutive promoters with unique sequences. Such promoters are not abundant in the *S. cerevisiae* molecular toolbox, and most often of glycolytic origin. The heterologous genes were therefore expressed from a selection of previously characterized *S. cerevisiae* promoters ^24, 25^ which also showed high and condition-independent expression in *S. cerevisiae* from a large transcriptome compendium ^26^, and promoters from *Saccharomyces eubayanus* and *Saccharomyces kudriavzevii* that have little sequence similarity but share functionality with their *S. cerevisiae* relatives ^27^ (Supplementary Table 1). The promoter regions of the transcription units on the NeoChr design did not display homologous sequences longer than 43 bp (*pScADH1*/*pSkADH1* and *pScTDH3*/*pSkTDH3*). Terminators, that generally have a milder impact on gene expression than promoters, especially when paired with strong promoters ^28, 29, 30^, were selected from a set of in-house central carbon metabolism terminators (Supplementary Table 1).

The second consideration for the genetic design of the NeoChrs (Fig. 3) was the spatial organization of transcriptional units and other elements along the chromosomes, which might impact both maintenance and gene expression of NeoChrs. To facilitate future modular pathway remodeling, genes encoding glycolysis, fermentation, PPP, shikimate and aromatic amino acids pathways were clustered per pathway. To prevent gene silencing resulting from chromatin structure near centromere and telomeres, transcriptional units were separated from these elements by 5 kb non-coding DNA fragments originating from *E. coli*. To facilitate the screening of correctly assembled chromosomes, fluorescent reporters (*mRuby2, mTurquoise2*) and auxotrophic markers (*HIS3* and *URA3* for circular and *HIS3* only for linear NeoChrs) were evenly spaced along the NeoChrs. Finally, while highly expressed genes are scattered across native chromosomes ^31^, the designed NeoChrs are transcriptional hotspots with high density of heavily transcribed genes. To prevent potential clashes between the replication and transcription machineries, the directionality of transcription units was chosen to follow the same orientation as replication ^11^.

To test the suitability of NeoChrs for both *de novo* assembly of pathways and as landing pads for pathways a two-steps construction approach was followed. In a first step, 100 kb circular and linear chromosomes carrying all yeast and bacterial genes were assembled *de novo* in yeast (NeoChr26 and 25, respectively) (Fig. 4A). In a second step, the genes for anthocyanin synthesis from plant and from *Rhodobacter capsulatus* (*coRcTAL1* only) were simultaneously *de novo* assembled and integrated as a module in the NeoChrs (Fig. 3).

*De novo* assembly of fully coding NeoChrs verified by fluorescence, PFE and sequencing, confirmed the higher assembly efficiency of circular over linear NeoChrs observed with the test NeoChrs (Table 1, Supplementary Fig. 7-8 and Supplementary Table 2-3). Remarkably, one linear NeoChr was faithful to the *in silico* design. Genome re-sequencing of a transformant with this NeoChr (NeoChr25, strain IMF27) and sequencing of a transformant with a correctly assembled circular NeoChr (NeoChr26, strain IMF29), identified few mutations, with a single non-synonymous mutation in a coding region of NeoChr26 (*RKI1*, Supplementary Table 2). Summarizing, the assembly of the linear and circular NeoChrs was successful and few mutations in the native and synthetic chromosomes were caused by the addition of the NeoChrs (Supplementary Table 2-3). Next, exclusive expression of glycolysis, fermentation and PPP from the specialized NeoChrs was achieved by Cas9-mediated excision of the single locus glycolysis from ChrIX and multiplex deletion of the seven genes of the minimized PPP (Fig. 4A). The targeted deletion of the seven PPP genes scattered over the native chromosomes was facilitated by the insertion of watermarks on their near-identical copy carried by the NeoChrs ^32^.

As previously observed, genetic relocalization of the glycolytic and fermentative pathways in SwYG caused a mild (IMX589, 10%) reduction in growth rate as compared to the control strain CEN.PK113-7D (Fig. 4B, ^4^). Additional genetic reduction of the PPP and deletion of *ARO10* (strain IMX2270) had no visible additional phenotypic effect (Fig. 4B). Remarkably, NeoChrs with 30 yeast and bacterial genes were as stable during cell division as native chromosomes in control strain CEN.PK113-7D, and strains IMF27 and IMF29 were as viable as the control strain. Their growth rate was slightly decreased (10-16%) with respect to their parental strain IMX2270 (Fig. 4B). The linear or circular topology affected neither NeoChr stability and viability nor specific growth rate. Finally, exclusive expression of the yeast glycolytic, fermentative and pentose phosphate pathway from the NeoChrs (strains IMF34 (linear) and IMF40 (circular) did not further decrease viability nor the specific growth rate, which remained high at ca. 0.30 h^-1^ (Fig. 4B and C)

### Expression of the anthocyanin biosynthetic pathway from a NeoChr

Inspired by earlier work ^7^, synthesis of P3G was attempted by integration of 11 anthocyanin genes from *Arabidopsis thaliana* (*AtPAL1, coAtCPR1, coAtC4H, At4CL3, coAtCHS3, AtCHI1, coAtF3H, coAt3GT, coAtANS), Gerbera hybrida* (*coGhDFR*) and *Rhodobacter capsulatus (coRcTAL1)*. As basal design, single copies of these 11 genes, interspaced by an ARS sequence, were integrated in the 16AB *E. coli* DNA chunk on the circular and linear NeoChrs (Fig. 3 and 4A), leading to NeoChr30 (IMF41) and NeoChr31 (IMF42), respectively. The two strains grew with a similar specific growth rate, but as compared to their parental strain devoid of anthocyanin pathway, they displayed a 15-17% slower specific growth rate (IMF34 and IMF40, *p*-values of 0.01 and 0.00 respectively, Fig. 4B), in agreement with previous observations ^7^. The viability of IMF41 was similar to that of IMF42, but IMF41 was slightly less viable than its ancestor IMF40 (7% lower, *p*-values of 0.03, Fig. 4C and D). Flow cytometric DNA quantification, short-and long-read sequencing showed that both linear and circular chromosomes were present in a single copy in IMF41 and IMF42 (Supplementary Fig. 5 and 6).

In both IMF41 and IMF42, insertion of the plant genes on the NeoChr caused the duplication and inversion of the last four genes in the anthocyanin pathway, *coAtF3H, coGhDFR, coAtANS* and *coAt3GT*. Sequence analysis revealed that this unexpected event resulted from homologous recombination between the *ScFBA1* promoter (upstream *ScFBA1*) and the *SeFBA1* promoter (upstream *coAtC4H*), although slightly different regions of the promoters recombined in IMF41 and IMF42 (Supplementary Fig. 9). The two promoters share 58% homology with identical stretches of 24 nucleotides. The absence of homology in the region where the second recombination event occurred suggested the non-homologous recombination of the two *pSePYK1*-SHR CJ ends (Supplementary Fig. 9). The presence of the same chromosomal rearrangement in both the linear and circular chromosome, involving rare micro-homology (7-24 bp of 100% homology) and non-homologous recombination events, and the absence of *ARS106*, suggested that the *ARS106* fragment was supplied in suboptimal concentration during transformation. As the same duplication occurred in the strains with linear and circular NeoChr, the impact of NeoChr shape can still be investigated. Plant proteins are typically difficult to express in a yeast environment, a positive rather than negative effect of this duplication on anthocyanin production was therefore anticipated. If desired the duplicated genes can be removed using CRISPR/Cas9 editing.

Short-and long-read sequencing revealed the occurrence of a 46 bp insertion at the beginning of the *coAtANS* gene. The presence of this mutation in the *coAtANS* expression cassette showed that it occurred prior to assembly in the NeoChr, and was not caused by *in vivo* assembly. This insertion led to an early stop codon, however the presence of a start codon further along the protein still enabled the synthesis of a 277 amino acids long peptide (Supplementary Fig. 10). This 80 amino acids shortening from the N-terminus most likely strongly affected the anthocyanidin synthase functionality in IMF41 and IMF42. With the exception of a single non-synonymous mutation in the gene encoding the 4-coumarate:CoA ligase (*At4CL3*, Thr-15-Ala) in IMF42, plant genes were exempt of mutations.

Grown in aerobic shake flask cultures in chemically defined medium, IMF41 and IMF42 produced detectable levels of anthocyanins. The total amount of aromatics produced by the two strains was similar (1.69±0.08 and 1.61±0.06 mM) and anthocyanins down to dihydrokaempferol (DHK) could be reliably detected and quantified (Fig. 1, Supplementary Table 4). The production of DHK revealed the functional expression of at least eight out of the eleven NeoChr-borne plant genes in *S. cerevisiae*. Considering that metabolites downstream *At*4Cl3 such as phloretic acid and dihydrokaempferol were detected in both IMF41 and IMF42, the mutation of *At4CL3* had no detectable effect (Fig. 1).

Over 98% of the produced aromatic products were intermediates (cinnamic and coumaric acid) or byproducts upstream chalcone synthase (phloretic acid)(Fig. 1), a result in line with previous reports of chalcone synthase being a bottleneck for anthocyanin production, and a typical target for gene-dosage engineering ^7, 21^. While the synthesis of the end products P3G and K3G is challenging, detectable levels of other metabolites downstream dihydrokaempferol were expected (kaempferol, pelargonidin and leucopelargonidin) ^7^. The absent or barely detectable levels of these metabolites, both in culture supernatants and cell extracts (Fig. 1 and Supplementary Table 4) confirmed the poor functionality of the truncated anthocyanidin synthase *At*Ans. Overall, these results showed that both, the linear and circular structure of the NeoChrs did not affect the physiology of the engineered strains or anthocyanin production, and that the two-step assembly of a NeoChr expressing the *de novo* anthocyanin synthesis pathway enabled anthocyanin production.

### Neochromosome engineering for improved de novo anthocyanin biosynthesis

The two engineering targets identified to improve anthocyanin synthesis were implemented by CRISPR/Cas9 editing of the NeoChrs. Targeting the linear NeoChr31 (strain IMF42), four additional *coAtCHS3* copies were integrated in four different loci on the NeoChr spaced by essential genes (in Chunk 7BC, chunk 15CD, SHR N and chunk 9CD, Fig 3). In an attempt to further boost anthocyanin synthesis, four additional *coAtCHS3* copies were integrated in native chromosomes (at *CAN1, X2, YPRCtau3* and *SPR3* loci^33, 34, 35, 36^), resulting in a total of nine *coAtCHS3* copies (IMF47). Finally, the *coAtANS* gene was repaired leading to strain IMF48. The structure of the NeoChr was not affected by these additional genetic interventions.

The performance of IMF48, IMF42 and IMF41 was compared in pH-controlled, aerated bioreactors, a culture tool previously shown to enable higher anthocyanin titers than shake flasks ^7^. The strains were grown in minimal, chemically defined medium without additional growth supplements (*e*.*g*. yeast extract or peptone), to purely evaluate *de novo* production of anthocyanins from glucose. The physiology of the three strains was similar, despite the eight additional *coAtCHS3* copies carried by IMF48 (Supplementary Table 5). Irrespective of the strain, flavonoids were produced both during exponential growth on glucose and after the diauxic shift (Supplementary Fig. 11). Confirming the results from shake flask cultures, the structure of the NeoChr in IMF41 and IMF42 affected neither yeast physiology nor flavonoid formation (Table 2, Supplementary Fig. 11 and Supplementary Table 5). Overall, strain IMF48 outperformed IMF41 and IMF42 carrying a single *coAtCHS3* copy and a truncated *coAtANS* (Table 2, Supplementary Fig. 11 and Supplementary Table 5). IMF48 produced 14-fold more flavonoids downstream chalcone synthase than IMF41 and IMF42, and the production of pelargonidin, kaempferol and K3G, metabolites downstream *At*Ans, was increased by ca. 30-, 350- and 500-fold, respectively in IMF48 as compared to IMF41 and IMF42 (Table 2). P3G, not detected intracellularly nor extracellularly in IMF41 and IMF42 cultures, was produced with a titer of 0.049 ± 0.007 μM by IMF48 at the end of the glucose consumption phase (Supplementary Fig. 11C). Notably, this is the first report of P3G extracellular production by yeast cultures. The concentration of P3G decreased by ca. three-fold at the end of the diauxic phase, presumably due to degradation by periplasmic β-glucosidases (e.g. - Exg1), upon glucose exhaustion (Supplementary Fig. 11C). These enzymes are interesting targets for further improvement of P3G production. IMF48 accumulated more coumaric acid and *p-*hydroxyphenylethanol, aromatic intermediates upstream *At*Chs3, an unexpected response to the increased chalcone synthase expression that suggested some yet unidentified regulation of the plant enzymes in the yeast environment (Table 2). No extracellular naringenin was detected for any of the tested strains. The absence of naringenin but increased intra- and extracellular dihydroxykaempferol concentration with increasing *coAtCHS3* copy number and flux through the anthocyanin pathway, showed that flavonoid hydroxylase (F3H) was most probably not limiting P3G production and was therefore not a target for genetic engineering at that stage. This apparent overcapacity of flavonoid hydroxylase might be related to its presence in two copies on the NeoChrs. Conversely, the strong increase in flavonoid concentration upstream the last enzyme, 3GT (kaempfarol and pelargonidin, Table 2), suggested that despite the duplication of its gene, this enzyme might be limiting for K3G and P3G production and a target for further improvement. The titers of produced flavonoids were modest. However, the fraction of flavonoids over all aromatic compounds produced by IMF48 was 15%, representing a substantial improvement compared to the 2% produced by IMF41 and IMF42. Additionally, the total production of aromatic compounds was increased by 2.3-fold in IMF48.

**Table 2.** Characterization of anthocyanin production in bioreactors. Determination of the intermediates of the anthocyanin pathway in *S. cerevisiae* strains IMF41, IMF42 and IMF48. Extracellular and intracellular metabolite concentration at the end of aerobic and pH controlled (pH 5.0) bioreactor batch cultures on glucose. The concentrations (mM) of metabolites upstream from, and including, naringenin (included) were measured by HPLC while metabolites downstream were quantified by LC-MS/MS. The data represents the average ± mean deviation of independent biological duplicates. Intermediates of the anthocyanin pathway coumaroyl-CoA, naringenin chalcone, and leucopelargonidin were not measured. An asterisk indicates statistical significance when comparing extracellular concentrations of IMF48 to concentrations of IMF41 and IMF42 (Student *t*-test, *p-*value threshold 0.05, two-tailed, homoscedastic).

|  | IMF41 |  | IMF42 |  | IMF48 |  |
| --- | --- | --- | --- | --- | --- | --- |
| | Extracellular (mM) | Intracellular (μmol/g dry weights) | Extracellular (mM) | Intracellular (μmol/g dry weights) | Extracellular (mM) | Intracellular (μmol/g dry weights) |
| Phenylpyruvate | 6.68E-02 ± 0.00E+00 | NM | 4.47E-02 ± 4.95E-03 | NM | BD | NM |
| 2-phenylethanol | BD | NM | BD | NM | BD | NM |
| *p*-hydroxyphenylethanol | 8.94E-02 ± 2.55E-03 | NM | 7.87E-02 ± 9.90E-04 | NM | 1.40E-01 ± 2.83E-03* | NM |
| Cinnamic acid | BD | NM | BD | NM | BD | NM |
| Coumaric acid | 8.46E-01 ± 2.38E-02 | NM | 7.89E-01 ± 6.22E-03 | NM | 2.05E+00 ± 1.98E-02* | NM |
| Phloretic acid | 5.46E-01 ± 3.24E-02 | NM | 5.92E-01 ± 1.15E-02 | NM | 5.98E-01 ± 2.42E-02 | NM |
| Naringenin | BD | NM | BD | NM | BD | NM |
| Dihydrokaempferol | 3.29E-02 ± 1.64E-04 | 7.26E-02 ± 4.44E-02 | 3.32E-02 ± 2.01E-04 | 1.41E-01 ± 1.93E-02 | 2.90E-01 ± 3.98E-03* | 1.15E+00 ± 2.00E-01 |
| Kaempferol | 1.04E-04 ± 8.51E-06 | 1.20E-03 ± 8.84E-04 | 8.61E-05 ± 1.64E-05 | 1.58E-03 ± 6.99E-04 | 3.55E-02 ± 5.63E-03* | 3.64E+00 ± 1.18E+00 |
| Pelargonidin | 1.46E-03 ± 1.99E-05 | 6.25E-03 ± 3.96E-03 | 1.64E-03 ± 2.73E-05 | 1.00E-02 ± 3.26E-03 | 4.14E-02 ± 5.63E-04* | 4.96E-01 ± 7.76E-02 |
| Kaempferol 3-O-glucoside | 2.74E-05 ± 5.38E-07 | 3.42E-05 ± 2.88E-05 | 2.17E-05 ± 6.21E-07 | 1.18E-04 ± 8.03E-05 | 9.77E-03 ± 1.09E-04* | 1.44E-02 ± 1.70E-03 |
| Pelargonidin 3-O-glucoside | BD | BD | BD | BD | 1.45E-05 ± 4.78E-06* | 2.51E-04 ± 1.95E-05 |
| Total aromatics (a) | 1.47 ± 0.04 | - | 1.14 ± 0.03 | - | 3.3 ± 0.23* | - |
| Total flavonoids (b) | 0.03 ± 0.002 | | 0.03 ± 0.002 | | 0.51 ± 0.19* | |
| Fraction of flavonoids (=b/a) | 2% | - | 2% | - | 15%* | - |

The combined co*AtCHS3* copy number increase and *coAtANS* repair markedly affected the flux distribution in the aromatic pathway. To estimate the respective contribution of chalcone and anthocyanidin synthase to these changes, the parental strain of IMF48, a strain carrying nine *coAtCHS3* copies, but with impaired *coAtANS* was tested (IMF47, Fig. 4A). In line with increased chalcone synthase activity, IMF47 produced three times more flavonoids than IMF42, while the production of metabolites upstream chalcone synthase was unchanged. However, P3G was still not detected in IMF47 cultures. The fraction of flavonoids over total aromatics was 6% in IMF47 (Supplementary Table 4). Repair of *coAtANS* in IMF48 further increased flavonoids production by ca. four-fold and slightly, but significantly, increased total aromatics production as compared to IMF47 (1.4-fold, Supplementary Table 4). Altogether these data revealed that both increased *coAtCHS3* copy number and *coAtANS* repair contributed to the improvement of anthocyanin production in IMF48.

## Discussion

The present study illustrates the amazing potential of *S. cerevisiae* for fast and extensive genome remodeling via synthetic chromosome engineering. Supernumerary, specialized chromosomes could be easily and rapidly assembled, in a single transformation round, from 30 transcription units and 13 accessory DNA parts. Beyond this technical ‘tour de force’, the NeoChrs resembled native chromosomes in terms of replication and segregation. The stable maintenance of the NeoChrs at one copy number and their harmlessness to the host are important features for their implementation as metabolic engineering platforms. Plant-derived chemicals have a broad range of biotechnological applications, but their *de novo* microbial production requires remodeling of the host metabolism, as well as functional transplantation of the plant pathway. Equipped with pathway swapping, NeoChrs enabled the facile implementation of three complex interventions: i) the remodeling of native metabolic networks (glycolysis and PPP), ii) the provision of an optimized metabolic route from prokaryotic origin (shikimate pathway) as surrogate for a native route, and iii) the implementation of a new, heterologous pathway from plant origin (anthocyanin synthesis). These modifications readily enabled the synthesis of anthocyanins from glucose, and the first report of extracellular P3G production in yeast or any other single microbial host. The implementation of NeoChrs not only accelerated strain construction, and thereby genome remodeling, but also prevented interferences with native chromosomes. While chromosome construction was carefully designed in two steps to probe the limits of *in vivo* NeoChr assembly, the present data suggest that single-step chromosome assembly would have readily enabled anthocyanin production in a shorter time frame. Once assembled, the NeoChrs could be edited using CRISPR/Cas9. CRISPR/Cas9 editing efficiency is locus-dependent, and a small set of robust genomic integration sites have been validated to date ^33, 34, 37^. Carefully designed, mostly non-coding NeoChrs, harboring strategically located, optimized CRISPR/Cas9 programmed sites, could become ideal landing pads for large sets of (heterologous) genes. Finally, the successful assembly of chromosomes from 43 parts suggests that the limit of *in vivo* assembly has not been reached and even larger chromosomes can be assembled if required.

Synthetic Genomics is a young research field ^38^, and the design principles for optimal, tailor-made NeoChrs are ill-defined. The synthetic chromosomes rebuilt in the Sc2.0 initiative uses native chromosomes as scaffolds and yet reproduces the native organization of the chromosomes, albeit omitting non-essential elements and adding some short sequences ^39, 40^. With the possibility to construct and test any genetic design, *de novo*-assembled NeoChrs are fantastic testbeds to explore the genetic and physiological impact of chromosome sequence and structure. In this study, some design guidelines were formulated and tested, leading to stable chromosomes, easy to screen and with functionally expressed transcription units. The 100-130 kb linear and circular NeoChrs in this study showed equal growth rate and mitotic stability. This is in good agreement with earlier studies showing approximate equal stability of linear and circular chromosomes of 100 kb, while beneath this size circular chromosomes appear more stable and above this size linear chromosomes are more stable ^11, 41, 42, 43, 44^. A remarkable observation was the low assembly efficiency of linear chromosomes as compared to their circular counterparts. As opposed to transcription units, telomeres and centromeres have specific, cell-cycle dependent localizations within the nucleus ^45, 46, 47, 48^. It is conceivable that, for linear chromosomes equipped with one centromere and two telomeres, accessibility and spatial organization of these DNA parts conflict with homologous recombination, a difficulty potentially alleviated for circular chromosomes equipped with a single centromere and no telomeres. Additionally, the frequent occurrence of non-homologous repair in the linear NeoChrs, an otherwise rare mechanism active throughout the entire cell cycle ^13^, might indicate temporal incompatibility between telomere and centromere availability and homologous recombination, mostly active during the S/G_2_ phase ^13^. For biotechnological applications, circular chromosomes, easier to isolate from yeast, are therefore recommended, and can be equipped with a telomerator to enable ulterior linearization if required ^11, 12^.

In their current design, the NeoChrs are extremely information-dense, with short intergenic regions and a concatenation of highly transcribed genes. While this genetic design leads to functional expression of the NeoChr-born genes and is not harmful for the host cells, many fundamental questions regarding optimal genetic design remain to be systematically explored, such as the cell’s tolerance to ‘transcriptional hotspots’ ^4, 5, 49^, the impact of transcription units localization, orientation and distancing on gene expression, or the requirement for multiple selection markers for chromosome stability. Furthermore, while NeoChrs harboring spaced homologous sequences were genetically stable, the presence of homology between DNA parts might lead to unwanted recombination events during *in vivo* assembly of the NeoChrs. In this work the DNA parts were designed as ‘homology-free’ as possible, a design principle difficult to apply considering the limited availability of strong, constitutive promoters. Homology between DNA parts could be kept to a minimum thanks to the implementation of promoters from *S. cerevisiae* relatives ^27^. Nevertheless, the highest homology between promoters was still 75% (with stretches of identical sequences up to 43 nucleotides (*pSkTDH3* and *pTDH3*)). Along this line the *S. cerevisiae* toolbox could be further enriched by mining non-*Saccharomyces* yeasts genetic diversity for functional but sequence divergent promoters ^27, 50^. In a more distant future, progress in the design of synthetic promoters should enable the construction of libraries of *in silico*-designed, artificial promoters with minimal homology ^51, 52, 53^.

From the present study, we can envisage future microbial cell factories with modular genomes in which core metabolic network and processes, localized on satellite, specialized NeoChrs can be swapped for alternative configurations. Following the 32% reduction of the 111 genes of yeast central carbon metabolism ^16, 54^, the present strategy for *in vivo* assembly of NeoChrs can be applied to construct yeast strains carrying specialized NeoChrs as exclusive expression platforms for central carbon metabolism. Combined with pathway swapping ^4^, such strains would enable fast and easy remodeling of large sets of core cellular functions. As yeast is tolerant to chromosome ploidy variation ^55^, other strategically designed NeoChr could carry other industrially-relevant pathways (e.g. nitrogen or fatty acids metabolism) and processes (e.g. protein secretion). Additional NeoChrs, similar to the test NeoChrs from the present study, but tailored for CRISPR/Cas9 targeting, could also serve as landing pads dedicated to the addition of functionalities.

## Materials and Methods

### Strains, growth medium and maintenance

All *S. cerevisiae* strains used and constructed in this study are derived from the CEN.PK family (Supplementary Table 6-7) ^56^. The detailed protocols for *S. cerevisiae* and *E. coli* growth and maintenance are detailed in Supplementary Methods 1.

### Molecular biology techniques

Genomic DNA extraction from *E*.*coli* and *S. cerevisiae*, different PCR techniques and gel-verification of the amplicons, methods for *E. coli* and yeast transformation and various kits used in this study are detailed in Supplementary Methods 2.

### Plasmid construction

All plasmids used in this study are listed in Supplementary Table 8.

**gRNA plasmids**. GuideRNA (gRNA) plasmids (Supplementary Table 8A) for targeting Cas9 to specific loci were constructed as described by Mans *et al*. ^37^. Primers to construct and verify the gRNA plasmid are listed in Supplementary Table 9-10. **Golden Gate part plasmids**. Part plasmids compatible with Golden Gate Assembly, harboring promoter, gene or terminator were constructed using the Yeast Toolkit principle ^24^. A range of part plasmids were present in-house and previously described (Supplementary Table 8B). Some of the promoters, genes and terminators flanked by BsaI and BsmBI restriction sites were ordered from GeneArt (Thermo Fisher Scientific, Waltham, MA). The promoters and terminators listed in Supplementary Table 8C were subcloned by GeneArt (Thermo Fisher Scientific) in the entry vector pUD565 and could be directly used for construction of expression plasmids. The pentose phosphate pathway genes were ordered from GeneArt but subcloned in-house into entry vector pUD565 (Supplementary Table 8D). Finally, some part plasmids were made by amplifying the target region with primers containing part type specific overhangs (Supplementary Table 11) and assembling in entry vector pUD565 by BsmBI cloning. These plasmids are listed in Supplementary Table 8E with the respective primers and template for target region amplification. Internal BsmBI or BsaI sites were removed as described by Hassing *et al*. ^57^ and verified by Sanger sequencing (Baseclear, Leiden, The Netherlands). For two parts (*coAtANS* #1 and *pSePYK1*) no correct *E*.*coli* part plasmid transformant was found. Therefore, the PCR fragments containing yeast toolkit flanks were directly assembled into expression cassettes (Supplementary Table 8G). Plasmids were verified by PCR and/or restriction analysis (Supplementary Table 12). **Golden Gate expression plasmids**. All expression plasmids were made using BsaI mediated Golden Gate assembly ^24, 57^ in the pGGKd012 GFP dropout plasmid. All expression cassettes and the part plasmids (or PCR fragments) used for construction are outlined in Supplementary Table 8F-G. Plasmids were verified by PCR and/or restriction analysis (Supplementary Table 13). Correct assembly of some of the expression plasmids was verified by Sanger sequencing (Baseclear). **Gibson assembly expression plasmids**. For two expression cassettes, namely: *pRPS3-coAtCPR1-tIDH2* and *pSeTPI1-At4CL3-tSDH2*, the individual parts contained too many BsaI/BsmBI sites, making Golden Gate assembly impossible. Therefore, the individual parts were amplified with PCR primers containing homology flanks and the expression cassettes were constructed via Gibson Assembly (Supplementary Table 8H and 11).

### Strain construction

#### Construction of strains harboring test NeoChrs

Test NeoChrs, NeoChr10 and NeoChr11 consisted of 43 fragments namely: 36 non-coding 2.5 kb *E*.*coli* DNA fragments, *CEN6/ARS4, ARS1, ARS417, mRuby2, mTurquoise2, HIS3* and two TeSS fragments (Supplementary Table S14). The TeSS fragments were amplified from the telomerator plasmid pLM092 ^12^. To prevent possible circularization of the linear NeoChr, in NeoChr11 the sequence of the right TeSS was changed by modifying a few base pairs (PCR amplification with a mutated primer). Fragments were amplified by PCR using primers with 60 bp SHR flanks ^58^ (Supplementary Table S15). NeoChrs were assembled by transforming 200 fmol of each non-coding 2.5 kb *E*.*coli* DNA fragment, 200 fmol of each fluorescent marker, 200 fmol of the TeSS fragments, and 100 fmol of *CEN6/ARS4, HIS3* and the two *ARS* fragments. Transformants were checked by FACS, PFE and long-read Nanopore sequencing. One transformant (NeoChr10.13) missing only 2113 bp of fragment 7A was stocked as strain IMF22.

#### Construction of strains harboring NeoChrs designed for anthocyanin production

**Construction of the host strain IMX2770**. The host strain for the anthocyanin NeoChrs, IMX2270, was constructed by engineering of the SwYG strain IMX589 in which the minor paralogs of glycolysis and fermentation have been deleted and the major paralogs relocalized as a cluster to the *sga1* locus on chromosome IX ^4^. Using CRISPR/Cas9 editing ^37^, the minor paralogs of the pentose phosphate pathway, *GND2, NQM1, SOL4* and *TKL2*, were deleted to facilitate the subsequent relocalization of the major homologues to the NeoChrs. Additionally, the *ARO10* gene was removed to minimize phenylethanol production ^21^, resulting in strain IMX2270. The strain construction strategy is represented in Fig. 4, and the construction steps leading from the SwYG strain to IMX2270 are detailed in Supplementary Methods 3, Supplementary table 16-19. **Assembly of NeoChr25 and NeoChr26**. In strain IMX2270 the circular NeoChr26 and the linear NeoChr25 were assembled. The fragments for assembly were identical between the two NeoChrs namely: 7 pentose phosphate pathway genes (*ZWF1, TKL1, GND1, RKI1, TAL1, RPE1, SOL3*), 13 glycolysis genes (*HXK2, PGK1, FBA1, TPI1, TDH3, GPM1, ENO2, PGI1, PYK1, PDC1, ADH1, PFK1, PFK2*), 10 codon optimized *E. coli* shikimate pathway genes (*coEcAroG*^*fbr*^, *coEcAroB, coEcAroE, coEcAroL, coEcAroA, coEcAroD, coEcAroC, coEcTyrA*^*fbr*^, *coEcPheA*^*fbr*^, *coEcTyrB*), *HIS3* selection marker, two fluorescent markers (*mRuby2, mTurquoise2*), *CEN6/ARS4* and two ARS sequences. For NeoChr26 (circular), the telomerator fragment was transformed, while for NeoChr25 (linear) two TeSS were used instead. The fragments were amplified by PCR from the corresponding expression cassettes for all transcriptional units or from genomic DNA and other plasmids for auxiliary parts using primers with 60 bp SHR flanks (Supplementary Table 20-21). For each fragment 200 fmol was used, with the exception of *HIS3* and *CEN6/ARS4* fragments, where 100 fmol were added instead. The pooled transformation mix were concentrated with Vivacon 500 (Sartorius AG, Gottingen, Germany) up to a final volume of 50 *μ*l and transformed in IMX2270. Both the NeoChr25 (linear) and NeoChr26 (circular) transformants were screened by FACS and long-read Nanopore sequencing, and NeoChr25 was additionally also verified by PFE. After sequence confirmation using short-read whole genome sequencing, the strains were stocked as IMF27 (NeoChr25) and IMF29 (NeoChr26). **Removal of single locus glycolysis from native chromosome**. In IMF29 (circular NeoChr) the glycolytic cassette was removed from the *sga1* locus by induction of two double strand breaks (DSB) in the flanks of the cassette with the gRNA plasmid pUDR413, and providing a 120 bp repair fragment homologous to the upstream and downstream region of *sga1* (Supplementary Table 8 and 22). For IMF27 (linear NeoChr) the glycolytic cassette was replaced by a *KlURA3* transcriptional-unit by providing a *KlURA3* expression unit as repair fragment, obtained by PCR amplification from pMEL10 with primers containing flanks homologous to *sga1* locus (Supplementary Table 22). After discarding the gRNA plasmids the strains were stocked as IMF31 (linear NeoChr) and IMF32 (circular NeoChr). The circular NeoChr in strain IMF32 contained a mutation in the *RKI1* ORF. The mutation was repaired by inducing a DSB in the vicinity of the mutation with the gRNA plasmid pUDR756, targeting only the watermarked gene but not the native copy, and repairing it with a 120 bp repair fragment (Supplementary Table 8 and 23). The repair fragment contained the non-mutated sequence and a silent mutation of the PAM sequence. A correct transformant was confirmed by Sanger sequencing (Baseclear) and named IMF35. **Deletion of major PPP paralogs from their native chromosomal loci**. In a first transformation round, *ZWF1, SOL3, GND1* and *RKI1* were removed from IMF31 and IMF35 using the gRNA plasmids pUDR703 and pUDR700 and 120 bp repair fragments (Supplementary Table 6 and 24). Plasmid removal resulted in strains IMF33 and IMF36 with linear and circular NeoChr, respectively. In a second transformation *RPE1, TKL1* and *TAL1* were removed using the gRNA plasmids pUDR701 and pUDR702 and 120 bp repair fragments (Supplementary Table 7 and 24). After plasmid recycling, the strains were stocked as IMF34 and IMF40, for the linear and circular NeoChr strains respectively. **Integration of plant anthocyanin pathway**. The next step in strain construction was the integration of single copies of the plant genes of the anthocyanin pathway on the NeoChrs, *AtPAL1, coRcTAL1, coAtC4H, coAtCPR1, At4CL3, AtCHI1, coAtCHS3, coAtF3H, coGhDFR, coAtANS, coAt3GT*, under expression of strong, constitutive yeast promoters. The fragments were amplified from their corresponding Golden Gate expression plasmids with primers containing 60 bp SHRs for *in vivo* assembly (Supplementary Table 25-26). They were integrated in a single locus of the linear (strain IMF34) and circular (strain IMF40) NeoChrs by induction of a DSB in the *E*.*coli* fragment 16AB using gRNA plasmid pUDR765 (Fig. 3, Supplementary Table 7). The two outer fragments contained 60 bp homology to a stretch of 60 bp in the *E*.*coli* fragment16AB and in the flanking SHR DL. After verification of correct integration by PCR (Supplementary Table 26), the gRNA plasmid was removed by unselective growth on SMD and the strains were stocked as IMF41 (circular NeoChr) and IMF42 (linear NeoChr). **Integration of multiple copies of *coAtCHS3***. To integrate multiple copies of *coAtCHS3*, the *coAtCHS3* expression unit was amplified from pUDC352 using primers adding 60 bp homology flanks to the integration sites Chunk 7BC, chunk 15CD, SHR N and chunk 9CD in the linear NeoChr31 and *CAN1, X2, YPRCtau3* and *SPR3* in native chromosomes (Supplementary Table 6, 7 and 27). The *coAtCHS3* expression units carrying homology with native chromosomes were transformed to IMF42 using the gRNA plasmids pUDR771 and pUDR772 for targeted editing. After recycling of the gRNA plasmids, strain IMF44 (Lin, 5x *CoAtCHS3* (4 copies in native genome)) was obtained. IMF44 was then transformed with suitable *coAtCHS3* cassettes and gRNA plasmids pUDR780 and pUDR781 targeting four loci in NeoChr31 resulting in strain IMF47, containing nine copies of the *coAtCHS3* (five on NeoChrs and four on native chromosomes). ***coAtANS* repair**. *coAtANS* was repaired by integration of a correct copy of *coAtANS* in the *mTurquoise* gene of NeoChr33 in IMF47. The sequence of pUDC398 (*pSeENO2-coAtANS-tFUM1)* was verified and the transcriptional unit was amplified using primers 18740/18741 (Supplementary Table 28) adding flanks with homology to *mTurquoise* on NeoChr33. IMF47 was transformed with the template DNA and pUDR400 (gRNA-*mTurquoise*) for targeting *mTurquoise*. After PCR confirmation (Supplementary Table 28), pUDR400 was recycled and the final strain was stocked as IMF48 (linear NeoChr34, 9x *coAtCHS3*, correct *coAtANS*).

#### Strain verification by PCR and whole genome sequencing

All strains constructed by CRISPR/Cas9 were verified by diagnostic PCR. In addition, whole genome short read sequencing was performed in-house for IMF27, IMF29 on an Illumina MiSeq Sequencer (Illumina, San Diego, CA) as described previously ^5, 32^. Strains IMF41, IMF42, and IMF47 were sequenced on a NovaSeq 6000 at GenomeScan Leiden (GenomeScan, Leiden, Netherlands). Long-read sequencing of IMF22, IMF27, IMF29, IMF41, IMF42, IMF47 and IMF48 was performed in-house on MinION flow cells using the SQK-LSK109 sequencing kit with the EXP-NBD104 expansion kit (Oxford Nanopore Technologies, Oxford, United Kingdom) (Supplementary Methods 4). The sequences were deposited to NCBI (https://www.ncbi.nlm.nih.gov/) as a BioProject under the accession number PRJNA738851.

### Characterization of NeoChrs fidelity, stability and toxicity

**Fluorescence-based sorting by flow cytometry**. Flow cytometry employing an BDFACSAria− II SORP Cell Sorter (BD Biosciences, Franklin Lakes, NJ), was used to sort single cells in the chromosome stability experiment, as well as for screening of fluorescent transformants bearing the constructed NeoChrs as both described earlier by Postma *et al*. ^5^. **Karyotyping with PFE analysis**. Chromosomes separation and size determination was performed using contour-clamped homogeneous electric field electrophoresis as described earlier by Postma *et al*. ^5^. **Quantification of strain viability and NeoChr stability**. NeoChr stability was assessed based on plating cells from shake flask cultures on selective and non-selective growth medium with respect to the NeoChr as described by Postma *et al*. ^5^. On the second and fifth day, strain viability and stability was measured, on the third and fourth day, only the cell density (OD_660_) of overnight cultures was measured. For the test-chromosome bearing strains (IMF22 and IMF23) viability and stability was based on sorting of 96 cells, while for CEN.PK113-7D, IMF27, IMF29, IMF34, IMF40, IMF41, IMF42 and IMF47, 384 cells were sorted on a microtiter plate. **Determination of specific growth rates**. For determination in shake flasks, strains were inoculated from a -80°C freezer stock in a 500 mL shake flask with 100 mL medium and grown until late-exponential phase. Cells were transferred to fresh 100 mL medium and grown until mid-exponential phase. Finally, from these cultures, shake flasks containing 100 mL medium were inoculated at an OD_660_ of 0.3. Cell density was measured with a 7200 Visible spectrophotometer (Cole-Parmer, Staffordshire, United Kingdom). A maximum specific growth rate (µ_max_) was calculated from at least biological duplicates, using five data points and at least two doublings in the exponential phase. For growth rate determination in multi titer plates, a Growth Profiler (Enzyscreen, Heemstede, NL) was used as previously described ^5^ using plate 06 and a: 0.084327, b:5.35 × 10^−8^, c: 4.398348, d: -0.41959 as constants. Growth rates were calculated from six biological replicates, using ODs between 1 and 8.

### Characterization of aromatics production by the engineered strains

**Shake flasks**. The strains were inoculated in biological triplicates in 500 mL shake flasks containing 100 mL synthetic medium with 20 g L^-1^ glucose. Urea was used as sole nitrogen source to prevent acidification of the medium, thereby allowing respiration of the produced ethanol. After 72 hours of cultivation, the optical density was measured using a Jenway 7200 spectrometer (Jenway, Staffordshire, United Kingdom) at 660 nm and aromatics were quantified as described below. **Aerobic batch bioreactors**. Strains IMF41, IMF42 and IMF48 were characterized in bioreactors. 2L bioreactors (Applikon, Delft, Netherlands) were filled with 1.0 L synthetic medium, containing 20 g L^-1^ glucose and ammonium sulfate as nitrogen source. Exponentially growing cells were used to inoculate the bioreactors at an initial biomass concentration of around 0.12 g L^-1^. The cultivations were performed at 30 °C, 800 RPM using 0.5 L min^-1^ pressurized air to sparge the bioreactor with oxygen. Automated addition of 2 M KOH and 2 M H_2_SO_4_ ensured maintenance of the culture pH at 5.0. Samples were taken at regular time intervals for optical density, metabolite concentrations and cell dry weights determinations. Cell dry weight, organic acids, sugars and ethanol concentrations were measured as previously described ^57, 59^. **Analysis of aromatics**. Extracellular naringenin and upstream aromatics were quantified by HPLC as previously described ^57^ and elaborated upon in Supplementary Methods 5A. Sample preparation and quantification of intracellular and extracellular flavonoids downstream naringenin using LC-MS/MS is described in Supplementary Methods 5B. Detection of P3G is illustrated in Supplementary Fig. 12.

## Abbreviations

Metabolites

2PE: 2-phenylethanol
CIN: cinnamic acid
COCOA: coumaroyl-CoA
COUM: coumaric acid
DHK: dihydrokaempferol
K3G: kaempferol 3-O-glucoside
KAE: kaempferol
LPE: leucopelargonidin
NAR: naringenin
NARCC: naringenin chalcone
P3G: pelargonidin 3-O-glucoside
PAA: phenylacetic acid
PEL: pelargonidin
PHE: L-phenylalanine
PHLOR: phloretic acid
*p*OHPPY: *p-*hydroxyphenylpyruvate
*p*OH2PE: *p*-hydroxyphenylethanol
*p*OHPAA: *p*-hydroxyphenylacetic acid
PPY: phenylpyruvate
TYR: tyrosine

Enzymatic reaction

3GT: anthocyanin 3-*O*-glucosyltransferase
ANS: anthocyanidin synthase
C4H: cinnamate 4-hydroxylase
4CL: 4-coumarate CoA ligase
CHI: chalcone isomerase
CHS: chalcone synthase
CPR: cytochrome P450 reductase
DFR: dihydroflavonol 4-reductase
F3H: flavanone 3-hydroxylase
PAL: phenylalanine ammonia lyase
TAL: tyrosine ammonia lyase

Methods

FACS: fluorescence-activated cell sorting
PFE: pulsed field electrophoresis

## Data availability

Genome sequence data are available at NCBI as Bioproject under the accession number PRJNA738851.

## Supporting information

**Supplementary Figure 1**: Reactions from glucose to pelargonidin 3-O-glucoside.

**Supplementary Figure 1**: Flow cytometric analysis of the test linear neochromosomes.

**Supplementary Figure 3**: Separation of the test linear neochromosomes with pulsed field electrophoresis.

**Supplementary Figure 4**: Sequencing results of test linear neochromosomes.

**Supplementary Figure 5**: NeoChr copy number estimation based on fluorescence.

**Supplementary Figure 6**: NeoChr copy number estimation based on sequencing.

**Supplementary Figure 7**: Flow cytometric analysis of (linear) NeoChr25 and (circular) NeoChr26 designed for anthocyanin production.

**Supplementary Figure 8**: Separation of (linear) NeoChr25 transformants on pulsed-field electrophoresis.

**Supplementary Figure 9**: Duplication and inversion of four plant genes in linear NeoChr25 and circular Neochr26.

**Supplementary Figure 10**: schematic representation of coAtANS mutation in strains IMF41, IMF42, IMF44 and IMF47.

**Supplementary Figure 11**: Substrates and products profiles during aerobic batch cultivation in bioreactors of IMF41, IMF42 and IMF48.

**Supplementary Figure 12**: Detection and quantification of pelargonidin and pelargonidin 3-O-glucoside by LC-MS/MS.

**Supplementary Table 1**: Promoter-gene-terminator combinations in the NeoChrs.

**Supplementary Table 2**: Sequence fidelity of NeoChrs.

**Supplementary Table 3**: Amino acid substitution in native genome of NeoChr strains.

**Supplementary Table 4**: Extracellular concentration of aromatic compounds produced by engineered **S. cerevisiae** strains in shake flask cultures.

**Supplementary Table 5**: Physiological characterization of anthocyanin-producing strains grown in bioreactors.

**Supplementary Table 6**: *S. cerevisiae* strains used in this study.

**Supplementary Table 7**: - Neochromosome configurations.

**Supplementary Table 8**: Plasmids.

**Supplementary Table 9**: pROS/pMEL gRNA primers.

**Supplementary Table 10**: Primers to check correct construction of gRNA plasmids.

**Supplementary Table 11** Primers to make golden gate part plasmids and expression plasmids with Gibson assembly.

**Supplementary Table 12**: Diagnostic primers to check golden gate part plasmids.

**Supplementary Table 13**: Diagnostic primers to check golden gate and Gibson assembly expression plasmids, for PCR and Sanger sequencing.

**Supplementary Table 14**: List of NeoChr10 and NeoChr11 chromosome parts.

**Supplementary Table 15**: List of primers for amplifying NeoChr10 and NeoChr11 chromosome parts.

**Supplementary Table 16**: Primers for of *amdSYM* deletion.

**Supplementary Table 17**: Primers for deletion of *GND2, NQM1, SOL4* and *TKL2*.

**Supplementary Table 18**: Primers for deletion of *ura3, his3* and *SpHIS5*.

**Supplementary Table 19**: Primers for deletion of *ARO10*.

**Supplementary Table 20**: List of NeoChr25 (linear) and NeoChr26 (circular) chromosome parts.

**Supplementary Table 21**: List of primers for amplifying NeoChr25 and NeoChr26 chromosome parts.

**Supplementary Table 22**1: Primers for glycolysis deletion.

**Supplementary Table 23**: Primers to repair *RKI1* mutation in IMF32.

**Supplementary Table 24**: Primers for deletion of native *ZWF1, GND1, SOL3, RKI1, TAL1, TKL1* and *RPE1* ORFs.

**Supplementary Table 25:** Parts of the “basic design” of the anthocyanin pathway.

**Supplementary Table 26**: List of primers for amplifying the fragments of the “basic design” of the anthocyanin pathway and diagnosing integration.

**Supplementary Table 27**: List of primers for amplifying the fragments of the “elaborate design” of the anthocyanin pathway with several copies of the chalcone synthase and diagnostic PCR.

**Supplementary Methods 1**: Strains, growth medium and maintenance.

**Supplementary Methods 2**: Molecular biology techniques.

**Supplementary Methods 3**: Detailed construction of the host strain IMX2770.

**Supplementary Methods 4**: MinION long-read sequencing.

**Supplementary Methods 5**: Analysis of aromatics.

## Acknowledgements

We thank Carol de Ram for technical support with LC-MS sample preparation.

## Author contributions

E.D.P. and E.-J.H., designed the research, carried out the experiments, analyzed the results, and wrote the paper. V.M. and J.G. assisted in chromosome design and construction. P.dlT. and M.vdB. assisted whole genome sequencing and analysis. C.M. assisted in strain characterization in bioreactor. M.P. developed and applied the method for LC-MS/MS detection of metabolites, and analyzed the results. P.D.-L. and J.-M.D. designed the research, supervised the project and wrote the paper. All authors approved the final manuscript.

