## Supplementary Material for "Modular, synthetic chromosomes as new tools for large scale engineering of metabolism"

### Supplementary Figure 1 - Reactions from glucose to pelargonidin 3-O-glucoside

Yellow: glycolysis an ethanolic fermentation. Blue: pentose phosphate pathway. Brown: *E. coli* shikimate pathway. Green: plant anthocyanin pathway. The gene names encoding the enzymes involved in the indicated reactions are indicated in italics. Genes deleted in the present study are indicated in red and underlined. *Ec E. coli, At Arabidopsis thaliana, Rc Rhodobacter capsulatus, co* codon optimized. *fbr* feedback resistant, Glc glucose, Glc-6P glucose 6-phosphate, Fru-6p fructose-6-phosphate, Fru-1,6-BP fructose 1,6-biphosphate, GAP glyceraldehyde 3-phosphate, DHAP dihydroxyacetone, 1,3-BPG 1,3-biphosphoglycerate, 3-PG 3-phosphoglycerate,2-PG 2-phosphoglycerate, PEP phosphoenolpyruvate, Pyr pyruvate, AcAL acetaldehyde, EtOH ethanol, 6p-GLCN-lac 6-phosphogluconolactone, 6p-GLCN 6-phosphoglucono, RL5P ribulose 5-phosphate, R5P ribose 5-phosphate, S7P sedoheptulose 7-phosphate, XUL-5P xylulose 5-phosphate, GAP glyceraldehyde 3-phosphate, Ery-4P erythrose 4-phosphate, DAHP 3-deoxy-D-arabino-heptulosonate-7-P, DHQ 3-dehydroquinate, DHS 3-dehydroshikimate, SHIK shikimate, SHP shikimate 3-phosphate, EP3P 5-enolpyruvoyl-shikimate 3-phosphate, CHA chorismate, PPA prephenate, PPY phenylpyruvate, PAC phenylacetaldehyde 2PE 2-phenylethanol, PAA phenylacetic acid, PHE L-phenylalanine, *p*OHPPY *p*-hydroxyphenylpyruvate, *p*OHPAC *p*-hydroxyphenylacetaldehyde, *p*OH2PE, *p*-hydroxyphenylethanol, *p*OHPAA, *p*-hydroxyphenylacetic acid, TYR tyrosine, COUM coumaric acid, CIN cinnamic acid, COCOA coumaroyl-CoA, NARCC naringenin chalcone, PHLOR phloretic acid, NAR naringenin, DHK dihydrokaempferol, KAE kaempferol, K3G kaempferol 3-O-glucoside, LPE, leucopelargonidin, PEL pelargonidin, P3G pelargonidin 3-O-glucoside

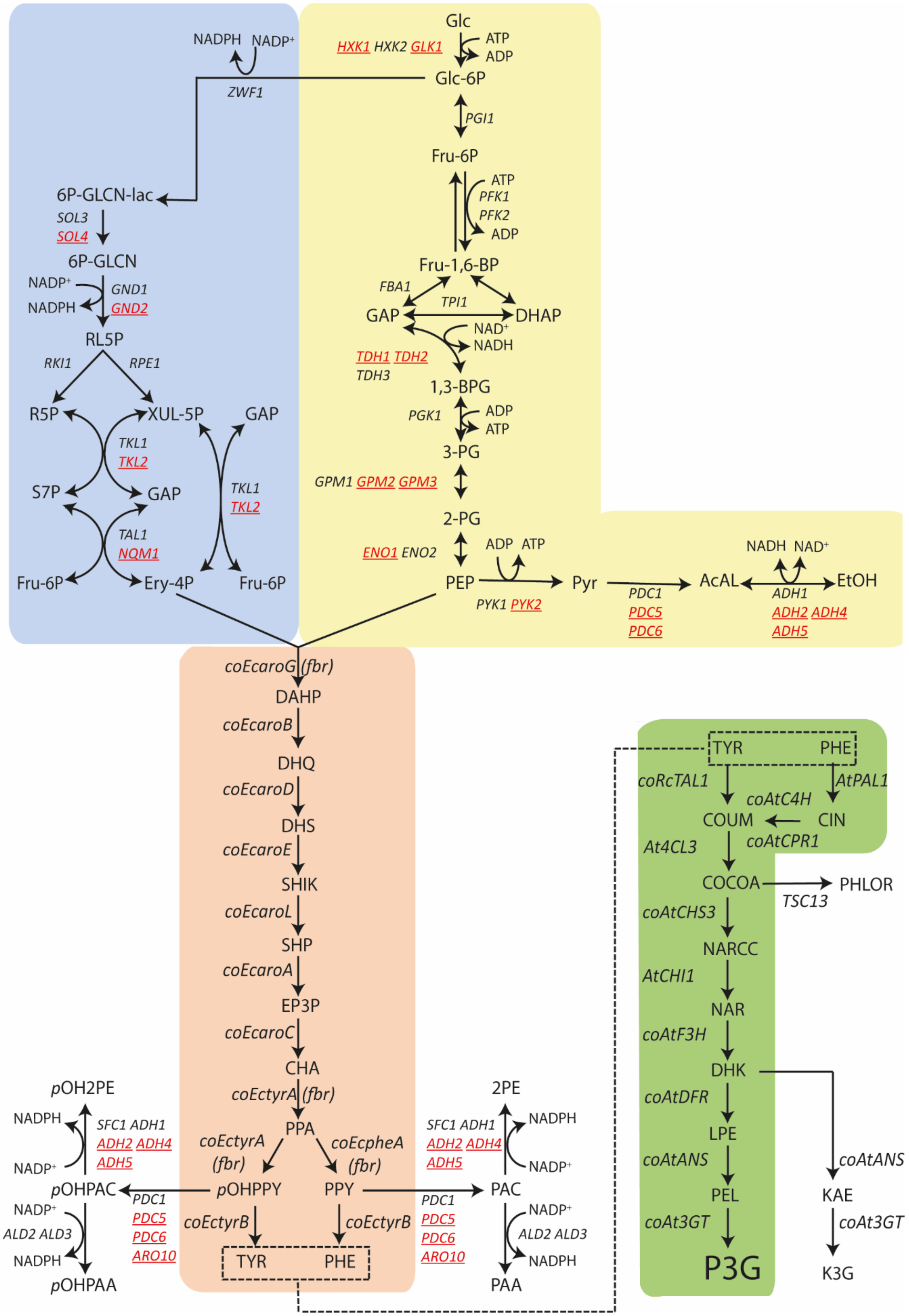

Glc
HXK1 HXK2 GLK1
ATP
ADP
Glc-6P
NADPH NADP+
ZWF1
PGI1
Fru-6P
ATP
PFK1
PFK2
ADP
Fru-1,6-BP
FBA1
TPI1
GAP
DHAP
NAD+
NADH
TDH1 TDH2
TDH3
1,3-BPG
PGK1
ADP
ATP
3-PG
GPM1 GPM2 GPM3
2-PG
ENO1 ENO2
ADP ATP
PEP
PYK1 PYK2
Pyr
PDC1
PDC5
PDC6
AcAL
NADH NAD+
ADH1
ADH2 ADH4
ADH5
EtOH
6P-GLCN-lac
SOL3
SOL4
6P-GLCN
NADP+
NADPH
GND1
GND2
RL5P
RKI1
RPE1
R5P
XUL-5P
GAP
TKL1
TKL2
S7P
GAP
TKL1
TKL2
TAL1
NQM1
Fru-6P
Ery-4P
Fru-6P
coEcaroG (fbr)
DAHP
coEcaroB
DHQ
coEcaroD
DHS
coEcaroE
SHIK
coEcaroL
SHP
coEcaroA
EP3P
coEcaroC
CHA
coEctyrA (fbr)
PPA
coEctyrA
(fbr)
coEcpheA
(fbr)
pOHPPY
PPY
coEctyrB
coEctyrB
TYR PHE
pOH2PE
NADPH
NADP+
SFC1 ADH1
ADH2 ADH4
ADH5
pOHPAC
PDC1
PDC5
PDC6
ARO10
NADP+
ALD2 ALD3
NADPH
pOHPAA
2PE
SFC1 ADH1
ADH2 ADH4
ADH5
NADPH
NADP+
PAC
PDC1
PDC5
PDC6
ARO10
NADP+
ALD2 ALD3
NADPH
PAA
TYR PHE
coRcTAL1
AtPAL1
coAtC4H
COUM
CIN
coAtCPR1
At4CL3
COCOA
PHLOR
TSC13
coAtCHS3
NARCC
AtCHI1
NAR
coAtF3H
DHK
coAtDFR
LPE
coAtANS
PEL
coAt3GT
P3G
coAtANS
KAE
coAt3GT
K3G

### Supplementary Figure 2 – Flow cytometric analysis of the test linear neochromosomes

Cells from shake flask cultures were analysed by FACS. The fluorescence is plotted on the y-axis and the forward scatter (FSC-A) on the x-axis. Negative control: CEN.PK113-7D. Positive controls: IMC111 (mRuby2), IMC112 (mTurquoise2) and IMF6 (mRuby2 and mTurquoise2). Gates for fluorescence of the two different fluorescent proteins were drawn based on the IMC111 and IMC112 controls. Approximately 10000 events are shown for each plot.

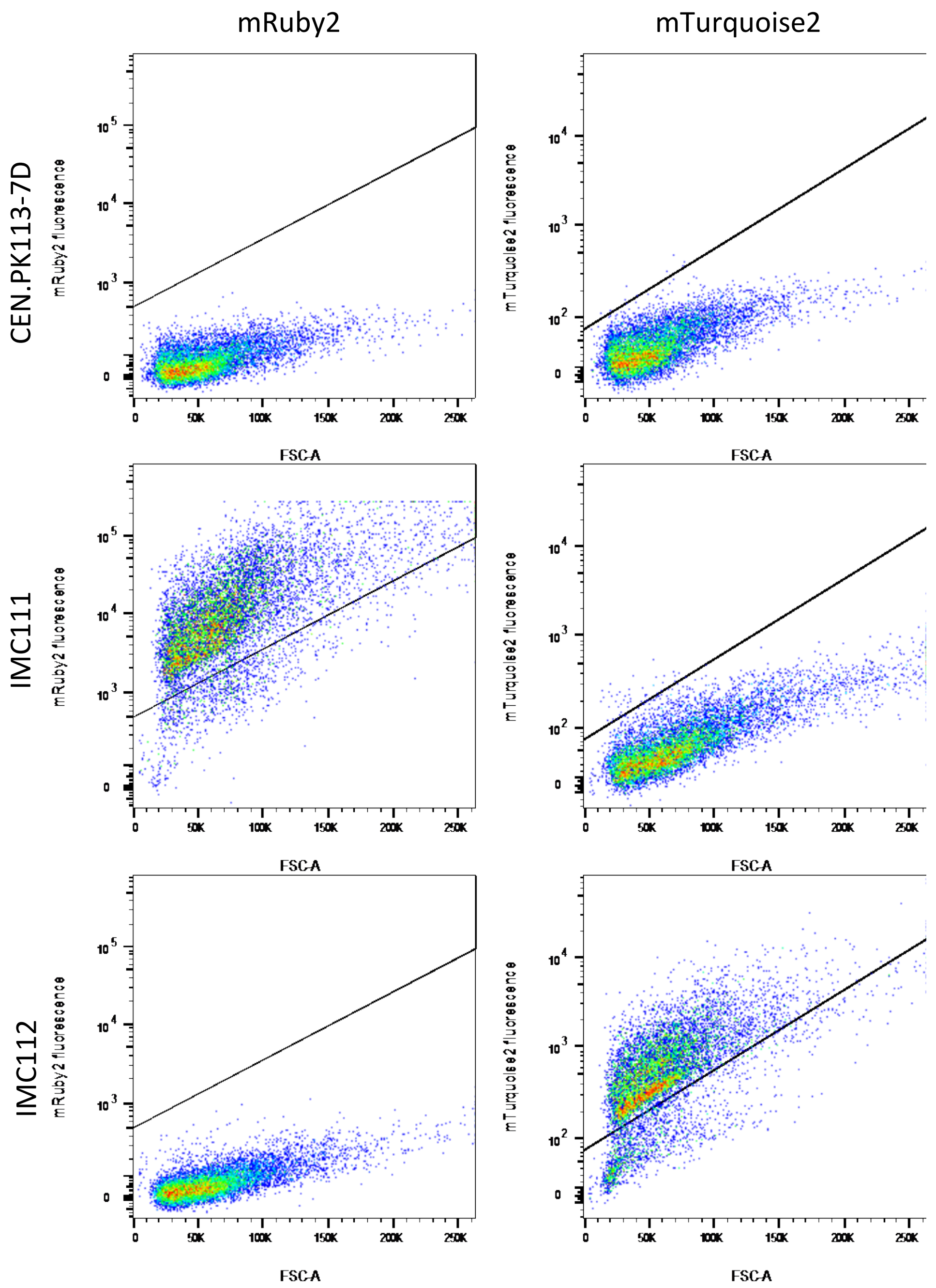

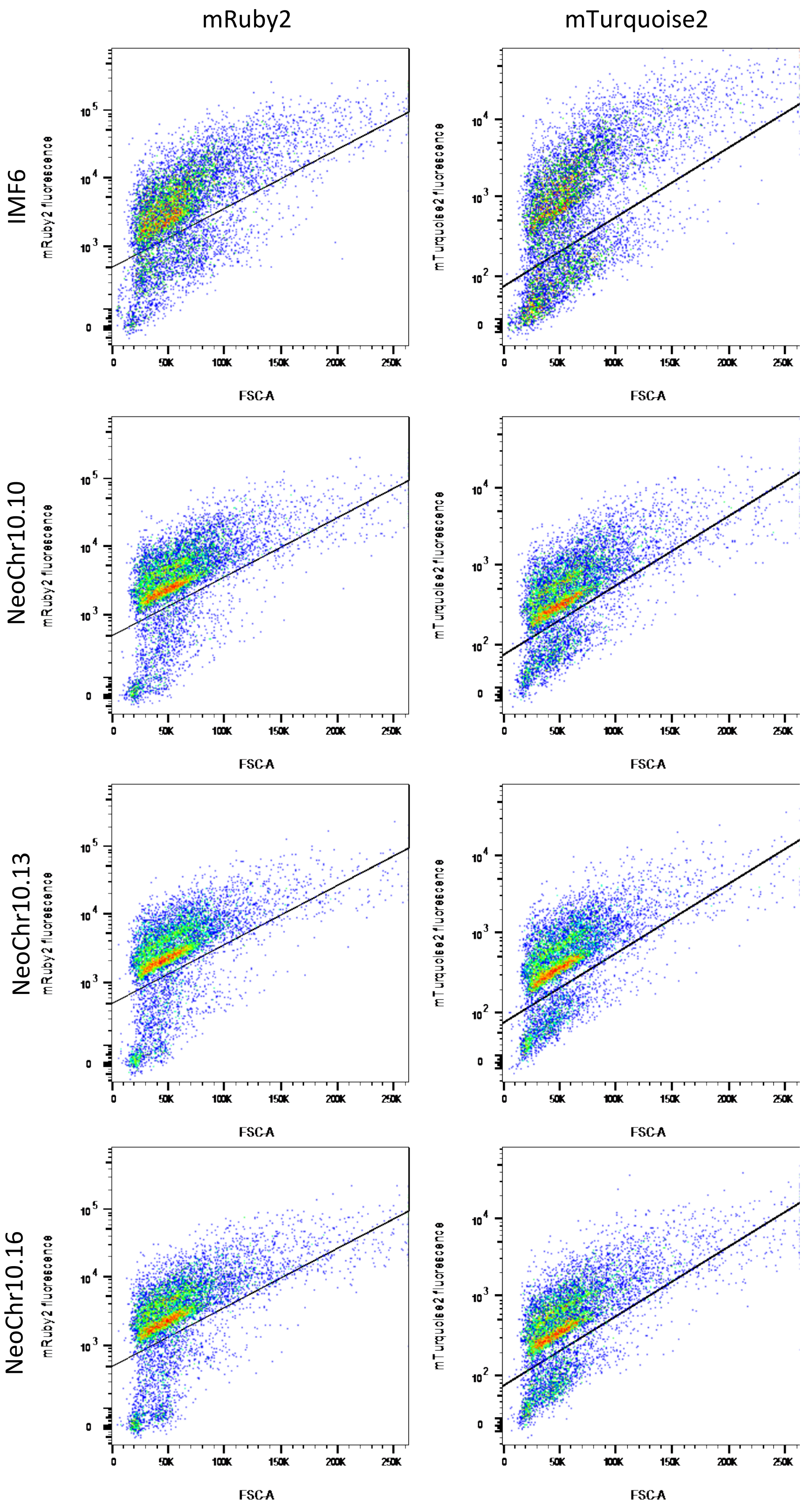
mRuby2
mTurquoise2
IMF6
NeoChr10.10
NeoChr10.13
NeoChr10.16
mRuby2 fluorescence
mTurquoise2 fluorescence
FSC-A
0
50K
100K
150K
200K
250K

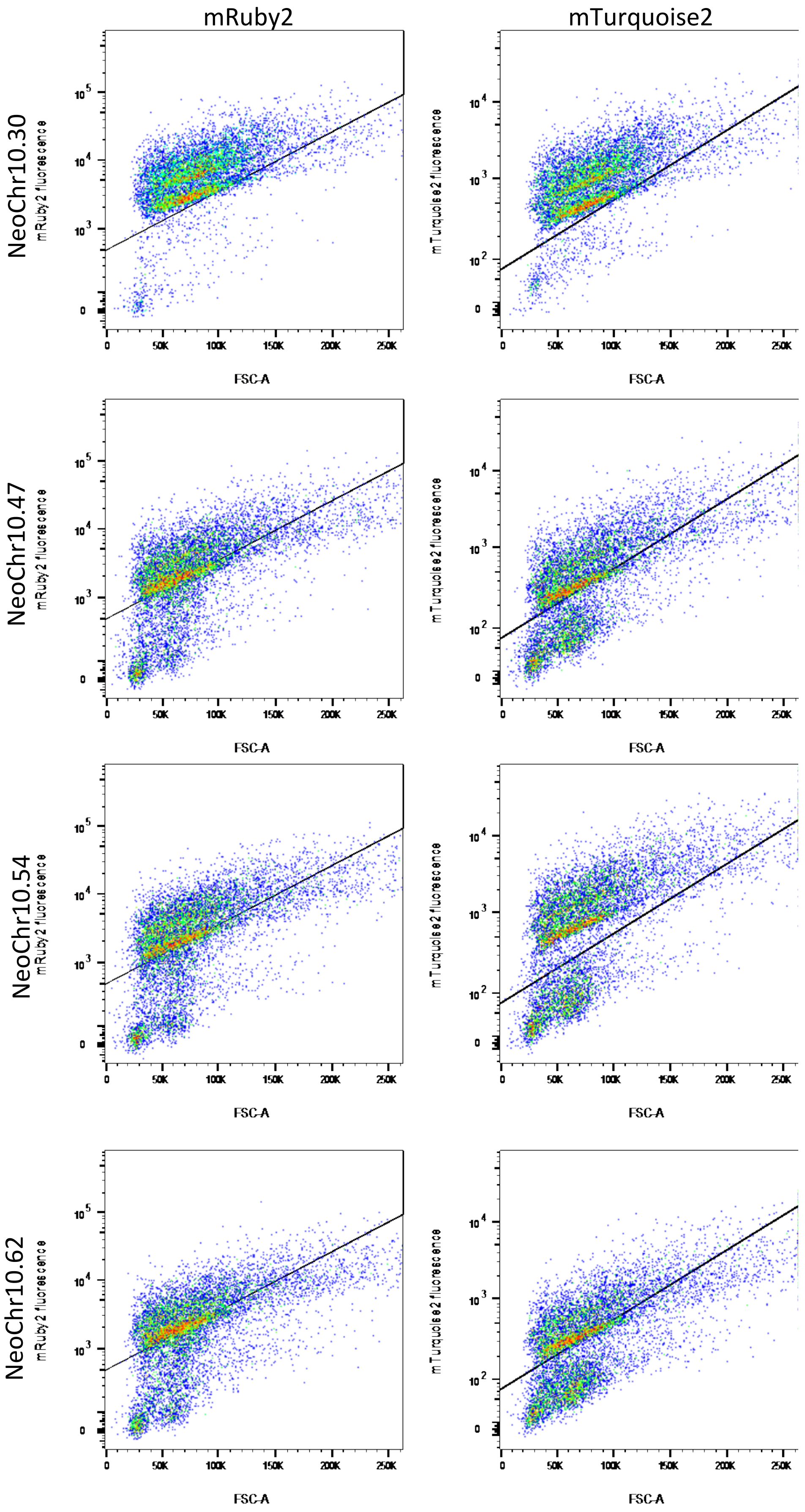
mRuby2
mTurquoise2
NeoChr10.30
NeoChr10.47
NeoChr10.54
NeoChr10.62
mRuby2 fluorescence
mTurquoise2 fluorescence
FSC-A
0
50K
100K
150K
200K
250K
$10^2$
$10^3$
$10^4$
$10^5$

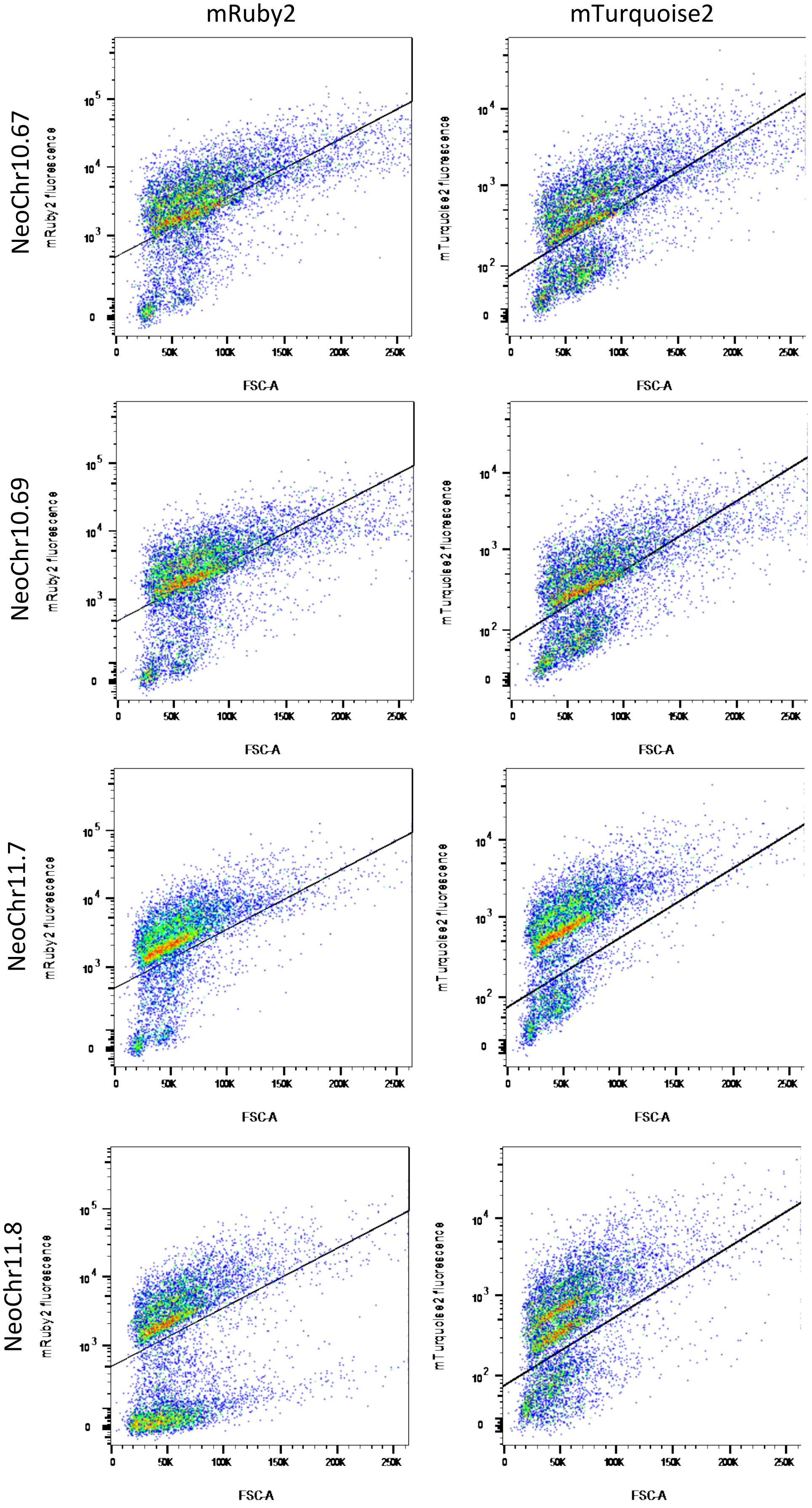
mRuby2
mTurquoise2
NeoChr10.67
NeoChr10.69
NeoChr11.7
NeoChr11.8
mRuby2 fluorescence
mTurquoise2 fluorescence
FSC-A
0
50K
100K
150K
200K
250K
$10^2$
$10^3$
$10^4$
$10^5$

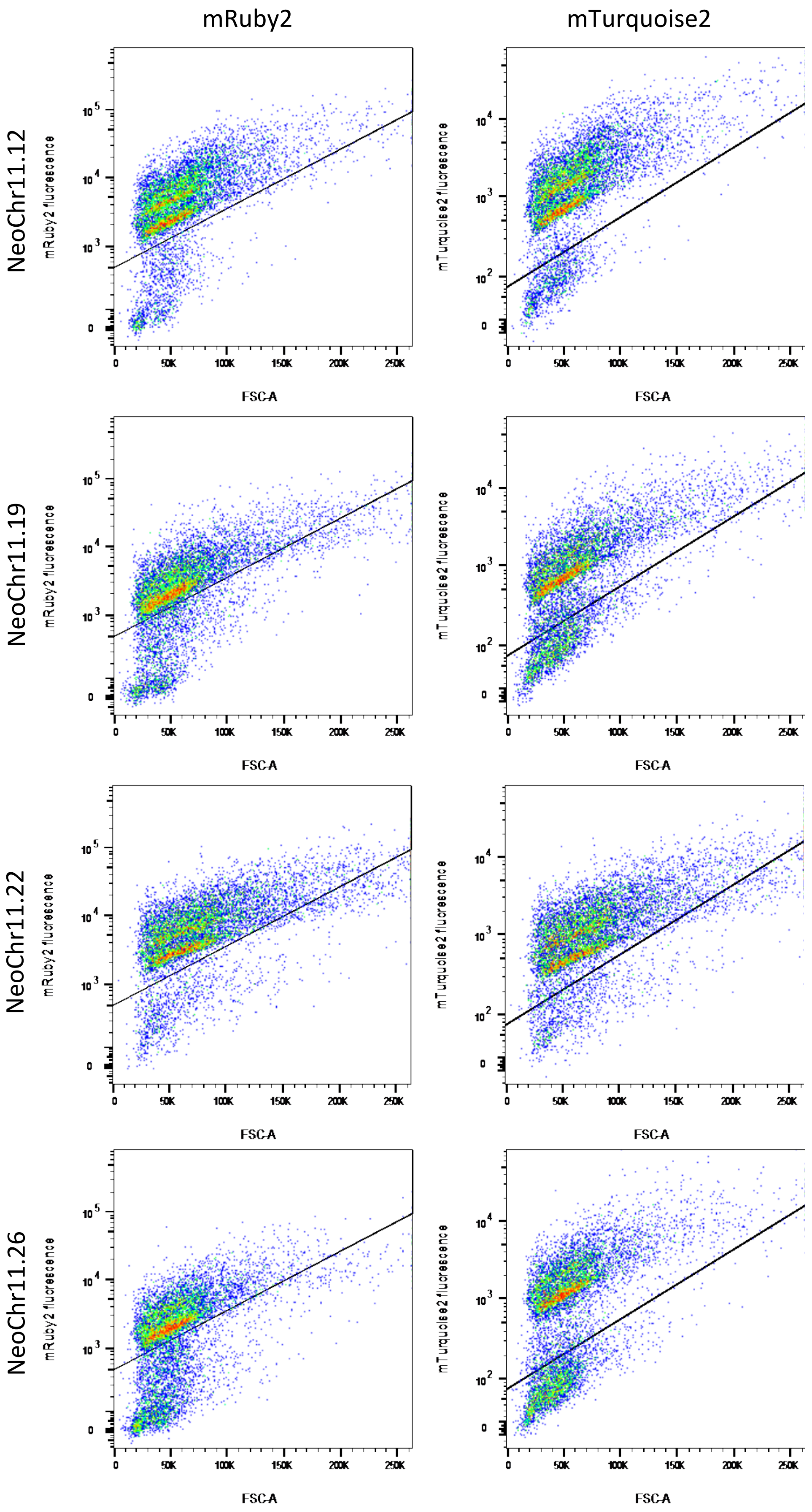
mRuby2
mTurquoise2
NeoChr11.12
NeoChr11.19
NeoChr11.22
NeoChr11.26
mRuby2 fluorescence
mTurquoise2 fluorescence
FSC-A
0
50K
100K
150K
200K
250K
$10^2$
$10^3$
$10^4$
$10^5$

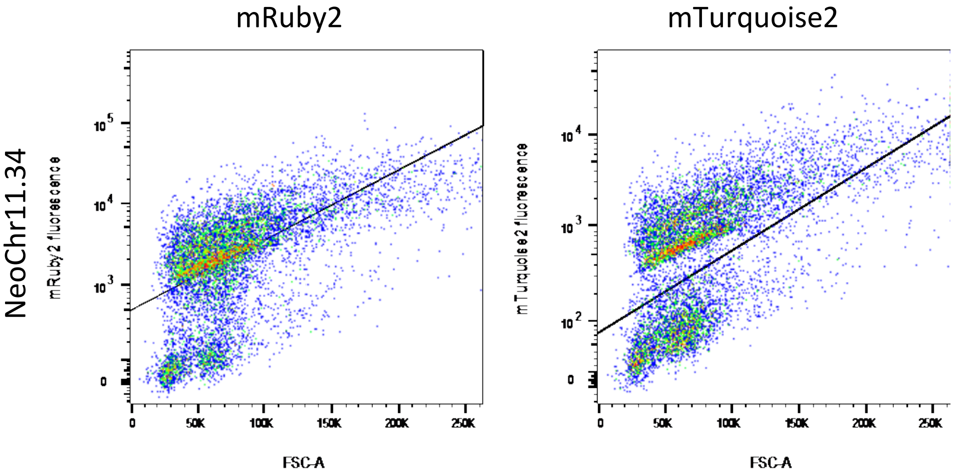
mRuby2
mTurquoise2
NeoChr11.34
mRuby2 fluorescence
mTurquoise2 fluorescence
FSC-A
FSC-A

### Supplementary Figure 3 - Separation of the test linear neochromosomes on pulsed-field electrophoresis

**A.** 1) IMX1338: control strain without neochromosome. 2) IMF6: control strain with 100 kb in plug linearized neochromosome NeoChr1. 3) IMF23: strain with 100 kb in plug linearized NeoChr12. 4) NeoChr10.10: correct size. 5)NeoChr10.13: correct size. 6) NeoChr10.16: wrong size. 7) NeoChr11.7: no visible neochromosome. 8) NeoChr11.8: wrong size. 9) NeoChr11.12: wrong size. 10) NeoChr11.19: correct size. 11)NeoChr11.22: wrong size. 12) NeoChr11.26: no visible neochromosome. 13) Size ladder

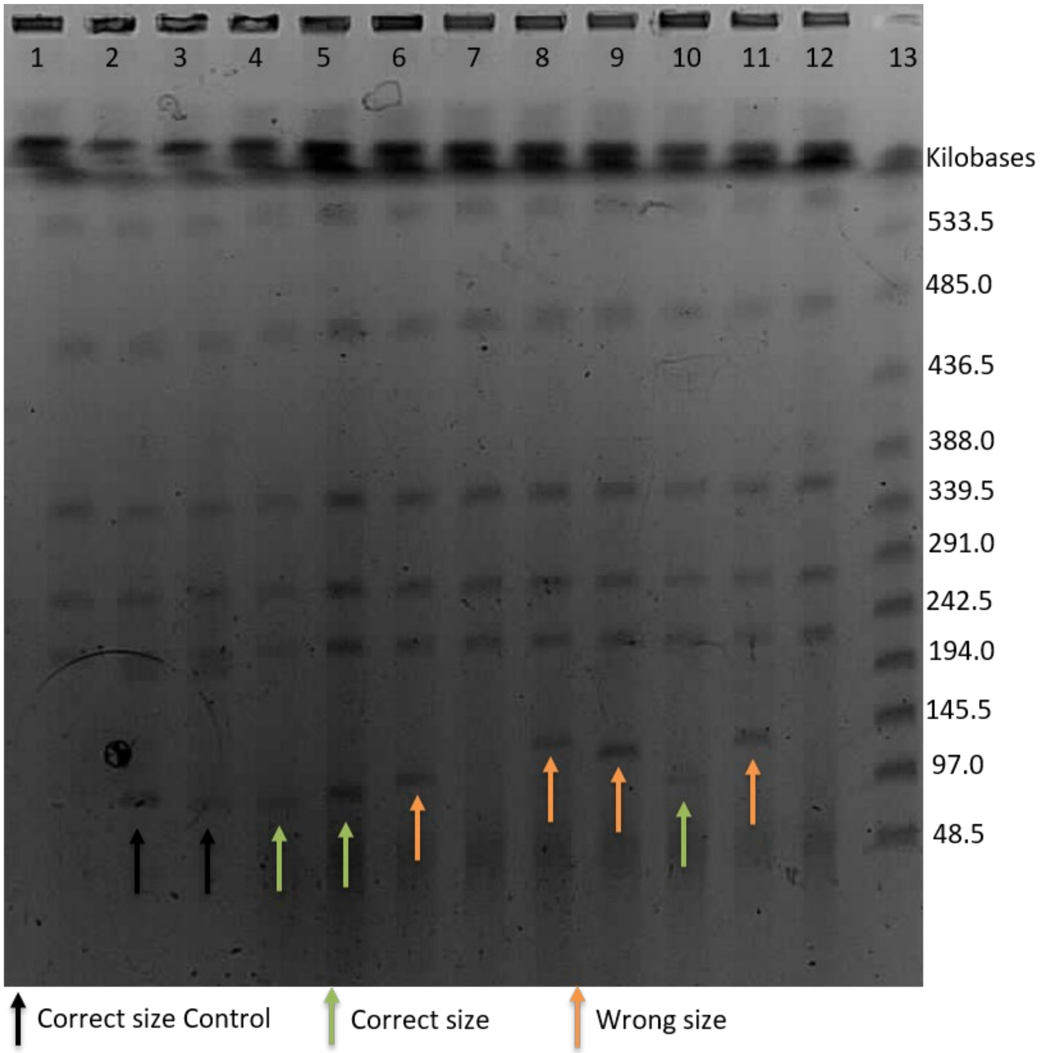

**B.** 1) IMX1338: control strain without neochromosome. 2) IMF6: control strain with 100 kb in plug linearized neochromosome (NeoChr1). 3) NeoChr10.30: no visible neochromosome. 4) NeoChr10.47: correct size. 5)NeoChr10.54: correct size. 6) NeoChr10.60: no visible neochromosome. 7) NeoChr10.62: correct size. 8) NeoChr10.67: correct size. 9) NeoChr10.69: correct size. 10) NeoChr11.29: no visible neochromosome. 11) NeoChr11.34: wrong size. 12) Size ladder

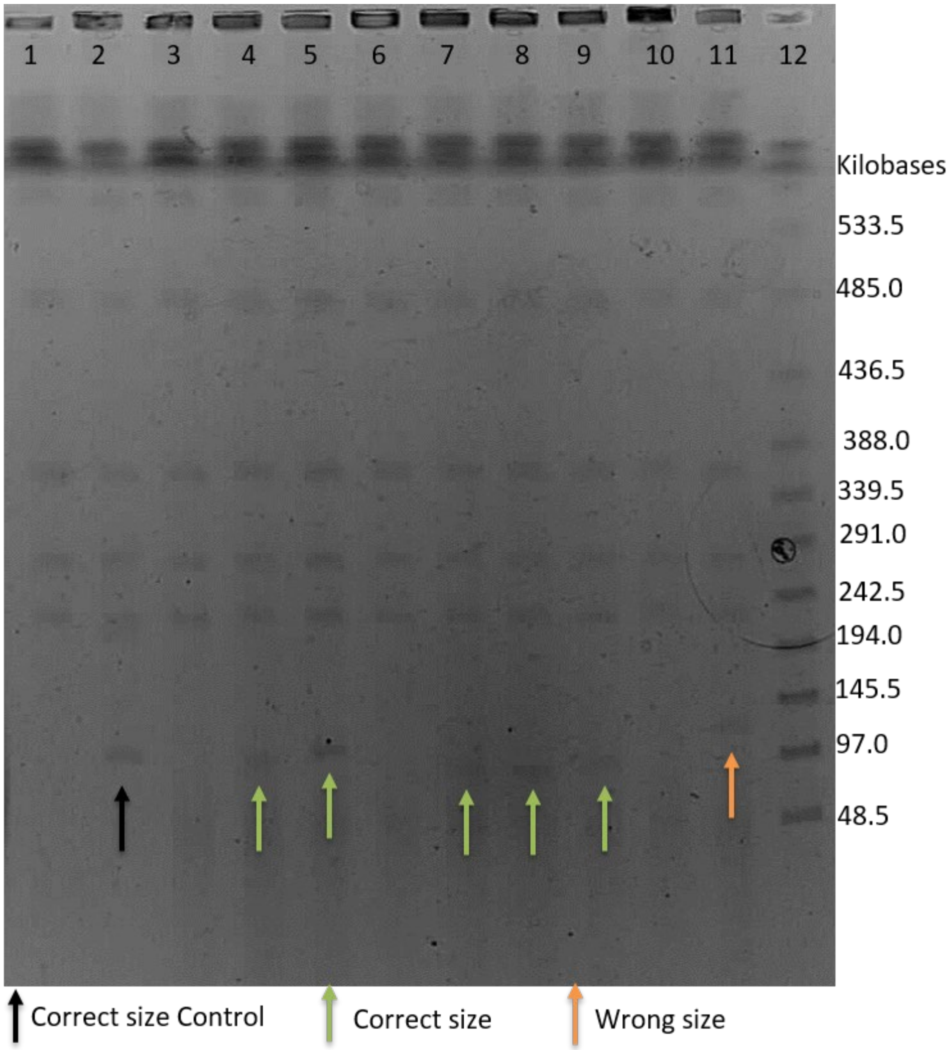

### Supplementary Figure 4 - Sequencing results of test linear neochromosomes

*in silico* fragment configuration in NeoChr10 / NeoChr11 as well as *in vivo* fragment configuration of the neochromosome transformants as measured by long-read nanopore sequencing. The fragments of the *in silico* design which are present in the neochromosome transformants are connected by the same colored line. A dotted line indicates an area which is inverted. Fragments are color coded as follows: blue represents a correctly assembled fragment; red represents a missing fragment; green represents a duplicated fragment; and yellow indicates an inverted fragment.

A) NeoChr10.13 and NeoChr10.47 are missing an internal part of chunk 7A. B) NeoChr10.69 is missing 3 chunks: 8A, 15B and 4D. C) NeoChr10.10 is missing 4 chunks: 7A, 15C, 16B and 4D. D) NeoChr10.67 is missing 6 chunks: 7A, 19D, 16B, 4B, 4D and 9D

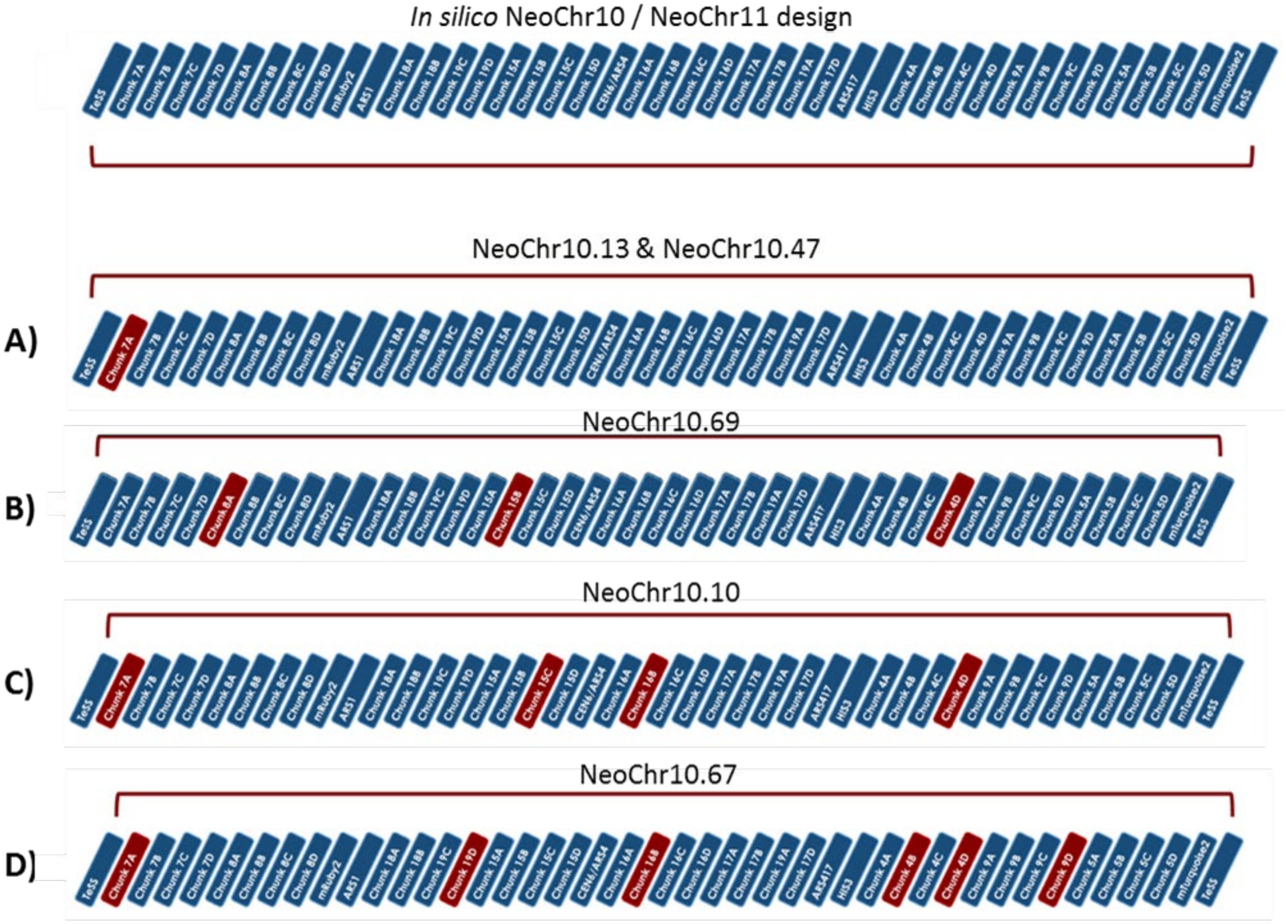

E) NeoChr10.62 is missing 4 chunks: 15B, 17B, 19A and 17D. In addition, a region containing the chunks 4A, 4B, 4C and 4D is duplicated

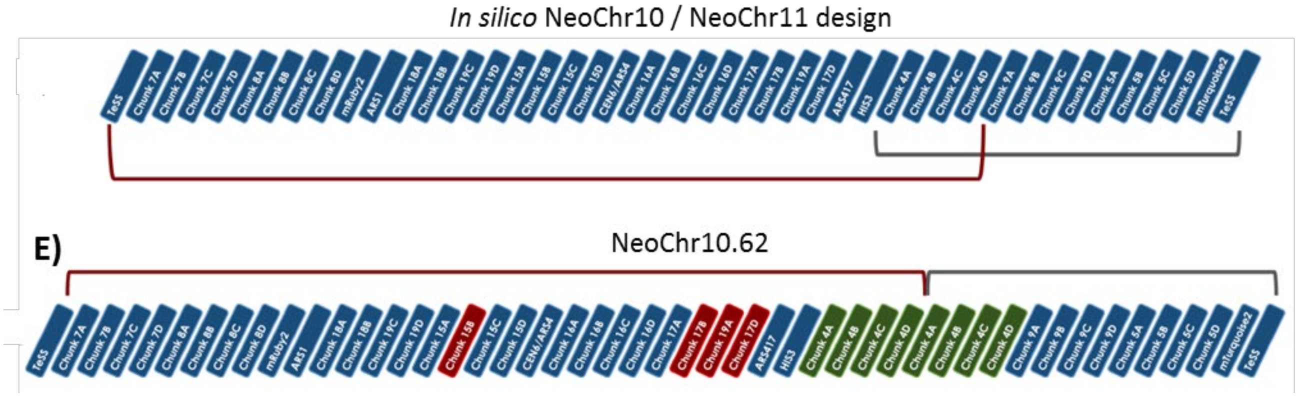


F) Neochr10.54 has a large inversion from 8D until 19A, from this region 2 chunks are missing: 15C and 18B. This region is link to a region spanning from 19A (which is thus duplicated) until the right telomere. From this region chunk 17D is missing

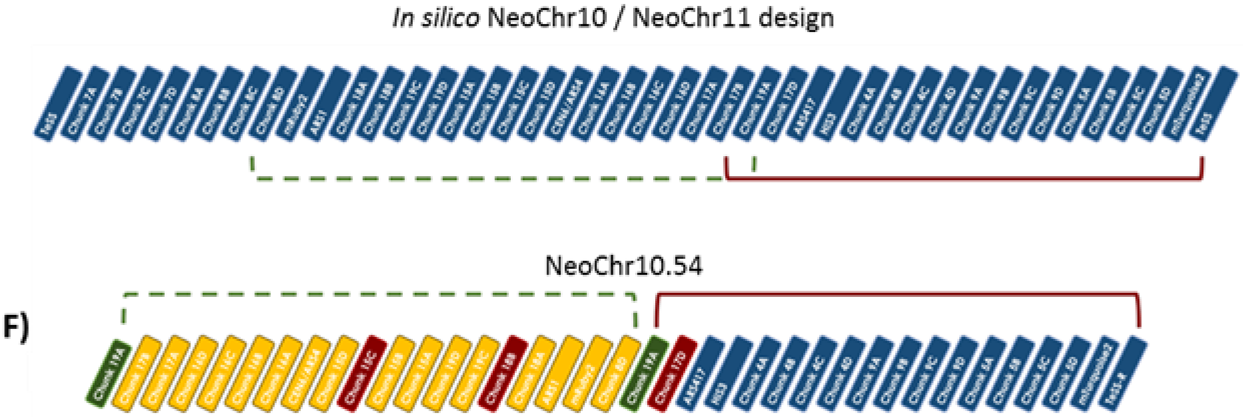


G) NeoChr11.19 contains several duplicated and inverted areas, from one area chunk 16A is missing.

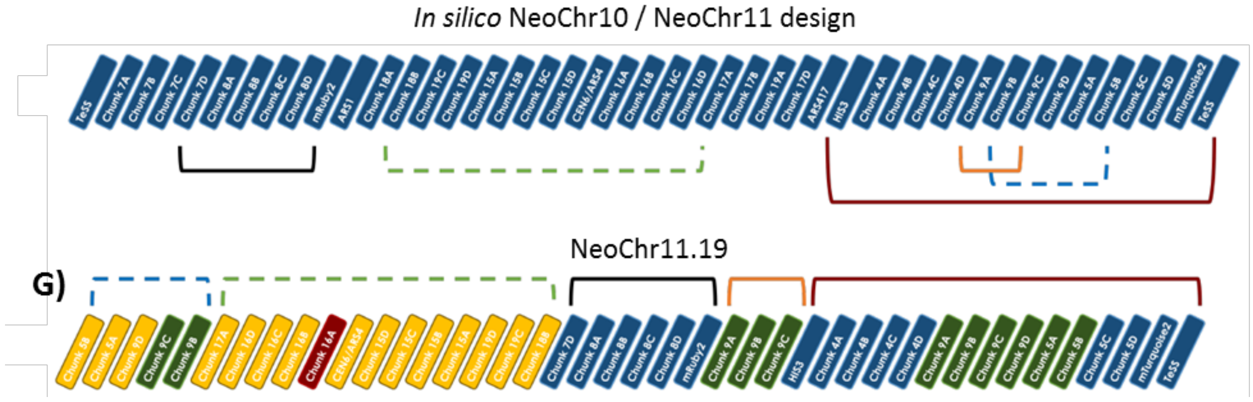

### Supplementary Figure 5 - NeoChr copy number estimation based on fluorescence

mRuby2 and mTurquoise2 fluorescence was measured by flow cytometry. CEN.PK113-7D with no fluorescent markers was used as negative control. IMX2224 and IMX2226 with a single copy of *mRuby2* and *mTurquoise2* integrated in the genome, respectively, were used as positive controls. All strains showed a fluorescence corresponding to the expected NeoChr. copy number.

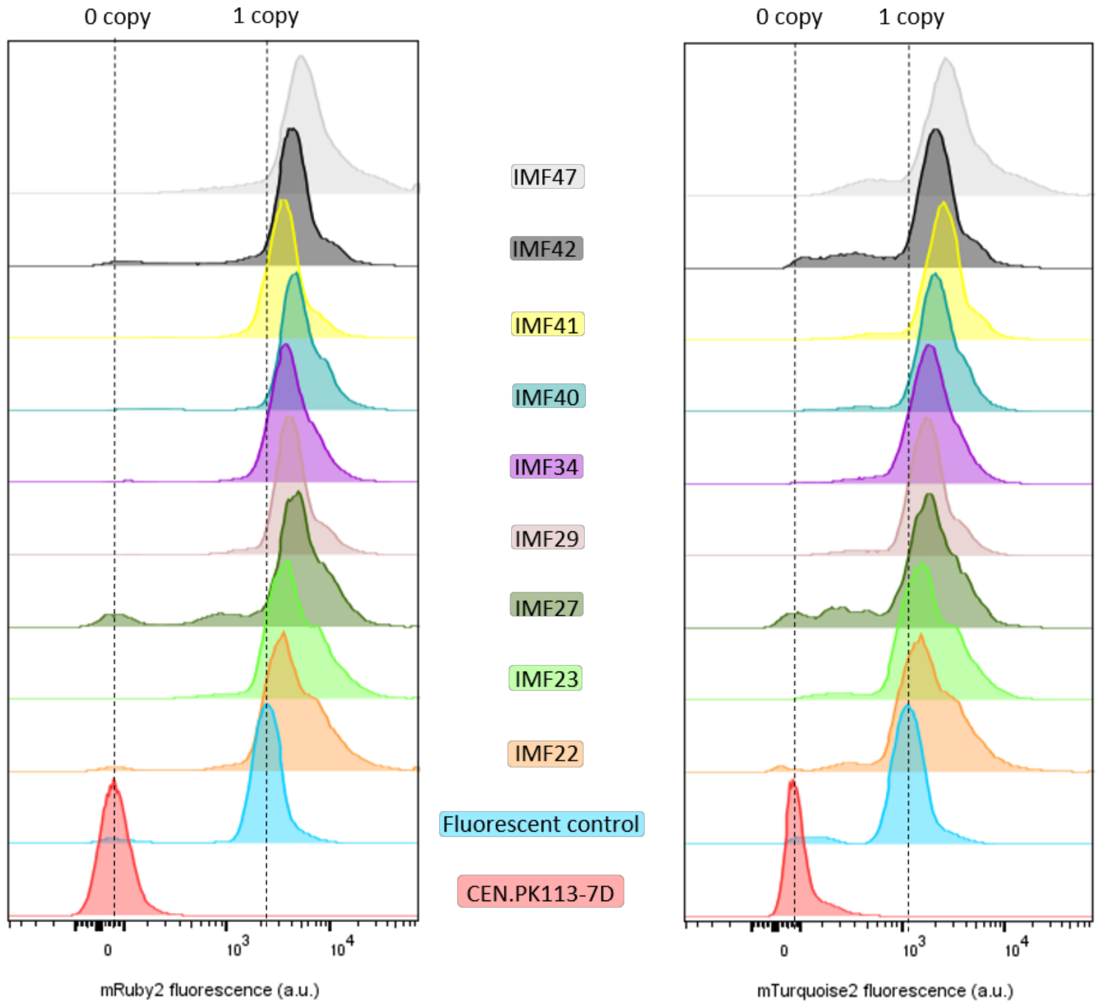

### Supplementary Figure 6 - NeoChr copy number estimation based on sequencing

IMF22 and IMF48 were analyzed by long-read Nanopore sequencing and IMF23, IMF41, IMF42 and IFM47 by short-read Miseq sequencing. Plots on the left represent the copy number of native chromosomes, while plots on the right show the NeoChrs copy number

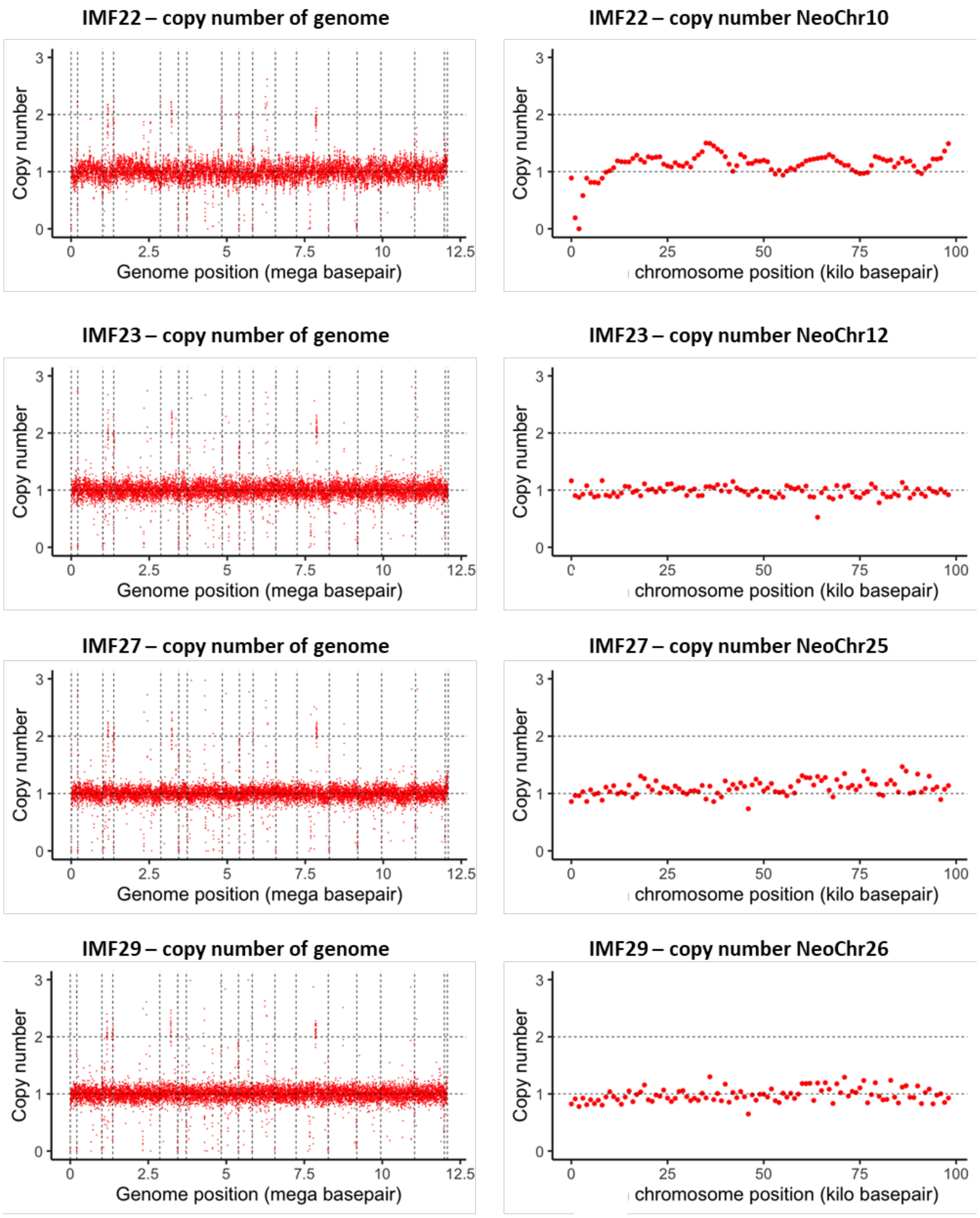

**IMF41 – copy number of genome**

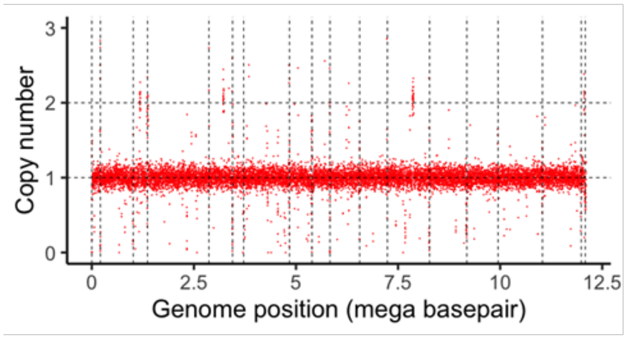


**IMF41 – copy number of NeoChr30**

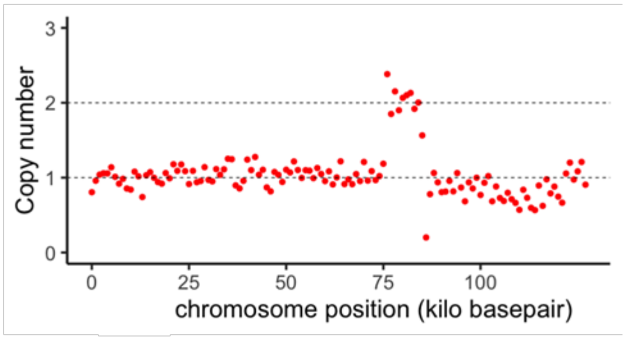


**IMF42 – copy number of genome**

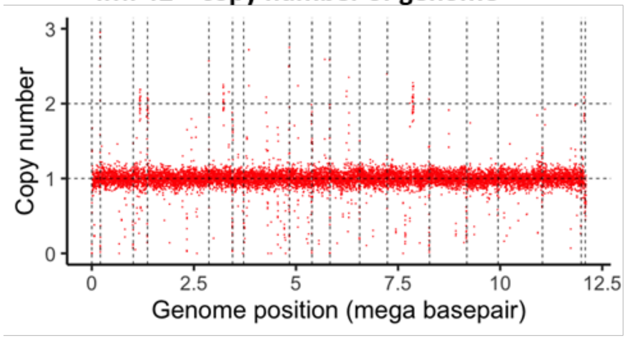


**IMF42 – copy number of NeoChr31**

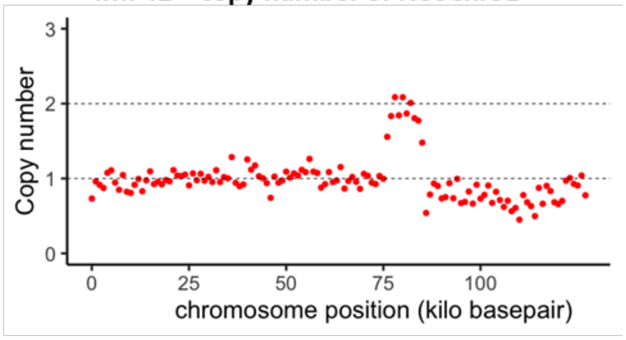


**IMF47 – copy number of genome**

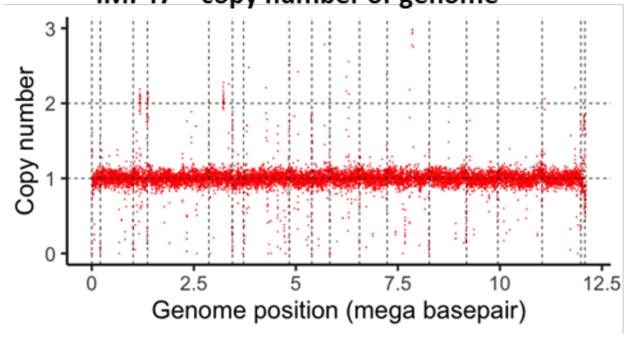


**IMF47 – copy number of NeoChr33**

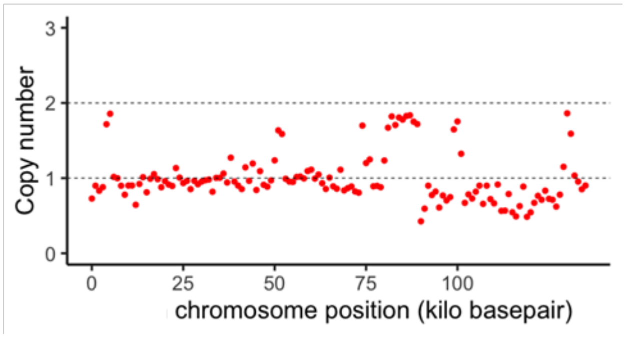


**IMF48 – copy number of genome**

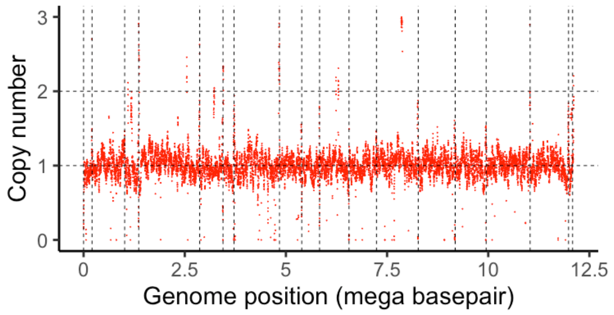


**IMF48 – copy number of NeoChr34**

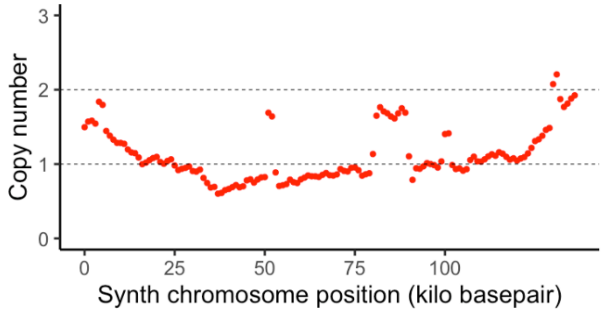

### Supplementary Figure 7 - Flow cytometric analysis of (linear) NeoChr25 and (circular) NeoChr26 designed for anthocyanin production

Cells from shake flask cultures were analyzed by FACS. The fluorescence is plotted on the y-axis and the FSC-A on the x-axis. Negative control: CEN.PK113-7D. Positive controls: IMX2224 (mRuby2), IMX2226 (mTurquoise2). Gates for fluorescence of the two different fluorescent proteins were drawn based on the IMX2224 and IMX2226 controls. Approximately 10000 or 100000 events are shown for each plot.

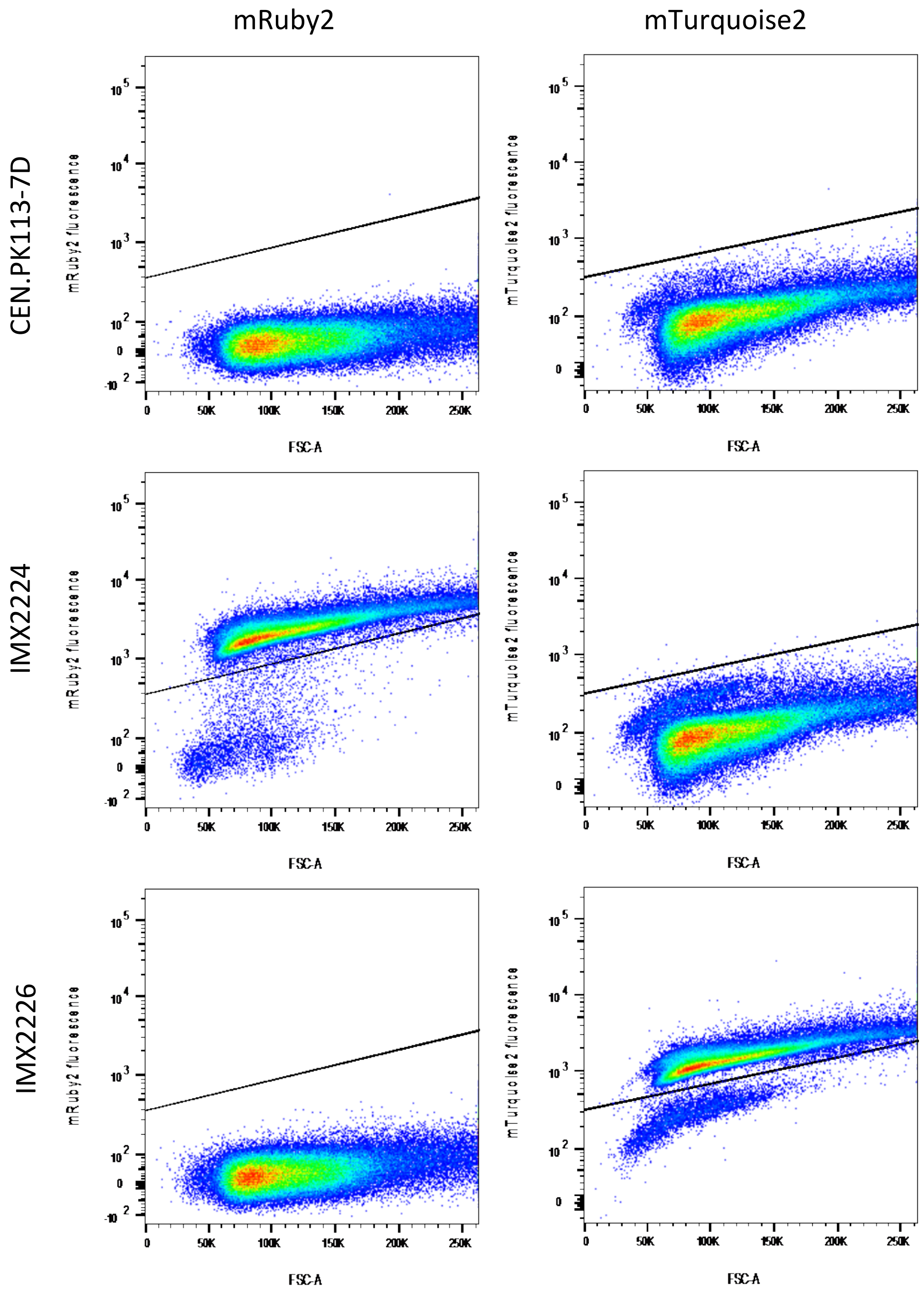

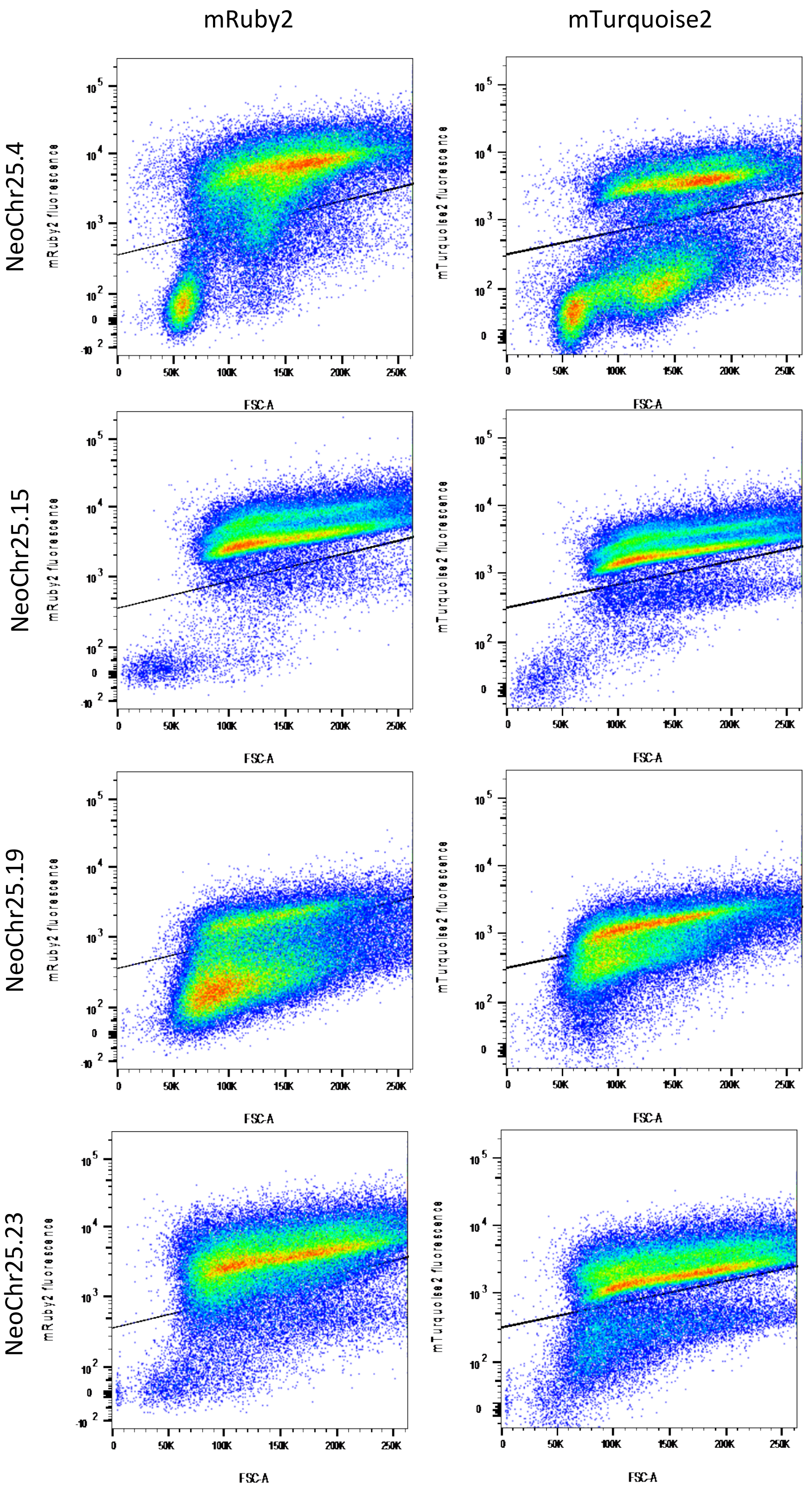
mRuby2
mTurquoise2
NeoChr25.4
NeoChr25.15
NeoChr25.19
NeoChr25.23
mRuby2 fluorescence
mTurquoise2 fluorescence
FSC-A

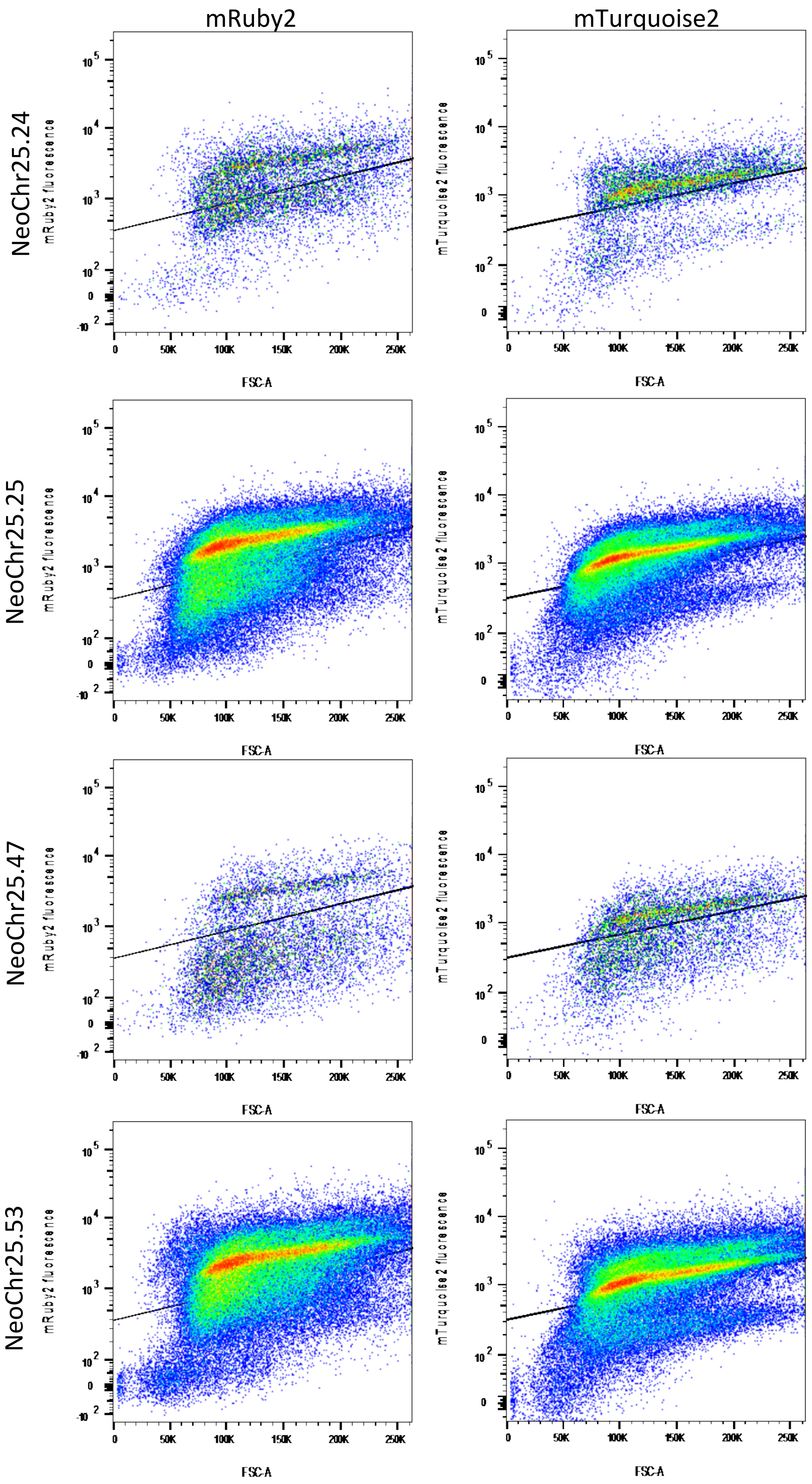
mRuby2
mTurquoise2
NeoChr25.24
NeoChr25.25
NeoChr25.47
NeoChr25.53
mRuby2 fluorescence
mTurquoise2 fluorescence
FSC-A

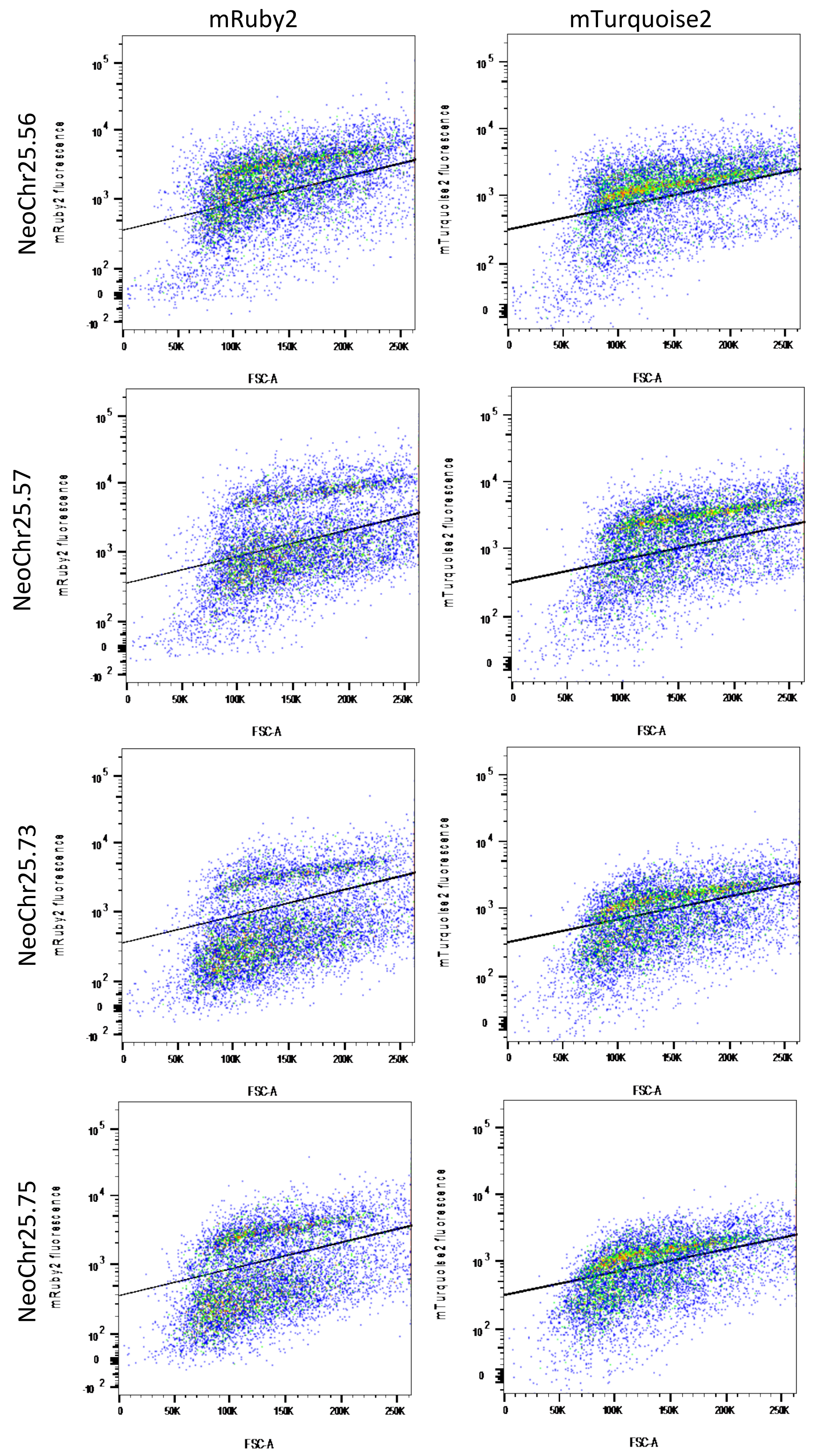
mRuby2
mTurquoise2
NeoChr25.56
NeoChr25.57
NeoChr25.73
NeoChr25.75
mRuby2 fluorescence
mTurquoise2 fluorescence
FSC-A
0
50K
100K
150K
200K
250K

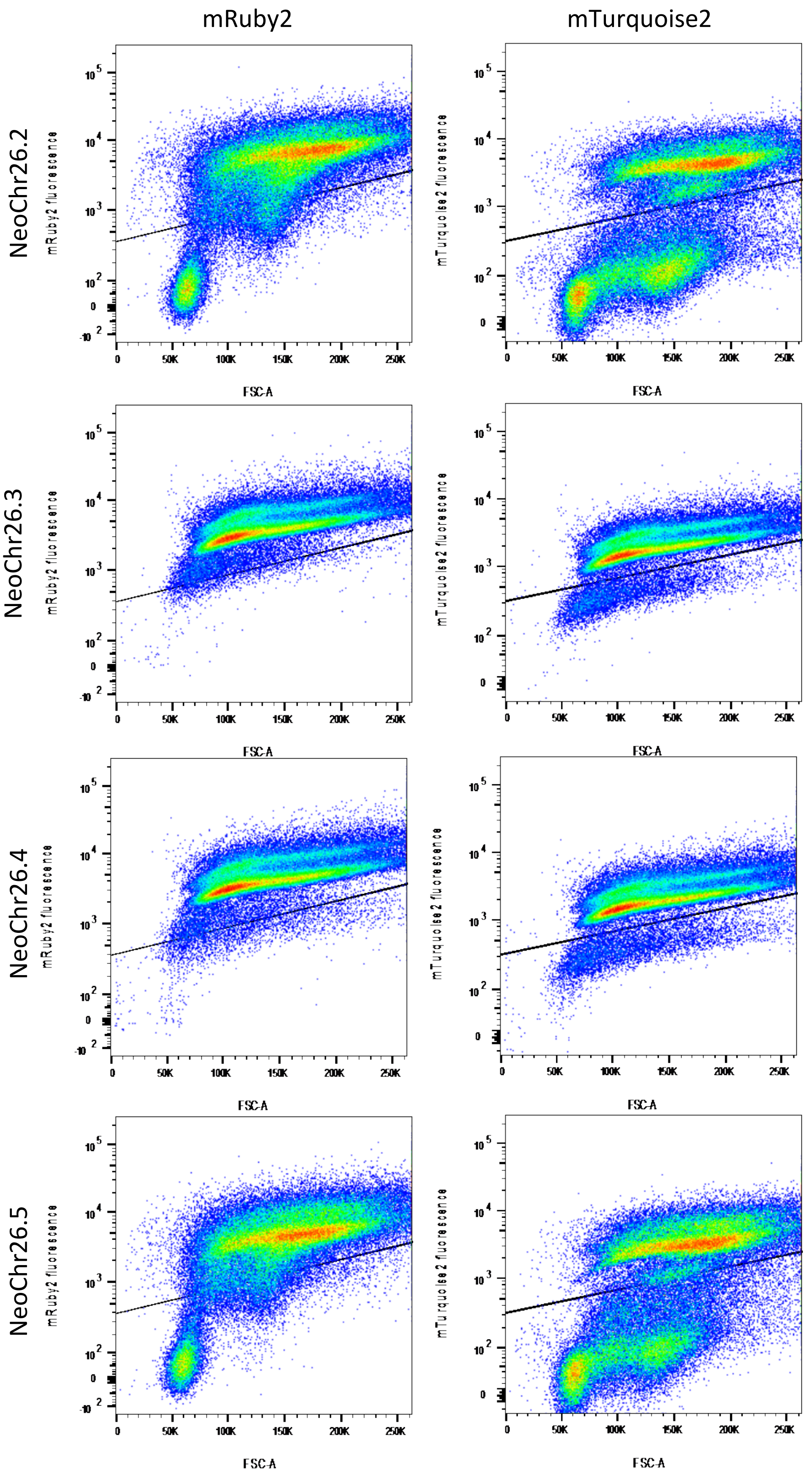
mRuby2
mTurquoise2
NeoChr26.2
NeoChr26.3
NeoChr26.4
NeoChr26.5
mRuby2 fluorescence
mTurquoise2 fluorescence
FSC-A

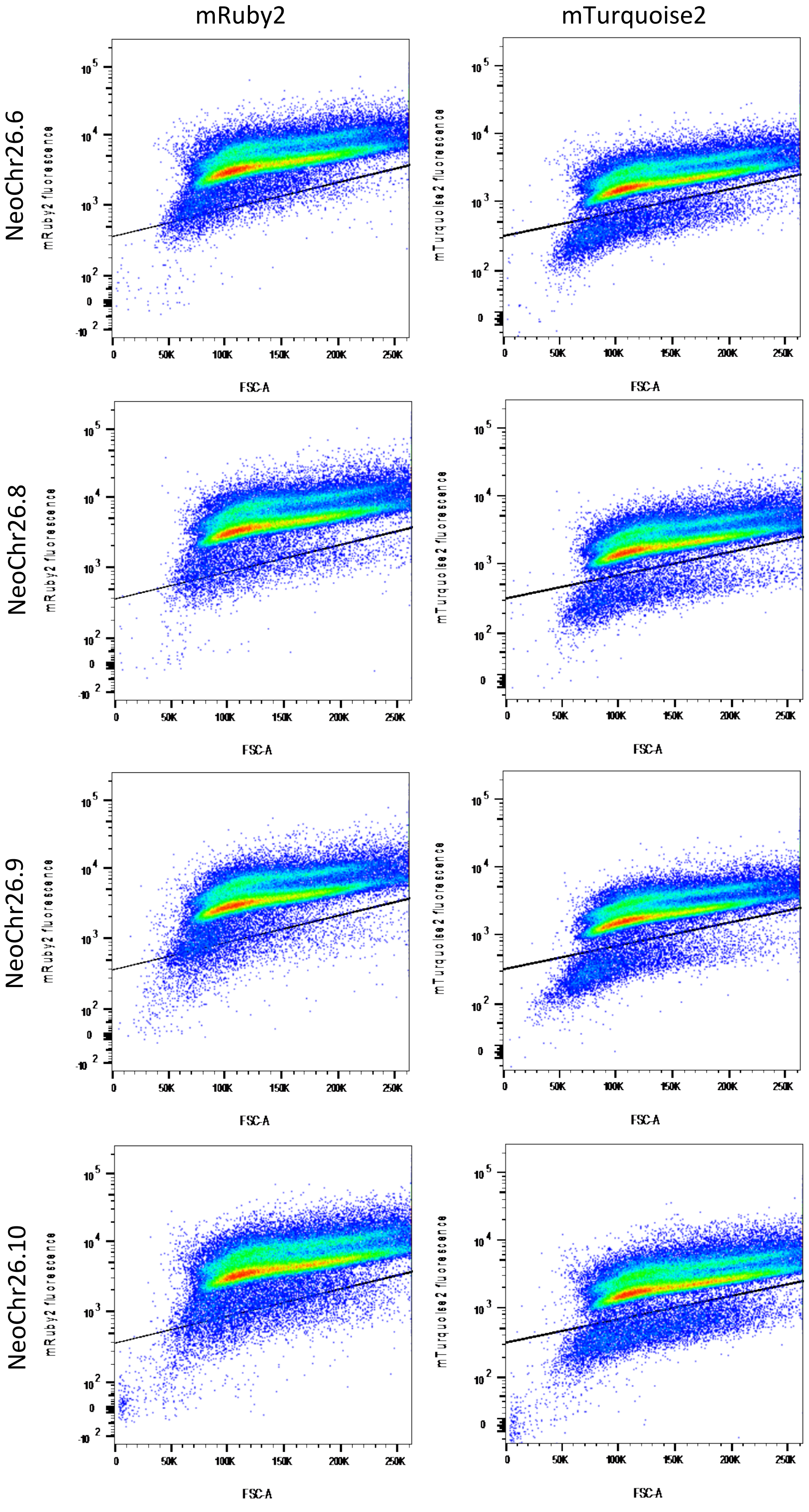
mRuby2
mTurquoise2
NeoChr26.6
NeoChr26.8
NeoChr26.9
NeoChr26.10
mRuby2 fluorescence
mTurquoise2 fluorescence
FSC-A
0
50K
100K
150K
200K
250K

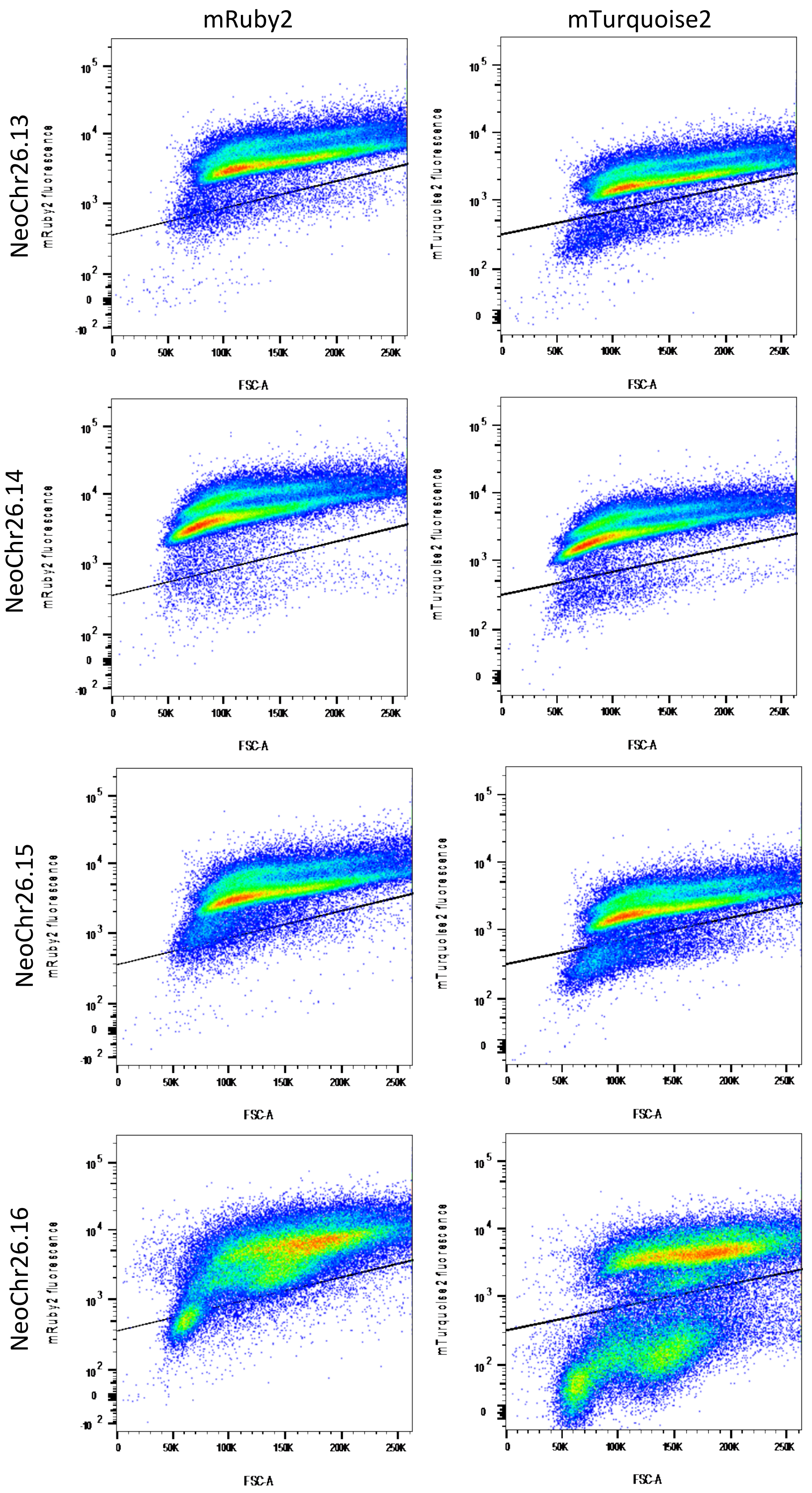

mRuby2
mTurquoise2
NeoChr26.13
NeoChr26.14
NeoChr26.15
NeoChr26.16
mRuby2 fluorescence
mTurquoise2 fluorescence
FSC-A

### Supplementary Figure 8 - Separation of (linear) NeoChr25 transformants on pulsed-field electrophoresis.

Pulsed-field electrophoresis was used to estimate the size of NeoChr25 in several yeast transformants. 1) Size ladder. 2) NeoChr25.4: correct size. 3) NeoChr25.15: no visible neochromosome. 4) NeoChr25.19: no visible neochromosome. 5) NeoChr25.23: no visible neochromosome. 6) NeoChr25.24: no visible neochromosome. 7) NeoChr25.25: correct size. 8) NeoChr25.47: correct size. 9) NeoChr25.53: correct size. 10) NeoChr25.56: correct size. 11) NeoChr25.57: no visible neochromosome. 12) NeoChr25.73: no visible neochromosome. 13) NeoChr25.75: correct size. 14) IMF22: positive control. 15) Size ladder.

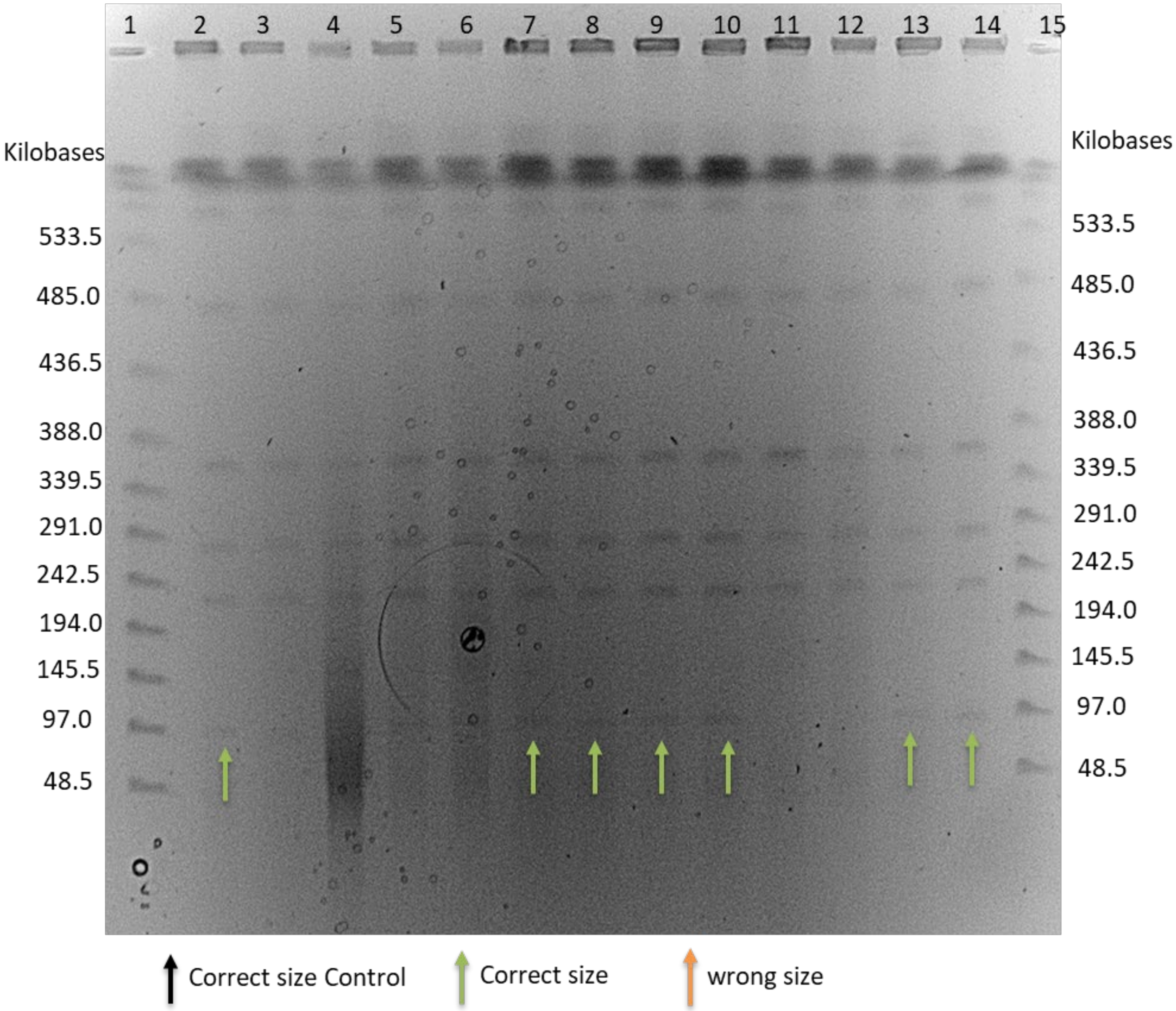

### Supplementary Figure 9 – Duplication and inversion of four plant genes in linear NeoChr25 and circular Neochr26.

An unexpected recombination was observed upon integration of the genes encoding the anthocyanin production pathway in the linear and circular NeoChrs. A) Schematic representation of the *in silico* design for the integration of the anthocyanin pathway in the circular NeoChr26 of IMF40 resulting in IMF41 and the linear NeoChr25 of IMF34 resulting in IMF42. B) Schematic representation of the genetic organization observed in IMF41 and IMF42. The last four genes in the anthocyanin pathway (*coAtF3H*, *coGhDFR*, *coAtANS* and *coAt3GT*) were duplicated and inversed, and *ARS106* was absent. The dashed boxes illustrate the recombination events that occurred on the left and right flank of this duplicated region. For the left flank, there was probably an exonuclease and subsequent Non-Homologous End Joining (NHEJ) event between the two SHR CJ, since there was no homology between the inverted and non-inverted sequences. In the sequenced IMF41 strain (circular) 57 bp of SHR CJ was retained and in the sequenced IMF42 (linear) 51 bp of SHR CJ was retained. For the right flank, in the IMF41 strain (circular) the first 649 bp showed exact homology to *pSeFBA1*, while the last 414 bp showed exact homology to *pScFBA1* (100% homology overlap of 7 bp). In the sequenced IMF42 strain (linear) the first 29 bp showed exact homology to *pSeFBA1* and the last 710 bp showed exact homology to *pSceFBA1* (overlap of 100% homology is 24 bp).

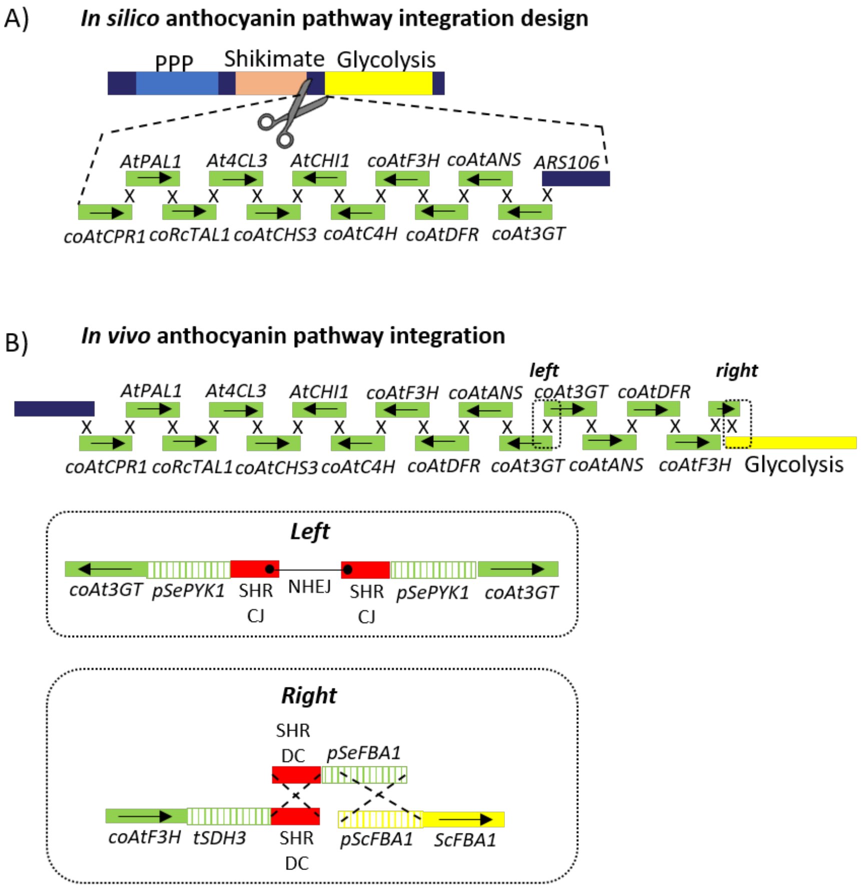

### Supplementary Figure 10 – schematic representation of *coAtANS* mutation in strains IMF41, IMF42, IMF44 and IMF47

**A)** The original coAtANS has a length of 1071 bp and encodes for an enzyme consisting of 356 amino acids. **B)** In strains IMF41, IMF42, IMF44 and IMF47, 21 nucleotides of non-homologous DNA (indicated in grey) together with 25 of the first 26 nucleotides of the *coAtANS* gene (indicated in dark green) were inserted right after the 26th nucleotide. **C)** This insertion resulted in a total insertion of 46 nucleotides disrupting the original ORF. However, this also resulted in a new ORF starting from the 284th nucleotide. **D)** The new ORF of the truncated coAtANS has a length of 834 bp and encodes for an enzyme consisting of 277 amino acids.

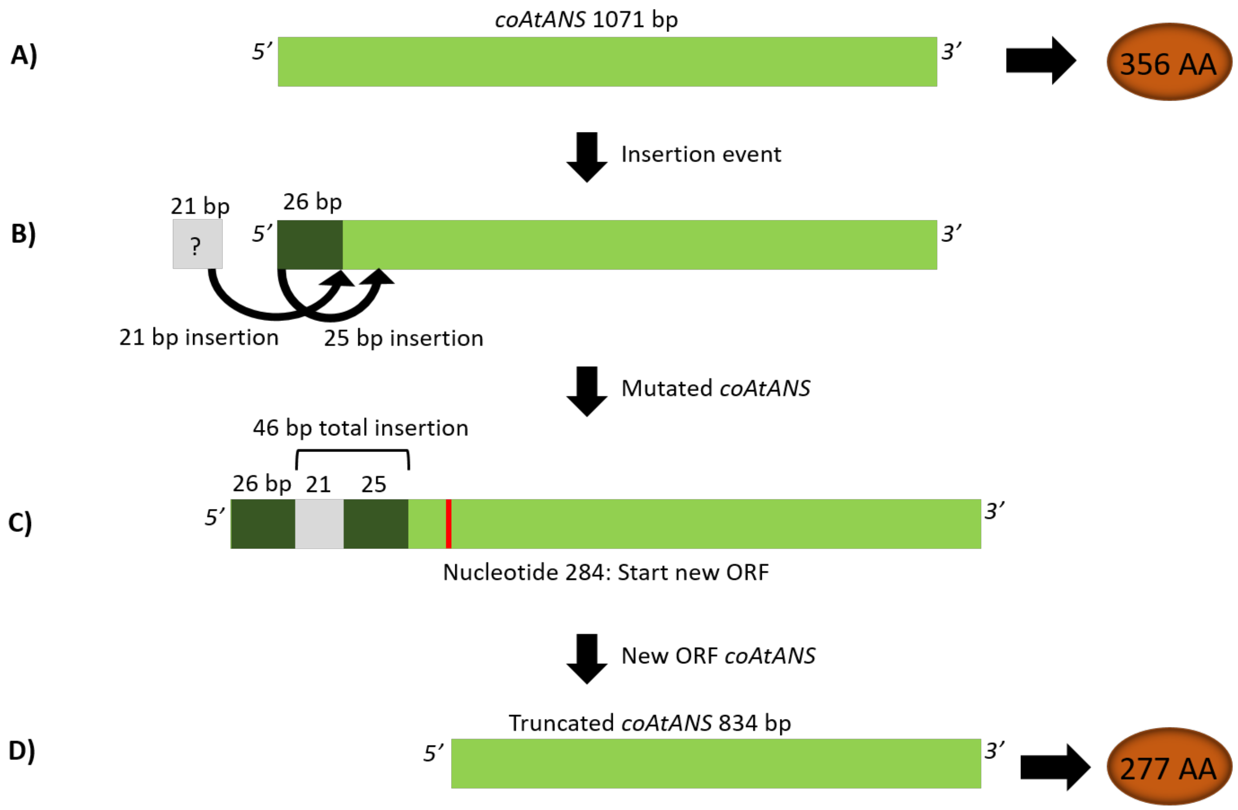

### Supplementary Figure 11 - Substrates and products profiles during aerobic batch cultivation in bioreactors of IMF41, IMF42 and IMF48.

**A)** IMF41 (Cir, 1x *coAtCHS3*), **B)** IMF42 (Lin, 1x *coAtCHS3*), and **C)** IMF48 (Lin, 9x *coAtCHS3, coAtANS*), were grown at 30°C in aerobic batch cultures in bioreactors, in chemically defined medium with 20 g $L^{-1}$ glucose as sole carbon source (SMD). Biological duplicates were performed and are shown in two columns as #1 and #2.

Row 1) ■ CDW ( g $L^{-1}$), ○ Glucose (mM), ● EtOH (mM), ▼ PYR (mM), △ Glyc (mM)
Row 2) ■ PPY (mM), ▲ COUM (mM), ● Phlor (mM), ▼ *p*OH2PE (mM)
Row 3) ■ DHK (mM), ▲ PEL (mM), ■ KEA (mM) ○ K3G (mM) ▼ P3G (µM)

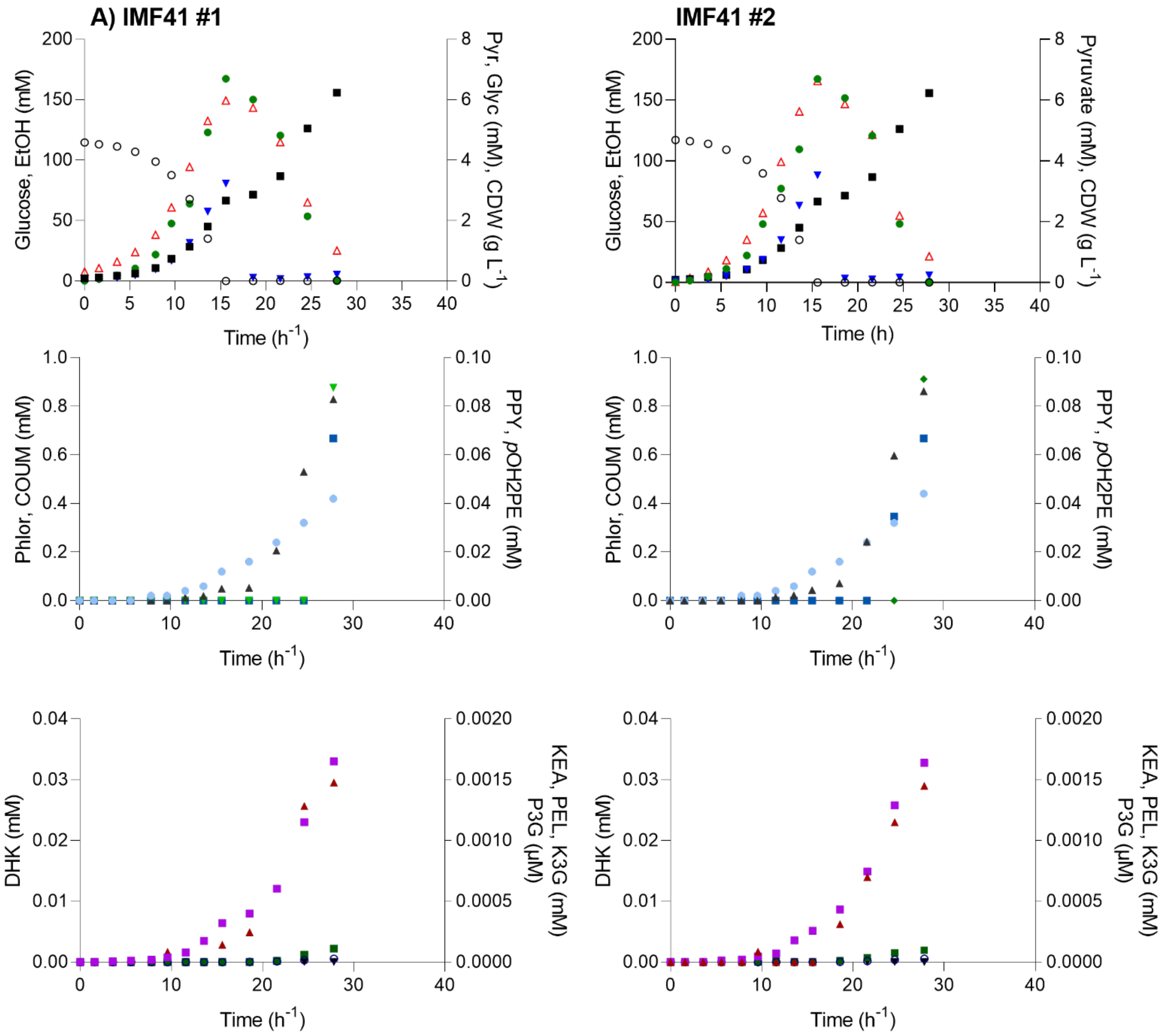

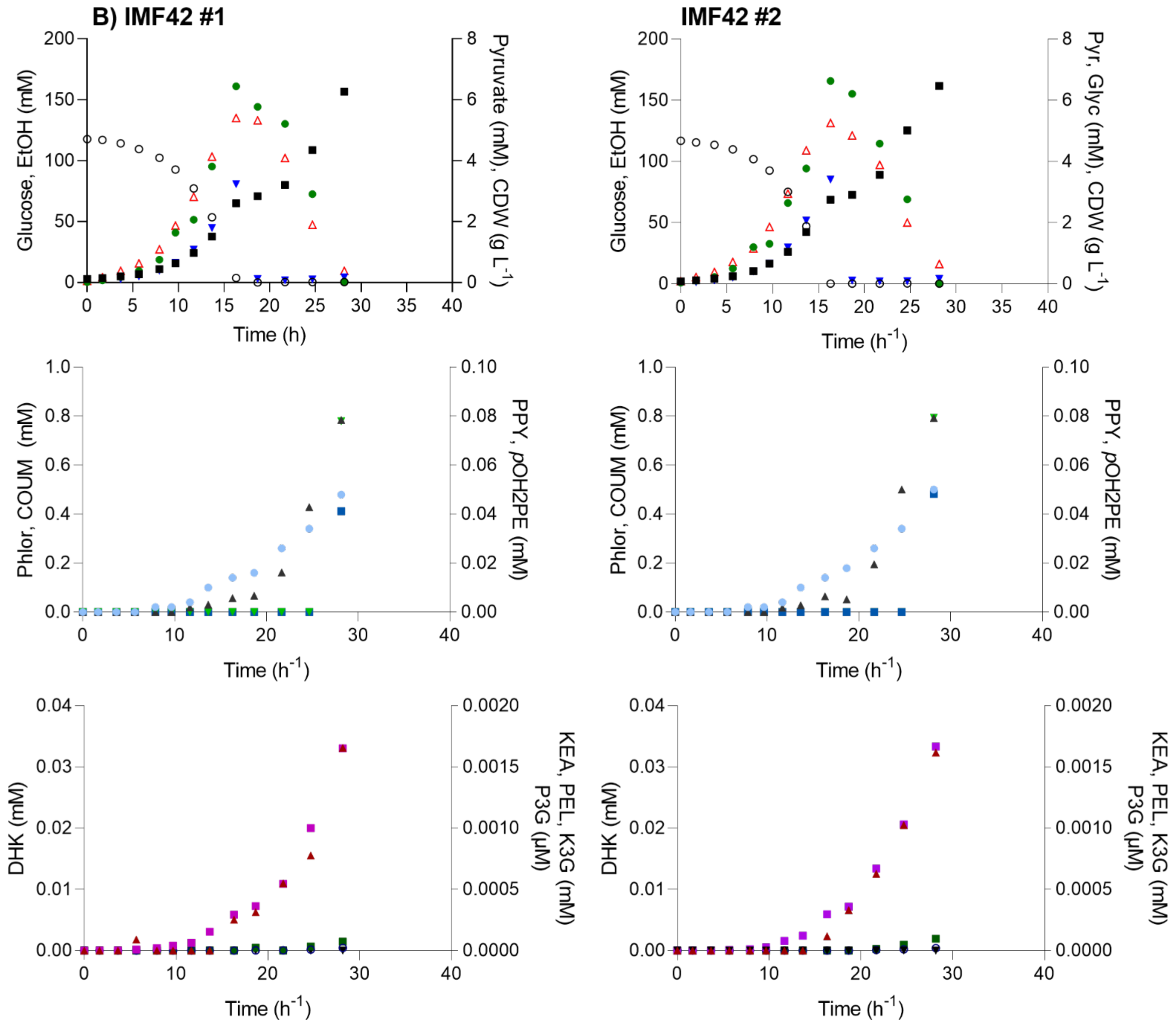

B) IMF42 #1
IMF42 #2
Glucose, EtOH (mM)
Pyruvate (mM), CDW (g L-1)
Pyr, Glyc (mM), CDW (g L-1)
Time (h)
Time (h-1)
Phlor, COUM (mM)
PPY, pOH2PE (mM)
DHK (mM)
KEA, PEL, K3G (mM)
P3G (µM)

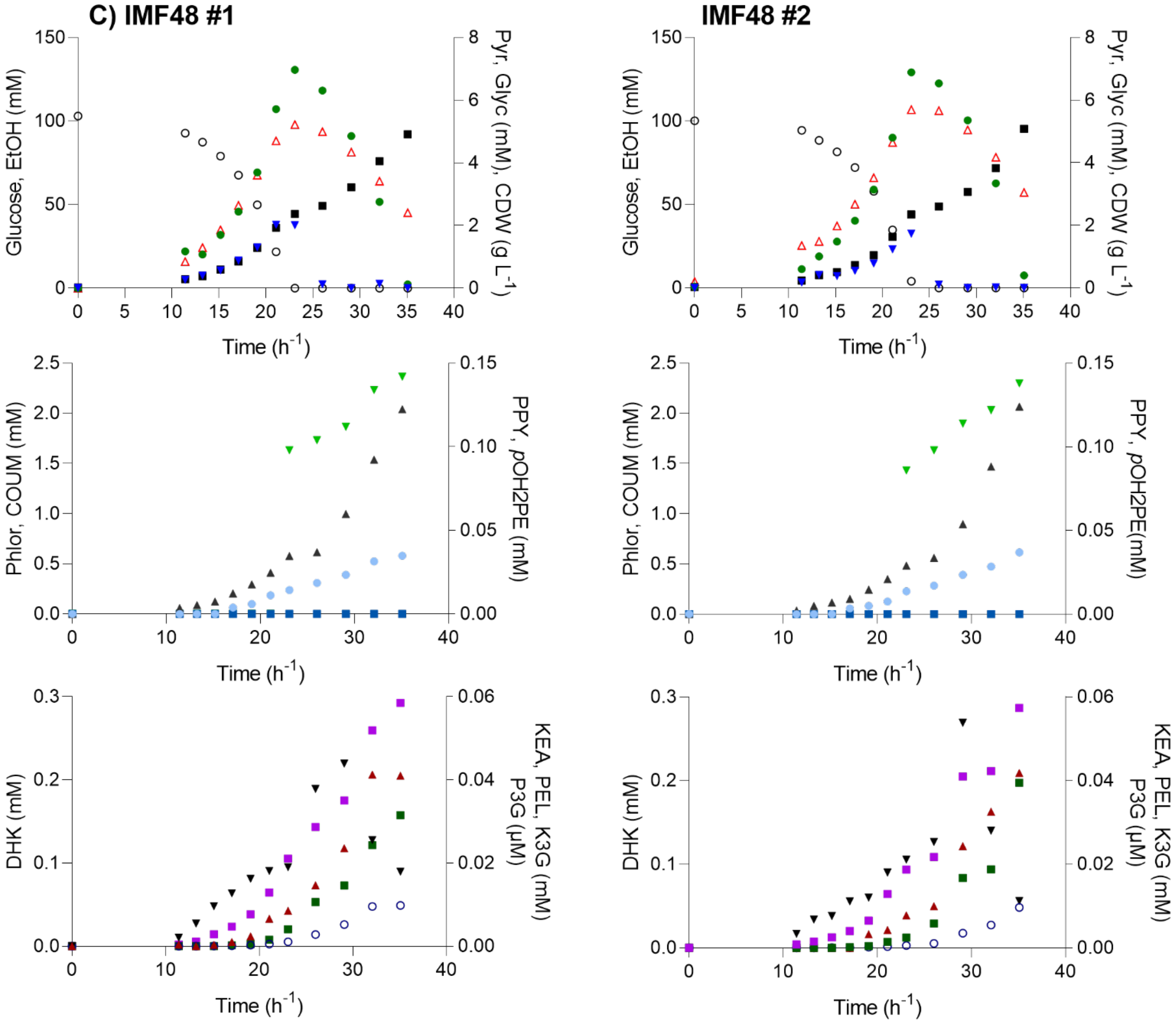

C) IMF48 #1
IMF48 #2
Glucose, EtOH (mM)
Pyr, Glyc (mM), CDW (g L-1)
Time (h-1)
Phlor, COUM (mM)
PPY, pOH2PE (mM)
DHK (mM)
KEA, PEL, K3G (mM)
P3G (µM)

### Supplementary Figure 12 – Detection and quantification of pelargonidin and pelargonidin 3-O-glucoside by LC-MS/MS

**A)** Extracted ion chromatogram for the pelargonidin 3-0-glucoside (P3G) mass peak with the composition $C_{21}H_{21}O_{10}^{+}$ and the m/z of 433.1. Data shown for the cell pellet extract of IMF48 duplicate #1 (Table 2), grown in aerobic bioreactor (sample, upper trace), for a blank injection (trace in the middle) analysed just before the sample and for a synthetic P3G standard shown in the lower trace (Pelargonidin 3-O-glucoside chloride, Sigma Aldrich, Cat No PHL89753).

**B)** The mass spectra show the accurate mass of P3G observed in the sample (upper mass spectrum) and the standard (lower mass spectrum). No corresponding P3G peak was observed for the blank injection (spectrum in the middle) analysed before the sample.

**C)** Extracted ion chromatogram of the pelargonidin (PEL) fragment with the composition $C_{15}H_{11}O_{5}$+, and a m/z of 271.06 Da. The corresponding fragment was observed in the sample (upper mass spectrum) and the standard (lower mass spectrum). No corresponding PEL fragment peak was observed for the blank injection analysed just before the sample. (Pelargonidin chloride, Sigma Aldrich, Cat No PHL80084).

**D)** The spectra show the accurate mass of the PEL major fragment with the composition $C_{15}H_{11}O_{5}^{+}$ and a m/z of 271.06 Da, as observed for the sample (upper spectrum) and the standard (lower spectrum). No corresponding fragment mass peak was observed for the blank injection (spectrum in the middle), which was performed just before the sample.

**E)** The table summarised the chemical compositions of P3G and the major fragment of pelargonidin (PEL) (loss of the sugar unit), the resulting theoretical m/z values, the sobered m/z values and the mass deviations (ppm). The observed mass deviations for standard and sample peaks were <5 ppm compared to their theoretical m/z values.

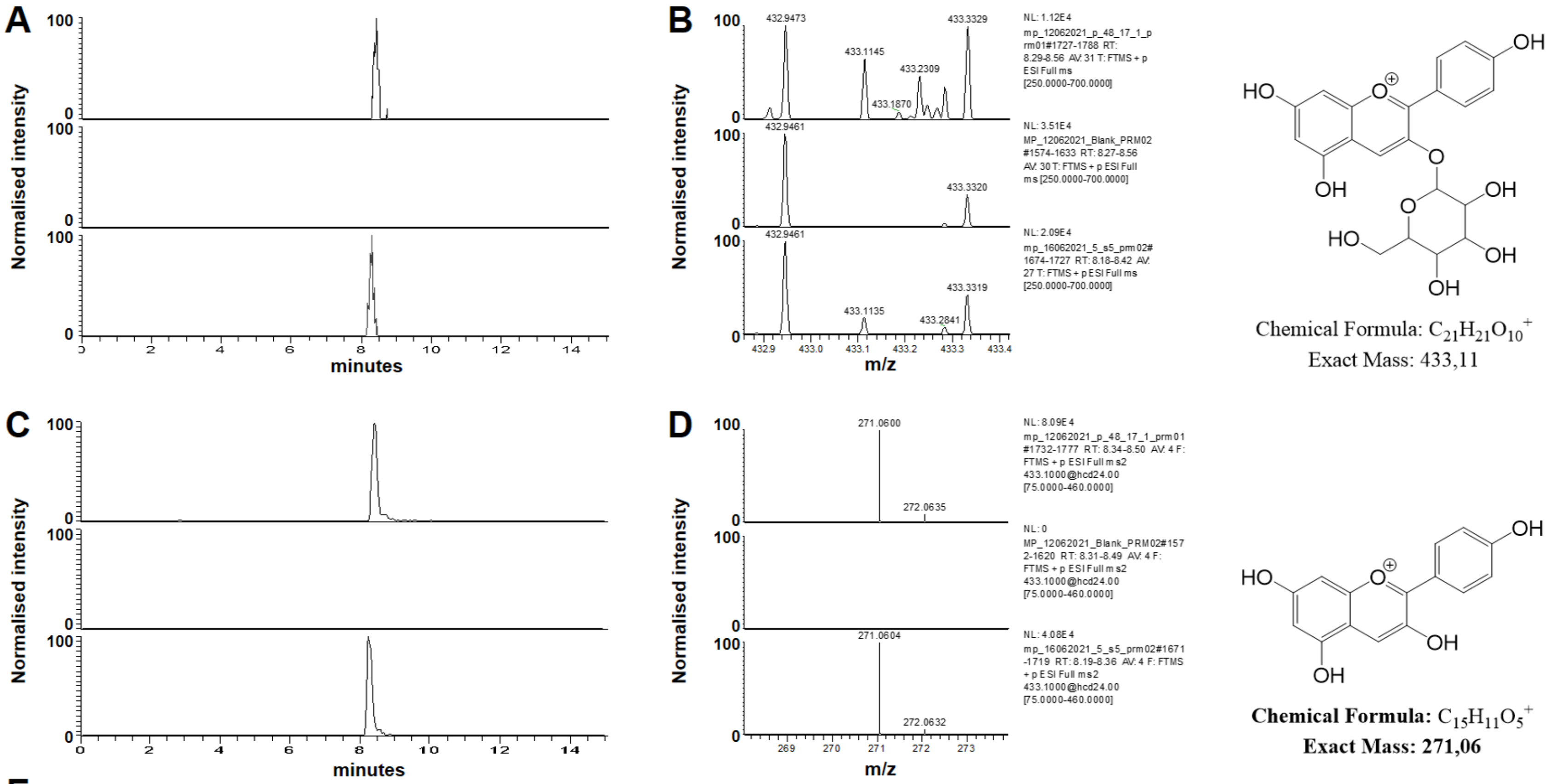


**E**

| COMPOUND | COMPOSITION | THEORETICAL M/Z | SAMPLE M/Z | Δppm | STANDARD M/Z | Δppm |
|---|---|---|---|---|---|---|
| P3G | $C_{21}H_{21}O_{10}^{+}$ | 433.1129 | 433.1145 | 3.69 | 433.1135 | 1.39 |
| PEL (fragment) | $C_{15}H_{11}O_{5}^{+}$ | 271.0601 | 271.0600 | -0.37 | 271.0604 | 1.11 |

### Supplementary Table 1 - Promoter-gene-terminator combinations in the NeoChrs.

Promoters, genes or terminators originate from *S. cerevisiae* unless indicated by: *Ec= Escherichia coli*, *At= Arabidopsis thaliana*, *Rc= Rhodobacter capsulatus*, *Gh= Gerbera hybrida Se= Saccharomyces eubayanus*, *Sk= Saccharomyces kudriavzevii*, *co= codon optimized.* Watermarked *S. cerevisiae* genes[1] are indicated with an *.

| Promoter | ORF | Terminator |
|---|---|---|
| **Genes from glycolysis and ethanolic fermentation are expressed from their native promoters and terminators** | | |
| *pFBA1* | *FBA1** | *tFBA1* |
| *pPGM1* | *PGM1** | *tPGM1* |
| *pHXK2* | *HXK2** | *tHXK2* |
| *pPDC1* | *PDC1** | *tPDC1* |
| *pPFK1* | *PFK1** | *tPFK1* |
| *pPFK2* | *PFK2** | *tPFK2* |
| *pPGK1* | *PGK1** | *tPGK1* |
| *pPYK1* | *PYK1** | *tPYK1* |
| *pTPI1* | *TPI1** | *tTPI1* |
| *pADH1* | *ADH1** | *tADH1* |
| *pTDH3* | *TDH3** | *tTDH3* |
| *pENO2* | *ENO2** | *tENO2* |
| *pPGI1* | *PGI1** | *tPGI1* |
| **Genes from the pentose phosphate pathway are expressed from their native promoters and terminators** | | |
| *pZWF1* | *ZWF1** | *tZWF1* |
| *pTKL1* | *TKL1** | *tTKL1* |
| *pGND1* | *GND1** | *tGND1* |
| *pRKI1* | *RKI1** | *tRKI1* |
| *pTAL1* | *TAL1** | *tTAL1* |
| *pRPE1* | *RPE1** | *tRPE1* |
| *pSOL3* | *SOL3** | *tSOL3* |
| **Auxotrophic markers are expressed from their native promoters and terminators** | | |
| *pHIS3* | *HIS3* | *tHIS3* |
| *pURA3* | *URA* | *tURA3* |
| **Fluorescent markers are expressed from *S. cerevisiae* promoters and terminators. Promoters identified from [2]** | | |
| *pCCW12* | *mRuby2* | *ENO1* |
| *pTEF2* | *mTurquoise2* | *tSSA1* |
| **Genes from the *E.coli* shikimate pathway are expressed from *S. cerevisiae* promoters and terminators. Promoters identified from [2, 3, 4]** | | |
| *pRPL3* | *coEcaroA* | *tSOL4* |
| *pRPL25* | *coEcaroD* | *tGPH1* |
| *pRPP0* | *coEcaroE* | *tCYC1* |
| *pHHF1* | *coEcaroG*$^{p150L}$ | *tTEF1* |
| *pHTB2* | *coEcaroL* | *tPGM2* |

| | | |
|---|---|---|
| *pRPL10* | *coEctyrA*$^{M53I\ A354V}$ | *tGDB1* |
| *pCWP2* | *coEctyrB* | *tGLC3* |
| *pHHF2* | *coEcaroB* | *tTEF2* |
| *pRPL8A* | *coEcaroC* | *tGPD2* |
| *pRPL18B* | *coEcpheA*$^{T326P}$ | *tGSY2* |
| **One gene from the anthocyanin pathway is expressed from a *S. cerevisiae* promoter and terminator. Promoter identified from [2, 3, 4]** | | |
| *pTEF1* | *coAtCHS3* | *tMDH1* |
| **Genes from the anthocyanin pathway are expressed from a *S. eubayanus* and *S. kudriavzevii* promoters [5] and *S. cerevisiae* terminators.** | | |
| *pSePDC1* | *AtPAL1* | *tLAT1* |
| *pSeGPM1* | *coRcTAL1* | *tCIT1* |
| *pSkADH1* | *AtCHI1* | *tSDH4* |
| *pSeFBA1* | *coAtC4H* | *tADH3* |
| *pSkTDH3* | *coAtF3H* | *tSDH3* |
| *pSePGK1* | *coGhDFR* | *tACO1* |
| *pSeENO2* | *coAtANS* | *tFUM1* |
| *pSePYK1* | *coAt3GT* | *tDIC* |

### Supplementary Table 2 - Sequence fidelity of NeoChrs

Mutation identified in the neochromosomes as compared to the *in silico* design and with the most relevant parental strain. The * indicates mutations which are the same in two separate transformations and therefore probably resulting from the template DNA and not during the *in vivo* assembly. Non-synonymous mutations are indicated in bold.

| Position | Region | Mutation type |
|---|---|---|
| **NeoChr25 (IMF27)** | | |
| 8648 | *pTKL1* | C to CT |
| 14466 | SHR BQ | C to CT |
| 20137* | *pTEF2 (mTurquoise 2)** | CAT to C* |
| 26993 | *pHHF2 (EcAroB)* | AT to A |
| 46676* | *pCWP2 (EcTyrB)** | A to G* |
| 52732 | SHR AE | G to GT |
| 66608 | SHR N | CA to C |
| 73854* | *tENO2** | C to A* |
| 86809 | *pPFK2* | GA to G |
| 90753 | SHR M | GC to G |
| 90762 | SHR M | AT to A |
| **NeoChr26 (IMF29)** | | |
| 14306 | *tGND1* | CT to C |
| **15181** | ***RKI1*** | **C to A (Glu-129-Gln)** |
| 20137* | *pTEF2 (mTurquoise2)** | CAT to C* |
| 22220 | SHR DF | TC to T |
| 22223 | SHR DF | TC to T |
| 46676* | *pCWP2 (EcTyrB)** | A to G* |
| 57795 | SHR DL | A to AG |
| 64374 | SHR Q | T to TG |
| 66632 | *pPYK1* | CT to C |
| 73854* | *tENO2** | C to A* |
| 73864 | *tENO2* | GT to G |
| 73925 | SHR B | T to C |
| 73926 | SHR B | A to T |
| 73928 | SHR B | G to A |
| 78556 | *pPGI1* | C to A |
| 88608 | *pHIS3* | GA to G |
| 90398 | *pGPM1* | C to CTA |
| **NeoChr30 (IMF41) as compared to NeoChr26 (IMF29)** | | |
| 8648 | *pTKL1* | C to CT |
| 35602 | *tPGM2 (coEcAroL)* | CT to C (In T stretch) |
| 67393 | *pSeTPI1 (At4CL3)* | G to GT (In T stretch) |
| 71689 | *tMDH1 (coAtCHS3)* | GA to G (In A stretch) |
| 73063* | *pSkADH1 (AtCHI1)** | AT to A* |
| 73209* | *pSkADH1 (AtCHI1)** | CT to C (In T stretch)* |
| 81107 | *tFUM1 (coAtANS)* | CG to C |
| **82459*** | ***coAtANS**** | **Insertion of 46 bp*** |

| | | |
|---|---|---|
| 115179 | *pPFK2* | G to GAA (In A stretch) |
| **NeoChr31 (IMF42) as compared to NeoChr25 (IMF27)** | | |
| 57432 | Chunk 16AB | A to AC |
| 64168 | *tLAT1 (AtPAL1)* | TAA to T (In A stretch) |
| 67393 | *pSeTPI1 (At4CL3)* | G to GT (In T stretch) |
| **67753** | ***At4CL3*** | **A to G (Thr-15-Ala)** |
| 70927 | *coAtCHS3* | G to A (Leu-155-Leu) |
| 70930 | *coAtCHS3* | A to G (Arg-156-Arg) |
| 73063* | *pSkADH1 (AtCHI1)** | AT to A* |
| 73209* | *pSkADH1 (AtCHI1)** | CT to C (In T stretch)* |
| **82459*** | ***coAtANS**** | **Insertion of 46 bp*** |
| **NeoChr33 (IMF47) as compared to NeoChr31 (IMF42)** | | |
| 48856 | *pCWP2 (coEcTyrB)* | A to G |
| 50730 | *tMDH1 (coAtCHS3)* | A to AT (In T stretch) |
| 85431 | SHR EB | AT to A |
| 85439 | SHR EB | TG to T |
| 85458 | SHR EB | GA to G |
| 96995 | *tPGK1* | T to A |

### Supplementary Table 3 - Amino acid substitution in native genome of NeoChr strains

Amino acid substitutions identified in the genome of the constructed strains as compared to most relevant parental strain.

| Systematic name | Name | Type | Amino acid change |
|---|---|---|---|
| **IMF27 compared to IMX589** | | | |
| YPL283W-A | - | Intron | - |
| YNL327W | *EGT2* | synonymous | Tyr-583-Tyr |
| YNL327W | *EGT2* | Non-synonymous | Thr-586-Ser |
| YNL161W | *CBK1* | Non-synonymous | Ser-711-Ala |
| **IMF29 compared to IMX589** | | | |
| YPL283W-A | - | intron | - |
| YPL283W-A | - | intron | - |
| YPL283W-A | - | intron | - |
| YPL283W-A | - | Non-synonymous | Gly-132-Ser |
| YMR160W | - | Non-synonymous | Gln-11-Arg |
| YNL327W | *EGT2* | Synonymous | Tyr-583-Tyr |
| YNL327W | *EGT2* | Non-synonymous | Thr-586-Ser |
| YNL161W | *CBK1* | Non-synonymous | Ser-711-Ala |
| **IMF41 compared to IMF29** | | | |
| YCR089W | FIG2 | Non-synonymous | Thr-1017-Arg |
| YCR089W | FIG2 | Non-synonymous | Ala-1020-Ser |
| YDR224C | HTB1 | Synonymous | Ala-121-Ala |
| YIL137C | *TMA108* | Non-synonymous | Ser-742-Leu |
| **IMF42 compared to IMF27** | | | |
| YCR089W | FIG2 | Non-synonymous | Thr-1017-Arg |
| YCR089W | FIG2 | Non-synonymous | Ala-1020-Ser |
| YBL113C | - | Non-synonymous | His-252-Asn |
| **IMF47 compared to IMF42** | | | |
| YEL075W-A | - | intron | - |
| YHR016C | *YSC84* | intron | - |
| YJR143C | *PMT4* | Non-synonymous | Met-1-Ile |

### Supplementary Table 4 - Extracellular concentration of aromatic compounds produced by engineered *S. cerevisiae* strains in shake flask cultures

Determination of the intermediates of the anthocyanin pathway in *S. cerevisiae* strains IMF41 (Cir NeoChr, 1X *coAtCHS3*), IMF42 (Lin NeoChr, 1X *coAtCHS3*), IMF47 (Lin NeoChr, 9X *coAtCHS3*) and IMF48 (Lin NeoChr, 9X *coAtCHS3* repaired *coAtANS*), grown in aerobic shake flask batch cultures on glucose (20 g $L^{-1}$) and urea. The data represents the average ± mean deviation of independent biological triplicates. Intermediates of the anthocyanin pathway coumaroyl-CoA, naringenin-chalcone, and leucopelargonidin were not measured. * Indicates statistical significance when comparing IMF47 or IMF48 to IMF42, and # when comparing IMF48 to IMF47 (Student *t*-test, two-tailed, homoscedastic, *p*-value threshold 0.05).

| **(mM)** | IMF41 | IMF42 | IMF47 | IMF48 |
|---|---|---|---|---|
| Phenylpyrurate | 2.00E-02 ± 0.00E+00 | 4.33E-02 ± 4.04E-02 | BD[a] | BD[a] |
| 2-Phenylethanol | 8.67E-02 ± 1.15E-02 | 8.67E-02 ± 2.52E-02 | 3.23E-01 ± 3.51E-02* | 1.97E-01 ± 2.62E-03*# |
| *p*-Hydroxyphenylethanol | 2.33E-02 ±2.08E-02 | 3.33E-02 ±5.77E-03 | 1.10E-01 ±1.73E-02* | BD[a] |
| Cinnamic acid | 3.80E-01 ±2.00E-02 | 1.60E-01 ± 2.00E-02 | 0.00E+00 ± 0.00E+00* | 1.46E-01 ± 3.06E-03# |
| Coumaric acid | 4.27E-01 ± 1.15E-02 | 5.40E-01 ± 2.00E-02 | 7.13E-01 ± 1.15E-02* | 7.64E-01 ± 6.24E-03*# |
| Phloretic acid | 7.25E-01 ± 1.21E-02 | 7.18E-01 ± 8.72E-03 | 5.09E-01 ± 1.15E-03* | 1.07E+00 ± 7.66E-03*# |
| Naringenin | BD[a] | BD[a] | BD[a] | BD[a] |
| Dihydrokaempferol | 2.25E-02 ± 2.57E-03 | 2.84E-02 ± 1.27E-03 | 9.49E-02 ± 4.29E-03* | 3.83E-01 ± 8.34E-02*# |
| Kaempferol | 6.33E-04 ± 3.44E-05 | 6.96E-04 ± 3.46E-05 | 2.14E-03 ± 3.06E-04* | 1.39E-02 ± 5.72E-03*# |
| Pelargonidin | 2.45E-03 ± 1.60E-04 | 1.81E-03 ± 1.98E-04 | 5.76E-03 ± 4.11E-04* | 3.65E-02 ± 7.64E-03*# |
| Kaempferol 3-O-glucoside | 2.24E-04 ± 1.83E-05 | 2.24E-04 ± 5.98E-06 | 4.39E-04 ± 2.41E-05* | 5.53E-03 ± 1.13E-03*# |
| Pelargonidin 3-O-glucoside | BD[a] | BD[a] | BD[a] | 2.14E-05 ± 3.93E-06*# |
| Total aromatics before CHS | 1.66 ± 0.06 | 1.58 ± 0.06 | 1.66 ± 0.05 | 1.98 ± 0.01*# |
| Total anthocyanins (after CHS) | 0.03 ± 0.00 | 0.03 ± 0.00 | 0.1 ± 0.0* | 0.45 ± 0.10*# |
| Total aromatics | 1.69 ± 0.06 | 1.61 ± 0.08 | 1.76 ± 0.05 | 2.43 ± 0.11*# |

[a]BD: below detection

### Supplementary Table 5 - Physiological characterization of anthocyanin-producing strains grown in bioreactors

A) The specific growth rate (μ) and the yield (Y) of biomass (X) and ethanol (ETOH) on glucose (S)

B) The overall yield (Y) of glycerol (GLYC), pyruvate (PYR), coumaric acid (COUM), phloretic acid (PHLOR) and dihydrokaempferol (DHK) on glucose and ethanol (S) during aerobic bioreactor batch cultivation of IMF41 (Cir NeoChr, 1x *coAtCHS3*), IMF42 (Lin NeoChr, 1x *coAtCHS3*),) and IMF48 (Lin NeoChr, 9x *coAtCHS3*, repaired *coAtANS*).

| **A)** | $^a\mu_{MAX}$ $h^{-1}$ | $^aY_{X/S}$ ($g\ g^{-1}$) | $^aY_{ETOH/S}$ ($mol\ mol^{-1}$) |
|---|---|---|---|
| **IMF41** (Cir) | 0.23 ± 0.00 | 0.12 ± 0.00 | 1.44 ± 0.05 |
| **IMF42** (Lin) | 0.22 ± 0.01 | 0.12 ± 0.00 | 1.41 ± 0.02 |
| **IMF48** (Lin, 9x *coAtCHS3, coAtANS*) | 0.20 ± 0.01 | 0.13 ± 0.00 | 1.29 ± 0.05 |

| **B)** | $Y_{GLYC/S}$ ($mol\ mol^{-1}$) | $Y_{PYR/S}$ ($mol\ mol^{-1}$) | $Y_{X/S}$ ($mol\ mol^{-1}$) | $Y_{coum/S}$ ($\mu mol\ mol^{-1}$) | $Y_{PHLOR/S}$ ($\mu mol\ mol^{-1}$) | $Y_{DHK/S}$ ($\mu mol\ mol^{-1}$) |
|---|---|---|---|---|---|---|
| **IMF41** (Cir) | 0.056 ± 0.005 | 0.029 ± 0.001 | 0.29 ± 0.01 | 7.30 ± 0.08 | 4.71 ± 0.20 | 0.28 ± 0.01 |
| **IMF42** (Lin) | 0.048 ± 0.002 | 0.029 ± 0.001 | 0.30 ± 0.01 | 6.70 ± 0.12 | 5.02 ± 0.15 | 0.28 ± 0.00 |
| **IMF48** (Lin, 9x *coAtCHS3, coAtANS*) | 0.052 ± 0.003 | 0.019 ± 0.003 | 0.27 ± 0.01 | 20.2 ± 0.59 | 5.89 ± 0.35 | 2.85 ± 0.02 |

[a] Determined for the glucose phase only

### Supplementary Table 6 - *S. cerevisiae* strains used in this study

Strains that were short-read or long-read sequenced in this study are marked with a *. SHRs are differently annotated than in Kuijpers *et al.* [6]. SHRs are annotated in bold subscript between de genetic fragments that they join together.

| Strain | Relevant Genotype | Source |
|---|---|---|
| CEN.PK113-7D | *MATa URA3 HIS3 LEU2 TRP1 MAL2-8c SUC2* | Entian and Kötter [7] |
| IMC111 | *MATa ura3-52 can1Δ::cas9-natNT2 TRP1 LEU2 HIS3* pUDC191 (mRuby2) | Postma, Dashko [8] |
| IMC112 | *MATa ura3-52 can1Δ::cas9-natNT2 TRP1 LEU2 HIS3* pUDC192 (mTurquoise2) | Postma, Dashko [8] |
| IMX589 | *MATa ura3-52 his3-1 leu2-3,112 MAL2-8c SUC2 glk1Δ:: (pAgTEF1-SpHIS5-tAgTEF1) hxk1Δ::KlLEU2 tdh1Δ tdh2Δ gpm2Δ gpm3Δ eno1Δ pyk2Δ pdc5Δ pdc6Δ adh2Δ adh5Δ adh4Δ sga1Δ::(* $_{\mathbf{G}}$ *tFBA1-FBA1-pFBA1* $_{\mathbf{H}}$ *pTPI1-TPI1-tTPI1* $_{\mathbf{P}}$ *tPGK1-PGK1-pPGK1* $_{\mathbf{Q}}$ *tADH1-ADH1-pADH1* $_{\mathbf{N}}$ *pPYK1-PYK1-tPYK1* $_{\mathbf{O}}$ *tTDH3-TDH3-pTDH3* $_{\mathbf{A}}$ *pENO2-ENO2-tENO2* $_{\mathbf{B}}$ *pHXK2-HXK2-tHXK2* $_{\mathbf{C}}$ *pPGI-PGI1-tPGI1 D pPFK1-PFK1-tPFK1* $_{\mathbf{J}}$ *tPFK2-PFK2-pPFK2* $_{\mathbf{K}}$ *pAgTEF1-AmdSYM-tAgTEF1* $_{\mathbf{L}}$ *tGPM1-GPM1-pPGM1* $_{\mathbf{M}}$ *pPDC1-PDC1-tPDC1-SYN* $_{\mathbf{F}}$ *) pyk1Δ pgi1Δ tpi1Δ tdh3Δ pfk2Δ::(pTEF1-Spcas9-tCYC1 natNT1) pgk1Δ gpm1Δ fba1Δ hxk2Δ pfk1Δ adh1Δ pdc1Δ eno2Δ* | Kuijpers, Solis-Escalante [6] |
| IMX1338 | *MATa ura3-52 his3-1 leu2-3,112 MAL2-8c SUC2 glk1Δ::(pAgTEF1-SpHIS5-tAgTEF1)Δ::(pGAL1-I SceI-tCYC1) hxk1Δ::KlLEU2 tdh1Δ tdh2Δ gpm2Δ gpm3Δ eno1Δ pyk2Δ pdc5Δ pdc6Δ adh2Δ adh5Δ adh4Δ sga1Δ::(* $_{\mathbf{G}}$ *tFBA1-FBA1-pFBA1* $_{\mathbf{H}}$ *pTPI1-TPI1-tTPI1 P tPGK1-PGK1-pPGK1* $_{\mathbf{Q}}$ *tADH1-ADH1-pADH1* $_{\mathbf{N}}$ *pPYK1-PYK1-tPYK1* $_{\mathbf{O}}$ *tTDH3-TDH3-pTDH3* $_{\mathbf{A}}$ *pENO2-ENO2-tENO2* $_{\mathbf{B}}$ *pHXK2-HXK2-tHXK2* $_{\mathbf{C}}$ *pPGI-PGI1-tPGI1* $_{\mathbf{D}}$ *pPFK1-PFK1-tPFK1* $_{\mathbf{J}}$ *tPFK2-PFK2-pPFK2* $_{\mathbf{K}}$ *pAgTEF1-AmdSYM-tAgTEF1* $_{\mathbf{L}}$ *tGPM1-GPM1-pPGM1* $_{\mathbf{M}}$ *pPDC1-PDC1-tPDC1-SYN* $_{\mathbf{F}}$*) pyk1Δ pgi1Δ tpi1Δ tdh3Δ pfk2Δ::(pTEF1-Spcas9-tCYC1 natNT1) pgk1Δ gpm1Δ fba1Δ hxk2Δ pfk1Δ adh1Δ pdc1Δ eno2Δ* | Postma, Dashko [8] |
| IMX1433 | *MATa ura3-52 his3-1 leu2-3,112 MAL2-8c SUC2 glk1Δ:: (pAgTEF1-SpHIS5-tAgTEF1) hxk1Δ::KlLEU2 tdh1Δ tdh2Δ gpm2Δ gpm3Δ eno1Δ pyk2Δ pdc5Δ pdc6Δ adh2Δ adh5Δ adh4Δ sga1Δ::(* $_{\mathbf{G}}$ *tFBA1-FBA1-pFBA1* $_{\mathbf{H}}$ *pTPI1-TPI1-tTPI1* $_{\mathbf{P}}$ *tPGK1-PGK1-pPGK1* $_{\mathbf{Q}}$ *tADH1-ADH1-pADH1* $_{\mathbf{N}}$ *pPYK1-PYK1-tPYK1* $_{\mathbf{O}}$ *tTDH3-TDH3-pTDH3* $_{\mathbf{A}}$ *pENO2-ENO2-tENO2* $_{\mathbf{B}}$ *pHXK2-HXK2-tHXK2* $_{\mathbf{C}}$ *pPGI-PGI1-tPGI1* $_{\mathbf{D}}$ *pPFK1-PFK1-tPFK1* $_{\mathbf{J}}$ *tPFK2-PFK2-pPFK2* $_{\mathbf{KL}}$ *tGPM1-GPM1-pPGM1* $_{\mathbf{M}}$ *pPDC1-PDC1-tPDC1-SYN* $_{\mathbf{F}}$*) pyk1Δ pgi1Δ tpi1Δ tdh3Δ pfk2Δ::(pTEF1-Spcas9-tCYC1 natNT1) pgk1Δ gpm1Δ fba1Δ hxk2Δ pfk1Δ adh1Δ pdc1Δ eno2Δ in vivo* recombined pMEL10 backbone with repair oligo 11588/11589 | This study |
| IMX1769 | *MATa ura3-52 his3-1 leu2-3,112 MAL2-8c SUC2 glk1Δ:: (pAgTEF1-SpHIS5-tAgTEF1) hxk1Δ::KlLEU2 tdh1Δ tdh2Δ gpm2Δ gpm3Δ eno1Δ pyk2Δ pdc5Δ pdc6Δ adh2Δ adh5Δ adh4Δ sga1Δ::(* $_{\mathbf{G}}$ *tFBA1-* | This study |

| | | |
|---|---|---|
| | *FBA1-pFBA1 $_{H}$ pTPI1-TPI1-tTPI1 $_{P}$ tPGK1-PGK1-pPGK1 $_{Q}$ tADH1-ADH1-pADH1 $_{N}$ pPYK1-PYK1-tPYK1 $_{O}$ tTDH3-TDH3-pTDH3 $_{A}$ pENO2-ENO2-tENO2 $_{B}$ pHXK2-HXK2-tHXK2 $_{C}$ pPGI-PGI1-tPGI1 $_{D}$ pPFK1-PFK1-tPFK1 $_{J}$ tPFK2-PFK2-pPFK2 $_{K L}$ tGPM1-GPM1-pPGM1 $_{M}$ pPDC1-PDC1-tPDC1-SYN $_{F}$) pyk1Δ pgi1Δ tpi1Δ tdh3Δ pfk2Δ::(pTEF1-Spcas9-tCYC1 natNT1) pgk1Δ gpm1Δ fba1Δ hxk2Δ pfk1Δ adh1Δ pdc1Δ eno2Δ* | |
| IMX2059 | *MATa ura3-52 his3-1 leu2-3,112 MAL2-8c SUC2 hxk1Δ::KlLEU2 tdh1Δ tdh2Δ gpm2Δ gpm3Δ eno1Δ pyk2Δ pdc5Δ pdc6Δ adh2Δ adh5Δ adh4Δ sga1Δ::(FBA1 $_{GH}$ TPI1 $_{HP}$ PGK1 $_{PQ}$ ADH1 $_{QN}$ PYK1 $_{NO}$ TDH3 $_{OA}$ ENO2 $_{AB}$ HXK2 $_{BC}$ PGI1 $_{CD}$ PFK1 $_{DJ}$ PFK2 $_{JK}$ AmdSYM $_{K L}$ GPM1 $_{LM}$ PDC1-SYN $_{MF}$) pyk1Δ pgi1Δ tpi1Δ tdh3Δ pfk2Δ::(pTEF-cas9-tCYC1 natNT1) pgk1Δ gpm1Δ fba1Δ hxk2Δ pfk1Δ adh1Δ pdc1Δ eno2Δ glk1Δ::Sphis5Δ::(pGAL1-I SceI-tCYC1) x2::pURA3-URA3-tURA3 $_{DT}$ pHIS3-HIS3-tHIS3* | Postma, Dashko [8] |
| IMX2154 | *MATa ura3-52 his3-1 leu2-3,112 MAL2-8c SUC2 glk1Δ:: (pAgTEF1-SpHIS5-tAgTEF1) hxk1Δ::KlLEU2 tdh1Δ tdh2Δ gpm2Δ gpm3Δ eno1Δ pyk2Δ pdc5Δ pdc6Δ adh2Δ adh5Δ adh4Δ sga1Δ::( $_{G}$ tFBA1-FBA1-pFBA1 $_{H}$ pTPI1-TPI1-tTPI1 $_{P}$ tPGK1-PGK1-pPGK1 $_{Q}$ tADH1-ADH1-pADH1 $_{N}$ pPYK1-PYK1-tPYK1 $_{O}$ tTDH3-TDH3-pTDH3 $_{A}$ pENO2-ENO2-tENO2 $_{B}$ pHXK2-HXK2-tHXK2 $_{C}$ pPGI-PGI1-tPGI1 $_{D}$ pPFK1-PFK1-tPFK1 $_{J}$ tPFK2-PFK2-pPFK2 $_{K L}$ tGPM1-GPM1-pPGM1 $_{M}$ pPDC1-PDC1-tPDC1-SYN $_{F}$) pyk1Δ pgi1Δ tpi1Δ tdh3Δ pfk2Δ::(pTEF1-Spcas9-tCYC1 natNT1) pgk1Δ gpm1Δ fba1Δ hxk2Δ pfk1Δ adh1Δ pdc1Δ eno2Δ gnd2Δ sol4Δ tkl2Δ nqm1Δ pUDR286 pUDR590* | This study |
| IMX2204 | *MATa ura3-52 his3-1 leu2-3,112 MAL2-8c SUC2 glk1Δ:: (pAgTEF1-SpHIS5-tAgTEF1) hxk1Δ::KlLEU2 tdh1Δ tdh2Δ gpm2Δ gpm3Δ eno1Δ pyk2Δ pdc5Δ pdc6Δ adh2Δ adh5Δ adh4Δ sga1Δ::( $_{G}$ tFBA1-FBA1-pFBA1 $_{H}$ pTPI1-TPI1-tTPI1 $_{P}$ tPGK1-PGK1-pPGK1 $_{Q}$ tADH1-ADH1-pADH1 $_{N}$ pPYK1-PYK1-tPYK1 $_{O}$ tTDH3-TDH3-pTDH3 $_{A}$ pENO2-ENO2-tENO2 $_{B}$ pHXK2-HXK2-tHXK2 $_{C}$ pPGI-PGI1-tPGI1 $_{D}$ pPFK1-PFK1-tPFK1 $_{J}$ tPFK2-PFK2-pPFK2 $_{K L}$ tGPM1-GPM1-pPGM1 $_{M}$ pPDC1-PDC1-tPDC1-SYN $_{F}$) pyk1Δ pgi1Δ tpi1Δ tdh3Δ pfk2Δ::(pTEF1-Spcas9-tCYC1 natNT1) pgk1Δ gpm1Δ fba1Δ hxk2Δ pfk1Δ adh1Δ pdc1Δ eno2Δ gnd2Δ sol4Δ tkl2Δ nqm1Δ* | This study |
| IMX2224 | *MATa ura3-52 his3-1 leu2-3,112 MAL2-8c SUC2 hxk1Δ::KlLEU2 tdh1Δ tdh2Δ gpm2Δ gpm3Δ eno1Δ pyk2Δ pdc5Δ pdc6Δ adh2Δ adh5Δ adh4Δ sga1Δ::(FBA1 $_{GH}$ TPI1 $_{HP}$ PGK1 $_{PQ}$ ADH1 $_{QN}$ PYK1 $_{NO}$ TDH3 $_{OA}$ ENO2 $_{AB}$ HXK2 $_{BC}$ PGI1 $_{CD}$ PFK1 $_{DJ}$ PFK2 $_{JK}$ AmdSYM $_{KL}$ GPM1 $_{LM}$ PDC1-SYN $_{MF}$) pyk1Δ pgi1Δ tpi1Δ tdh3Δ pfk2Δ::(pTEF-cas9-tCYC1 natNT1) pgk1Δ gpm1Δ fba1Δ hxk2Δ pfk1Δ adh1Δ pdc1Δ eno2Δ glk1Δ::Sphis5Δ::(pGAL1-I SceI-tCYC1) x2::pURA3-URA3-tURA3 $_{DT}$ pHIS3-HIS3-tHIS3 YPRCtau3Δ::pCCW12-mRuby2-tENO1* | Postma, Dashko [8] |
| IMX2226 | *MATa ura3-52 his3-1 leu2-3,112 MAL2-8c SUC2 hxk1Δ::KlLEU2 tdh1Δ tdh2Δ gpm2Δ gpm3Δ eno1Δ pyk2Δ pdc5Δ pdc6Δ adh2Δ adh5Δ adh4Δ sga1Δ::(FBA1 $_{GH}$ TPI1 $_{HP}$ PGK1 $_{PQ}$ ADH1 $_{QN}$* | Postma, Dashko [8] |

| | | |
|---|---|---|
| | *PYK1 **NO** TDH3 **OA** ENO2 **AB** HXK2 **BC** PGI1 **CD** PFK1 **DJ** PFK2 **JK** AmdSYM **KL** GPM1 LM PDC1-SYN **MF**) pyk1Δ pgi1Δ tpi1Δ tdh3Δ pfk2Δ::(pTEF-cas9-tCYC1 natNT1) pgk1Δ gpm1Δ fba1Δ hxk2Δ pfk1Δ adh1Δ pdc1Δ eno2Δ glk1Δ::Sphis5Δ::(pGAL1-I SceI-tCYC1) x2::pURA3-URA3-tURA3-SHR DT-pHIS3-HIS3-tHIS3 YPRCtau3Δ::pTEF1-Venus-tTDH1* | |
| IMX2234 | *MATa ura3Δ his3Δ leu2-3,112 MAL2-8c SUC2 glk1Δ hxk1Δ::KlLEU2 tdh1Δ tdh2Δ gpm2Δ gpm3Δ eno1Δ pyk2Δ pdc5Δ pdc6Δ adh2Δ adh5Δ adh4Δ sga1Δ::( **G** tFBA1-FBA1-pFBA1 **H** pTPI1-TPI1-tTPI1 **P** tPGK1-PGK1-pPGK1 **Q** tADH1-ADH1-pADH1 **N** pPYK1-PYK1-tPYK1 **O** tTDH3-TDH3-pTDH3 **A** pENO2-ENO2-tENO2 **B** pHXK2-HXK2-tHXK2 **C** pPGI-PGI1-tPGI1 **D** pPFK1-PFK1-tPFK1 **J** tPFK2-PFK2-pPFK2 **KL** tGPM1-GPM1-pPGM1 **M** pPDC1-PDC1-tPDC1-SYN F) pyk1Δ pgi1Δ tpi1Δ tdh3Δ pfk2Δ::(pTEF1-Spcas9-tCYC1 natNT1) pgk1Δ gpm1Δ fba1Δ hxk2Δ pfk1Δ adh1Δ pdc1Δ eno2Δ gnd2Δ sol4Δ tkl2Δ nqm1Δ* | This study |
| IMX2270 | *MATa ura3Δ his3Δ leu2-3,112 MAL2-8c SUC2 glk1Δ hxk1Δ::KlLEU2 tdh1Δ tdh2Δ gpm2Δ gpm3Δ eno1Δ pyk2Δ pdc5Δ pdc6Δ adh2Δ adh5Δ adh4Δ sga1Δ::( **G** tFBA1-FBA1-pFBA1 **H** pTPI1-TPI1-tTPI1 **P** tPGK1-PGK1-pPGK1 **Q** tADH1-ADH1-pADH1 **N** pPYK1-PYK1-tPYK1 **O** tTDH3-TDH3-pTDH3 **A** pENO2-ENO2-tENO2 **B** pHXK2-HXK2-tHXK2 **C** pPGI-PGI1-tPGI1 **D** pPFK1-PFK1-tPFK1 **J** tPFK2-PFK2-pPFK2 **K L** tGPM1-GPM1-pPGM1 **M** pPDC1-PDC1-tPDC1-SYN **F**) pyk1Δ pgi1Δ tpi1Δ tdh3Δ pfk2Δ::(pTEF1-Spcas9-tCYC1 natNT1) pgk1Δ gpm1Δ fba1Δ hxk2Δ pfk1Δ adh1Δ pdc1Δ eno2Δ gnd2Δ sol4Δ tkl2Δ nqm1Δ aro10Δ* | This study |
| IMF2 | *MATa ura3-52 his3-1 leu2-3,112 MAL2-8c SUC2 glk1Δ::(pAgTEF1-SpHIS5-tAgTEF1)Δ::(pGAL1-I SceI-tCYC1) hxk1Δ::KlLEU2 tdh1Δ tdh2Δ gpm2Δ gpm3Δ eno1Δ pyk2Δ pdc5Δ pdc6Δ adh2Δ adh5Δ adh4Δ sga1Δ::( **G** tFBA1-FBA1-pFBA1 **H** pTPI1-TPI1-tTPI1 **P** tPGK1-PGK1-pPGK1 **Q** tADH1-ADH1-pADH1 **N** pPYK1-PYK1-tPYK1 **O** tTDH3-TDH3-pTDH3 **A** pENO2-ENO2-tENO2 **B** pHXK2-HXK2-tHXK2 **C** pPGI-PGI1-tPGI1 **D** pPFK1-PFK1-tPFK1 **J** tPFK2-PFK2-pPFK2 **K** pAgTEF1-AmdSYM-tAgTEF1 **L** tGPM1-GPM1-pPGM1 **M** pPDC1-PDC1-tPDC1-SYN **F**) pyk1Δ pgi1Δ tpi1Δ tdh3Δ pfk2Δ::(pTEF1-Spcas9-tCYC1 natNT1) pgk1Δ gpm1Δ fba1Δ hxk2Δ pfk1Δ adh1Δ pdc1Δ eno2Δ NeoChr2* | Postma, Dashko [8] |
| IMF6 | *MATa ura3-52 his3-1 leu2-3,112 MAL2-8c SUC2 glk1Δ::(pAgTEF1-SpHIS5-tAgTEF1)Δ::(pGAL1-I SceI-tCYC1) hxk1Δ::KlLEU2 tdh1Δ tdh2Δ gpm2Δ gpm3Δ eno1Δ pyk2Δ pdc5Δ pdc6Δ adh2Δ adh5Δ adh4Δ sga1Δ::( **G** tFBA1-FBA1-pFBA1 **H** pTPI1-TPI1-tTPI1 P tPGK1-PGK1-pPGK1 **Q** tADH1-ADH1-pADH1 **N** pPYK1-PYK1-tPYK1 **O** tTDH3-TDH3-pTDH3 **A** pENO2-ENO2-tENO2 **B** pHXK2-HXK2-tHXK2 **C** pPGI-PGI1-tPGI1 **D** pPFK1-PFK1-tPFK1 **J** tPFK2-PFK2-pPFK2 **K** pAgTEF1-AmdSYM-tAgTEF1 **L** tGPM1-GPM1-pPGM1 **M** pPDC1-PDC1-tPDC1-SYN **F**) pyk1Δ pgi1Δ tpi1Δ tdh3Δ pfk2Δ::(pTEF1-Spcas9-tCYC1 natNT1) pgk1Δ gpm1Δ fba1Δ hxk2Δ pfk1Δ adh1Δ pdc1Δ eno2Δ NeoChr1* | Postma, Dashko [8] |

| | | |
|---|---|---|
| IMF22* | *MATa ura3-52 his3-1 leu2-3,112 MAL2-8c SUC2 glk1Δ::(pAgTEF1-SpHIS5-tAgTEF1)Δ::(pGAL1-I SceI-tCYC1) hxk1Δ::KlLEU2 tdh1Δ tdh2Δ gpm2Δ gpm3Δ eno1Δ pyk2Δ pdc5Δ pdc6Δ adh2Δ adh5Δ adh4Δ sga1Δ::( G tFBA1-FBA1-pFBA1 H pTPI1-TPI1-tTPI1 P tPGK1-PGK1-pPGK1 Q tADH1-ADH1-pADH1 N pPYK1-PYK1-tPYK1 O tTDH3-TDH3-pTDH3 A pENO2-ENO2-tENO2 B pHXK2-HXK2-tHXK2 C pPGI-PGI1-tPGI1 D pPFK1-PFK1-tPFK1 J tPFK2-PFK2-pPFK2 K pAgTEF1-AmdSYM-tAgTEF1 L tGPM1-GPM1-pPGM1 M pPDC1-PDC1-tPDC1-SYN F) pyk1Δ pgi1Δ tpi1Δ tdh3Δ pfk2Δ::(pTEF1-Spcas9-tCYC1 natNT1) pgk1Δ gpm1Δ fba1Δ hxk2Δ pfk1Δ adh1Δ pdc1Δ eno2Δ NeoChr10* | This study |
| IMF23* | *MATa ura3-52 his3-1 leu2-3,112 MAL2-8c SUC2 glk1Δ::(pAgTEF1-SpHIS5-tAgTEF1)Δ::(pGAL1-I SceI-tCYC1) hxk1Δ::KlLEU2 tdh1Δ tdh2Δ gpm2Δ gpm3Δ eno1Δ pyk2Δ pdc5Δ pdc6Δ adh2Δ adh5Δ adh4Δ sga1Δ::( G tFBA1-FBA1-pFBA1 H pTPI1-TPI1-tTPI1 P tPGK1-PGK1-pPGK1 Q tADH1-ADH1-pADH1 N pPYK1-PYK1-tPYK1 O tTDH3-TDH3-pTDH3 A pENO2-ENO2-tENO2 B pHXK2-HXK2-tHXK2 C pPGI-PGI1-tPGI1 D pPFK1-PFK1-tPFK1 J tPFK2-PFK2-pPFK2 K pAgTEF1-AmdSYM-tAgTEF1 L tGPM1-GPM1-pPGM1 M pPDC1-PDC1-tPDC1-SYN F) pyk1Δ pgi1Δ tpi1Δ tdh3Δ pfk2Δ::(pTEF1-Spcas9-tCYC1 natNT1) pgk1Δ gpm1Δ fba1Δ hxk2Δ pfk1Δ adh1Δ pdc1Δ eno2Δ NeoChr12* | Postma, Dashko [8] |
| IMF27* | *MATa ura3Δ his3Δ leu2-3,112 MAL2-8c SUC2 glk1Δ hxk1Δ::KlLEU2 tdh1Δ tdh2Δ gpm2Δ gpm3Δ eno1Δ pyk2Δ pdc5Δ pdc6Δ adh2Δ adh5Δ adh4Δ sga1Δ::( G tFBA1-FBA1-pFBA1 H pTPI1-TPI1-tTPI1 P tPGK1-PGK1-pPGK1 Q tADH1-ADH1-pADH1 N pPYK1-PYK1-tPYK1 O tTDH3-TDH3-pTDH3 A pENO2-ENO2-tENO2 B pHXK2-HXK2-tHXK2 C pPGI-PGI1-tPGI1 D pPFK1-PFK1-tPFK1 J tPFK2-PFK2-pPFK2 K L tGPM1-GPM1-pPGM1 M pPDC1-PDC1-tPDC1-SYN F) pyk1Δ pgi1Δ tpi1Δ tdh3Δ pfk2Δ::(pTEF1-Spcas9-tCYC1 natNT1) pgk1Δ gpm1Δ fba1Δ hxk2Δ pfk1Δ adh1Δ pdc1Δ eno2Δ gnd2Δ sol4Δ tkl2Δ nqm1Δ aro10Δ NeoChr25* | This study |
| IMF29* | *MATa ura3Δ his3Δ leu2-3,112 MAL2-8c SUC2 glk1Δ hxk1Δ::KlLEU2 tdh1Δ tdh2Δ gpm2Δ gpm3Δ eno1Δ pyk2Δ pdc5Δ pdc6Δ adh2Δ adh5Δ adh4Δ sga1Δ::( G tFBA1-FBA1-pFBA1 H pTPI1-TPI1-tTPI1 P tPGK1-PGK1-pPGK1 Q tADH1-ADH1-pADH1 N pPYK1-PYK1-tPYK1 O tTDH3-TDH3-pTDH3 A pENO2-ENO2-tENO2 B pHXK2-HXK2-tHXK2 C pPGI-PGI1-tPGI1 D pPFK1-PFK1-tPFK1 J tPFK2-PFK2-pPFK2 K L tGPM1-GPM1-pPGM1 M pPDC1-PDC1-tPDC1-SYN F) pyk1Δ pgi1Δ tpi1Δ tdh3Δ pfk2Δ::(pTEF1-Spcas9-tCYC1 natNT1) pgk1Δ gpm1Δ fba1Δ hxk2Δ pfk1Δ adh1Δ pdc1Δ eno2Δ gnd2Δ sol4Δ tkl2Δ nqm1Δ aro10Δ NeoChr26* | This study |
| IMF31 | *MATa ura3Δ his3Δ leu2-3,112 MAL2-8c SUC2 glk1Δ hxk1Δ::KlLEU2 tdh1Δ tdh2Δ gpm2Δ gpm3Δ eno1Δ pyk2Δ pdc5Δ pdc6Δ adh2Δ adh5Δ adh4Δ sga1Δ::pKlURA3-KlURA3-tKlURA3 pyk1Δ pgi1Δ tpi1Δ tdh3Δ pfk2Δ::(pTEF1-Spcas9-tCYC1 natNT1) pgk1Δ gpm1Δ fba1Δ hxk2Δ pfk1Δ adh1Δ pdc1Δ eno2Δ gnd2Δ sol4Δ tkl2Δ nqm1Δ aro10Δ NeoChr25* | This study |

| | | |
|---|---|---|
| IMF32 | *MATa ura3Δ his3Δ leu2-3,112 MAL2-8c SUC2 glk1Δ hxk1Δ::KlLEU2 tdh1Δ tdh2Δ gpm2Δ gpm3Δ eno1Δ pyk2Δ pdc5Δ pdc6Δ adh2Δ adh5Δ adh4Δ sga1Δ pyk1Δ pgi1Δ tpi1Δ tdh3Δ pfk2Δ::(pTEF1-Spcas9-tCYC1 natNT1) pgk1Δ gpm1Δ fba1Δ hxk2Δ pfk1Δ adh1Δ pdc1Δ eno2Δ gnd2Δ sol4Δ tkl2Δ nqm1Δ aro10Δ NeoChr26* | This study |
| IMF33 | *MATa ura3Δ his3Δ leu2-3,112 MAL2-8c SUC2 glk1Δ hxk1Δ::KlLEU2 tdh1Δ tdh2Δ gpm2Δ gpm3Δ eno1Δ pyk2Δ pdc5Δ pdc6Δ adh2Δ adh5Δ adh4Δ sga1Δ::pKlURA3-KlURA3-tKlURA3 pyk1Δ pgi1Δ tpi1Δ tdh3Δ pfk2Δ::(pTEF1-Spcas9-tCYC1 natNT1) pgk1Δ gpm1Δ fba1Δ hxk2Δ pfk1Δ adh1Δ pdc1Δ eno2Δ gnd2Δ sol4Δ tkl2Δ nqm1Δ aro10Δ NeoChr25 zwf1Δ sol3Δ gnd1Δ rki1Δ* | This study |
| IMF34 | *MATa ura3Δ his3Δ leu2-3,112 MAL2-8c SUC2 glk1Δ hxk1Δ::KlLEU2 tdh1Δ tdh2Δ gpm2Δ gpm3Δ eno1Δ pyk2Δ pdc5Δ pdc6Δ adh2Δ adh5Δ adh4Δ sga1Δ::pKlURA3-KlURA3-tKlURA3 pyk1Δ pgi1Δ tpi1Δ tdh3Δ pfk2Δ::(pTEF1-Spcas9-tCYC1 natNT1) pgk1Δ gpm1Δ fba1Δ hxk2Δ pfk1Δ adh1Δ pdc1Δ eno2Δ gnd2Δ sol4Δ tkl2Δ nqm1Δ aro10Δ NeoChr25 zwf1Δ sol3Δ gnd1Δ rki1Δ tkl1Δ tal1Δ rpe1Δ* | This study |
| IMF35 | *MATa ura3Δ his3Δ leu2-3,112 MAL2-8c SUC2 glk1Δ hxk1Δ::KlLEU2 tdh1Δ tdh2Δ gpm2Δ gpm3Δ eno1Δ pyk2Δ pdc5Δ pdc6Δ adh2Δ adh5Δ adh4Δ sga1Δ pyk1Δ pgi1Δ tpi1Δ tdh3Δ pfk2Δ::(pTEF1-Spcas9-tCYC1 natNT1) pgk1Δ gpm1Δ fba1Δ hxk2Δ pfk1Δ adh1Δ pdc1Δ eno2Δ gnd2Δ sol4Δ tkl2Δ nqm1Δ aro10Δ NeoChr26::(rki1::RKI1)* | This study |
| IMF36 | *MATa ura3Δ his3Δ leu2-3,112 MAL2-8c SUC2 glk1Δ hxk1Δ::KlLEU2 tdh1Δ tdh2Δ gpm2Δ gpm3Δ eno1Δ pyk2Δ pdc5Δ pdc6Δ adh2Δ adh5Δ adh4Δ sga1Δ pyk1Δ pgi1Δ tpi1Δ tdh3Δ pfk2Δ::(pTEF1-Spcas9-tCYC1 natNT1) pgk1Δ gpm1Δ fba1Δ hxk2Δ pfk1Δ adh1Δ pdc1Δ eno2Δ gnd2Δ sol4Δ tkl2Δ nqm1Δ aro10Δ NeoChr26::(rki1::RKI1) zwf1Δ sol3Δ gnd1Δ rki1Δ* | This study |
| IMF40 | *MATa ura3Δ his3Δ leu2-3,112 MAL2-8c SUC2 glk1Δ hxk1Δ::KlLEU2 tdh1Δ tdh2Δ gpm2Δ gpm3Δ eno1Δ pyk2Δ pdc5Δ pdc6Δ adh2Δ adh5Δ adh4Δ sga1Δ pyk1Δ pgi1Δ tpi1Δ tdh3Δ pfk2Δ::(pTEF1-Spcas9-tCYC1 natNT1) pgk1Δ gpm1Δ fba1Δ hxk2Δ pfk1Δ adh1Δ pdc1Δ eno2Δ gnd2Δ sol4Δ tkl2Δ nqm1Δ aro10Δ NeoChr26::(rki1::RKI1) zwf1Δ sol3Δ gnd1Δ rki1Δ tkl1Δ tal1Δ rpe1Δ* | This study |
| IMF41* | *MATa ura3Δ his3Δ leu2-3,112 MAL2-8c SUC2 glk1Δ hxk1Δ::KlLEU2 tdh1Δ tdh2Δ gpm2Δ gpm3Δ eno1Δ pyk2Δ pdc5Δ pdc6Δ adh2Δ adh5Δ adh4Δ sga1Δ pyk1Δ pgi1Δ tpi1Δ tdh3Δ pfk2Δ::(pTEF1-Spcas9-tCYC1 natNT1) pgk1Δ gpm1Δ fba1Δ hxk2Δ pfk1Δ adh1Δ pdc1Δ eno2Δ gnd2Δ sol4Δ tkl2Δ nqm1Δ aro10Δ zwf1Δ sol3Δ gnd1Δ rki1Δ tkl1Δ tal1Δ rpe1Δ NeoChr30* | This study |
| IMF42* | *MATa ura3Δ his3Δ leu2-3,112 MAL2-8c SUC2 glk1Δ hxk1Δ::KlLEU2 tdh1Δ tdh2Δ gpm2Δ gpm3Δ eno1Δ pyk2Δ pdc5Δ pdc6Δ adh2Δ adh5Δ adh4Δ sga1Δ::pKlURA3-KlURA3-tKlURA3 pyk1Δ pgi1Δ tpi1Δ tdh3Δ pfk2Δ::(pTEF1-Spcas9-tCYC1 natNT1) pgk1Δ gpm1Δ fba1Δ hxk2Δ pfk1Δ adh1Δ pdc1Δ eno2Δ gnd2Δ sol4Δ tkl2Δ nqm1Δ aro10Δ zwf1Δ sol3Δ gnd1Δ rki1Δ tkl1Δ tal1Δ rpe1Δ NeoChr31* | This study |

| | | |
|---|---|---|
| IMF44 | *MATa ura3Δ his3Δ leu2-3,112 MAL2-8c SUC2 glk1Δ hxk1Δ::KlLEU2 tdh1Δ tdh2Δ gpm2Δ gpm3Δ eno1Δ pyk2Δ pdc5Δ pdc6Δ adh2Δ adh5Δ adh4Δ sga1Δ::pKlURA3-KlURA3-tKlURA3 pyk1Δ pgi1Δ tpi1Δ tdh3Δ pfk2Δ::(pTEF1-Spcas9-tCYC1 natNT1) pgk1Δ gpm1Δ fba1Δ hxk2Δ pfk1Δ adh1Δ pdc1Δ eno2Δ gnd2Δ sol4Δ tkl2Δ nqm1Δ aro10Δ zwf1Δ sol3Δ gnd1Δ rki1Δ tkl1Δ tal1Δ rpe1Δ x2Δ::pTEF1-coAtCHS3-tMDH1 yprctau3Δ::pTEF1-coAtCHS3-tMDH1 spr3Δ::pTEF1-coAtCHS3-tMDH1 can1::pTEF1-coAtCHS3-tMDH NeoChr31* | This study |
| IMF47* | *MATa ura3Δ his3Δ leu2-3,112 MAL2-8c SUC2 glk1Δ hxk1Δ::KlLEU2 tdh1Δ tdh2Δ gpm2Δ gpm3Δ eno1Δ pyk2Δ pdc5Δ pdc6Δ adh2Δ adh5Δ adh4Δ sga1Δ::pKlURA3-KlURA3-tKlURA3 pyk1Δ pgi1Δ tpi1Δ tdh3Δ pfk2Δ::(pTEF1-Spcas9-tCYC1 natNT1) pgk1Δ gpm1Δ fba1Δ hxk2Δ pfk1Δ adh1Δ pdc1Δ eno2Δ gnd2Δ sol4Δ tkl2Δ nqm1Δ aro10Δ zwf1Δ sol3Δ gnd1Δ rki1Δ tkl1Δ tal1Δ rpe1Δ x2Δ::pTEF1-coAtCHS3-tMDH1 yprctau3Δ::pTEF1-coAtCHS3-tMDH1 spr3Δ::pTEF1-coAtCHS3-tMDH1 can1::pTEF1-coAtCHS3-tMDH NeoChr33* | This study |
| IMF48* | *MATa ura3Δ his3Δ leu2-3,112 MAL2-8c SUC2 glk1Δ hxk1Δ::KlLEU2 tdh1Δ tdh2Δ gpm2Δ gpm3Δ eno1Δ pyk2Δ pdc5Δ pdc6Δ adh2Δ adh5Δ adh4Δ sga1Δ::pKlURA3-KlURA3-tKlURA3 pyk1Δ pgi1Δ tpi1Δ tdh3Δ pfk2Δ::(pTEF1-Spcas9-tCYC1 natNT1) pgk1Δ gpm1Δ fba1Δ hxk2Δ pfk1Δ adh1Δ pdc1Δ eno2Δ gnd2Δ sol4Δ tkl2Δ nqm1Δ aro10Δ zwf1Δ sol3Δ gnd1Δ rki1Δ tkl1Δ tal1Δ rpe1Δ x2Δ::pTEF1-coAtCHS3-tMDH1 yprctau3Δ::pTEF1-coAtCHS3-tMDH1 spr3Δ::pTEF1-coAtCHS3-tMDH1 can1::pTEF1-coAtCHS3-tMDH NeoChr34* | This study |

### Supplementary Table 7 - Neochromosome configurations

SHRs are differently annotated than in Kuijpers *et al.* [6]. SHRs are annotated in subscript between the genetic fragments that they join together.

| Name | Size | Notes | Stocked name | Neochromosome configuration |
|---|---|---|---|---|
| **NeoChr1** | 100 kb | Circular | IMF6 | AO' 7A BJ 7B BK 7C BL 7D AP 8A BM 8B BN 8C BO 8D AC *pCCW12-mRuby2-tENO1* AD *CEN6/ARS4* AE 1A AT 1B AS 1C AU 1D AF 2A AV 2B AW 2C AX 2D AG 3A AY 3B AZ 3C BA 3D AH *pTEF2-mTurquoise2-tSSA 1* AI ARS417 BU *pHIS3-HIS3-tHIS3* AJ 4A BC 4B BD 4C BE 4D AK 9A BF 9B BS 9C BT 9D AQ 5A BG 5B BH 5C BI 5D AL *pTEF1-Venus-tTDH1* AM ARS1 AN 6A BP 6B BQ 6C BR 6D AO` Telomerator AO` |
| **NeoChr2** | 50 kb | Circular | IMF2 | AO' 7A BJ 7B BK 7C BL 7D AC *pCCW12-mRuby2-tENO1* AD *CEN6/ARS4* AE 1A AT 1B AS 1C AU 1Ds AH *pTEF2 - mTurquoise2 - tSSA1* AI *pHIS3-HIS3-tHIS3* AJ 4A BC 4B BD 4C BE 4D AL *pTEF1 - Venus - tTDH1* AM *ARS1* AN 6A BP 6B BQ 6C BR 6D AO' Telomerator AO' |
| **NeoChr10** | 100 kb | Linear | IMF22 | Telomere AO 7A BJ 7B BK 7C BL 7D AP 8A BM 8B BN 8C BO 8D AC *pCCW12-mRuby2-tENO1* AD *ARS1* AN 18A BP 18B BQ 19C BR 19D DE 15A DF 15B DH 15C DI 15D DJ *CEN6/ARS4* AE 16A DK 16B DL 16C DM 16D DN 17A DO 17B DP 19A DQ 17D DR *ARS417* BU *pHIS3-HIS3-tHIS3* AJ 4A BC 4B BD 4C BE 4D AK 9A BF 9B BS 9C BT 9D AQ 5A BG 5B BH 5C BI 5D AL *tSSA1-mTurquoise2- pTEF2* DS Telomere |
| **NeoChr11** | 100 kb | Linear. Few bases changed in right telomere to prevent recirculari-sation | - | Telomere AO 7A BJ 7B BK 7C BL 7D AP 8A BM 8B BN 8C BO 8D AC *pCCW12-mRuby2-tENO1* AD *ARS1* AN 18A BP 18B BQ 19C BR 19D DE 15A DF 15B DH 15C DI 15D DJ *CEN6/ARS4* AE 16A DK 16B DL 16C DM 16D DN 17A DO 17B DP 19A DQ 17D DR *ARS417* BU *pHIS3-HIS3-tHIS3* AJ 4A BC 4B BD 4C BE 4D AK 9A BF 9B BS 9C BT 9D AQ 5A BG 5B BH 5C BI 5D AL *tSSA1-mTurquoise2- pTEF2* DS Telomere |
| **NeoChr12** | 100 kb | Circular | IMF23 | AO 7A BJ 7B BK 7C BL 7D AP 8A BM 8B BN 8C BO 8D AC *pCCW12-mRuby2-tENO1* AD *ARS1* AN 18A BP 18B BQ 19C BR 19D DE 15A DF 15B DH 15C DI 15D DJ *CEN6/ARS4* AE 16A DK 16B DL 16C DM 16D DN 17A DO 17B DP 19A DQ 17D DR *ARS417* BU *pHIS3-HIS3-tHIS3* AJ 4A BC 4B BD 4C BE 4D AK 9A BF 9B BS 9C BT 9D AQ 5A BG 5B BH 5C BI 5D AL *tSSA1-mTurquoise2- pTEF2* DS Telomerator AO |
| **NeoChr25** | 100 kb | Linear | IMF27, IMF31, IMF33, IMF34 | telomere BJ 7BC BL *ARS1* AN *pZWF1-ZWF1-tZWF1* BP *pTKL1-TKL1-tTKL1* DE *pGND1-GND1-tGND1* BQ *tRKI1-RKI1-pRKI1* BR *tTAL1-TAL1-pTAL1* AL *tSSA1-mTurquoise2-pTEF2* DS *tRPE1-RPE1-pRPE1* DF *tSOL3-SOL3-pSOL3* DI *ARS417* BE *pHHF1-coEcaroG$^{(P150L)}$-tTEF1* DK *pHHF2-coEcaroB-tTEF2* AC *pCCW12-mRuby2-tENO1* AD *pRPL25-coEcaroD-tGPH1* DM |

| | | | | |
|---|---|---|---|---|
| | | | | pRPP0-coEcaroE-tCYC1 DN pHTB2-coEcaroL-tPGM2 DO pRPL3-coEcaroA-tSOL4 DP tGPD2-coEcaroC-pRPL8A DQ tGDB1-coEctyrA$^{(M53I,A354V)}$-pRPL10 DR tGSY2-coEcpheA$^{(T326P)}$ -pRPL18B AJ tGLC3-coEctyrB-pCWP2 DH 15CD DJ CEN6/ARS4 AE 16AB DL pFBA1-FBA1-tFBA1 H pTPI1-TPI1-tTPI1 P pPGK1-PGK1-tPGK1 Q pADH1-ADH1-tADH1 N pPYK1-PYK1-tPYK1 O pTDH3-TDH3-tTDH3 A pENO2-ENO2-tENO2 B tHXK2-HXK2-pHXK2 C tPGI1-PGI1-pPGI1 D tPFK1-PFK1-pPFK1 J tPFK2-PFK2-pPFK2 BU tHIS3-HIS3-pHIS3 L tGPM1-GPM1-pGPM1 M tPDC1-PDC1-pPDC1 AR ARS1211 BS 9CD AQ telomere |
| **NeoChr26** | 100 kb | Circular | IMF29, IMF32, IMF35, IMF36, IMF40 | BJ 7BC BL ARS1 AN pZWF1-ZWF1-tZWF1 BP pTKL1-TKL1-tTKL1 DE pGND1-GND1-tGND1 BQ tRKI1-RKI1-pRKI1 BR tTAL1-TAL1-pTAL1 AL tSSA1-mTurquoise2-pTEF2 DS tRPE1-RPE1-pRPE1 DF tSOL3-SOL3-pSOL3 DI ARS417 BE pHHF1-coEcaroG$^{(P150L)}$-tTEF1 DK pHHF2-coEcaroB-tTEF2 AC pCCW12-mRuby2-tENO1 AD pRPL25-coEcaroD-tGPH1 DM pRPP0-coEcaroE-tCYC1 DN pHTB2-coEcaroL-tPGM2 DO pRPL3-coEcaroA-tSOL4 DP tGPD2-coEcaroC-pRPL8A DQ tGDB1-coEctyrA$^{(M53I,A354V)}$-pRPL10 DR tGSY2-coEcpheA$^{(T326P)}$ -pRPL18B AJ tGLC3-coEctyrB-pCWP2 DH 15CD DJ CEN6/ARS4 AE 16AB DL pFBA1-FBA1-tFBA1 H pTPI1-TPI1-tTPI1 P pPGK1-PGK1-tPGK1 Q pADH1-ADH1-tADH1 N pPYK1-PYK1-tPYK1 O pTDH3-TDH3-tTDH3 A pENO2-ENO2-tENO2 B tHXK2-HXK2-pHXK2 C tPGI1-PGI1-pPGI1 D tPFK1-PFK1-pPFK1 J tPFK2-PFK2-pPFK2 BU tHIS3-HIS3-pHIS3 L tGPM1-GPM1-pGPM1 M tPDC1-PDC1-pPDC1 AR ARS1211 BS 9CD AQ telomerator |
| **NeoChr30** | 128 kb | Circular. Insertion of anthocyanin pathway in NeoChr26 of strain IMF40 | IMF41 | BJ 7BC BL ARS1 AN pZWF1-ZWF1-tZWF1 BP pTKL1-TKL1-tTKL1 DE pGND1-GND1-tGND1 BQ tRKI1-RKI1-pRKI1 BR tTAL1-TAL1-pTAL1 AL tSSA1-mTurquoise2-pTEF2 DS tRPE1-RPE1-pRPE1 DF tSOL3-SOL3-pSOL3 DI ARS417 BE pHHF1-coEcaroG$^{(P150L)}$-tTEF1 DK pHHF2-coEcaroB-tTEF2 AC pCCW12-mRuby2-tENO1 AD pRPL25-coEcaroD-tGPH1 DM pRPP0-coEcaroE-tCYC1 DN pHTB2-coEcaroL-tPGM2 DO pRPL3-coEcaroA-tSOL4 DP tGPD2-coEcaroC-pRPL8A DQ tGDB1-coEctyrA$^{(M53I,A354V)}$-pRPL10 DR tGSY2-coEcpheA$^{(T326P)}$ -pRPL18B AJ tGLC3-coEctyrB-pCWP2 DH 15CD DJ CEN6/ARS4 AE 16AB pRPS3-coAtCPR1-tIDH2 F pSePDC1-AtPAL1-tLAT1 DW pSeGPM1-coRcTAL1-tCIT1 DX pSeTPI1-At4CL3-tSDH2 DY pTEF1-coAtCHS3-tMDH1 AM tSDH4-AtCHI1-pSkADH1 AB tADH3-coAtC4H-pSeFBA1 DC tSDH3-coAtF3H-pSkTDH3 EA tACO1-coGhDFR-pSePGK1 EB tFUM1-coAtANS-pSeENO2 EC tDIC1-coAt3GT-pSePYK1 CJ ARS106 DL pFBA1-FBA1-tFBA1 H pTPI1-TPI1-tTPI1 P pPGK1-PGK1-tPGK1 Q pADH1-ADH1-tADH1 N pPYK1-PYK1-tPYK1 O pTDH3-TDH3-tTDH3 A pENO2-ENO2-tENO2 B tHXK2-HXK2-pHXK2 C tPGI1-PGI1-pPGI1 D tPFK1-PFK1- |

| | | | | |
|---|---|---|---|---|
| | | | | pPFK1 J tPFK2-PFK2-pPFK2 BU tHIS3-HIS3-pHIS3 L tGPM1-GPM1-pGPM1 M tPDC1-PDC1-pPDC1 AR ARS1211 BS 9CD AQ Telomerator |
| **NeoChr31** | 128 kb | Linear. Insertion of anthocyanin pathway in NeoChr25 of strain IMF34 | IMF42, IMF44 | Telomere BJ 7BC BL ARS1 AN pZWF1-ZWF1-tZWF1 BP pTKL1-TKL1-tTKL1 DE pGND1-GND1-tGND1 BQ tRKI1-RKI1-pRKI1 BR tTAL1-TAL1-pTAL1 AL tSSA1-mTurquoise2-pTEF2 DS tRPE1-RPE1-pRPE1 DF tSOL3-SOL3-pSOL3 DI ARS417 BE pHHF1-coEcaroG$^{(P150L)}$-tTEF1 DK pHHF2-coEcaroB-tTEF2 AC pCCW12-mRuby2-tENO1 AD pRPL25-coEcaroD-tGPH1 DM pRPP0-coEcaroE-tCYC1 DN pHTB2-coEcaroL-tPGM2 DO pRPL3-coEcaroA-tSOL4 DP tGPD2-coEcaroC-pRPL8A DQ tGDB1-coEctyrA$^{(M53I,A354V)}$-pRPL10 DR tGSY2-coEcpheA$^{(T326P)}$ -pRPL18B AJ tGLC3-coEctyrB-pCWP2 DH 15CD DJ CEN6/ARS4 AE 16AB pRPS3-coAtCPR1-tIDH2 F pSePDC1-AtPAL1-tLAT1 DW pSeGPM1-coRcTAL1-tCIT1 DX pSeTPI1-At4CL3-tSDH2 DY pTEF1-coAtCHS3-tMDH1 AM tSDH4-AtCHI1-pSkADH1 AB tADH3-coAtC4H-pSeFBA1 DC tSDH3-coAtF3H-pSkTDH3 EA tACO1-coGhDFR-pSePGK1 EB tFUM1-coAtANS-pSeENO2 EC tDIC1-coAt3GT-pSePYK1 CJ ARS106 DL pFBA1-FBA1-tFBA1 H pTPI1-TPI1-tTPI1 P pPGK1-PGK1-tPGK1 Q pADH1-ADH1-tADH1 N pPYK1-PYK1-tPYK1 O pTDH3-TDH3-tTDH3 A pENO2-ENO2-tENO2 B tHXK2-HXK2-pHXK2 C tPGI1-PGI1-pPGI1 D tPFK1-PFK1-pPFK1 J tPFK2-PFK2-pPFK2 BU tHIS3-HIS3-pHIS3 L tGPM1-GPM1-pGPM1 M tPDC1-PDC1-pPDC1 AR ARS1211 BS 9CD AQ Telomere |
| **NeoChr33** | 137 kb | Insertion of *AtCHS* at Chunk 7BC, chunk 15CD, SHR N and chunk 9CD in NeoChr31 of strain IMF42 and IMF44. | IMF47 (from IMF44) | Telomere BJ 7BC tMDH1-coAtCHS3-pTEF1 7BC BL ARS1 AN pZWF1-ZWF1-tZWF1 BP pTKL1-TKL1-tTKL1 DE pGND1-GND1-tGND1 BQ tRKI1-RKI1-pRKI1 BR tTAL1-TAL1-pTAL1 AL tSSA1-mTurquoise2-pTEF2 DS tRPE1-RPE1-pRPE1 DF tSOL3-SOL3-pSOL3 DI ARS417 BE pHHF1-coEcaroG$^{(P150L)}$-tTEF1 DK pHHF2-coEcaroB-tTEF2 AC pCCW12-mRuby2-tENO1 AD pRPL25-coEcaroD-tGPH1 DM pRPP0-coEcaroE-tCYC1 DN pHTB2-coEcaroL-tPGM2 DO pRPL3-coEcaroA-tSOL4 DP tGPD2-coEcaroC-pRPL8A DQ tGDB1-coEctyrA$^{(M53I,A354V)}$-pRPL10 DR tGSY2-coEcpheA$^{(T326P)}$ -pRPL18B AJ tGLC3-coEctyrB-pCWP2 DH 15CD tMDH1-coAtCHS3-pTEF1 15 CD DJ CEN6/ARS4 AE 16AB pRPS3-coAtCPR1-tIDH2 F pSePDC1-AtPAL1-tLAT1 DW pSeGPM1-coRcTAL1-tCIT1 DX pSeTPI1-At4CL3-tSDH2 DY pTEF1-coAtCHS3-tMDH1 AM tSDH4-AtCHI1-pSkADH1 AB tADH3-coAtC4H-pSeFBA1 DC tSDH3-coAtF3H-pSkTDH3 EA tACO1-coGhDFR-pSePGK1 EB tFUM1-coAtANS-pSeENO2 EC tDIC1-coAt3GT-pSePYK1 CJ ARS106 DL pFBA1-FBA1-tFBA1 H pTPI1-TPI1-tTPI1 P pPGK1-PGK1-tPGK1 Q pADH1-ADH1-tADH1 N pTEF1-coAtCHS3-tMDH1 pPYK1-PYK1-tPYK1 O pTDH3-TDH3-tTDH3 A pENO2-ENO2-tENO2 B tHXK2-HXK2-pHXK2 C tPGI1-PGI1-pPGI1 D tPFK1-PFK1-pPFK1 J |

| | | | | |
|---|---|---|---|---|
| | | | | *tPFK2-PFK2-pPFK2* BU *tHIS3-HIS3-pHIS3* L *tGPM1-GPM1-pGPM1* M *tPDC1-PDC1-pPDC1* AR *ARS1211* BS 9CD *pTEF1-coAtCHS3-tMDH1 9CD* AQ Telomere |
| **NeoChr34** | 137 kb | Insertion of *CoAtANS* in *mTurquoise* | IMF48 (from IMF47) | Telomere BJ 7BC *tMDH1-coAtCHS3-pTEF1* 7BC BL *ARS1* AN *pZWF1-ZWF1-tZWF1* BP *pTKL1-TKL1-tTKL1* DE *pGND1-GND1-tGND1* BQ *tRKI1-RKI1-pRKI1* BR *tTAL1-TAL1-pTAL1* AL *tSSA1-mTurquoise2-pTEF2Δ::tFUM1-CoAtANS-pSeENO2* DS *tRPE1-RPE1-pRPE1* DF *tSOL3-SOL3-pSOL3* DI *ARS417* BE *pHHF1-coEcaroG*$^{(P150L)}$*-tTEF1* DK *pHHF2-coEcaroB-tTEF2* AC *pCCW12-mRuby2-tENO1* AD *pRPL25-coEcaroD-tGPH1* DM *pRPP0-coEcaroE-tCYC1* DN *pHTB2-coEcaroL-tPGM2* DO *pRPL3-coEcaroA-tSOL4* DP *tGPD2-coEcaroC-pRPL8A* DQ *tGDB1-coEctyrA*$^{(M53I,A354V)}$*-pRPL10* DR *tGSY2-coEcpheA*$^{(T326P)}$ *-pRPL18B* AJ *tGLC3-coEctyrB-pCWP2* DH 15CD *tMDH1-coAtCHS3-pTEF1 15 CD* DJ *CEN6/ARS4* AE 16AB *pRPS3-coAtCPR1-tIDH2* F *pSePDC1-AtPAL1-tLAT1* DW *pSeGPM1-coRcTAL1-tCIT1* DX *pSeTPI1-At4CL3-tSDH2* DY *pTEF1-coAtCHS3-tMDH1* AM *tSDH4-AtCHI1-pSkADH1* AB *tADH3-coAtC4H-pSeFBA1* DC *tSDH3-coAtF3H-pSkTDH3* EA *tACO1-coGhDFR-pSePGK1* EB *tFUM1-coAtANS-pSeENO2* EC *tDIC1-coAt3GT-pSePYK1* CJ *ARS106* DL *pFBA1-FBA1-tFBA1* H *pTPI1-TPI1-tTPI1* P *pPGK1-PGK1-tPGK1* Q *pADH1-ADH1-tADH1* N *pTEF1-coAtCHS3-tMDH1 pPYK1-PYK1-tPYK1* O *pTDH3-TDH3-tTDH3* A *pENO2-ENO2-tENO2* B *tHXK2-HXK2-pHXK2* C *tPGI1-PGI1-pPGI1* D *tPFK1-PFK1-pPFK1* J *tPFK2-PFK2-pPFK2* BU *tHIS3-HIS3-pHIS3* L *tGPM1-GPM1-pGPM1* M *tPDC1-PDC1-pPDC1* AR *ARS1211* BS 9CD *pTEF1-coAtCHS3-tMDH1 9CD* AQ Telomere |

### Supplementary Table 8 - Plasmids

**Table 8A gRNA plasmids**

| Plasmid | Relevant characteristics | Primer(s) used for gRNA | Source |
|---|---|---|---|
| **pMEL10** | 2µm ampR *KlURA3* gRNA-*CAN1*.Y | N.A. | Mans, *et al.* [9] |
| **pROS10** | 2µm ampR *URA3* gRNA-*CAN1.Y* gRNA-*ADE2.Y* | N.A. | Mans, *et al.* [9] |
| **pROS11** | 2µm ampR *amdSYM* gRNA-*CAN1*.Y gRNA-*ADE2*.Y | N.A. | Mans, *et al.* [9] |
| **pROS12** | 2µm ampR *hphNT1* gRNA-*CAN1.Y* gRNA-*ADE2.Y* | N.A. | Mans, *et al.* [9] |
| **pROS13** | 2µm ampR *kanMX* gRNA-*CAN1*.Y gRNA-*ADE2*.Y | N.A. | Mans, *et al.* [9] |
| **pUDR286** | 2µm ampR *URA3* gRNA-*TKL2* gRNA-*SOL4* | 9508 & 9503 | Postma, *et al.*[10] |
| **pUDR400** | 2µm ampR *hphNT1* gRNA-*mTurquoise* | 12911 & 12912 | This study |
| **pUDR406** | 2µm ampR *URA3* gRNA-*ARO10* gRNA-*PDC5/6* | 13614 & 7246 | This study |
| **pUDR413** | 2µm ampR *kanMX* gRNA- *RECYCLE SinLoG* | N.A. | Boonekamp, *et al.* [1] |
| **pUDR426** | 2µm ampR *KanMX* gRNA-*spHIS5 (2x)* | 10641 | This study |
| **pUDR546** | 2µm ampR *hphNT1* gRNA-*URA3* gRNA-*HIS3* | 14756 & 8314 | This study |
| **pUDR590** | 2µm ampR *amdS* gRNA-*NQM1* gRNA-*GND2* | 12569 &7231 | This study |
| **pUDR700** | 2µm ampR *KanMX* gRNA-*GND1* gRNA-*RKI1* | 16895 & 16897 | This study |
| **pUDR701** | 2µm ampR *KanMX* gRNA-*TAL1 (2x)* | 16909 | This study |
| **pUDR702** | 2µm ampR *hphNT1* gRNA-*RPE1* gRNA-*TKL1* | 16903 & 16905 | This study |
| **pUDR703** | 2µm ampR *hphNT1* gRNA-*ZWF1* gRNA-*SOL3* | 8564 & 16889 | This study |
| **pUDR756** | 2µm ampR *hphNT1* gRNA-*RKI1_WM (2x)* | 17613 | This study |
| **pUDR765** | 2µm ampR *hphNT1* gRNA-*Chunk16AB (2x)* | 17868 | This study |
| **pUDR771** | 2µm ampR *hphNT1* gRNA-*CAN1* gRNA-*X2* | 6008 & 10866 | This study |
| **pUDR772** | 2µm ampR *kanMX* gRNA-*YPRCtau3* gRNA-*SPR3* | 12985 & 12034 | This study |
| **pUDR780** | 2µm ampR *hphNT1* gRNA-*Chunk7BC* gRNA-*Chunk15CD)* | 18226 & 18277 | This study |
| **pUDR781** | 2µm ampR *kanMX* gRNA-*shrN* gRNA-*Chunk9CD)* | 18228 & 18299 | This study |

**Table 8B in-house golden gate part plasmids**

| Plasmid | Relevant characteristics | Source |
|---|---|---|
| **pUD565** | camR *GFP* entry vector | Boonekamp, *et al.*[1] |
| **pYTK012** | camR *pHHF2* | Lee, *et al.* [2] |
| **pYTK013** | camR *pTEF1* | Lee, *et al.* [2] |
| **pYTK015** | camR *pHHF1* | Lee, *et al.* [2] |
| **pYTK016** | camR *pHTB2* | Lee, *et al.* [2] |
| **pYTK017** | camR *pRPL18B* | Lee, *et al.* [2] |
| **pGGKp038** | camR *tTEF2* | Hassing, *et al.* [11] |
| **pGGKp039** | camR *tTEF1* | Hassing, *et al.* [11] |
| **pGGKp062** | kanR *pSkADH1* | Hassing, *et al.* [11] |
| **pGGKp063** | kanR *pSkTDH3* | Hassing, *et al.* [11] |
| **pGGkp074** | camR *pSePDC1* | Hassing, *et al.* [11] |
| **pGGKp075** | camR *pSeFBA1* | Hassing, *et al.* [11] |
| **pGGkp095** | camR *pSeGPM1* | Hassing, *et al.* [11] |
| **pGGKp113** | camR *tADH3* | Hassing, *et al.* [11] |
| **pGGKp119** | camR *coEcaroG*$^{(p150L)}$ | Hassing, *et al.* [11] |

| | | |
|---|---|---|
| **pGGKp120** | camR *coEcaroB* | Hassing, *et al.* [11] |
| **pGGKp121** | camR *coEcaroD* | Hassing, *et al.* [11] |
| **pGGKp122** | camR *coEcaroE* | Hassing, *et al.* [11] |
| **pGGKp123** | camR *coEcaroL* | Hassing, *et al.* [11] |
| **pGGKp124** | camR *coEcaroA* | Hassing, *et al.* [11] |
| **pGGKp125** | camR *coEcaroC* | Hassing, *et al.* [11] |
| **pGGKp126** | camR $coEcpheA^{T326P}$ | Hassing, *et al.* [11] |
| **pGGKp182** | camR *tCYC1* | This study |
| **pGGKp327** | camR *coRcTAL1* | This study |

**Table 8C Part plasmids subcloned by GeneArt in entry vector pUD565**

| Plasmid | Relevant characteristics | Source |
|---|---|---|
| pGGKp131 | camR *AtPAL1* | GeneArt |
| pGGKp135 | camR $coEctyrA^{M53I\ A354V}$ | GeneArt |
| pGGKp245 | ampR *coEctyrB* | GeneArt |
| pUD262 | camR *pCWP2* | GeneArt |
| pUD819 | camR *tACO1* | GeneArt |
| pUD826 | camR *tCIT1* | GeneArt |
| pUD829 | camR *tDIC1* | GeneArt |
| pUD832 | camR *tIDH2* | GeneArt |
| pUD836 | camR *tFUM1* | GeneArt |
| pUD841 | camR *tGDB1* | GeneArt |
| pUD842 | camR *tGLC3* | GeneArt |
| pUD844 | camR *tTAL1* | GeneArt |
| pUD845 | camR *tGPD2* | GeneArt |
| pUD846 | camR *tGPH1* | GeneArt |
| pUD848 | camR *tGSY2* | GeneArt |
| pUD850 | camR *tSDH2* | GeneArt |
| pUD851 | camR *tSDH3* | GeneArt |
| pUD852 | camR *tSDH4* | GeneArt |
| pUD858 | camR *tLAT1* | GeneArt |
| pUD862 | camR *tMDH1* | GeneArt |
| pUD873 | camR *tPGM2* | GeneArt |
| pUD875 | camR *tSOL3* | GeneArt |
| pUD876 | camR *tRKI1* | GeneArt |
| pUD879 | camR *tGND1* | GeneArt |
| pUD880 | camR *tSOL4* | GeneArt |
| pUD881 | camR *tRPE1* | GeneArt |
| pUD882 | camR *tTKL1* | GeneArt |
| pUD884 | camR *tZWF1* | GeneArt |
| pUD911 | camR *pTAL1* | GeneArt |
| pUD933 | camR *pSOL3* | GeneArt |
| pUD934 | camR *pRKI1* | GeneArt |
| pUD944 | camR *pGND1* | GeneArt |
| pUD946 | camR *pRPE1* | GeneArt |
| pUD947 | camR *pTKL1* | GeneArt |
| pUD949 | camR *pZWF1* | GeneArt |

**Table 8D Part plasmids ordered from GeneArt and subcloned in house in entry vector pUD565**

| Plasmid | Relevant characteristics | Source |
|---|---|---|
| **pUD1038** | ampR *ZWF1 ** | GeneArt |
| **pUD1039** | ampR *TKL1 ** | GeneArt |
| **pUD1040** | kanR *TAL1 ** | GeneArt |
| **pUD1041** | ampR *SOL3 ** | GeneArt |
| **pUD1042** | ampR *RPE1 ** | GeneArt |
| **pUD1043** | ampR *RKI1 ** | GeneArt |
| **pUD1044** | kanR *GND1 ** | GeneArt |
| **pGGKp245** | Amp *coEctyrB* | GeneArt |
| **pGGKp247** | camR *ZWF1 ** | This study |
| **pGGKp248** | camR *TKL1 ** | This study |
| **pGGKp249** | camR *TAL1 ** | This study |
| **pGGKp250** | camR *SOL3 ** | This study |
| **pGGKp251** | camR *RPE1 ** | This study |
| **pGGKp252** | camR *RKI1 ** | This study |
| **pGGKp253** | camR *GND1 ** | This study |
| **pGGKp293** | camR *coEctyrB* | This study |

**Table 8E Part plasmids made in house by PCR**

*****For *pSEPYK1* and initially the *coAtANS* no correct *E.coli* part plasmid transformant was found and expression cassettes were thus assembled with PCR fragments with yeast toolkit flanks.

| Plasmid | Relevant characteristics | Template | Primers | Source |
|---|---|---|---|---|
| **pUD257** | camR *pRPP0* | CEN.PK113-7D genomic DNA | 16294 & 16295 | This study |
| **pUD258** | camR *pRPL3* | CEN.PK113-7D genomic DNA | 16300 & 16301 | This study |
| **pUD259** | camR *pRPL8A* | CEN.PK113-7D genomic DNA | 16298 & 16299 | This study |
| **pUD260** | camR *pRPL10* | CEN.PK113-7D genomic DNA | 16296 & 16297 | This study |
| **pUD261** | camR *pRPL25* | CEN.PK113-7D genomic DNA | 16292 & 16293 | This study |
| **pGGKp324** | camR *AtCHI1* | pUDI065 | 17829 & 17830 | This study |
| **pGGKp325** | camR *coAtCHS3* | pUDE185 | 17827 & 17828 & 17834 & 17835 | This study |
| **pGGKp326** | camR *coAtC4H* | pUDE172 | 17823 & 17824 | This study |
| **pGGKp328** | camR *coGhDFR* | Genomic DNA PATW076 [12] | 17872 & 17873 | This study |
| **pGGKp329** | camR *coAt3GT* | Genomic DNA PATW076 [12] | 17876 & 17877 | This study |
| **pGGKp330** | camR *pSePGK1* | pUDI102 | 9413 & 9414 | This study |
| **pGGKp331** | camR *pSeENO2* | *Saccharomyces eubayanus* genomic DNA | 9743 & 9744 | This study |
| **pGGKp332** | CamR *coAtF3H* | Genomic DNA PATW076 | 17870 & 17871 | This study |
| **pGGKp340** | camR *coAtANS* | Genomic DNA PATW076 | 17874 & 17875 | This study |
| **PCR fragment*** | *coAtANS* | Genomic DNA PATW076 | 17874 & 17875 | This study |

| | | | | |
|---|---|---|---|---|
| **PCR fragment*** | *pSePYK1* | pUDI129 | 10610 & 10611 | This study |

**Table 8F in house expression plasmids described in other studies**

| Plasmid | Relevant characteristics | Source |
|---|---|---|
| **pGGKd005** | CEN6/ARS4 ampR *hphNT1 GFP* | Hassing, *et al.* [11] |
| **pGGKd012** | CEN6/ARS4 *cloNAT$^R$ GFP* | Boonekamp, *et al.* [1] |
| **pUDC212** | CEN6/ARS4 *ampR natNT2 pFBA1-FBA1*-tFBA1* | Boonekamp, *et al.* [1] |
| **pUDC213** | CEN6/ARS4 *ampR natNT2 pPGM1-PGM1*-tPGM1* | Boonekamp, *et al.* [1] |
| **pUDC214** | CEN6/ARS4 *ampR natNT2 pHXK2-HXK2*-tHXK2* | Boonekamp, *et al.*[1] |
| **pUDC215** | CEN6/ARS4 *ampR natNT2 pPDC1-PDC1*-tPDC1* | Boonekamp, *et al.* [1] |
| **pUDC216** | CEN6/ARS4 *ampR natNT2 pPFK1-PFK1*-tPFK1* | Boonekamp, *et al.*[1] |
| **pUDC217** | CEN6/ARS4 *ampR natNT2 pPFK2-PFK2*-tPFK2* | Boonekamp, Dashko [1] |
| **pUDC219** | CEN6/ARS4 *ampR natNT2 pPGK1-PGK1*-tPGK1* | Boonekamp, *et al.* [1] |
| **pUDC220** | CEN6/ARS4 *ampR natNT2 pPYK1-PYK1*-tPYK1* | Boonekamp, *et al.*[1] |
| **pUDC222** | CEN6/ARS4 *ampR natNT2 pTPI1-TPI1*-tTPI1* | Boonekamp, *et al.*[1] |
| **pUDC229** | CEN6/ARS4 *ampR natNT2 pADH1-ADH1*-tADH1* | Boonekamp, *et al.*[1] |
| **pUDC230** | CEN6/ARS4 *ampR natNT2 pTDH3-TDH3*-tTDH3* | Boonekamp, *et al.*[1] |
| **pUDC231** | CEN6/ARS4 *ampR natNT2 pENO2-ENO2*-tENO2* | Boonekamp, *et al.*[1] |
| **pUDC232** | CEN6/ARS4 *ampR natNT2 pPGI1-PGI1*-tPGI1* | Boonekamp, *et al.*[1] |

**Table 8G Expression plasmids constructed in this study**

*****For *pSEPYK1* and initially *coAtANS* no correct *E.coli* part plasmid transformant was found and expression cassettes were thus assembled with PCR fragments with yeast toolkit flanks

** the ORF of pUDC357 turned out to be mutated after sanger sequencing.

| Plasmid | Relevant characteristics | Parts used | Source |
|---|---|---|---|
| **pUDC275** | CEN6/ARS4 ampR *natNT2 pZWF1-ZWF1*-tZWF1* | pGGKd012, pUD949, pGGKp247, pUD884 | This study |
| **pUDC276** | CEN6/ARS4 ampR *natNT2 pTKL1-TKL1*-tTKL1* | pGGKd012, pUD947, pGGKp248, pUD882 | This study |
| **pUDC277** | CEN6/ARS4 ampR *natNT2 pGND1-GND1*-tGND1* | pGGKd012, pUD911, pGGKp253, pUD844 | This study |
| **pUDC278** | CEN6/ARS4 ampR *natNT2 pRKI1-RKI1*-tRKI1* | pGGKd012, pUD933, pGGKp252, pUD875 | This study |
| **pUDC279** | CEN6/ARS4 ampR *natNT2 pTAL1-TAL1*-tTAL1* | pGGKd012, pUD946, pGGKp249, PUD881 | This study |
| **pUDC280** | CEN6/ARS4 ampR *natNT2 pRPE1-RPE1*-tRPE1* | pGGKd012, pUD934, pGGKp251, pUD876 | This study |
| **pUDC281** | CEN6/ARS4 ampR *natNT2 pSOL3-SOL3*-tSOL3* | pGGKd012, pUD944, pGGKp250, pUD879 | This study |
| **pUDC293** | CEN6/ARS4 ampR *natNT2 pRPL3-coEcaroA-tSOL4* | pGGKd012, pUD258, pGGKp124, pUD880 | This study |
| **pUDC296** | CEN6/ARS4 ampR *natNT2 pRPL25-coEcaroD-tGPH1* | pGGKd012, pUD261, pGGKp121, pUD846 | This study |

| | | | |
|---|---|---|---|
| **pUDC297** | CEN6/ARS4 ampR *natNT2 pRPP0-coEcaroE-tCYC1* | pGGKd012, pUD257, pGGKp122, pGGKp182 | This study |
| **pUDC298** | CEN6/ARS4 ampR *natNT2 pHHF1-coEcaroG*$^{(p150L)}$*-tTEF1* | pGGKd012, pYTK015, pGGKp119, pGGKp039 | This study |
| **pUDC299** | CEN6/ARS4 ampR *natNT2 pHTB2-coEcaroL-tPGM2* | pGGKd012, pYTK016, pGGKp123, pUD873 | This study |
| **pUDC301** | CEN6/ARS4 ampR *natNT2 pRPL10-coEctyrA*$^{M53I\ A354V}$*-tGDB1* | pGGKd012, pUD260, pGGKp135, pUD841 | This study |
| **pUDC302** | CEN6/ARS4 ampR *natNT2 pCWP2-coEctyrB-tGLC3* | pGGKd012, pUD262, pGGKp293, pUD842 | This study |
| **pUDC294** | CEN6/ARS4 ampR *natNT2 pHHF2-coEcaroB-tTEF2* | pGGKd012, pYTK012, pGGKp120, pGGKp038 | This study |
| **pUDC295** | CEN6/ARS4 ampR *natNT2 pRPL8A-coEcaroC-tGPD2* | pGGKd012, pUD259, pGGKp125, pUD845 | This study |
| **pUDC300** | CEN6/ARS4 ampR *natNT2 pRPL18B-coEcpheA*$^{T326P}$*-tGSY2* | pGGKd012, pYTK017, pGGKp126, pUD848 | This study |
| **pUDC349** | CEN6/ARS4 ampR *natNT2 pSePDC1-AtPAL1-tLAT1* | pGGKd012, pGGkp074, pGGKp131, pUD858 | This study |
| **pUDC350** | CEN6/ARS4 ampR *natNT2 pSeGPM1-coRcTAL1-tCIT1* | pGGKd012, pGGkp095, pGGKp327, pUD826 | This study |
| **pUDC352** | CEN6/ARS4 ampR *natNT2 pTEF1-coAtCHS3-tMDH1* | pGGKd012, pYTK013, pGGKp325, pUD862 | This study |
| **pUDC353** | CEN6/ARS4 ampR *natNT2 pSkADH1-AtCHI-tSDH4* | pGGKd012, pGGKp062, pGGKp324, pUD852 | This study |
| **pUDC354** | CEN6/ARS4 ampR *natNT2 pSeFBA1-coAtC4H-tADH3* | pGGKd012, pGGKp075, pGGKp326, pGGKp113 | This study |
| **pUDC355** | CEN6/ARS4 ampR *natNT2 pSkTDH3-coAtF3H-tSDH3* | pGGKd012, pGGKp063, pGGKp332, pUD851 | This study |
| **pUDC356** | CEN6/ARS4 ampR *natNT2 pSePGK1-coGhDFR-tACO1* | pGGKd012, PGGKp330, pGGKp328, pUD819 | This study |
| **pUDC357**** | CEN6/ARS4 ampR *natNT2 pSeENO2-coAtANS-tFUM1* | pGGKd012, pGGKp331, *coAtANS* | This study |

| | | | |
|---|---|---|---|
| | | PCR fragment*, pUD836 | |
| **pUDC358** | CEN6/ARS4 ampR *natNT2 pSePYK1-coAt3GT-tDIC* | pGGKd012, *coAt3GT* PCR fragment *, pGGKp329, pUD829 | This study |
| **pUDC398** | CEN6/ARS4 ampR *natNT2 pSeENO2-coAtANS-tFUM1* | pGGKd012, pGGKp331, pGGKp340, pUD836 | This study |

**Table 8H Expression plasmids made by Gibson assembly in this study**

| Plasmid | Relevant characteristics | Source |
|---|---|---|
| pUDC348 | CEN6/ARS4 ampR *natNT2 pRPS3-coAtCPR1-tIDH2* | This study |
| pUDC351 | CEN6/ARS4 ampR *natNT2 pSeTPI1-At4CL3-tSDH2* | This study |

| Part | Source | Primers |
|---|---|---|
| | **pUDC348** | |
| ***pRPS3*** | CEN.PK113-7D genomic DNA | 17811 &17812 |
| ***coAtCPR1*** | pUDE172 | 17813 & 17814 |
| ***tIDH2*** | pUD832 | 17815 & 17816 |
| **Backbone** | pGGKd012 | 12377 & 12378 |
| | **pUDC351** | |
| ***pSeTPI1*** | pUDI116 | 17817, 17818 |
| ***At4CL3*** | pUDI065 | 17819, 17820 |
| ***tSDH2*** | pUD850 | 17821, 17822 |
| **Backbone** | pGGKd012 | 12377, 12378 |

**Table 8I Other plasmids**

| Plasmid | Relevant characteristics | Source |
|---|---|---|
| **pUDI065** | Integration plasmid *LEU2 pTDH3-AtCHI1-tCYC1 pTPI-AtCHS3-tADH pTEF-At4CL3-tTEF PYK2*(1-710) | Koopman, *et al.*[13] |
| **pUDI102** | *pSePGK1-mRuby2-tENO2* | Boonekamp, *et al.*[5] |
| **pUDI116** | *pSeTPI1-mRuby2-tENO2* | Boonekamp, *et al.*[5] |
| **pUDI129** | *pSePYK1-mRuby2-tENO2* | Boonekamp, *et al.*[5] |
| **pUDE172** | *CEN6/ARS4, URA3, pTDH3-AtPAL1-tCYC1, pTPI-coC4H-tADH, pPGI-coCPR1-tPGI* | Koopman, *et al.*[13] |
| **pUDE185** | 2 µm *HIS3 pTDH3-coCHS3-tCYC1* | Koopman, *et al.*[13] |
| **pLM092** | CEN6/ARS4, ampR, *HIS3*, 5'*URA3-ACT1*intron[Tess-I-SceI-Tess]-3'*URA3* | Mitchell and Boeke [14] |
| **pUDC191** | CEN6/ARS4, ampR, *URA3, pCCW12-mRuby2-tENO1* | Postma, *et al.*[8] |
| **pUDC192** | CEN6/ARS4, ampR, *URA3, pTEF2-mTurquoise2-tSSA1* | Postma, *et al.*[8] |

### Supplementary Table 9 - pROS/pMEL gRNA primers.

gRNA sequence is underlined.

| Primer number | Primer name | Sequence (5' to 3') |
|---|---|---|
| **6008** | CAN1_targetRNA FW | GTGCGCATGTTTCGGCGTTCGAAACTTCTCCGCAGTGAAAGATAAATGATCGATACGTTCTCTATGGAGGAGTTTTAGAGCTAGAAATAGCAAGTTAAAATAAG |
| **9508** | TKL2_targetRNA FW | TGCGCATGTTTCGGCGTTCGAAACTTCTCCGCAGTGAAAGATAAATGATCTCAAAAACTTAATGAGGAATGTTTTAGAGCTAGAAATAGCAAGTTAAAATAAG |
| **7231** | RV_gnd2_gRNA | GTTGATAACGGACTAGCCTTATTTTAACTTGCTATTTCTAGCTCTAAAACTATGATCTGGCAGCTTCGCGGATCATTTATCTTTCACTGCGGAGAAGTTTCGAACGCCGAAACATGCGCA |
| **7246** | ARO10 CRISPR KO seq | TGCGCATGTTTCGGCGTTCGAAACTTCTCCGCAGTGAAAGATAAATGATCATTTACAAGTATTCTAAACCGTTTTAGAGCTAGAAATAGCAAGTTAAAATAAGGCTAGTCCGTTATCAAC |
| **8314** | Sc_URA3_2gRNA_primer | TGCGCATGTTTCGGCGTTCGAAACTTCTCCGCAGTGAAAGATAAATGATCTTGACTGATTTTTCCATGGAGTTTTAGAGCTAGAAATAGCAAGTTAAAATAAG |
| **8564** | ZWF1_targetRNA FW | TGCGCATGTTTCGGCGTTCGAAACTTCTCCGCAGTGAAAGATAAATGATCTTAGATTCAGATCTGTGACTGTTTTAGAGCTAGAAATAGCAAGTTAAAATAAG |
| **9503** | SOL4_targetRNA FW | TGCGCATGTTTCGGCGTTCGAAACTTCTCCGCAGTGAAAGATAAATGATCACATTTTTCCACATATTAAGGTTTAGAGCTAGAAATAGCAAGTTAAAATAAG |
| **10641** | Sphis5_targetRNA FW | TGCGCATGTTTCGGCGTTCGAAACTTCTCCGCAGTGAAAGATAAATGATCTTCCAAGCATGCAAACCAAAGTTTTAGAGCTAGAAATAGCAAGTTAAAATAAGGCTAGTCCGTTATCAAC |
| **10866** | X2_targetRNA FW | TGCGCATGTTTCGGCGTTCGAAACTTCTCCGCAGTGAAAGATAAATGATCGGCGACTAGGAAGAGAGTAGGTTTTAGAGCTAGAAATAGCAAGTTAAAATAAG |
| **12034** | SPR3_targetRNA FW | TGCGCATGTTTCGGCGTTCGAAACTTCTCCGCAGTGAAAGATAAATGATCATGCTTTTATAACGAATAATGTTTTAGAGCTAGAAATAGCAAGTTAAAATAAGGCTAGTCCGTTATCAAC |
| **12569** | NQM1_targetRNA RV | GTTGATAACGGACTAGCCTTATTTTAACTTGCTATTTCTAGCTCTAAAACTCTAGAACAGTTATATGAATGATCATTTATCTTTCACTGCGGAGAAGTTTCGAACGCCGAAACATGCGCA |
| **12911** | mTurquoise2_gRNA1_fw | TGCGCATGTTTCGGCGTTCGAAACTTCTCCGCAGTGAAAGATAAATGATCTACTGCTGCTGGTATTACCTGTTTTAGAGCTAGAAATAGCAAGTTAAAATAAG |
| **12912** | mTurquoise2_gRNA2_fw | TGCGCATGTTTCGGCGTTCGAAACTTCTCCGCAGTGAAAGATAAATGATCCCTTAGTCACTACTTTATCTGTTTTAGAGCTAGAAATAGCAAGTTAAAATAAG |
| **12985** | YPRCtau3_targetRNA FW | TGCGCATGTTTCGGCGTTCGAAACTTCTCCGCAGTGAAAGATAAATGATCAAACATTCAAATATATTCCAGTTTTAGAGCTAGAAATAGCAAGTTAAAATAAG |

| | | |
|---|---|---|
| **13614** | PDC5_PDC6_targetRNA FW | TGCGCATGTTTCGGCGTTCGAAACTTCTCCGCAGTGAAAGATAAATGATCATTGTTGTTGCATCATACCTGTTTTAGAGCTAGAAATAGCAAGTTAAAATAAG |
| **14756** | HIS3_targetRNA2 FW | TGCGCATGTTTCGGCGTTCGAAACTTCTCCGCAGTGAAAGATAAATGATCTTAACGTCCACACAGGTATAGTTTTAGAGCTAGAAATAGCAAGTTAAAATAAG |
| **16889** | SOL3_targetRNA FW | TGCGCATGTTTCGGCGTTCGAAACTTCTCCGCAGTGAAAGATAAATGATCCTCATGCATTATATTTTGTTGTTTTAGAGCTAGAAATAGCAAGTTAAAATAAGGCTAGTCCGTTATCAAC |
| **16895** | GND1_targetRNA FW | TGCGCATGTTTCGGCGTTCGAAACTTCTCCGCAGTGAAAGATAAATGATCTTACGAAGAATTGAAGAAGAGTTTTAGAGCTAGAAATAGCAAGTTAAAATAAGGCTAGTCCGTTATCAAC |
| **16897** | RKI1_targetRNA FW | TGCGCATGTTTCGGCGTTCGAAACTTCTCCGCAGTGAAAGATAAATGATCAATGCGAGGATACTGTTCAAGTTTTAGAGCTAGAAATAGCAAGTTAAAATAAGGCTAGTCCGTTATCAAC |
| **16903** | RPE1_targetRNA FW | TGCGCATGTTTCGGCGTTCGAAACTTCTCCGCAGTGAAAGATAAATGATCCGACTTGGATATTCAAATGGGTTTTAGAGCTAGAAATAGCAAGTTAAAATAAGGCTAGTCCGTTATCAAC |
| **16905** | TKL1_targetRNA FW | TGCGCATGTTTCGGCGTTCGAAACTTCTCCGCAGTGAAAGATAAATGATCTAACCCAGATATTATTTTAGGTTTTAGAGCTAGAAATAGCAAGTTAAAATAAGGCTAGTCCGTTATCAAC |
| **16909** | TAL1_targetRNA FW | TGCGCATGTTTCGGCGTTCGAAACTTCTCCGCAGTGAAAGATAAATGATCAACTAACCCATCATTGATCTGTTTTAGAGCTAGAAATAGCAAGTTAAAATAAGGCTAGTCCGTTATCAAC |
| **17613** | RKI1_targetRNA_SNP1_FW | TGCGCATGTTTCGGCGTTCGAAACTTCTCCGCAGTGAAAGATAAATGATCATGTTTTGGGGACTTTTTTCGTTTTAGAGCTAGAAATAGCAAGTTAAAATAAGGCTAGTCCGTTATCAAC |
| **18010** | SHR AM_targetRNA _FW | TGCGCATGTTTCGGCGTTCGAAACTTCTCCGCAGTGAAAGATAAATGATCACTCGTATCTTACATGACGTGTTTTAGAGCTAGAAATAGCAAGTTAAAATAAGGCTAGTCCGTTATCAAC |
| **18266** | Chunk7BC_targetRNA _FW | TGCGCATGTTTCGGCGTTCGAAACTTCTCCGCAGTGAAAGATAAATGATCTTTGCGGAATATCGACCACGGTTTTAGAGCTAGAAATAGCAAGTTAAAATAAG |
| **18227** | Chunk15CD_targetRNA _FW | TGCGCATGTTTCGGCGTTCGAAACTTCTCCGCAGTGAAAGATAAATGATCATATAAGTGTCCAGCCAGAGGTTTTAGAGCTAGAAATAGCAAGTTAAAATAAG |
| **18228** | shrN_targetRNA _FW | TGCGCATGTTTCGGCGTTCGAAACTTCTCCGCAGTGAAAGATAAATGATCTCTTCGTTAGGACTCAATCGGTTTTAGAGCTAGAAATAGCAAGTTAAAATAAG |
| **18229** | Chunk9CD_targetRNA _FW | TGCGCATGTTTCGGCGTTCGAAACTTCTCCGCAGTGAAAGATAAATGATCCAGATCAAAATCCACCAGTGGTTTTAGAGCTAGAAATAGCAAGTTAAAATAAG |

### Supplementary Table 10 - Primers to check correct construction of gRNA plasmids

| Primer number | Primer name | Sequence (5' to 3') |
|---|---|---|
| **7257** | RV_gnd2_gRNA_check | TATGATCTGGCAGCTTCGCG |
| **9708** | TKL2_beta_dg rv | ATTCCTCATTAAGTTTTTGA |
| **9709** | SOL4_alpha_dg rv | CTTAATATGTGGAAAAATGT |
| **12684** | DG spHIS5 targetRNA | TTTGGTTTGCATGCTTGGAAG |
| **12729** | NQM1_pMEL_dg fw | ATTCATATAACTGTTCTAGA |
| **13040** | YPRCtau3_pROS_dg rv | CTGGAATATATTTGAATGTTTGAT |
| **13263** | PDC5 and 6 gRNA dg | AGGTATGATGCAACAACAATG |
| **13264** | ARO10 gRNA DG | CGGTTTAGAATACTTGTAAAT |
| **14602** | zwf1 gRNA dg RV | AGTCACAGATCTGAATCTAAG |
| **14757** | HIS3_pROS_dg rv | TATACCTGTGTGGACGTTAA |
| **14758** | URA3_pROS_dg fw | TCCATGGAAAAATCAGTCAA |
| **15821** | X2_pROS_dg rv | CTACTCTCTTCCTAGTCGCC |
| **15968** | SPR3_pROS_dg rv | CATTATTCGTTATAAAAGCATGAT |
| **16890** | SOL3_pROS_dg rv | AACAAAATATAATGCATGAGGATC |
| **16896** | GND1_pROS_dg rv | TCTTCTTCAATTCTTCGTAAGAT |
| **16898** | RKI1_pROS_dg rv | TTGAACAGTATCCTCGCATT |
| **16904** | RPE1_pROS_dg rv | CCATTTGAATATCCAAGTCGG |
| **16906** | TKL1_pROS_dg rv | CTAAAATAATATCTGGGTTAGATCA |
| **16910** | TAL1_pROS_dg rv | AGATCAATGATGGGTTAGTTG |
| **17621** | RKI1_SNP1_pROS_dg rv | CGAAAAAAGTCCCCAAAACATG |
| **18010** | SHR_AM_pROS_dg rv | CACGTCATGTAAGATACGAGTG |
| **18012** | CAN1_pROS_dg rv | CTCCTCCATAGAGAACGTATCG |
| **18230** | diag_gRNA_chunk7BC rv | CGTGGTCGATATTCCGCAAAG |
| **18231** | diag_gRNA_chunk15CD rv | CCTCTGGCTGGACACTTATATG |
| **18232** | diag_gRNA_SHR-N rv | CCGATTGAGTCCTAACGAAGAG |
| **18233** | diag_gRNA_chunk9CD rv | CCACTGGTGGATTTTGATCTGG |

### Supplementary Table 11 - Primers to make golden gate part plasmids and expression plasmids with Gibson assembly

| Primer number | Primer name | Sequence (5’ to 3’) |
|---|---|---|
| **9413** | PGK1 se prom fw Ytoolkit | AAGCATCGTCTCATCGGTCTCAAACGGCTTCAATTCAAGATACACAGATATAC |
| **9414** | PGK1 se prom rev Ytoolkit | TTATGCCGTCTCAGGTCTCACATATGTTTTATATTTGTTGCAAAAAGTAG |
| **9743** | ENO1 se prom fw Ytoolkit | AAGCATCGTCTCATCGGTCTCAAACGCCAAGAAGATGCCGGCTAC |
| **9744** | ENO1 se prom rev Ytoolkit | TTATGCCGTCTCAGGTCTCACATATATTATTGTTTGATATAGTATTAGTTGCTT GGT |
| **10610** | PYK1 se prom fw Ytoolkit | AAGCATCGTCTCATCGGTCTCAAACGTGTAAATACCGGTTTTAGCC |
| **10611** | PYK1 se prom rv Ytoolkit | TTATGCCGTCTCAGGTCTCACATATGTGATGATGTTTTATTTGTTTTG |
| **16292** | GG_pRPL25_fw | GCATCGTCTCATCGGTCTCAAACGAGGTATGTTAGTGCTAAAAGCAAAATG |
| **16293** | GG_pRPL25_rv | ATGCCGTCTCAGGTCTCACATATTTATCTTATTGATCTTCTTTGTTTAGCCTTTT C |
| **16294** | GG_pRPP0_fw | GCATCGTCTCATCGGTCTCAAACGTTCAACAATTCGTTATATATATGGTAGGCT |
| **16295** | GG_pRPP0_rv | ATGCCGTCTCAGGTCTCACATATTCAAACTTATTATACGTATTTATTAGACTGTT |
| **16296** | GG_pRPL10_fw | GCATCGTCTCATCGGTCTCAAACGTCACTTGTCTGTGTGTTTAACTGCC |
| **16297** | GG_pRPL10_rv | ATGCCGTCTCAGGTCTCACATACTTGAATTAGTTATTTGATATACTGTACTT |
| **16298** | GG_pRPL8A_fw | GCATCGTCTCATCGGTCTCAAACGACATAAATAATTTCTATTAACAATGTAATT TC |
| **16299** | GG_pRPL8A_rv | ATGCCGTCTCAGGTCTCACATATTCGAATTAGTTGTTTTGATGTG |
| **16300** | GG_pRPL3_fw | GCATCGTCTCATCGGTCTCAAACGAGAGTCTTGGAGATTTTCGACCTG |
| **16301** | GG_pRPL3_rv | ATGCCGTCTCAGGTCTCACATAGATTGATTGTTGTAGTAACTGTGTTGTTC |
| **12377** | Backbone pGGKd017 FW | AAATCTGCTCGTCAGTGGTG |
| **12378** | Backbone pGGKd017 REV | ATTGCGACGAATTGCCACG |
| **17811** | pGGKd012-pRPS3 GA fw | CGACAACGTGGCAATTCGTCGCAATTCTGCTACTTTCCATTATCTGG |
| **17812** | coAtCPR1-pRPS3 GA rv | AGCAGAAGTCATTTTTGTAGTTTGTTTGCTGTTTTATTTTC |
| **17813** | pRPS3-coAtCPR1 GA fw | ACAAACTACAAAAATGACTTCTGCTTTGTACGC |
| **17814** | tIDH2-coAtCPR1 GA rv | AAGAATAGGACTTTTACCAAACGTCTCTCAAGTATC |
| **17815** | *ATCPR1*-tIDH2 GA fw | GACGTTTGGTAAAAGTCCTATTCTTTTCCCTCTC |

| | | |
|---|---|---|
| **17816** | pGGKd012-tIDH2 GA rv | GTGAGCACCACTGACGAGCAGATTTTCCACTGAGGGACATTTTG |
| **17817** | pGGKd012-pSeTPI1 GA fw | CGACAACGTGGCAATTCGTCGCAATGGATGTCGTTGTTCTTGTTAC |
| **17818** | At4CL3-pSeTPI1 GA rv | TGCAGTGATCATTTTTAGTGTATGTGTATGTGTGTTTG |
| **17819** | SeTPI1-At4CL3 GA fw | ACATACACTAAAAATGATCACTGCAGCTCTAC |
| **17820** | tSDH2-At4CL3 GA rv | TTTTTCTGATAGTTCAACAAAGCTTAGCTTTGAG |
| **17821** | At4CL3-tSDH2 GA fw | AAGCTTTGTTGAACTATCAGAAAAACAGCTAGCC |
| **17822** | pGGkd012-tSDH2 GA rv | GTGAGCACCACTGACGAGCAGATTTAAGCCAAAAGGCCCTTCAAAAAC |
| **17823** | coAtC4H YTKpart fw | GCATCGTCTCATCGGTCTCATATGGACTTGTTGTTGTTGGAAAAGTC |
| **17824** | coAtC4H YTKpart rv | ATGCCGTCTCAGGTCTCAGGATTTAACAGTTTCTTGGCTTCATAACG |
| **17827** | coAtCHS3 YTKpart fw | GCATCGTCTCATCGGTCTCATATGGTTATGGCTGGTGCTTCTTC |
| **17828** | coAtCHS3 YTKpart rv | ATGCCGTCTCAGGTCTCAGGATTTACAATGGAACAGAGTGCAAAAC |
| **17829** | AtCHI1 YTKpart fw | GCATCGTCTCATCGGTCTCATATGATGTCTTCATCCAACGCCTGCG |
| **17830** | AtCHI1 YTKpart rv | ATGCCGTCTCAGGTCTCAGGATTCAGTTCTCTTTGGCTAGTTTTTCCTC |
| **17834** | CHS4 internal BsaI removal RV | CACGTCTCACCTTAAACCCAACAACTTAGTCAA |
| **17835** | CHS4 internal BsaI removal FW | TTCGTCTCTAAGGCCATCTGTTAAGAGATTGATGATGTA |
| **17870** | coAtF3H YTKpart fw | GCATCGTCTCATCGGTCTCATATGGCTCCAGGTACTTTGAC |
| **17871** | coAtF3H YTKpart rv | ATGCCGTCTCAGGTCTCAGGATTTAAGCGAAGATTTGGTCAACTG |
| **17872** | coGhDFR YTKpart fw | GCATCGTCTCATCGGTCTCATATGGAAGAAGACTCTCCAGCTACTGTTTG |
| **17873** | coGhDFR YTKpart rv | ATGCCGTCTCAGGTCTCAGGATCTATTGACCTTCCTTAGAACAACACAAC |
| **17874** | *coAtANS* YTKpart fw | GCATCGTCTCATCGGTCTCATATGGTTGCTGTTGAAAGAGTTGAATC |
| **17875** | *coAtANS* YTKpart rv | ATGCCGTCTCAGGTCTCAGGATTTAGTCGTTCTTTTCAGAAACCAATTC |
| **17876** | coAt3GT YTKpart fw | GCATCGTCTCATCGGTCTCATATGACTAAGCCATCTGACCCAACTAGAG |
| **17877** | coAt3GT YTKpart rv | ATGCCGTCTCAGGTCTCAGGATTTAGATGATGTTAACAACAGCGTCCAAC |

### Supplementary Table 12 - Diagnostic primers to check golden gate part plasmids

| Primer number | Primer name | Sequence (5' to 3') |
|---|---|---|
| 1642 | MF fbas | TTTCCCAGTCACGACGTTG |
| 2012 | m132 | GGAAACAGCTATGACCATG |
| 2397 | FW pMA-RQ | AGACCGAGATAGGGTTGAGTG |
| 4941 | I-sceI inside rv n | |
| 5394 | TKL1 fw (ol pTHD3) | CGAATAAACACACATAAACAAACAAAATGACTCAATTCACTGACATTGAT AAGC |
| 7613 | FW_gnd1_inside | GATTGGTTTGGCCGTCATGG |
| 7868 | RPE1_F | CAACTTGGGTTGCGAATGTC |
| 7869 | RKI1_F | CTTTGGGCAATCCTTTGGAG |
| 7871 | TAL1_F | GGTGATTTCGGCTCTATTGC |
| 8953 | FW_pADH1_ZWF1 | CCAAGCATACAATCAACTATCTCATATACAATGAGTGAAGGCCCCGTCAA |
| 8988 | FW_diag_3'_SOL3 | TTGGGCTGTGGTCCTGATGG |
| 12611 | pUD565_fw1 | |

### Supplementary Table 13 - Diagnostic primers to check golden gate and Gibson assembly expression plasmids, for PCR and Sanger sequencing

| Primer number | Primer name | Sequence (5' to 3') |
|---|---|---|
| 1047 | TAL1Fw1 | CTGTACACTAGGAAGCCCTGTT |
| 1858 | FK050 | CGGATGGATGTCTCAAAC |
| 2122 | BG26-DF | GCTGCAGTATTGTTCCTGAG |
| 2123 | BG26-DR | CCTGTTTGCCTTTCCTTACG |
| 2557 | FK117-MP1 | GTGGACGCTATGTTATGC |
| 2558 | FK118-MP2 | AGTCTCACCACCAAGATTC |
| 2559 | FK119-MP3 | CCAGGTCCAATCCCAATC |
| 2560 | FK120-MP4 | CAGTGTTCACCGTAAACAG |
| 2561 | FK121-MP5 | TCTGCTTTGTACGCTTCTG |
| 2562 | FK122-MP6 | CGTATTGGTCGTCGTCAG |
| 2564 | FK124-MP8 | GATGAAGAACGCTGTTCC |
| 4494 | RPE1 DG fw | TATCCAAGTCGAGCTGGGAAAG |
| 4495 | RPE1 DG rv | CCCATGAGTTAGGCACTTACG |
| 5598 | TKL1 DG fw | CGTTCCGTTCGCAATCTC |
| 5599 | TKL1 DG rv | GGTGTGATTCTCTCGAAGG |
| 5807 | DT2 | ATGTCTTCATCCAACGCC |
| 6023 | CHI knockout cassette rv | CAGTTCTCTTTGGCTAGTTTTTC |
| 8566 | FW_zwf1_outside | GGGTGGCGAATTCTTCAATG |
| 10335 | ConRE Rv | GGCTGTCTTGCTTAGTTGTG |
| 12220 | ABZ1_SNO1_THI4 REV | GTGTGGTTCATGGGTGCGTTAGTCATCGGTATGATCTGTACATG |
| 12612 | PAL1_fw2 | CAGTTCTCTTTGGCTAGTTTTTC |
| 12614 | PAL1_rv1 | TCGAATCTAACCGCTTCGAG |
| 12615 | PAL1_rv2 | ACAAATCGCAACGAGGAACG |
| 13483 | ConL_pGGK_fw | TCTCCAGGACCATCTGAATC |
| 13668 | pGPM1 s.e dg fw | GAGGGCGGTTCTCATATTTC |
| 14788 | nadABhigh_seq2_fw | GCCGATAATTGCAGACGAAC |
| 17634 | TDH3p pGGKd017 fw | CTGGCCGATAATTGCAGACG |
| 17636 | CYC1t pGGKd017 rv | GATTTCCGTCTCATGCTCAG |
| 17819 | SeTPI1-At4CL3 GA fw | ACATACACTAAAAATGATCACTGCAGCTCTAC |
| 17820 | tSDH2-At4CL3 GA rv | TTTTTCTGATAGTTCAACAAAGCTTAGCTTTGAG |
| 17948 | coAtCHS3 dg fw | TATCGACGGTCACTTGAGAG |
| 17949 | coAtCHS3 dg rv | CGTGTCTAGTAGCTCTCATC |
| 17975 | coGhDFR dg rv1 | AAACCAGCAGCACCAGTAAC |
| 17976 | coGhDFR dg rv2 | CACAAGTCGTCCAAGTGAAC |
| 17977 | coGhDFR dg fw | AGTTGTGGAAGGCTGACTTG |
| 17978 | pSkTDH3 dg fw1 | CGGACATAACCTCAATGGAGTG |
| 17979 | coAtF3H dg rv1 | GTCTGGTTGTGGACACTTTG |
| 17980 | coAtF3H dg fw1 | ACGCTTGTGTTGACATGG |

### Supplementary Table 14 - List of NeoChr10 and NeoChr11 chromosome parts

* Size of the fragments does not include the SHR sequences.

| SHR 5’ | part | SHR 3’ | Size (bp)* | Template | Primer Fw | Primer Rv |
|---|---|---|---|---|---|---|
| | Telomere left | AO | 813 | pLM092 | 13395 | 13396 |
| AO | Chunk 7A | BJ | 2472 | *E.coli* BL21/*E.coli* XL1 Blue | 10004 | 11535 |
| BJ | Chunk 7B | BK | 2435 | *E.coli* BL21/*E.coli* XL1 Blue | 11536 | 11537 |
| BK | Chunk 7C | BL | 2478 | *E.coli* BL21/*E.coli* XL1 Blue | 11538 | 11539 |
| BL | Chunk 7D | AP | 2526 | *E.coli* BL21/*E.coli* XL1 Blue | 11540 | 10005 |
| AP | Chunk 8A | BM | 2477 | *E.coli* BL21/*E.coli* XL1 Blue | 10006 | 11541 |
| BM | Chunk 8B | BN | 2567 | *E.coli* BL21/*E.coli* XL1 Blue | 11542 | 11543 |
| BN | Chunk 8C | BO | 2454 | *E.coli* BL21/*E.coli* XL1 Blue | 11544 | 11545 |
| BO | Chunk 8D | AC | 2452 | *E.coli* BL21/*E.coli* XL1 Blue | 11546 | 10007 |
| AC | *mRuby2* | AD | 1667 | pUDC191 | 11365 | 11366 |
| AD | *ARS1* | AN | 56 | Annealing of complementary primers | 13397 | 9989 |
| AN | Chunk 18A | BP | 2525 | *E.coli* BL21/*E.coli* XL1 Blue | 13398 | 13399 |
| BP | Chunk 18B | BQ | 2510 | *E.coli* BL21/*E.coli* XL1 Blue | 13400 | 13401 |
| BQ | Chunk 19C | BR | 2495 | *E.coli* BL21/*E.coli* XL1 Blue | 13509 | 13510 |
| BR | Chunk 19D | DE | 2496 | *E.coli* BL21/*E.coli* XL1 Blue | 13511 | 13512 |
| DE | Chunk 15A | DF | 2515 | *E.coli* BL21/*E.coli* XL1 Blue | 13406 | 13407 |
| DF | Chunk 15B | DH | 2504 | *E.coli* BL21/*E.coli* XL1 Blue | 13408 | 13409 |
| DH | Chunk 15C | DI | 2489 | *E.coli* BL21/*E.coli* XL1 Blue | 13410 | 13411 |
| DI | Chunk 15D | DJ | 2497 | *E.coli* BL21/*E.coli* XL1 Blue | 13412 | 13413 |
| DJ | *CEN6/ARS4* | AE | 519 | pLM092 | 13414 | 9991 |
| AE | Chunk 16A | DK | 2520 | *E.coli* BL21/*E.coli* XL1 Blue | 13415 | 13416 |
| DK | Chunk 16B | DL | 2470 | *E.coli* BL21/*E.coli* XL1 Blue | 13417 | 13418 |
| DL | Chunk 16C | DM | 2517 | *E.coli* BL21/*E.coli* XL1 Blue | 13419 | 13420 |
| DM | Chunk 16D | DN | 2520 | *E.coli* BL21/*E.coli* XL1 Blue | 13421 | 13422 |
| DN | Chunk 17A | DO | 2499 | *E.coli* BL21/*E.coli* XL1 Blue | 13423 | 13424 |
| DO | Chunk 17B | DP | 2498 | *E.coli* BL21/*E.coli* XL1 Blue | 13425 | 13426 |
| DP | Chunk 19A | DQ | 2509 | *E.coli* BL21/*E.coli* XL1 Blue | 13427 | 13428 |
| DQ | Chunk 17D | DR | 2543 | *E.coli* BL21/*E.coli* XL1 Blue | 13429 | 13430 |
| DR | *ARS417* | BU | 60 | Annealing of complementary primers | 13431 | 11508 |
| BU | *HIS3* | AJ | 1250 | pLM092 | 11509 | 11032 |
| AJ | Chunk 4A | BC | 2526 | *E.coli* BL21/*E.coli* XL1 Blue | 9998 | 11510 |
| BC | Chunk 4B | BD | 2488 | *E.coli* BL21/*E.coli* XL1 Blue | 11511 | 11512 |
| BD | Chunk 4C | BE | 2470 | *E.coli* BL21/*E.coli* XL1 Blue | 11513 | 11514 |
| BE | Chunk 4D | AK | 2419 | *E.coli* BL21/*E.coli* XL1 Blue | 11515 | 9999 |
| AK | Chunk 9.2A | BF | 2405 | *E.coli* BL21/*E.coli* XL1 Blue | 10008 | 11516 |
| BF | Chunk 9.2B | BS | 2565 | *E.coli* BL21/*E.coli* XL1 Blue | 11517 | 11518 |

| | | | | | | |
|---|---|---|---|---|---|---|
| BS | Chunk 9.2C | BT | 2487 | *E.coli* BL21/*E.coli* XL1 Blue | 11519 | 11520 |
| BT | Chunk 9.2D | AQ | 2502 | *E.coli* BL21/*E.coli* XL1 Blue | 11521 | 11522 |
| AQ | Chunk 5A | BG | 2485 | *E.coli* BL21/*E.coli* XL1 Blue | 10000 | 11523 |
| BG | Chunk 5B | BH | 2521 | *E.coli* BL21/*E.coli* XL1 Blue | 11524 | 11525 |
| BH | Chunk 5C | BI | 2513 | *E.coli* BL21/*E.coli* XL1 Blue | 11526 | 11527 |
| BI | Chunk 5D | AL | 2441 | *E.coli* BL21/*E.coli* XL1 Blue | 11528 | 10001 |
| AL | *mTurquoise2* | DS | 1683 | pUDC192 | 13432 | 13433 |
| DS | Telomerator | | 807 | pLM092 | 13434 | 13435 / 13436 |

### Supplementary Table 15 - List of primers for amplifying NeoChr10 and NeoChr11 chromosome parts

| Primers number | Primer Name | Sequence 5' - 3' |
|---|---|---|
| **9989** | ARS1_rv | ATACATCATGCACGCCTGAAAGCATCCCTGACGCGAGTATGACGCAGTTCACACATCTTACTTGTTATTTTACAGATTTTATGTTTAGATCTTTTATGCTTGCTTTTCAAAA |
| **9991** | CEN6_ARS4_rv | CAACGCATGAGGATGATGACAGCAGCACTCGTACCAGATAGAGACAGCTCTTCCGAACATGGACGGATCGCTTGCCTGTAAC |
| **9998** | Ecoli_ch4_fw | CACACGCACGAATTTGCATACAGATAGTTTGAGACACTCGCACGATGGCCGATATTGCGTCCGCGCATATTCCAGCAAGGAG |
| **9999** | Ecoli_ch4_rv | TCAGACAATTCTATACGCGGACTGATATGGCAGAAGCTAGGAGACGTTATGCGATCTTAGTGGTACGGTTTACTCCTTCACCTGTG |
| **10000** | Ecoli_ch5_fw | GCGCAGAAGGCAATGCTATACATCTGATTGAAGCAGCGTCGCGCGTGCATCATTCCTTATCTGGGGAAACTGGCGAGCG |
| **10001** | Ecoli_ch5_rv | GCGTCCTCTGTATTAAGATGTCATGGTGTTGAGTCTGCACATGCAATGGCAATGAGTCAGTTAGCCCGCAGTGGATCCTCC |
| **10004** | Ecoli_ch7_fw | TCATTGATGCCAGGTACGTGGCTAATCTGAAATCCCAGCAACATGGTTGAAAGCGCGTCAATGGTTAGCTTCCCGCTCATTCTCC |
| **10005** | Ecoli_ch7_rv | CTCAAGGCTGTGCTGACGTAGGACTGATTCGAGCTATCTCTTGGTGTATTTGAGCAGACCCAGACATGGCGTAACCCCGTG |
| **10006** | Ecoli_ch8_fw | GGTCTGCTCAAATACACCAAGAGATAGCTCGAATCAGTCCTACGTCAGCACAGCCTTGAGGCATTTACGCTGTACGGACACCTT |
| **10007** | Ecoli_ch8_rv | GCTACATCTTCCGTACTATGCTGTAGTCTCATGGTCGAGTTCTATTGCTGTTCGGCGGCAGGGCAATATTACCAACCCGTTTTGC |
| **10008** | Ecoli_ch9_fw | CTAAGATCGCATAACGTCTCCTAGCTTCTGCCATATCAGTCCGCGTATAGAATTGTCTGACATTACCGGCGTAGCCCCATG |
| **11032** | terHIS3_rv | ACGCAATATCGGCCATCGTGCGAGTGTCTCAAACTATCTGTATGCAAATTCGTGCGTGTGGCATCTGTGCGGTATTTCACAC |
| **11365** | prCCW12_mRuby_tENO1_fw | TGCCGCCGAACAGCAATAGAACTCGACCATGAGACTACAGCATAGTACGGAAGATGTAGCAACGCACCCATGAACCACAC |
| **11366** | prCCW12_mRuby_tENO1_rv | CTCCACTGTACTGCATGTAGCATTCGCCGATCTGCATGATGTGTGACATTCTGCTATCGGGGCAGCATACATGGGTGACCAAA |
| **11508** | ARS417_rv | AATCATGTGACCCAGGCTTGCGCATACATGATCCTTCTTGCGCTGCATGGGCGACTATATCTTACGCTCAATTCCTTTATTTTTTTATATTTATGTAGCTTTTT |
| **11509** | BU-His3_fw | ATATAGTCGCCCATGCAGCGCAAGAAGGATCATGTATGCGCAAGCCTGGGTCACATGATTTGCGGCATCAGAGCAGATTG |
| **11510** | Chunk_4A_rv | CTAGGCTCTGCTGCATGTCAGTGATTTCTATTAGGCAGCGCTTACCCATGATTAGCGCAGCTACGGTCAGTTCGCCTTTC |
| **11511** | Chunk_4B_fw | CTGCGCTAATCATGGGTAAGCGCTGCCTAATAGAAATCACTGACATGCAGCAGAGCCTAGGTGTCAGCTTTCGTGGTGTG |
| **11512** | Chunk_4B_rv | AGTCACGCTGAGTCCATGCTGACCATGATTCACACTCAGTGCCGATAATTCCATAGTCTGCTCTCCGGCATTGACGGAAC |
| **11513** | Chunk_4C_fw | CAGACTATGGAATTATCGGCACTGAGTGTGAATCATGGTCAGCATGGACTCAGCGTGACTGCCAATTTCCGTGTTGTAGG |

| | | |
|---|---|---|
| **11514** | Chunk_4C_rv | TCAATCATTCGTTCTCGCAGATCTACAATCGTCCTGAGCTCTGTGAGTGATGTACGCTCCCTAACGCGTCAATGCACTCC |
| **11515** | Chunk_4D_fw | GGAGCGTACATCACTCACAGAGCTCAGGACGATTGTAGATCTGCGAGAACGAATGATTGATACCGTCACGCCACGTCCAC |
| **11516** | Chunk_9.2A_rv | GCGCGACGTGTCTCGTATATTAGTGAAGTTGGATCTGTCCATGAATCCTCGGCTCTGGTGTGCAAGCGGTATGAGGAAAG |
| **11517** | Chunk_9.2B_fw | CACCAGAGCCGAGGATTCATGGACAGATCCAACTTCACTAATATACGAGACACGTCGCGCTATACTGCGGGTAGGAAAGG |
| **11518** | Chunk_9.2B_rv | GTTCAGGATTCTGTCGATGCCACATCGAGTCAGTCGTAGTAACATGGAACGCAGTGCATCTCACGTTCTGGTATTGGGTGC |
| **11519** | Chunk_9.2C_fw | GATGCACTGCGTTCCATGTTACTACGACTGACTCGATGTGGCATCGACAGAATCCTGAACGCGTCGCTTTACGCCAGGTC |
| **11520** | Chunk_9.2C_rv | CAGATACTGGGCAGGCTCTATAGGAGCTTGTACCGCATTGGCTTTGCCACTCATTCGAGAGCTGCCGCCGATGAGATCGC |
| **11521** | Chunk_9.2D_fw | TCTCGAATGAGTGGCAAAGCCAATGCGGTACAAGCTCCTATAGAGCCTGCCCAGTATCTGGACTATCTGCTGACTGAGTTGCTGTTG |
| **11522** | Chunk_9.2D_rv | ATAAGGAATGATGCACGCGCGACGCTGCTTCAATCAGATGTATAGCATTGCCTTCTGCGCGGTTGCATACTGTGGCAACTGAC |
| **11523** | Chunk_5A_rv | GAGGCTTCACAGTGCTTTATTAGTATGATTGCCTAGCTGGTATATGTGTTCCTGGAGCGCTGTGGATCTGGCGGTTACGG |
| **11524** | Chunk_5B_fw | GCGCTCCAGGAACACATATACCAGCTAGGCAATCATACTAATAAAGCACTGTGAAGCCTCGCTGATTACCGCAGCCTGAA |
| **11525** | Chunk_5B_rv | AGGATCGCTCGCGTACTCATGCATTCTCCCACATATTGAGGCCCTGATTCCATGCAATGTGGAAAATCTCCGCCATTCCC |
| **11526** | Chunk_5C_fw | ACATTGCATGGAATCAGGGCCTCAATATGTGGGAGAATGCATGAGTACGCGAGCGATCCTATGGCTTACGGCAGCATTGG |
| **11527** | Chunk_5C_rv | TCTGTCAGTTGGTTAAGCGCCGCTACGATTACTACACATGCCACAGACTGATCTACAATGGTACCGCTCTGCACCACAGG |
| **11528** | Chunk_5D_fw | CATTGTAGATCAGTCTGTGGCATGTGTAGTAATCGTAGCGGCGCTTAACCAACTGACAGAACAAAGTACCGCCAGCCAGG |
| **11535** | Chunk_7A_rv | GGCGCACATGGTATATTATGATCGGAGATGCGGCAACATAGCTGGGTGTGATCCTCTCTACGCCATCCGTGGGTCTTTTC |
| **11536** | Chunk_7B_fw | TAGAGAGGATCACACCCAGCTATGTTGCCGCATCTCCGATCATAATATACCATGTGCGCCTGGGCGTCATTGTCCGGAGT |
| **11537** | Chunk_7B_rv | GAGCATACTGTCCTATCATGTCGACTCTTGTCACATCTGACGCCTCTCTGCGATAGGATTTCCGGCGGCAGCCATCAAAG |
| **11538** | Chunk_7C_fw | AATCCTATCGCAGAGAGGCGTCAGATGTGACAAGAGTCGACATGATAGGACAGTATGCTCCCAGGCGGAAGAAGTCTTTGAAGAC |
| **11539** | Chunk_7C_rv | CAGCAAGTGCGTAGAGATCAGCATTATCTGACTGTGGATGATCCTACATCGTCATCAGAGCGCCGCTTCATAAGCGCCAA |
| **11540** | Chunk_7D_fw | CTCTGATGACGATGTAGGATCATCCACAGTCAGATAATGCTGATCTCTACGCACTTGCTGAATGGCGATCCCCGAGCAAC |
| **11541** | Chunk_8A_rv | ACAATGAGAATCGAGCGCCGCTGCTTAATCTGTCAGTCGATCCTATGGTTGCTGCTGAGCACAGTGCAGCGCGTTTGGTC |
| **11542** | Chunk_8B_fw | GCTCAGCAGCAACCATAGGATCGACTGACAGATTAAGCAGCGGCGCTCGATTCTCATTGTCTGGCGGGTTACTGGCTGTG |
| **11543** | Chunk_8B_rv | GGCCGCTGTGTAGTCTCTATGCATGTACTTAGATCCTAGCGCATCTTCGCCAGCTATATTTGCCGACTTACGCCGTGGTT |

| | | |
|---|---|---|
| **11544** | Chunk_8C_fw | AATATAGCTGGCGAAGATGCGCTAGGATCTAAGTACATGCATAGAGACTACACAGCGGCCTGATCGGACTGGGCGATCAC |
| **11545** | Chunk_8C_rv | GGCATTCGGCGTGATTCCATCATGCTATGCACTGATCTCGCACATAATCTCGGTCGGCTGCGGTATGACCCTGGCGGAAG |
| **11546** | Chunk_8D_fw | CAGCCGACCGAGATTATGTGCGAGATCAGTGCATAGCATGATGGAATCACGCCGAATGCCTTTCAGCGTGCTCTGTTTACCC |
| **13395** | Telomerator_l_fw | AGGGTAATCACCCACCACAC |
| **13396** | Telomerator_l_rv | TGACGCGCTTTCAACCATGTTGCTGGGATTTCAGATTAGCCACGTACCTGGCATCAATGACCAAAGCTGGAGCTCCACCG |
| **13397** | ARS1-AD_fw | CCGATAGCAGAATGTCACACATCATGCAGATCGGCGAATGCTACATGCAGTACAGTGGAGGGCCTTTTGAAAAGCAAGCATAAAAGATCTAAAC |
| **13398** | Chunk_18A_fw | TAAGATGTGTGAACTGCGTCATACTCGCGTCAGGGATGCTTTCAGGCGTGCATGATGTAT |
| **13399** | Chunk_18A_rv | GAGATGACTGGGTCCACTCTTTCGTGTATTTCGAGAGAGCGATACGCATGTCTCCATCGTGCTAACTGTCACCCAACATAC |
| **13400** | Chunk_18B_fw | ACGATGGAGACATGCGTATCGCTCTCTCGAAATACACGAAAGAGTGGACCCAGTCATCTCGATCCGCAAGTTCTTCATCG |
| **13401** | Chunk_18B_rv | CAGATCAGTGTCAATGAAGGTAGGCTGCTTGGCAATGCTTCTGGTGACTGGTAGATCATCGCGATGTGCAATGTTCTTTGTTAC |
| **13406** | Chunk_15A_fw | TCCTCGACGCGATGGCATATCCAGTGTGATAACGTATGAGAAGGTACTGGAAGCTACTGCTGCGAGCTGAATGCCATGAC |
| **13407** | Chunk_15A_rv | GATGAACGTGCCTTCGATTTATAGAAACTGCGCTGCCCTGTGATGAATTGTCTTAGCGCGAAAGCGGCAGGTTGAGGTCC |
| **13408** | Chunk_15B_fw | CGCGCTAAGACAATTCATCACAGGGCAGCGCAGTTTCTATAAATCGAAGGCACGTTCATCCTGCGACCACGCAGTTTGAG |
| **13409** | Chunk_15B_rv | CGTGCCGGTTAATGAGCTATGCGTGTCATGTATCCTTAGGCATATCCTTAACACGCAGTGCGCCTTTGGCATGATCGAACAG |
| **13410** | Chunk_15C_fw | CACTGCGTGTTAAGGATATGCCTAAGGATACATGACACGCATAGCTCATTAACCGGCACGAACCGGCAGGTTATAGCTGATG |
| **13411** | Chunk_15C_rv | CGGGTCATTAGAGATAGTCTCTCAGGATTCAACTAGATGGTGATCTATTGTCTACGCGGCATGGCCCATATACACTTCGAGCAC |
| **13412** | Chunk_15D_fw | GCCGCGTAGACAATAGATCACCATCTAGTTGAATCCTGAGAGACTATCTCTAATGACCCGATGCGTGAATGGCTGGCAGAG |
| **13413** | Chunk_15D_rv | CGCTGACCTGTCTAACGTATCAACAGAATGCACGTCAGTCGTATGCTTGACGTGTCTGCCCAGTATCAACCACCGGGTAAC |
| **13414** | CEN6_ARS4_DJ_fw | GGCAGACACGTCAAGCATACGACTGACGTGCATTCTGTTGATACGTTAGACAGGTCAGCGGGTCCTTTTCATCACGTGCTATAAAAATAATTATAATTTAAATTTTTAATATAAATATA |
| **13415** | Chunk_16A_fw | ATGTTCGGAAGAGCTGTCTCTATCTGGTACGAGTGCTGCTGTCATCATCCTCATGCGTTGTGATGCGCGATGCTTATCAGG |
| **13416** | Chunk_16A_rv | CGCCGCTCTTAGAAGGCTATACGAGCTATGAGAGAGACTCGCTATCCATTCCGCTGAGTTCGTCAGCGATGAGACGTTAC |
| **13417** | Chunk_16B_fw | AACTCAGCGGAATGGATAGCGAGTCTCTCTCATAGCTCGTATAGCCTTCTAAGAGCGGCGGCATCGGTGAACAGGGTGCTAAG |
| **13418** | Chunk_16B_rv | CGCAAATGTCCCATCGTATTTCAGAACCTTGTCACTCATGCGAGCAAGTGTGACAGCTATATGGCGTTCTCCGCCCAGTATG |

| | | |
|---|---|---|
| **13419** | Chunk_16C_fw | ATAGCTGTCACACTTGCTCGCATGAGTGACAAGGTTCTGAAATACGATGGGACATTTGCGCCGGCGCAGATCACTTTCATAG |
| **13420** | Chunk_16C_rv | CGACAAAGTGCTCGTCACGTGCGTCAGCTTCGAGGCATATCAAGCACCTGCCGGATGATTATGCCCGTGAATGGCAAAGCG |
| **13421** | Chunk_16D_fw | AATCATCCGGCAGGTGCTTGATATGCCTCGAAGCTGACGCACGTGACGAGCACTTTGTCGAACACCGGACGGCCTTTGCTAC |
| **13422** | Chunk_16D_rv | CCTCCGCTGCGTAGAGTAATCCTGGCTCTCGCGTGTATATTGATAGATTGTCTGTCAGGCTCGCGCAGGTATGGTTCAGG |
| **13423** | Chunk 17A_fw | GCCTGACAGACAATCTATCAATATACACGCGAGAGCCAGGATTACTCTACGCAGCGGAGGTACGCAGTTTATCGGCCAGTTG |
| **13424** | Chunk 17A_rv | CAACCACCTGACTAGAGTGTCAAAGCGTGCTCCTACATAGGTAGAGTTGCATAATCTGGCAAATCGCTGAAGCGTTCC |
| **13425** | Chunk 17B_fw | GCCAGATTATGCAACTCTACCTATGTAGGAGCACGCTTTGACACTCTAGTCAGGTGGTTGTTGATAATCGCGGATGGACG |
| **13426** | Chunk 17B_rv | ATTCAGCGGGTGATCCGACTTGACTACATTTAGGTGTGGCCTCCTTACTACTCTGAGATGCATCCGGTGAAAGCGTACCC |
| **13427** | Chunk 19A_fw | CATCTCAGAGTAGTAAGGAGGCCACACCTAAATGTAGTCAAGTCGGATCACCCGCTGAATGCGATGGTCATTATTTACGGTAG |
| **13428** | Chunk 19A_rv | ATGCCGGTGGCCGAATCTATGGTCCACATTATTTGCTGCACAAGATAGTGCAGTAGCGTTCCATTATTGGCAGGATACTTTGAG |
| **13429** | Chunk 17D_fw | AACGCTACTGCACTATCTTGTGCAGCAAATAATGTGGACCATAGATTCGGCCACCGGCATTGGGTGTTTATGCCCGGACTAGC |
| **13430** | Chunk 17D_rv | ATCGACGGTCCTCGCAAGATCTCAATGTGCAGTGGTATGCTGATAACTTGTGCCTGTGGCGGGATTTAGATCCCACATTAACG |
| **13431** | ARS417_DR_fw | GCCACAGGCACAAGTTATCAGCATACCACTGCACATTGAGATCTTGCGAGGACCGTCGATACAAGTCCTTAAGAATATACAAAAAGCTACATAAATATAA |
| **13432** | Turquoise_AL_rv | CTGACTCATTGCCATTGCATGTGCAGACTCAACACCATGACATCTTAATACAGAGGACGCCGTCTCATTGGCAGCATAAA |
| **13433** | Turquoise_DS_fw | ATCAAGACTGAGGAGTACGTCAGGTTGCAGAGGATCACTTGTAATGAATGTGTGCTCGCTGAACGTTGATAGGTCAAGATCAATG |
| **13434** | Telomerator_r_fw | AGCGAGCACACATTCATTACAAGTGATCCTCTGCAACCTGACGTACTCCTCAGTCTTGATCCGGGGGATCCGGTGATTG |
| **13435** | Telomerator_r_rv | GTTATCCCTACCCACACAC |
| **13436** | rm_Telomerator_r_rv | GTTATCCCTACCCACACACCCACACACCCCAACACACCCACACACCACACACTCGAGCAATTGGGACCGTGCAATTC |
| **13509** | Chunk 19C_fw | GATGATCTACCAGTCACCAGAAGCATTGCCAAGCAGCCTACCTTCATTGACACTGATCTGGGTGCTGCAGTTGACCAGAC |
| **13510** | Chunk 19C_rv | AGCTGGCGTCGCGCATAAATGCATGATCTGTCCTGGCTGACACGCATCTGCACTAATGATGCGGCAAGAGAATTGGTTAG |
| **13511** | Chunk 19D_fw | ATCATTAGTGCAGATGCGTGTCAGCCAGGACAGATCATGCATTTATGCGCGACGCCAGCTCTGTCGTCCATGCCGGATAC |
| **13512** | Chunk 19D_rv | GCAGTAGCTTCCAGTACCTTCTCATACGTTATCACACTGGATATGCCATCGCGTCGAGGAAATCACGCGGAAATAGCTGG |

### Supplementary Table 16 - Primers for of *amdSYM* deletion

| Primer number | Primer name | Sequence (5' to 3') |
|---|---|---|
| **gRNA construction** | | |
| 6005 | p426 CRISP rv | GATCATTTATCTTTCACTGCGGAGAAG |
| 11588 | targetAmdS FW | TGCGCATGTTTCGGCGTTCGAAACTTCTCCGCAGTGAAAGATAAATGATCATCACATCCGAACATAAACAGTTTTAGAGCTAGAAATAGCAAGTTAAAATAAGGCTAGTCCGTTATCAAC |
| 11589 | targetAmdS RV | GTTGATAACGGACTAGCCTTATTTTAACTTGCTATTTCTAGCTCTAAAACTGTTTATGTTCGGATGTGATGATCATTTATCTTTCACTGCGGAGAAGTTTCGAACGCCGAAACATGCGCA |
| **Primers to make repair fragment** | | |
| 11590 | Repair AmdS FW | AAGATAGTCGCCGAACTCGCAAGAGTCATTAACACCTCGCAATTGATGGGAAGTCCTCGCATATGACCTGAACCGACGGCAAATGCTCTTCAACTACGGCATACTTGCGGAAGCTACGGC |
| 11591 | Repair AmdS RV | GCCGTAGCTTCCGCAAGTATGCCGTAGTTGAAGAGCATTTGCCGTCGGTTCAGGTCATATGCGAGGACTTCCCATCAATTGCGAGGTGTTAATGACTCTTGCGAGTTCGGCGACTATCTT |
| **Diagnostic primers to check gene deletion** | | |
| 2433 | K glycolysis Fw | GACGCCATTTGGAACGAAAAAAAG |
| 3366 | L glycolysis Rv | AATGAGTGGTAATTAATGGTGACATGAC |

### Supplementary Table 17 - Primers for deletion of *GND2, NQM1, SOL4* and *TKL2*

| Primer number | Primer name | Sequence (5' to 3') |
|---|---|---|
| **Primers to make repair fragments** | | |
| **7299** | Gnd2_repair_FW_new | AAGAATTCGTAGGTGCAGGTGAGCATATTGCCGGATAAGTGTAGTTACGCAACTACAATTGTTACTAAGGCCCAATCCGGTTGGAGAAGAACTATTGCCCTTGCTGCTACTTACGGTATT |
| **7300** | Gnd2_repair_RV_new | AATACCGTAAGTAGCAGCAAGGGCAATAGTTCTTCTCCAACCGGATTGGGCCTTAGTAACAATTGTAGTTGCGTAACTACACTTATCCGGCAATATGCTCACCTGCACCTACGAATTCTT |
| **9504** | SOL4_repair oligo fw | CAGCAGTTTTCCAAACAAAGAATGCCATTCATCAAATAATCCACAACCACCTCAAGAAAATTACACTCGTCTTTATACGAAACTGGCTCCGTTAATCACGACAGACAACCTTAATTACAT |
| **9505** | SOL4_repair oligo rv | ATGTAATTAAGGTTGTCTGTCGTGATTAACGGAGCCAGTTTCGTATAAAGACGAGTGTAATTTTCTTGAGGTGGTTGTGGATTATTTGATGAATGGCATTCTTTGTTTGGAAAACTGCTG |
| **9509** | TKL2_repair oligo fw | TTGTTGGGAGGAGTCCTGAATAAGGAGTGTCGAATATAGGGAGCTTCATTCGTTGTCAAGGAAGTAAACAGTTCTTTGCTATTTCACACTTCCTGGTTGATGGTCACTTGCTGCCTGAAA |
| **9510** | TKL2_repair oligo rv | TTTCAGGCAGCAAGTGACCATCAACCAGGAAGTGTGAAATAGCAAAGAACTGTTTACTTCCTTGACAACGAATGAAGCTCCCTATATTCGACACTCCTTATTCAGGACTCCTCCCAACAA |

| | | |
|---|---|---|
| **12570** | NQM1_repair oligo fw | TTCTTGCTAGCGTAAGTCATAAAAAATAGGAAATAATCACATATATAC AAGAAATTAAATTCATTAAGAGTAGAGGTACCTACTTATATATATAAA TATATATATACCACTTTCCTTTTC |
| **12571** | NQM1_repair oligo rv | GAAAAGGAAAGTGGTATATATATATTTATATATATAAGTAGGTACCTC TACTCTTAATGAATTTAATTTCTTGTATATATGTGATTATTTCCTATTTT TTATGACTTACGCTAGCAAGAA |
| **Diagnostic primers to check gene deletion** | | |
| **7258** | FW_gnd2KO_check | TCTGACAGGTGGCAGTTTCC |
| **7259** | RV_gnd2KO_check | ATCCGAAAGGCGGCAATAGG |
| **9506** | SOL4_dg fw | GGGTGGACGTTTAAGCATAC |
| **9507** | SOL4_dg rv | GTATCACCGGGTGAGCTATG |
| **1360** | TKL2 dis500 fw | TCTTAATGGTGGCTCGCTGTC |
| **1361** | TKL2 dis500 rv | TCAATGCAGCCCATACACTC |
| **12572** | NQM1_dg fw | CCTTGATCTGGCTCTGGCTC |
| **12573** | NQM1_dg rv | CGCAAGGTAATTACGCCACG |

### Supplementary Table 18 - Primers for deletion of *ura3, his3* and *SpHIS5*

| Primer number | Primer name | Sequence (5' to 3') |
|---|---|---|
| **Primers to make repair fragments** | | |
| **10521** | HIS3_repair oligo fw | AATGTGATTTCTTCGAAGAATATACTAAAAAATGAGCAGGCAAGATA AACGAAGGCAAAGTGACACCGATTATTTAAAGCTGCAGCATACGATA TATATACATGTGTATATATGTATACC |
| **10522** | HIS3_repair oligo rv | GGTATACATATATACACATGTATATATATCGTATGCTGCAGCTTTAAAT AATCGGTGTCACTTTGCCTTCGTTTATCTTGCCTGCTCATTTTTTAGTAT ATTCTTCGAAGAAATCACATT |
| **12685** | spHIS5 Repair oligo FW | GCCCAACTCAGCTTCCGTAAACCACAACACCACCACTAATACAACTCT ATCATACACAAGTCTTTTTACATTTTTTTGGTTTGTGTACGTATCCCACC GTACTTACCATCTTCTCTCCTT |
| **12686** | spHIS5 Repair oligo RV | AAGGAGAGAAGATGGTAAGTACGGTGGGATACGTACACAAACCAAA AAAATGTAAAAAGACTTGTGTATGATAGAGTTGTATTAGTGGTGGTG TTGTGGTTTACGGAAGCTGAGTTGGGC |
| **13807** | URA3repair_FW | CGGTTTCCTTGAAATTTTTTTGATTCGGTAATCTCCGAACAGAAGGAA GAACGAAGGAAGGGAATCTCGGTCGTAATGATTTCTATAATGACGAA AAAAAAAAAATTGGAAAGAAAAAGC |
| **13808** | URA3repair_RV | GCTTTTTCTTTCCAATTTTTTTTTTTTCGTCATTATAGAAATCATTACGA CCGAGATTCCCTTCCTTCGTTCTTCCTTCTGTTCGGAGATTACCGAATC AAAAAAATTTCAAGGAAACCG |
| **Diagnostic primers to check gene deletion** | | |
| **111** | URA3 CTRL RV | TATACGCCAGTACACCTTATCGGC |
| **1026** | HIS3 outside fw | CGCTTTGTCTTCATTCAACGTTTCC |
| **1024** | HIS3 outside rv | CCACTTGCCACCTATCAC |
| **1460** | GLK1FW1 | CGCGCCCATATAAATATCC |
| **1461** | GLK1RV1 | CCCGTTTCCGATGATATTG |
| **1521** | URA3 Ctrl Fw2 | GCTACTGCGCCAATTGATGAC |

### Supplementary Table 19 - Primers for deletion of *ARO10*

| Primer number | Primer name | Sequence (5' to 3') |
|---|---|---|
| **Primers to make repair fragments** | | |
| **7247** | ARO10 CRISPR repair upper | ACAAGTTGACGCGACTTCTGTAAAGTTTATTTACAAGATAACAAAGAAACTCCCTTAAGCAAACTTGTGGGCGCAATTATAAAACACTGCTACCAATTGTTCGTTTTCTGTTCATTAACA |
| **7248** | ARO10 CRISPR repair lower | TGTTAATGAACAGAAAACGAACAATTGGTAGCAGTGTTTTATAATTGCGCCCACAAGTTTGCTTAAGGGAGTTTCTTTGTTATCTTGTAAATAAACTTTACAGAAGTCGCGTCAACTTGT |
| **Diagnostic primers to check gene deletion** | | |
| **2359** | Aro10 KO CHK for | TGCTTGTACACCTCATGTAG |
| **2360** | Aro 10 KO chk rev | GCAGACATTTAGCAGATGTAG |

### Supplementary Table 20 - List of NeoChr25 (linear) and NeoChr26 (circular) chromosome parts

#The +, - or 0 signifies the orientation of the part with respect to the neochromosome
* Size of the fragments does not include the SHR sequences.

| SHR Fw | Component# | SHR Rv | Size* | Template | Primer Fw | Primer Rv |
|---|---|---|---|---|---|---|
| **Unique to NeoChr25** | | | | | | |
| | Left TeSS (0) | BJ | 813 | pLM092 | 13395 | 16577 |
| **Common to Neochr25 and NeoChr26** | | | | | | |
| **BJ** | chunk 7BC (0) | BL | 4909 | *E.coli* (migula) Castellani and Chalmers (ATCC 47076) | 11536 | 11539 |
| **BL** | ARS1 (0) | AN | 56 | Annealing of complementary primers | 16578 | 9989 |
| **AN** | ZWF1 (+) | BP | 2623 | pUDC275 | 16526 | 16527 |
| **BP** | TKL1 (+) | DE | 3148 | pUDC276 | 16528 | 16529 |
| **DE** | GND1 (+) | BQ | 2575 | pUDC277 | 16530 | 16531 |
| **BQ** | RKI1 (-) | BR | 1882 | pUDC278 | 16532 | 16533 |
| **BR** | TAL1 (-) | AL | 2113 | pUDC279 | 16534 | 16535 |
| **AL** | mTurqouise2 (-) | DS | 1683 | pUDC192 | 13432 | 13433 |
| **DS** | RPE1 (-) | DF | 1822 | pUDC280 | 16536 | 16537 |
| **DF** | SOL3 (-) | DI | 1855 | pUDC281 | 16538 | 16539 |
| **DI** | ARS417 (0) | BE | 64 | Annealing of complementary primers | 16579 | 16580 |
| **BE** | aroG$^{fbr}$ (+) | DK | 2089 | pUDC298 | 16540 | 16541 |
| **DK** | aroB (+) | AC | 2104 | pUDC294 | 16542 | 16543 |
| **AC** | mRuby2 (+) | AD | 1667 | pUDC191 | 11365 | 11366 |
| **AD** | aroD (+) | DM | 1931 | pUDC296 | 16544 | 16545 |
| **DM** | aroE (+) | DN | 1860 | pUDC297 | 16546 | 16547 |
| **DN** | aroL (+) | DO | 1534 | pUDC299 | 16548 | 16549 |
| **DO** | aroA (+) | DP | 2718 | pUDC293 | 16550 | 16551 |

| | | | | | | |
|---|---|---|---|---|---|---|
| **DP** | aroC (-) | DQ | 1746 | pUDC295 | 16552 | 16553 |
| **DQ** | tyrA$^{fbr}$ (-) | DR | 1874 | pUDC301 | 16554 | 16555 |
| **DR** | pheA$^{fbr}$ (-) | AJ | 2171 | pUDC300 | 16556 | 16557 |
| **AJ** | tyrB (-) | DH | 2499 | pUDC302 | 16558 | 16559 |
| **DH** | chunk 15CD (0) | DJ | 4996 | *E.coli* (migula) Castellani and Chalmers (ATCC 47076) | 13410 | 13413 |
| **DJ** | CEN6/ARS4 (0) | AE | 519 | | 13414 | 9991 |
| **AE** | chunk 16AB (0) | DL | 5019 | *E.coli* (migula) Castellani and Chalmers (ATCC 47076) | 13415 | 16560 |
| **DL** | FBA1 (+) | H | 2185 | pUDC212 | 16561 | 16562 |
| **H** | TPI1 (+) | P | 1856 | pUDC222 | 12957 | 12958 |
| **P** | PGK1 (+) | Q | 2356 | pUDC219 | 16563 | 16564 |
| **Q** | ADH1 (+) | N | 2152 | pUDC229 | 16565 | 16566 |
| **N** | PYK1 (+) | O | 2612 | pUDC220 | 12963 | 12964 |
| **O** | TDH3 (+) | A | 2104 | pUDC230 | 16567 | 16568 |
| **A** | ENO2 (+) | B | 2423 | pUDC231 | 12967 | 13227 |
| **B** | HXK2 (-) | C | 2566 | pUDC214 | 16569 | 16570 |
| **C** | PGI1 (-) | D | 2774 | pUDC232 | 12972 | 12971 |
| **D** | PFK1 (-) | J | 4069 | pUDC216 | 16571 | 16572 |
| **J** | PFK2 (-) | BU | 3989 | pUDC217 | 12976 | 13900 |
| **BU** | HIS3 (-) | L | 1254 | pLM092 | 16573 | 16574 |
| **L** | GPM1 (-) | M | 1853 | pUDC213 | 12980 | 12979 |
| **M** | PDC1 (-) | AR | 2797 | pUDC215 | 16575 | 16576 |
| **AR** | ARS1211 (0) | BS | 251 | *S. cerevisiae* CEN.PK113-7D | 12983 | 13901 |
| **BS** | chunk 9CD (0) | AQ | 4994 | *E.coli* (migula) Castellani and Chalmers (ATCC 47076) | 11519 | 11522 |
| **Unique to NeoChr25** | | | | | | |
| AQ | right TeSS (0) | | 807 | pLM092 | 16581 | 13435 |
| **Unique to NeoChr26** | | | | | | |
| AQ | telomerator (+) | BJ | 1620 | pLM092 | 16581 | 16577 |

### Supplementary Table 21 - List of primers for amplifying NeoChr25 and NeoChr26 chromosome parts

| Primer Number | Primer Name | Sequence (5' to 3') |
|---|---|---|
| **13395** | Telomerator_l_fw | AGGGTAATCACCCACCACAC |
| **16577** | BJ_tel_le _RV | GGCGCACATGGTATATTATGATCGGAGATGCGGCAACATAGCTGGGTGTGATCCTCTCTACCAAAGCTGGAGCTCCACCG |
| **11536** | Chunk_7B_fw | TAGAGAGGATCACACCCAGCTATGTTGCCGCATCTCCGATCATAATATACCATGTGCGCCTGGGCGTCATTGTCCGGAGT |
| **11539** | Chunk_7C_rv | CAGCAAGTGCGTAGAGATCAGCATTATCTGACTGTGGATGATCCTACATCGTCATCAGAGCGCCGCTTCATAAGCGCCAA |
| **16578** | BL_ARS1_FW | CTCTGATGACGATGTAGGATCATCCACAGTCAGATAATGCTGATCTCTACGCACTTGCTGGGCCTTTTGAAAAGCAAGCATAAAAGATCTAAAC |
| **9989** | ARS1_rv | ATACATCATGCACGCCTGAAAGCATCCCTGACGCGAGTATGACGCAGTTCACACATCTTACTTGTTATTTTACAGATTTTATGTTTAGATCTTTTATGCTTGCTTTTCAAAA |
| **16526** | AN_pZWF1_FW | TAAGATGTGTGAACTGCGTCATACTCGCGTCAGGGATGCTTTCAGGCGTGCATGATGTATGTCCGGGTGCATGCATGAA |
| **16527** | BP_tZWF1_RV | GAGATGACTGGGTCCACTCTTTCGTGTATTTCGAGAGAGCGATACGCATGTCTCCATCGTATATATTCAATTCATATTTTATCTCTTTTT |
| **16528** | BP_pTKL1_FW | ACGATGGAGACATGCGTATCGCTCTCTCGAAATACACGAAAGAGTGGACCCAGTCATCTCTAAGCGCTTTTTTTTTTTTT |
| **16529** | DE_tTKL1_RV | GCAGTAGCTTCCAGTACCTTCTCATACGTTATCACACTGGATATGCCATCGCGTCGAGGAATATTCTTTATTGGCTTTATACTTG |
| **16530** | DE_pGND1_FW | TCCTCGACGCGATGGCATATCCAGTGTGATAACGTATGAGAAGGTACTGGAAGCTACTGCTCACTTCCGCCGCAAACAATAC |
| **16531** | BQ_tGND1_RV | CAGATCAGTGTCAATGAAGGTAGGCTGCTTGGCAATGCTTCTGGTGACTGGTAGATCATCCTACTCTACTTCTATCATGATAATAGGCAC |
| **16532** | BQ_tRKI1_FW | GATGATCTACCAGTCACCAGAAGCATTGCCAAGCAGCCTACCTTCATTGACACTGATCTGCTTGGTGTGTCATCGGTAGTAACG |
| **16533** | BR_pRKI1_RV | AGCTGGCGTCGCGCATAAATGCATGATCTGTCCTGGCTGACACGCATCTGCACTAATGATTTGCGAACGCATTAAGTTGAG |
| **16534** | BR_tTAL1_FW | ATCATTAGTGCAGATGCGTGTCAGCCAGGACAGATCATGCATTTATGCGCGACGCCAGCTGACGTTGATTTAAGGTGGTTC |
| **16535** | AL_pTAL1_RV | GCGTCCTCTGTATTAAGATGTCATGGTGTTGAGTCTGCACATGCAATGGCAATGAGTCAGGAAAAGCTAGAAAAGGAATTAGAC |
| **13433** | Turquoise_DS_fw | ATCAAGACTGAGGAGTACGTCAGGTTGCAGAGGATCACTTGTAATGAATGTGTGCTCGCTGAACGTTGATAGGTCAAGATCAATG |
| **13432** | Turquoise_AL_rv | CTGACTCATTGCCATTGCATGTGCAGACTCAACACCATGACATCTTAATACAGAGGACGCCGTCTCATTGGCAGCATAAA |
| **16536** | DS_tRPE1_FW | AGCGAGCACACATTCATTACAAGTGATCCTCTGCAACCTGACGTACTCCTCAGTCTTGATAAATGGATATTGATCTAGATGGCGG |

| | | |
|---|---|---|
| **16537** | DF_pRPE1_RV | GATGAACGTGCCTTCGATTTATAGAAACTGCGCTGCCCTGTGATGAATTGT CTTAGCGCGACCACTTGACAACGGTCTTG |
| **16538** | DF_tSOL3_FW | CGCGCTAAGACAATTCATCACAGGGCAGCGCAGTTTCTATAAATCGAAGG CACGTTCATCAGGATGCACTCTACAAATAC |
| **16539** | DI_pSOL3_RV | CGGGTCATTAGAGATAGTCTCTCAGGATTCAACTAGATGGTGATCTATTGT CTACGCGGCCTGACTGCAATAGGAAACTG |
| **16579** | DI_ARS417_FW | GCCGCGTAGACAATAGATCACCATCTAGTTGAATCCTGAGAGACTATCTCT AATGACCCGACAAGTCCTTAAGAATATACAAAAAGCTACATAAATATAA |
| **16580** | BE_ARS417_RV | TCAATCATTCGTTCTCGCAGATCTACAATCGTCCTGAGCTCTGTGAGTGAT GTACGCTCCCTTACGCTCAATTCCTTTATTTTTTTATATTTATGTAGCTTTTT |
| **16540** | BE_pHHF1_FW | GGAGCGTACATCACTCACAGAGCTCAGGACGATTGTAGATCTGCGAGAAC GAATGATTGATCTTGGGGCCTTACCACC |
| **16541** | DK_tTEF1_RV | CGCCGCTCTTAGAAGGCTATACGAGCTATGAGAGAGACTCGCTATCCATTC CGCTGAGTTGGTATCACCATAGATTTCGAAAC |
| **16542** | DK_pHHF2_FW | AACTCAGCGGAATGGATAGCGAGTCTCTCTCATAGCTCGTATAGCCTTCTA AGAGCGGCGTGTGGAGTGTTTGCTTGG |
| **16543** | AC_tTEF2_RV | GCTACATCTTCCGTACTATGCTGTAGTCTCATGGTCGAGTTCTATTGCTGTT CGGCGGCAAGGAAACGTAAATTACAAGG |
| **11365** | prCCW12_mRuby _tENO1_fw | TGCCGCCGAACAGCAATAGAACTCGACCATGAGACTACAGCATAGTACGG AAGATGTAGCAACGCACCCATGAACCACAC |
| **11366** | prCCW12_mRuby _tENO1_rv | CTCCACTGTACTGCATGTAGCATTCGCCGATCTGCATGATGTGTGACATTC TGCTATCGGGGCAGCATACATGGGTGACCAAA |
| **16544** | AD_pRPL25_FW | CCGATAGCAGAATGTCACACATCATGCAGATCGGCGAATGCTACATGCAG TACAGTGGAGAGGTATGTTAGTGCTAAAAG |
| **16545** | DM_tGPH1_RV | CGACAAAGTGCTCGTCACGTGCGTCAGCTTCGAGGCATATCAAGCACCTG CCGGATGATTAAACGTCAGTACATCCTACC |
| **16546** | DM_pRPP0_FW | AATCATCCGGCAGGTGCTTGATATGCCTCGAAGCTGACGCACGTGACGAG CACTTTGTCGTTCAACAATTCGTTATATATATGGTAGGCT |
| **16547** | DN_tCYC1_RV | CCTCCGCTGCGTAGAGTAATCCTGGCTCTCGCGTGTATATTGATAGATTGT CTGTCAGGCAAGCTTGTCCCAAAACCTTCTC |
| **16548** | DN_pHTB2_FW | GCCTGACAGACAATCTATCAATATACACGCGAGAGCCAGGATTACTCTAC GCAGCGGAGGTATATATTAAATTTGCTCTTGTTCTG |
| **16549** | DO_tPGM2_RV | CAACCACCTGACTAGAGTGTCAAAGCGTGCTCCTACATAGGTAGAGTTGC ATAATCTGGCAACTCGGGGTAGGTAATC |
| **16550** | DO_pRPL3_FW | GCCAGATTATGCAACTCTACCTATGTAGGAGCACGCTTTGACACTCTAGTC AGGTGGTTGAGAGTCTTGGAGATTTTCGACCTG |
| **16551** | DP_tSOL4_RV | ATTCAGCGGGTGATCCGACTTGACTACATTTAGGTGTGGCCTCCTTACTAC TCTGAGATGAGTCATAGCATTAAGATTAACGCGTTG |
| **16552** | DP_tGPD2_FW | CATCTCAGAGTAGTAAGGAGGCCACACCTAAATGTAGTCAAGTCGGATCA CCCGCTGAATTTAAGGGCTATAGATAACAG |

| | | |
|---|---|---|
| **16553** | DQ_pRPL8A_RV | ATGCCGGTGGCCGAATCTATGGTCCACATTATTTGCTGCACAAGATAGTGCAGTAGCGTTAACGACATAAATAATTTCTATTAAC |
| **16554** | DQ_tGDB1_FW | AACGCTACTGCACTATCTTGTGCAGCAAATAATGTGGACCATAGATTCGGCCACCGGCATCAAATACGTACGTGGCAACCCTTTC |
| **16555** | DR_pRPL10_RV | ATCGACGGTCCTCGCAAGATCTCAATGTGCAGTGGTATGCTGATAACTTGTGCCTGTGGCTCACTTGTCTGTGTGTTTAACTGCC |
| **16556** | DR_tGSY2_FW | GCCACAGGCACAAGTTATCAGCATACCACTGCACATTGAGATCTTGCGAGGACCGTCGATGTATGACTATATGTTGATAACTG |
| **16557** | AJ_pRPL18A_RV | ACGCAATATCGGCCATCGTGCGAGTGTCTCAAACTATCTGTATGCAAATTCGTGCGTGTGAAGAGGATGTCCAATATTTTTTT |
| **16558** | AJ_tGLC3_FW | CACACGCACGAATTTGCATACAGATAGTTTGAGACACTCGCACGATGGCCGATATTGCGTTAGGTTAAACCTTGGAAGAG |
| **16559** | DH_pCWP2_RV | CGTGCCGGTTAATGAGCTATGCGTGTCATGTATCCTTAGGCATATCCTTAACACGCAGTGCTAATAGACAAGGTGCTATGAG |
| **13410** | Chunk_15C_fw | CACTGCGTGTTAAGGATATGCCTAAGGATACATGACACGCATAGCTCATTAACCGGCACGAACCGGCAGGTTATAGCTGATG |
| **13413** | Chunk_15D_rv | CGCTGACCTGTCTAACGTATCAACAGAATGCACGTCAGTCGTATGCTTGACGTGTCTGCCCAGTATCAACCACCGGGTAAC |
| **13414** | CEN6_ARS4_DJ_fw | GGCAGACACGTCAAGCATACGACTGACGTGCATTCTGTTGATACGTTAGACAGGTCAGCGGGTCCTTTTCATCACGTGCTATAAAAATAATTATAATTTAAATTTTTTAATATAAATATA |
| **9991** | CEN6_ARS4_rv | CAACGCATGAGGATGATGACAGCAGCACTCGTACCAGATAGAGACAGCTCTTCCGAACATGGACGGATCGCTTGCCTGTAAC |
| **13415** | Chunk_16A_fw | ATGTTCGGAAGAGCTGTCTCTATCTGGTACGAGTGCTGCTGTCATCATCCTCATGCGTTGTGATGCGCGATGCTTATCAGG |
| **16560** | DL_Chunk 16B_rv | CGCAAATGTCCCATCGTATTTCAGAACCTTGTCACTCATGCGAGCAAGTGTGACAGCTATATGGCGTTCTCCGCCCGGTATG |
| **16561** | DL_pFBA1_FW | ATAGCTGTCACACTTGCTCGCATGAGTGACAAGGTTCTGAAATACGATGGGACATTTGCGTGAACAACAATACCAGCCTTCC |
| **16562** | H_tFBA1_RV | GTCACGGGTTCTCAGCAATTCGAGCTATTACCGATGATGGCTGAGGCGTTAGAGTAATCTAATGAGCTATCAAAAACGATAGATC |
| **12957** | TPI FW + H | AGATTACTCTAACGCCTCAGCCATCATCGGTAATAGCTCGAATTGCTGAGAACCCGTGACAACGAAGACCCAGAGATGTTGTTGT |
| **12958** | TPI Rv + P | CTGATAGTGCTGTAAGTCGCCTCCATCTTAGCAGAGCTGTCCCTGAATGCGTACTCGTGATGAGTAACCCATATAGAGATCGTAC |
| **16563** | P_pPGK1_FW | TCACGAGTACGCATTCAGGGACAGCTCTGCTAAGATGGAGGCGACTTACAGCACTATCAGTCTTTTTATTAACCTTAATTTTTAT |
| **16564** | Q_tPGK1_RV | GAGCTGAATGTATATGCTGCGGGATCATTGCACAGCTCTGAGAGCCCTGCAACGCGATATAAATAATATCCTTCTCGAAAGC |

| | | |
|---|---|---|
| **16565** | Q_pADH1_FW | ATATCGCGTTGCAGGGCTCTCAGAGCTGTGCAATGATCCCGCAGCATATAC ATTCAGCTCAAGTCCAATGCTAGTAGAGAAG |
| **16566** | N_tADH1_RV | TTCTAGGCTTTGATGCAAGGTCCACATATCTTCGTTAGGACTCAATCGTGG CTGCTGATCTTGTCCTCTGAGGACATAAAATAC |
| **12963** | PYK1 Fw + N | GATCAGCAGCCACGATTGAGTCCTAACGAAGATATGTGGACCTTGCATCA AAGCCTAGAAAACGTGGTCAAACTTCAGAACTAAG |
| **12964** | PYK1 Rv + O | ATACTCCCTGCACAGATGAGTCAAGCTATTGAACACCGAGAACGCGCTGA ACGATCATTCATAATCATGATAACCTTGAGGGAAG |
| **16567** | O_pTDH3_FW | GAATGATCGTTCAGCGCGTTCTCGGTGTTCAATAGCTTGACTCATCTGTGC AGGGAGTATATACTAGCGTTGAATGTTAGCGTC |
| **16568** | A_tTDH3_RV | GTGCCTATTGATGATCTGGCGGAATGTCTGCCGTGCCATAGCCATGCCTTC ACATATAGTATCCTGGCGGAAAAAATTCATTTG |
| **12967** | ENO2 Fw + A | ACTATATGTGAAGGCATGGCTATGGCACGGCAGACATTCCGCCAGATCAT CAATAGGCACAACGGATGATGAAAACACTAAACGA |
| **13227** | ENO2 Rv + B | GTTGAACATTCTTAGGCTGGTCGAATCATTTAGACACGGGCATCGTCCTCT CGAAAGGTGTAACGAAGACGTTACCAGCTGATTG |
| **16569** | B_tHXK2_FW | CACCTTTCGAGAGGACGATGCCCGTGTCTAAATGATTCGACCAGCCTAAG AATGTTCAACACTTGAACAATAAATACGAAATCC |
| **16570** | C_pHXK2_RV | CTAGCGTGTCCTCGCATAGTTCTTAGATTGTCGCTACGGCATATACGATCC GTGAGACGTACGCTGGTAAAGTACAGCTA |
| **12971** | PGI1 Fw + D | AATCACTCTCCATACAGGGTTTCATACATTTCTCCACGGGACCCACAGTCGT AGATGCGTAACGTATTCTTAGTGGATAACATGC |
| **12972** | PGI1 Rv + C | ACGTCTCACGGATCGTATATGCCGTAGCGACAATCTAAGAACTATGCGAG GACACGCTAGTTTTAAACAGTTGATGAGAACCTTT |
| **16571** | D_tPFK1_FW | ACGCATCTACGACTGTGGGTCCCGTGGAGAAATGTATGAAACCCTGTATG GAGAGTGATTATTCCATAGCTTAGTTTAATCAAGG |
| **16572** | J_pPFK1_RV | CGACGAGATGCTCAGACTATGTGTTCTACCTGCTTGGACATCTTCGCGTAT ATGACGGCCCGGCTAGTAAAAAGAAAATTAATA |
| **12976** | PFK2 Rv + J | GGCCGTCATATACGCGAAGATGTCCAAGCAGGTAGAACACATAGTCTGAG CATCTCGTCGAAATCGTCTATATCACATATTCCAG |
| **13900** | PFK2_fw + BU | AATCATGTGACCCAGGCTTGCGCATACATGATCCTTCTTGCGCTGCATGGG CGACTATATAACGATTCTCTGCTGCTTTGTTGCA |
| **16573** | BU_tHIS3_FW | ATATAGTCGCCCATGCAGCGCAAGAAGGATCATGTATGCGCAAGCCTGGG TCACATGATTGCATCTGTGCGGTATTTCACAC |
| **16574** | L_pHIS3_RV | GCCGTAGCTTCCGCAAGTATGCCGTAGTTGAAGAGCATTTGCCGTCGGTTC AGGTCATATTGCGGCATCAGAGCAGATTG |
| **12980** | GPM1 Rv + L | ATATGACCTGAACCGACGGCAAATGCTCTTCAACTACGGCATACTTGCGGA AGCTACGGCTATTGCTATAACATGTCATGTCACC |
| **12979** | GPM1 Fw + M | ACGAGAGATGAAGGCTCACCGATGGACTTAGTATGATGCCATGCTGGAAG CTCCGGTCATAACGGTGATACTTTGACAGGAGCTA |

| | | |
|---|---|---|
| **16575** | M_tPDC1_FW | ATGACCGGAGCTTCCAGCATGGCATCATACTAAGTCCATCGGTGAGCCTTC ATCTCTCGTACAGTGTTCCTTAATCAAGGATAC |
| **16576** | AR_pPDC1_RV | TGACGAGATTTGAGAAGTCCCCAATATCGACTCGTGATGTGCCATGCGTG CTGTCAGTATCATGCGACTGGGTGAGCATA |
| **12983** | ARS1211_fw +AR | ATACTGACAGCACGCATGGCACATCACGAGTCGATATTGGGGACTTCTCA AATCTCGTCAGACATAGTATTTCGCAACCTTTCAG |
| **13901** | ARS1211_rv + BS | GTTCAGGATTCTGTCGATGCCACATCGAGTCAGTCGTAGTAACATGGAAC GCAGTGCATCGACAGGCGTTTCTGTACCGCTGTTA |
| **11519** | Chunk_9.2C_fw | GATGCACTGCGTTCCATGTTACTACGACTGACTCGATGTGGCATCGACAGA ATCCTGAACGCGTCGCTTTACGCCAGGTC |
| **11522** | Chunk_9.2D_rv | ATAAGGAATGATGCACGCGCGACGCTGCTTCAATCAGATGTATAGCATTG CCTTCTGCGCGGTTGCATACTGTGGCAACTGAC |
| **16581** | AQ_tel_right_FW | GCGCAGAAGGCAATGCTATACATCTGATTGAAGCAGCGTCGCGCGTGCAT CATTCCTTATCCGGGGGATCCGGTGATTG |
| **13435** | Telomerator_r_rv | GTTATCCCTACCCACACAC |

### Supplementary Table 225 - Primers for glycolysis deletion

| Primer number | Primer name | Sequence (5' to 3') |
|---|---|---|
| **Primers for repair fragment IMF27 transformation** | | |
| 13273 | URA3 repair SGA1 Fw | TTTTCTCATCTCTTGGCTCTGGATCCGTTATCTGTTCTGTTACACAA GAAATCGTACATAACTGTCATCCTGCGTGAAGATTAA |
| 13274 | URA3 repair SGA1 Rv | TCTCGCTTTTCTTTTATTTTTTTTTGTCTACAAACTCTGTAAAACTTC TTGTCTTATTTGAGTGTTGCACCGTGCCAATGCAGGT |
| **Primers to make repair fragment IMF29 transformation** | | |
| 6075 | COUNTER SELECT oligo fw | TTTTTCTCATCTCTTGGCTCTGGATCCGTTATCTGTTCTGTTACACA AGAAATCGTACATACTAGAGCAAGATTTCAAATAAGTAACAGCA GCCATACGTTGAAACTACGGCAAAGGATT |
| 6076 | COUNTER SELECT oligo rv | AATCCTTTGCCGTAGTTTCAACGTATGGCTGCTGTTACTTATTTGA AATCTTGCTCTAGTATGTACGATTTCTTGTGTAACAGAACAGATA ACGGATCCAGAGCCAAGAGATGAGAAAAA |
| **Diagnostic primers** | | |
| 3751 | sequence primer right - fw | GGTCAGCAGTACAGAACCGTCG |
| 4229 | Sequence SGA1 2 rv | TGGTCGACAGATACAATCCTGG |
| 4880 | c I-SceI inside rv | GCCAATCAAACCCTTCTTCTC |
| 7298 | FW_sga1u_check | TTGTTCAATGGATGCGGTTC |

### Supplementary Table 23 - Primers to repair *RKI1* mutation in IMF32

| Primer number | Primer name | Sequence (5' to 3') |
|---|---|---|
| **Primers to make repair fragment** | | |
| 17614 | RKI1_repair_SNP1_fw | GAGAATTTACAATTAATTAAAGGTGGTGGTGCTTGTCTATTTCAAGAA AAATTGGTTAGCACTAGCGCTAAAACATTCATTGTCGTTGCTGATTCA AGAAAAAAGTCCCCAAAACATCTA |
| 17615 | RKI1_repair_SNP1_Rv | TAGATGTTTTGGGGACTTTTTTCTTGAATCAGCAACGACAATGAATGT TTTAGCGCTAGTGCTAACCAATTTTTCTTGAAATAGACAAGCACCACC ACCTTTAATTAATTGTAAATTCTC |
| **Diagnostic primers to check gene deletion** | | |
| 17623 | RKI1_SNPT_dg_fw | GGTGGTGCTTGTCTATTTCAAT |
| 17624 | RKI1_SNPG_dg_fw | GGTGGTGCTTGTCTATTTCAAG |
| 17625 | RKI1_SNP_dg_rv | GGTTTCCACCACTCCCACTAAA |

### Supplementary Table 24 - Primers for deletion of native *ZWF1, GND1, SOL3, RKI1, TAL1, TKL1* and *RPE1* ORFs

| Primer number | Primer name | Sequence (5' to 3') |
|---|---|---|
| **Primers to make repair fragment** | | |
| 7363 | FW_Gnd1_repair | TAAACCTGTATTGTTGCCATTACAGAAAAAAGCCACTTTCTATACAAAAACTACAATAAATTCAGAGTGTTGCCAGAATGTGCTTCTGACAACTTGCCAGTAGACAAGGATATCCATATC |
| 7364 | RV_Gnd1_repair | GATATGGATATCCTTGTCTACTGGCAAGTTGTCAGAAGCACATTCTGGCAACACTCTGAATTTATTGTAGTTTTTGTATAGAAAGTGGCTTTTTTCTGTAATGGCAACAATACAGGTTTA |
| 8868 | FW_zwf1_repair | CAATTGGCTGTATAGACAGAAAGAGTAAATCCAATAGAATAGAAAACCACATAAGGCAAGAGATACGAAGGATAATTAGAAAAATGCAAGCACATTCATTTATCGGCTAAGTCACTGAAA |
| 8869 | RV_zwf1_repair | TTTCAGTGACTTAGCCGATAAATGAATGTGCTTGCATTTTTCTAATTATCCTTCGTATCTCTTGCCTTATGTGGTTTTCTATTCTATTGGATTTACTCTTTCTGTCTATACAGCCAATTG |
| 9281 | RPE1_repair oligo fw | CAATTTCATGCAAGAAGGCCATTTGCTAATTCCAAGAGCGAGGTAAACACACAAGAAAAATTGTACATATGCGGCATTTCTTATATTTATACTCTCTATACTATACGATATGGTATTTTT |
| 9282 | RPE1_repair oligo rv | AAAAATACCATATCGTATAGTATAGAGAGTATAAATATAAGAAATGCCGCATATGTACAATTTTTCTTGTGTGTTTACCTCGCTCTTGGAATTAGCAAATGGCCTTCTTGCATGAAATTG |
| 16891 | SOL3_repair oligo fw | GCCTCGAGGATAATAGAAGGCAATGCACCATCAATTGCTTTACCCCTGGTCCGCGACCAAAAAAGACACACATGCGAGCTTTCGAACCTCAGATGCTAATATTACGTGTTATATATACCA |
| 16892 | SOL3_repair oligo rv | TGGTATATATAACACGTAATATTAGCATCTGAGGTTCGAAAGCTCGCATGTGTGTCTTTTTTGGTCGCGGACCAGGGGTAAAGCAATTGATGGTGCATTGCCTTCTATTATCCTCGAGGC |
| 16899 | RKI1_repair oligo fw | TGTTACATAAACTTGGTTACCGCATACTGCAACCTCATATAAATACAACATAGGAAAGAAGCAGATCAAAGGCAAAGACAGAAACCGTAGTAAAGGTTGACTTTTCACAACAGTGTCTCC |
| 16900 | RKI1_repair oligo rv | GGAGACACTGTTGTGAAAAGTCAACCTTTACTACGGTTTCTGTCTTTGCCTTTGATCTGCTTCTTTCCTATGTTGTATTTATATGAGGTTGCAGTATGCGGTAACCAAGTTTATGTAACA |
| 16901 | TKL1_repair oligo fw | ACAACAGAGAAGGAAGCTCATCCCAAGCAACTCTACATAGTTACCTCTTTAGCAAACAAAATTCTGATCGTAGATCATCAGATTTGATATGATATTATTTGTGAAAAAATGAAATAAAAC |
| 16902 | TKL1_repair oligo rv | GTTTTATTTCATTTTTTCACAAATAATATCATATCAAATCTGATGATCTACGATCAGAATTTTGTTTGCTAAAGAGGTAACTATGTAGAGTTGCTTGGGATGAGCTTCCTTCTCTGTTGT |
| 16907 | TKL1_repair oligo fw | ACAACAGAGAAGGAAGCTCATCCCAAGCAACTCTACATAGTTACCTCTTTAGCAAACAAAATTCTGATCGTAGATCATCAGATTTGATATGATATTATTTGTGAAAAAATGAAATAAAAC |

| | | |
|---|---|---|
| 16908 | TKL1_repair oligo rv | GTTTTATTTCATTTTTTCACAAATAATATCATATCAAATCTGATGATCTACGATCAGAATTTTGTTTGCTAAAGAGGTAACTATGTAGAGTTGCTTGGGATGAGCTTCCTTCTCTGTTGT |
| 16911 | TAL1_repair oligo fw | AGGTAAAATTTAGTACGATAGTAAAATACTTCTCGAACTCGTCACATATACGTGTACATAGGAAGTATCTCGGAAATATTAATTTAGGCCATGTCCTTATGCACGTTTCTTTTGATACTT |
| 16912 | TAL1_repair oligo rv | AAGTATCAAAAGAAACGTGCATAAGGACATGGCCTAAATTAATATTTCCGAGATACTTCCTATGTACACGTATATGTGACGAGTTCGAGAAGTATTTTACTATCGTACTAAATTTTACCT |
| **Diagnostic primers to check gene deletion** | | |
| 1046 | TAL1Rv1 | AAGAACACCGAGCGGCTTTG |
| 1047 | TAL1Fw1 | CTGTACACTAGGAAGCCCTGTT |
| 2122 | BG26-DF | GCTGCAGTATTGTTCCTGAG |
| 2123 | BG26-DR | CCTGTTTGCCTTTCCTTACG |
| 4494 | RPE1 DG fw | TATCCAAGTCGAGCTGGGAAAG |
| 4495 | RPE1 DG rv | CCCATGAGTTAGGCACTTACG |
| 5598 | TKL1 DG fw | CGTTCCGTTCGCAATCTC |
| 5599 | TKL1 DG rv | GGTGTGATTCTCTCGAAGG |
| 8566 | FW_zwf1_outside | GGGTGGCGAATTCTTCAATG |
| 8567 | RV_zwf1_outside | ATTGCGTACGATGCGGTATG |
| 16893 | SOL3_dg fw | TGTCGCTGCTATCTACTGCG |
| 16894 | SOL3_dg rv | GATGAGGCACGCAAAGGTTG |
| 16901 | RKI1_dg fw | CATGGCCCAGATTGCTTGTG |
| 16902 | RKI1_dg rv | ATCCGGACAGGGTCTTGTTG |

### Supplementary Table 25 - Parts of the “basic design” of the anthocyanin pathway

* Size of the fragments does not include the SHR sequences.

| SHR Fw | Component | SHR Rv | Size* | Template | Primer Fw | Primer Rv |
|---|---|---|---|---|---|---|
| Chunk 16AB | *pRPS3-coAtCPR1-tIDH2* | F | 3559 | pUDC348 | 17908 | 17909 |
| F | *pSePDC1-AtPAL1-tLAT1* | DW | 3282 | pUDC349 | 14612 | 17910 |
| DW | *pSeGPM1-coRcTAL1-tCIT1* | DX | 2403 | pUDC350 | 17911 | 17912 |
| DX | *pSeTPI1-At4CL3-tSDH2* | DY | 2785 | pUDC351 | 17913 | 17914 |
| DY | *pTEF1-coAtCHS3-tMDH1* | AM | 2292 | pUDC352 | 17915 | 17916 |
| AM | *tSDH4-AtCHI1-pSkADH1* | AB | 1849 | pUDC353 | 17917 | 15587 |
| AB | *tADH3-coAtC4H-pSeFBA1* | DC | 2603 | pUDC354 | 15168 | 14460 |
| DC | *tSDH3-coAtF3H-pSkTDH3* | EA | 2186 | pUDC355 | 17918 | 17919 |
| EA | *tACO1-coGhDFR-pSePGK1* | EB | 2202 | pUDC356 | 17920 | 17921 |
| EB | *tFUM1-coAtANS-pSeENO2* | EC | 2180 | pUDC357 | 17922 | 17923 |
| EC | *tDIC1-coAt3GT-pSePYK1* | CJ | 2498 | pUDC358 | 17924 | 17925 |
| CJ | ARS106 | DL | 236 | CEN.PK113-7D genomic DNA | 13183 | 17926 |

### Supplementary Table 26 - List of primers for amplifying the fragments of the "basic design" of the anthocyanin pathway and diagnosing integration

| Primer number | Primer name | Sequence (5' to 3') |
|---|---|---|
| 13183 | ARS106 + Tag CJ Fw | TCGACCCATGTTTATCGCTAGCAGTCGCTTCAGCTAGATTCACAGAGTGGCCGTGACAATCAATGTTTTATCTACGTTGGAGTAA |
| 14460 | DC - pFBA1 - FW NEW | TGAGCCAGTGCATTCCATCGATGCAGATTCGCGTCCACGTAACGTATCGGAAGCATAGGCCTTTTCCCATGTTTCCAATG |
| 14612 | flank F - pPDC1 (Se) - FW | CATACGTTGAAACTACGGCAAAGGATTGGTCAGATCGCTTCATACAGGGAAAGTTCGGCAGATGAAGTGACGCGCGCCCGGA |
| 15168 | AB-tADH3-rv | TCAGCGTGTTGTAATGATGCGCCATGAATTAGAATGCGTGATGATGTGCAAAGTGCCGTCTCTCTTCGGCCCTTTTATCGTG |
| 15587 | AB-SkADH1p_fwd | GACGGCACTTTGCACATCATCACGCATTCTAATTCATGGCGCATCATTACAACACGCTGAACTCCCAAATAATCAAGGG |
| 17908 | Chunk16AB_pRPS3_fw | CGTAAGAACCGACTAACGTCCCCATATTGATGTTTACCGCCGAAGTGGGATCGGCCATCTTCTGCTACTTTCCATTATCTGGTC |
| 17909 | F_tIDH2_rv | TGCCGAACTTTCCCTGTATGAAGCGATCTGACCAATCCTTTGCCGTAGTTTCAACGTATGTCCACTGAGGGACATTTTGAG |
| 17910 | DW_tLAT1_rv | ACCCACAGTCGTAGATGCGTTGTCAGAATTTCCAGGTGTGGCTACATCTTCCGTACTATGAAACTTTATGCGTTATATCCTATATCCCACTCC |
| 17911 | DW_pSeGPM1_fw | CATAGTACGGAAGATGTAGCCACACCTGGAAATTCTGACAACGCATCTACGACTGTGGGTTAAACCTGATCTTTCACCTCAGTAAC |
| 17912 | DX_tCIT1_rv | ACAGGTCCTCAGGGCGATATTAATGGGATTGATGTCTGCCCTCCACTGTACTGCATGTAGCTTGACGTAGTATATCGACTACAGGC |
| 17913 | DX_pSeTPI1_fw | CTACATGCAGTACAGTGGAGGGCAGACATCAATCCCATTAATATCGCCCTGAGGACCTGTGGATGTCGTTGTTCTTGTTACAC |
| 17914 | DY_tSDH2_rv | ACACAGTCTAAGGAGAGTCTGCAATCCCTTATGAGTCAGTCAACGCATGAGGATGATGACAAGCCAAAAGGCCCTTCAA |
| 17915 | DY_pTEF1_fw | GTCATCATCCTCATGCGTTGACTGACTCATAAGGGATTGCAGACTCTCCTTAGACTGTGTCCTTGCCAACAGGGAGTTC |
| 17916 | AM_tMDH1_rv | CAGTGACATGCCGCTCAGTACTCGTATCTTACATGACGTGGGCATGGGTTCCGCTCATATGTTTATTCATCATTATCATCATCATCATC |
| 17917 | AM_tSDH4_fw | ATATGAGCGGAACCCATGCCCACGTCATGTAAGATACGAGTACTGAGCGGCATGTCACTGAATTGAAAATCCGCGAGTG |
| 17918 | DC(i)_tSDH3_fw | GCCTATGCTTCCGATACGTTACGTGGACGCGAATCTGCATCGATGGAATGCACTGGCTCAGCAGAAATTATCTTGATATCTGT |
| 17919 | EA_pSkTDH3_rv | AAGTAGGTCAGAGTAGCACTGGCTATGATTCGCAATGCTTGGTGAATTGAGAGCTATCCTAACGGCGAATTTTTACTAACC |
| 17920 | EA_tACO1_fw | AGGATAGCTCTCAATTCACCAAGCATTGCGAATCATAGCCAGTGCTACTCTGACCTACTTGCTCAGCCTTATTACTTAATTT |
| 17921 | EB_pSePGK1_rv | AAGCCTCGGACTCGAAGCATGAATCATGTATCATAGGCGGCTCAGCCTTAGCCAATATGAGCTTCAATTCAAGATACACAG |
| 17922 | EB_tFUM1_fw | TCATATTGGCTAAGGCTGAGCCGCCTATGATACATGATTCATGCTTCGAGTCCGAGGCTTTGCGGGTAATACTAGGTCC |
| 17923 | EC_pSeENO2_rv | TGAAATTATTCTGTGCCGGGCAGCGAAATGGCAGTATGCTCAGTGACGTGAGTGCCATCTAACGCCAAGAAGATGCCG |
| 17924 | EC_tDIC1_fw | AGATGGCACTCACGTCACTGAGCATACTGCCATTTCGCTGCCCGGCACAGAATAATTTCAGCCCAGCAAAATTCGAAA |

| | | |
|---|---|---|
| 17925 | CJ(i)_pSePYK1_rv | ATTGTCACGGCCACTCTGTGAATCTAGCTGAAGCGACTGCTAGCGATA AACATGGGTCGAAACGTGTAAATACCGGTTTTAGC |
| 17926 | DL_ARS106_rv | CGCAAATGTCCCATCGTATTTCAGAACCTTGTCACTCATGCGAGCAAG TGTGACAGCTATGCCGAAAAGGAGGTTTTCTTCTTATTC |
| **Diagnostic primers** | | |
| 18079 | Chunk 16AB fw | GCCTCGACATACTGTTCATC |
| 18080 | pRPS3 rv | CTTACATCAGCGCAGCAC |
| 18081 | ARS106 fw | GGGTCTGTCCAGCGAATAAG |
| 18082 | pFBA1 rv | TAACGTGGGCGAAGAAGAAG |
| 18083 | *CoAtF3H* fw | CGTCGATACCAGCCAAAGAG |
| 18084 | tACO1 rv | GTTCGGCTGGAGAAGTCAAG |
| 18085 | pSkADH1 fw | GCGGGTATGGTGAGGTAAC |
| 18086 | coAtC4H rv | GCTAACGGTAACGACTTCAG |
| 18087 | tCIT1 fw | AGACCCTCCAGCCTAAATCC |
| 18088 | pSeTPI1 rv | AACTGGATGCCGAAACAGAG |
| 18089 | tLAT11 fw | CAAACGGTGCGTCAACATC |
| 18090 | coRcTAL1 rv | TCTAGCTTCGGCCCAAGAC |

Supplementary Table 27 - List of primers for amplifying the fragments of the "elaborate design" of the anthocyanin pathway with several copies of the chalcone synthase and diagnostic PCR.

| Primer number | Primer name | Sequence (5' to 3') |
|---|---|---|
| **14823** | YPRCtau3_pTEF1_fw | ACAGTTTTGACAACTGGTTACTTCCCTAAGACTGTTTATATTAGGATT GTCAAGACACTCCCTTGCCAACAGGGAGTTC |
| **18003** | can1_pTEF1_fw | GATGAGAAAAGTAAAGAATTGTATCCATTGCGCTCTTTCCCGACGAGA GTAAATGGCGAGCCTTGCCAACAGGGAGTTC |
| **18004** | can1_tMDH1_rv | GGTGTATGACTTATGAGGGTGAGAATGCGAAATGGCGTGGGAATGTGA TTAAAGGTAATAGTTTATTCATCATTATCATCATCATCATC |
| **18005** | X2_pTEF1_fw | TCACAGAGGGATCCCGTTACCCATCTATGCTGAAGATTTATCATACTA TTCCTCCGCTCGCCTTGCCAACAGGGAGTTC |
| **18006** | X2_tMDH1_rv | GTCATAACTCAATTTGCCTATTTCTTACGGCTTCTCATAAAACGTCCC ACACTATTCAGGGTTTATTCATCATTATCATCATCATCATC |
| **18007** | YPRCtau3_tMDH1_rv | ATAATTATAATATCCTGGACACTTTACTTATCTAGCGTATGTTATTAC TCGATAAGTGCTGTTTATTCATCATTATCATCATCATCATC |
| **18008** | SPR3_pTEF1_fw | AGAAATAAATAAATAAATAATAAAAACCTAAAATTCCTTTTGCGTCAT TGAATTTTTATTCCTTGCCAACAGGGAGTTC |
| **18009** | SPR3_tMDH1_rv | TTTATTATGTAGAGCAAAGCTTGCGCGAAATTATTGGCTTTTTTTTTT TTTTAATTAATAGTTTATTCATCATTATCATCATCATCATC |
| **18225** | CAN1_pAgTEF1_fw | GATGAGAAAAGTAAAGAATTGTATCCATTGCGCTCTTTCCCGACGAGA GTAAATGGCGAGGACATGGAGGCCCAGAATACC |
| **18234** | chunk7BC_pTEF1_fw | CCGCTGCCACCACGCCAAGCGCGGTTAAACCGACGGTATCGGCGGTGG AATGAGTACCCACCTTGCCAACAGGGAGTTC |
| **18235** | chunk7BC_tMHD1_rv | TCACGGCTGCCTGGGCTGTGCGGAAAATGGTTTCTGGGATCGCGGTTC GTTCTACAGCCGGCGTTTATTCATCATTATCATCATCATC |
| **18236** | chunk15CD_pTEF1_fw | AGATATACCAGACAAATCAATGTCAGAATCCAAATCAGATATTCCCGG CGTATTTATCCGCCTTGCCAACAGGGAGTTC |
| **18237** | chunk15CD_tMHD1_rv | ATATTGAAAACATTAATGCGTGTGATGATGTTTTTTCTGAGTATTGTT TTGATGATGAAAGCGTTTATTCATCATTATCATCATCATC |
| **18238** | tADH1/SHR-N_pTEF1_fw | CAATGAGTTGATGAATCTCGGTGTGTATTTTATGTCCTCAGAGGACAA GATCAGCAGCCACCTTGCCAACAGGGAGTTC |
| **18239** | pPYK1/SHR-N_tMHD1_rv | TATTTTTCTTAGTTCTGAAGTTTGACCACGTTTTCTAGGCTTTGATGC AAGGTCCACATAGCGTTTATTCATCATTATCATCATCATC |
| **18240** | chunk9CD_pTEF1_fw | AACCAGCGACGGATATTGCTGTGCCAGTTGTCGGGCAAGCGTAATGCC GTCGATATCCGGCCTTGCCAACAGGGAGTTC |
| **18241** | chunk9CD_tMHD1_rv | CTGTTGGCAATGCCGCGCAGGCTTTAGAGACACTGCAAAATAGCGAAC CGTTTGCTGCCGGCGTTTATTCATCATTATCATCATCATC |
| **Diagnostic primers** | | |
| **92** | YGR059w CTRL RV | ATGATGTCGCGCATTTGATGCCTTAAATAC |
| **2496** | FW-conf-upstrm | CGGGAGCAAGATTGTTGTG |
| **2497** | RV-conf-dwnstrm | GGTTGCGAACAGAGTAAACC |
| **2820** | Probe AmdS fw | AGCTTCTGCTGCTGACTTGG |
| **2908** | D_FW PDH construct ctrl | GGATTGGGTGTGATGTAAGGATTCGC |
| **3853** | Fus GF cassette fw | GCTGCATCCTTCCCATGCAAAGTG |
| **6028** | AmdS ORF rv (DT37) | TGTCAGCAGCCAATTCTTC |

| | | |
|---|---|---|
| **7376** | FW_x-2_outside | GGTCTAGGCCTGCATAATCG |
| **7377** | RV_X-2_outside | TGCGGCATCATGTCTACTTG |
| **13261** | YPRCtau3 dg FW | AATACGAGGCGAATGTCTAGG |
| **13262** | YPRCtau3 dg RV | GCCTCCCCTAGCTGAACAAC |
| **13336** | TEF1p seq | GCTCATTAGAAAGAAAGCATAGCAATC |
| **17360** | CHS_Citrus_reticulat_fw | CGGTTTCGGTCCAGGTTTGAC |
| **17950** | AtCHI1 dg - FW | CCCGTTCTTCCGTGAAATAG |
| **18156** | AM_dg_fw | AACATATGAGCGGAAGAC |
| **18157** | DZ_dg_rv | GAATCACAGTCGCCCTTG |
| **18158** | DZ_dg_fw | CAAGGGCGACTGTGATTC |
| **18159** | ED_dg_rv | ATATCGGCCATCGTGCCTTG |
| **18160** | ED_dg_fw | ATGGCCGATATTGCGTTGAG |
| **18161** | EE_dg_rv | CGATATTGCCAGTCAGGTCAG |
| **18162** | EE_dg_fw | CCTGACTGGCAATATCGTTAC |
| **18163** | EF_dg_rv | ATGGTCGTGGACTCTATCTG |
| **18164** | EF_dg_fw | GATAGAGTCCACGACCATCC |
| **18165** | tSDH4_dg_rv | CGCCGGTATATTCCTTTGC |
| **18378** | diag_7BC fw | GAACAAACCGCGCATTCC |
| **18379** | diag_15CD fw | GGCAGAGCTTCAGAGTCTATC |
| **18380** | diag_15CD rv | GACGTGTCGGTATCTAAAGC |
| **18381** | diag 9CD fw | TCCCGCGGAATAATGAACTG |
| **18382** | diag 9CD rv | CGCTAACCCAGCGAATTAC |

### Supplementary Table 28 List of primers for amplifying the correct *CoAtANS* transcriptional unit and the diagnostic primers used to confirm correct integration.

| Primer number | Primer name | Sequence (5' to 3') |
|---|---|---|
| **18740** | SHR AL_tFUM1_fw | CTGACTCATTGCCATTGCATGTGCAGACTCAACACCATGACATCTTAA TACAGAGGACGCTGCGGGTAATACTAGGTCC |
| **18741** | SHR DS_pSeENO2_rv | ATCAAGACTGAGGAGTACGTCAGGTTGCAGAGGATCACTTGTAATG AATGTGTGCTCGCTAACGCCAAGAAGATGCCG |
| **3537** | ilv5 flanking rv | AATCGTAGCTGTCCCGATGAGG |
| **17730** | 17730_RPE1_dg_rv | GTGGTTTGGGCAAGGAGACAATC |
| **17973** | *coAtANS* diag rv | GGTTCCAAACCCAAACCAACAG |
| **17974** | *coAtANS* diag fw | GGCCAAAGACTCCATCTGAC |

### Supplementary Methods 1: Strains, growth medium and maintenance.

For liquid cultures yeast was cultivated in 50-/100-/500 mL shakeflasks containing, respectively 10-/20-/100 mL media in an Innova 44 Incubator shaker (New Brunswick Scientific, Edisan, NJ, USA) at 30 °C and 200 rpm. Cultures on solid media were incubated at 30°C until single colonies were visible. For non-selective growth, yeast strains were cultivated on Yeast extract Peptone Dextrose (YPD) medium containing: 10 g $L^{-1}$ Bacto yeast extract, 20 g $L^{-1}$ Bacto peptone and 20 g $L^{-1}$ glucose. For selective growth to maintain plasmids or NeoChrs, Synthetic Medium (SM) was used, consisting of: 3 g $L^{-1}$ $KH_2PO_4$, 0.5 g $L^{-1}$ $MgSO_4 \cdot 7H_2O$, 5 g $L^{-1}$ $(NH_4)_2SO_4$ and 1 mL $L^{-1}$ of a trace element solution [15]. Alternatively, synthetic medium with urea as sole nitrogen source was used, consisting of 3 g $L^{-1}$ $KH_2PO_4$, 0.5 g $L^{-1}$ $MgSO_4 \cdot 7H_2O$, 5 g $L^{-1}$ $K_2SO_4$, 2.3 g $L^{-1}$ urea and 1 mL $L^{-1}$ of a trace element solution [16]. Media were set to pH 6 by 1M KOH addition and for solid media, 20 g $L^{-1}$ Bacto agar was added. Autoclaving was performed for 20 min at 110°C and 120°C for YPD and SM medium, respectively. Thereafter SM medium was supplemented with 1 mL $L^{-1}$ of a filter sterilized vitamin solution and 20 g $L^{-1}$ of glucose separately autoclaved for 20 min at 110°C. For auxotrophic strains, SM was supplemented with 125 mg $L^{-1}$ histidine and/or 150 mg $L^{-1}$ uracil. Disruption of the *URA3* marker was verified by growth on SMD with 150 mg $L^{-1}$ uracil and 1 g $L^{-1}$ 5-FluoroOrotic Acid (SMD–5-FOA). For the selection based on the markers *hphNT1, KanMX* and *amdS,* SM medium without nitrogen source was prepared by replacing $(NH_4)_2SO_4$ with 6.6 g $L^{-1}$ $K_2SO_4$. For *hphNT1* and *KanMX,* 2.3 g $L^{-1}$ urea was used as nitrogen source and 200 mg $L^{-1}$ hygromycin (Hyg) and 200 mg $L^{-1}$ G418 were added to the medium, respectively. For *amdS,* 1.8 g $L^{-1}$ filter sterilized acetamide was employed as nitrogen source.

All *E. coli* strains were cultivated in Lysogeny Broth (LB) medium containing: 10 g $L^{-1}$ tryptone, 5.0 g $L^{-1}$ yeast extract and 5 g $L^{-1}$ NaCl. For plasmid selection 100 mg $mL^{-1}$ ampicillin (ampR), 50 mg $mL^{-1}$ kanamycin (kanR), or 25 mg $mL^{-1}$ chloramphenicol (camR), was supplemented to the medium. Liquid cultivation was performed in 5 mL medium in a 15 ml Greiner Tubes at 37°C and 200 rpm in an Innova 4000 shaker (New Brunswick Scientific). Cultures on solid media were incubated at 37°C until single colonies were visible.

*S. cerevisiae* and *E.coli* strains were stored at -80˚C in 1 mL vials containing cultures mixed with glycerol (30% v/v).

### Supplementary Methods 2: Molecular biology techniques

Genomic DNA from *E.coli* (migula) Castellani and Chalmers (ATCC 47076) or *S. cerevisiae* used for strain construction purposes was isolated using the QIAGEN Blood & Cell Culture Kit with 100/G Genomic-tips (Qiagen, Hilden, Germany) or alternatively for yeast with the YeaStar genomic DNA kit (Zymo Research, Irvine, CA). *E.coli* DNA from a mixed population of *E.coli* XL1-Blue and *E.coli* BL21 used for construction of the test NeoChrs was isolated as described by Postma *et al.* [8]. Plasmids were isolated from *E.coli* using the GenElute Plasmid Miniprep Kit (Sigma-Aldrich, St. Louis, MO) or the GeneJET Plasmid Miniprep Kit (Thermo Fisher Scientific, Waltham, MA), according to the manufacturer's instructions.

All PCRs for strain construction purposes were performed with Phusion High-Fidelity DNA Polymerase (Thermo Fisher Scientific) using either desalted or PAGE purified (in case of ORFs) primers (Sigma-Aldrich). PCR products were verified by separation on 1% (w/v) or 2% (w/v) agarose (TopVision Agarose, Thermo Fisher Scientific) gels in 1x Tris-acetate-EDTA (TAE) buffer (Thermo Fisher Scientific) or 1x Tris-Borate-EDTA (TBE) (Thermo Fisher Scientific) buffer. For size determination GeneRuler DNA Ladder mix (Sigma-Aldrich) or GeneRuler DNA Ladder 50bp (Sigma-Aldrich) were used. For DNA staining 10 µL $L^{-1}$ SERVA (SERVA Electrophoresis GmbH, Heidelberg, Germany) was added to the agarose gel solution. DNA was purified using either the Zymoclean Gel DNA Recovery kit (Zymo Research), the GenElute PCR Clean-Up kit (Sigma-Aldrich), the GeneJET PCR Purification Kit (Thermo Fisher Scientific) or using AMPure XP beads (Beckman Coulter, Brea, CA) according to the suppliers' protocols. Purity of DNA was checked using the NanoDrop 2000 spectrophotometer (Thermo Fisher Scientific) and the concentration was measured either by the NanoDrop 2000 (Thermo Fisher Scientific) or by the Qubit dsDNA BR Assay kit (Thermo Fisher Scientific) using the Qubit 2.0 Fluorometer (Invitrogen, Carlsbad, CA). Gibson assembly used to construct gRNA plasmids and some expression plasmids was performed with the NEBuilder® HiFi DNA Assembly Master Mix (New England Biolabs, Ipswich, MA) in a final volume of 5 µL according to the supplier's instruction.

Chemical *E.coli* XL1-Blue transformation was performed as described by Inoue *et al.* [17] and correct assembly of plasmids was verified by diagnostic PCR or restriction analysis. *S. cerevisiae* was transformed using the lithium acetate/polyethylene glycol method [18]. For diagnostic PCR, DNA was isolated by resuspending some culture in 0.2 M NaOH or by using the method described by Looke *et al.* [19]. All diagnostic PCRs were performed using DreamTaq PCR Master Mix (Thermo Fisher Scientific) according to the manufacturer's instruction. For yeast, single colony isolates were obtained by three consecutive re-streaks on solid selective medium.

### Supplementary Methods 3: Detailed construction of the host strain IMX2770

Before assembly of coding NeoChrs a suitable starting strain was engineered by several rounds of CRISPR/Cas9 gene deletions as described by Mans *et al.* [9] . As parental strain, the SwYG strain IMX589 from Kuijpers *et al.* [6] was used. In this strain the minor paralogs of glycolysis are deleted and the major paralogs are centralized at the *sga1* locus on chromosome IX.  From this strain, the *amdSYM* marker located between the major paralogs of glycolysis was deleted using *in vivo* assembly of a pMEL10 gRNA plasmid backbone (amplified with primer 6005, Supplementary Table 16) and a gRNA insert made from annealing primers (11588 & 11589, Suppl. Table S15). The DSB was repaired with a 120 bp repair fragment homologous to the flanking SHRs K and L, made by annealing of complementary primers (11590 & 11591, Supplementary Table 16), resulting in strain IMX1433 and IMX1769 before and after plasmid recycling, respectively.

Subsequently the minor paralogs of the pentose phosphate pathway, *GND2, NQM1, SOL4* and *TKL2*, were deleted by transformation of two gRNA plasmids (pUDR286 & pUDR590, Supplementary Table 8A) and 120 bp repair fragments (Supplementary Table 17) homologous to the 60 bp upstream and downstream of the ORF. The strains were stocked before and after discarding the gRNA plasmids, resulting in respectively IMX2154 and IMX2204

Next as much as possible of the promoter, gene and terminator of the *ura3* and *his3* as well as of the functional *SpHIS5* gene were removed using gRNA plasmids pUDR426 and pUDR546 and repair fragments (Supplementary Tables 8A and 18) obtaining strain IMX2234 after plasmid recycling.

Finally, in the last round of deletion, the *ARO10* gene was removed with gRNA plasmid pUDR406 and a 120 bp repair fragment (Supplementary Tables 8A and 19). Again, the plasmid was removed and the strain was stocked as IMX2270.

### Supplementary Methods 4: MinION long-read sequencing

Average DNA size and integrity were verified with the TapeStation 2200 (Agilent Technologies, Santa Clara, CA). Before sequencing, flow cell quality was assessed by running the MinKNOW platform QC. All samples were sequenced in-house on a MinION (Oxford Nanopore). Samples NeoChr10.10, NeoChr10.13 (IMF22), Neochr10.47, NeoChr10.54, NeoChr10.16, NeoChr10.62, NeoChr10.67, NeoChr10.69, NeoChr11.19, NeoChr11.22, NeoChr25.25, NeoChr25.47, NeoChr25.53, NeoChr25.56 (IMF27), NeoChr26.2, NeoChr26.4 (IMF29), NeoChr26.6, Neochr26.9 and NeoChr26.1 were sequenced on a FLO-MIN106 flowcell with sequencing kit SQK-LSK108.

Samples NeoChr12 (IMF23), NeoChr30 (IMF41), NeoChr31 (IMF42), NeoChr33 (IMF47) and NeoChr34 (IMF48) were sequenced on a FLO-MIN111 with sequencing kit SQK-LSK109. Basecalling was performed for samples with NeoChr10 and Neochr11 by using Albacore (version 2.3.1, Oxford Nanopore). Demultiplexing of the fastq files of the NeoChr10 and NeoChr11 samples was performed with Porechop (https://github.com/rrwick/Porechop). Basecalling and demultiplexing was performed with Guppy (Oxford Nanopore) for samples with NeoChr25 and NeoChr26 with version 3.1.5, samples IMF41, IMF42, IMF47 with version 4.4.2 and IMF48 with version 4.5.4. All resulting fastq files were filtered on length (> 1kb) followed by *de novo* assembly by Canu version 2.0 [20].

### Supplementary Methods 5: Analysis of aromatics

#### A. HPLC analysis of aromatic compounds up until naringenin

For extracellular aromatic compounds, a sample containing broth was mixed 1:1 with 96% ethanol, vortexed thoroughly, spun down for 5 minutes at 14800 rpm and the supernatant was used for further analysis. The aromatic compounds up until naringenin (2-phenylethanol (2PE), *p*-hydroxyphenylethanol (*p*OH2PE), phenylacetic acid (PAA), *p*-hydroxyphenylacetic acid (*p*OHPAA), phenylpyruvic acid (PPY), coumaric acid (COUM), cinnamic acid (CIN), phloretic acid (PHLOR) and naringenin (NAR) were measured using an Agilent Zorbax Eclipse plus C18 column (4.6 x 100mm, 3.5 μm) (Agilent). As mobile phase, 0.020 M $KH_2PO_4$ set at pH 2.0 containing 1% acetonitrile was used at a flow rate of 0.8 mL $min^{-1}$ at an operating temperature of 40°C. The amount of acetonitrile was gradually increased to 10% within 6 minutes, then to 40% after 23 minutes, followed by a decrease in amount to 1% after 30 minutes. The compounds were detected using a diode array and a multiple wavelength detector (Agilent G1315C) at different wavelengths: 200 nm for PAA, 210 nm for PPY, 214 nm for 2PE, *p*OH2PE, *p*OHPAA and PHLOR, 270 nm for CIN and finally 280 nm for NAR and COUM.

The extracellular concentrations in the supernatant of the aromatic compounds kaempferol (KEA), dihydrokaempferol (DHK), kaempferol 3-O-glucoside (K3G), pelargonidin (PEL) and pelargonidin 3-O-glucoside (P3G) were detected using LS-MS/MS, as described in the next section. Additionally, since P3G has never been measured extracellular before, the intracellular concentrations of P3G, and its precursors kaempferol, dihydrokaempferol, K3G and pelargonidin were also measured. A certain amount of cell culture was spun down for 5 minutes at 5000 rpm, washed once with $dH_2O$, resuspended in 0.5-1 ml methanol (0.75% HCL) and the samples were stored overnight at -80°C. Next, the samples were lyophilized for 24 h using a Mini Lyotrap freeze-dryer (LTE Scientific TLD, UK) operated at -80 °C, connected to a Pirani 501 manometer (Edwards Vacuum, UK) using a RV8 pump (Edwards Vacuum, UK). Finally, the pellet was resuspended in 1 mL methanol (2.0% HCL) and stored overnight at -80 °C.

#### B. Mass spectrometric analysis of anthocyanin pathway compounds

Identification and quantification of compounds from the anthocyanin pathway downstream of naringenin was performed using an ACQUITY UPLC chromatography system (Waters, UK) coupled online to a high-resolution Orbitrap mass spectrometer (Q-Exactive Focus, Thermo Fisher Scientific, Germany). For chromatographic separation, a reverse phase separation column (ACQUITY UPLC BEH C18, 1.0 mm × 100 mm, 3 μm particle size, part No 186002346, Waters UK) was operated at room temperature using $H_2O$ plus 0.1% formic as mobile phase A, and acetonitrile plus 0.1% formic acid as mobile phase B. A gradient was maintained at 50 µL/min at 7.5% B over 5 minutes. Solvent B was then increased to 80% over 4 minutes, and kept constant for additional 3 minutes before equilibrating back to the starting conditions. The metabolite extracts were taken from -80°C immediately before injection, brought to room temperature, vortexed and 15 µL crude extract were mixed with 85 µL 1 mM HCl. The mixture was carefully vortexed and centrifuged using a bench top centrifuge for 1 minute to remove insoluble materials. 5 µL were subsequently injected onto the UPLC reverse phase separation system. The mass spectrometer was operated alternating in full scan and PRM mode. Full scan was acquired from 250–700 m/z in ESI positive mode (+ 3.25 kV), at a resolution of 70 K. Parallel reaction monitoring was performed for the precursor masses for dihydrokaempferol (DHK, Cas No. 104486-98-8) 289.07 m/z $[M+H]^+$ using a NCE of 26, kaempferol (KEA, Cas No. 520-18-3) 287.05 m/z $[M+H]^+$ using a NCE of 30, kaempferol 3-O-glucoside (K3G, Cas No. 480-10-4) 449.10 m/z $[M+H]^+$ using a NCE of 24, pelargonidin (PEL, Cas No. 134-04-3) 271.06 m/z $[M]^+$ using a NCE of 30 and pelargonidin 3-O-glucoside (Cas No. 18466-51-8) m/z 433.10 $[M]^+$ using a NCE of 24. Fragment ions were measured at fixed first mass of 75 m/z, a resolution of 35K, a max IT of 100 ms and an AGC target of 1e5, by acquiring 2 microscans. Raw data were analyzed using XCalibur 4.1 (Thermo) where retention and unique fragments for each individual compound were compared to commercial standards. For quantification, peak intensities of identified compounds from the samples were summed using Matlab 2020b, and compared against an external calibration curve established using commercial standards. The standards were purchased from Sigma Aldrich (dihydrokaempferol Cat No. 91216, kaempferol Cat No. 60010, kaempferol 3-O-glucoside Cat No. PHL89237, pelargonidin chloride Cat No. PHL80084, pelargonidin 3-O-glucoside chloride Cat No. PHL89753). The mass spectrometer was calibrated using the Pierce™ LTQ ESI positive ion calibration solution (Thermo Fisher Scientific, Germany).
